# Genome-wide analysis of polymorphisms identified domestication-associated polymorphism desert carrying important rice grain size/weight QTL

**DOI:** 10.1101/725242

**Authors:** Angad Kumar, Anurag Daware, Arvind Kumar, Vinay Kumar, Gopala Krishnan S, Subhasish Mondal, Bhaskar Chandra Patra, Ashok. K. Singh, Akhilesh K. Tyagi, Swarup K. Parida, Jitendra K. Thakur

**Affiliations:** National Institute of Plant Genome Research, ArunaAsaf Ali Marg, New Delhi 110067, India; Division of Genetics, ICAR-Indian Agricultural Research Institute, Pusa, New Delhi 110012, India; Crop Improvement Division, ICAR-National Rice Research Institute, Cuttack 753006, India; Department of Plant Molecular Biology, University of Delhi South Campus, New Delhi 110021, India

**Keywords:** rice, genome sequence, grain size/weight, polymorphism desert, domestication, selection sweep, association analysis

## Abstract

Rice grain size and weight are major determinant of grain quality and yield and so have been under rigorous selection since domestication. However, genetic basis for contrasting grain size/weight trait among indian germplasm, and their association with domestication shaped evolutionary region is not studied before. To identify genetic basis of grain size/weight two long (LGG) and two short grain genotypes (SGG) were resequenced. LGG (LGR and Pusa Basmati 1121) differentiated from SGG (Sonasal and Bindli) by 504439 SNPs and 78166 InDels. The *LRK* gene cluster was significantly affected and a truncation mutation in the LRK8 kinase domain was uniquely associated with LGG. Phylogeny with 3000 diverse rice accessions revealed four sequenced genotypes belonged to *japonica* group and were at the edge of clades indicating source of genetic diversity available in Asian rice population. Five SNPs significantly were associated with grain size/weight and top three SNPs were validated in RIL mapping populations, suggesting this study as a valuable resource for high-throughput genotyping. A contiguous ∼6 Mb polymorphism desert region carrying a major grain weight QTL was identified on chromosome 5 in four sequenced genotypes. Further, among 3000, this region was identified as evolutionary important site with significant positive selection, elevated LD, and multiple selection sweeps, stabilising many domestication-related traits including grain size/weight. The *aus* group genotype retained more allelic variations in the desert region than *japonica* and *indica*, and likely to be one of the differentiation point for *aus* group. We suggest this desert region as an important evolutionary node that can be selected in breeding programs for improvement of grain yield and quality. All data and analysis can be accessed from RiceSzWtBase database.

**Significance statement:** Being an important trait, rice grain size/weight has been under rigorous selection since domestication. However, a link between this trait and domestication is not so directly established. In addition to characterization of novel grain size/weight-associated SNPs, in this study, ∼ 6 Mb polymorphism desert region harboring major grain weight QTL was identified on chromosome 5, which turned out to be an evolutionary important site with multiple selection sweeps and introgression events, significantly correlated with domestication-related traits.

## Introduction

More than half of the world population use rice (*Oryza sativa* L.) as the main staple food. In Southeast Asia, it provides 75% of the calorie intake of the whole population (Khush, 1997; Fitzgerald et al., 2009). As world population is increasing and productive cultivable land areas is decreasing due to a variety of factors including climate change, there is an urgency to increase the rice production. Therefore, increasing the potential of rice yield has been the prime objective in rice breeding, and genomics research enabling dissection of potential genes and variations which could be utilized in molecular breeding. Rice grain yield is mainly determined by panicle number per plant, filled grain number per panicle and grain weight traits (Huang et al., 2011). Among these, the most important trait is grain weight which is measured as the 1000- grain weight (Jianlong et al., 2002). All the parameters of grain shape like length, width and thickness are positively associated with grain weight and attributes to the major yield determinant in cultivated rice (Tan et al., 2000; Jianlong et al., 2002). In addition to yield, grain size especially the seed length-to-width ratio is considered as desired trait that influences the market value of rice. Moreover, indian subcontinent flaunts the rich germplasm diversity and status of second largest producer of rice. Therefore, exploring the genetic potential and molecular basis in the determination of grain size/weight and yield is of great significance for rice breeding programs.

Map-based gold standard genome of rice (Nipponbare) has been released by the International Rice Genome Sequencing Project (2005). Also, the complete physical map of chromosomes of ‘Kasalath’ and an updated version of the whole genome sequence of ‘93-11’ rice variety have been published (Gao et al., 2013; Kanamori et al., 2013). All these high-quality genome sequences serve as references for resequencing of genomes of diverse rice accessions with contrasting agronomic traits. The main DNA polymorphisms, single nucleotide polymorphisms (SNPs) and insertion-and-deletion polymorphisms (InDels), have been routinely used as DNA markers (Subbaiyan et al., 2012). Though, some of the previous studies have emphasised on genome-wide DNA polymorphism discovery and association analysis to reveal high genetic variability (McNally et al., 2009; Jain et al., 2014; Alexandrov et al., 2015; Duitama et al., 2015; Wang et al., 2018), their effective deployment in marker-assisted breeding for yield improvement in different genetic background of rice cultivars (*indica*, *japonica*, *aus* and aromatic) is very limited. This may be because the available SNP resources are restricted to cultivars that are adapted to specific agro-climatic regions. Thus, discovery of sequence variations from rice cultivars with different genetic backgrounds is necessary and a comprehensive analysis of genetic-based grain size/weight diversity is required.

The interaction between selection and recombination, or linked selection, can have a profound impact on the levels of genetic variation across the genome. Genomic region of severely reduced polymorphism density are considered polymorphism desert and one such desert region has been found on chromosome 5 in previous studies (Krishnan S et al., 2014; Rathinasabapathi et al., 2015; Singhabahu et al., 2017; Trinh et al., 2017). However, evolutionary forces shaping the patterns of such polymorphism desert across genomes remain elusive.

In this study, whole genome of four indian rice genotypes, two large grain [LGR (IC 301206) and Pusa Basmati 1121] and two short grain (Sonasal and Bindli) were re-sequenced. Analysis of sequences led to the identification of large number of DNA polymorphisms, including SNPs and InDels. Different genes and processes significantly enriched in large-effect nucleotide variations that might be associated with the regulation of grain size/weight trait were characterized. These can be utilized for improving rice yields through molecular-marker assisted pyramiding of relevant SNPs. GWAS (Genome wide association study) performed with 1142 rice accessions including the four genotypes identified five SNPs significantly associated with grain size/weight. Three SNPs, including a novel one were validated in a couple of Recombinant Inbred Lines (RILs) underlying the importance of this study. Analysis of SNPs among four indian genotypes identified one long ∼6Mb low diversity region (polymorphism desert) over the centromeric region of chromosome 5. Extended analysis among 3004 diverse rice accessions revealed presence of this desert region in almost all accessions except most of *aus* rice. We found this region as the major site of selection sweep, elevated Tajima’s D (positive selection), stable linkage disequilibrium and region with high genetic differentiation (*F*ST) among *japonica* and *indica* from *aus* genotypes. Interestingly, this desert region showed potential association with grain size/weight and other domestication-related traits that might have played important role during rice adaptive evolution. Mapping population created by PB 1121 (LGG) x Bindli (SGG) revealed presence of a major QTL of grain size/weight in the polymorphism desert region of chromosome 5. These all evidences shows a correlation between grain size/weight trait and domestication linked through polymorphism desert region, which we suggest to be vertically selected from the standing variation among the wild progenitors of *japonica* and *indica*. Substantial introgression/gene flow from modern cultivars to traditional *aus* varieties could possibly explain the intermittent presence of desert in few *aus* genotypes. Overall, this study reveals that the long low diversity region on chromosome 5 is an evolutionary important site for domestication-associated multiple selection sweeps, but is contrastingly crafted among different varietal groups. More importantly, sequence variations within this site found to be associated with the regulation of rice seed size/weight trait that can be utilized for various genomics-assisted breeding applications in rice crop improvement. All the data and analyses are available on RiceSzWtBase database.

## Results

### Whole genome resequencing of low and high grain weight rice genotypes

In the present study, four rice genotypes from different regions of India with significant differences in grain length, width and 1000-grain weight (dehusked) namely, LGR and PB 1121 possessing longer grains (Long grain genotypes/LGG) with comparatively higher seed weight, and Sonasal and Bindli possessing shorter grains (Short grain genotypes/SGG) with lower seed weight, were selected (Figure 1A). Width-wise, Sonasal was found to be thinnest and LGR was the widest (Figure 1B). Thus, LGR seeds are heavier than the other three selected genotypes and Sonasal has the least 1000-grain weight (Figure 1C; Supplemental Table 1). The genome sequencing of four genotypes using Illumina platform (Hiseq 2000) generated over 180 million high-quality 90-bp long paired-end reads from each genotype. More than 92% of clean reads uniformly covering 87-97% of the total genome were successfully mapped to the Nipponbare reference genome. Uniquely mapped reads provided average sequencing depth from 44X to 65X. Summary of reference based alignment of sequencing data of four genotypes is provided in Supplemental Table 2.

**Figure 1:**
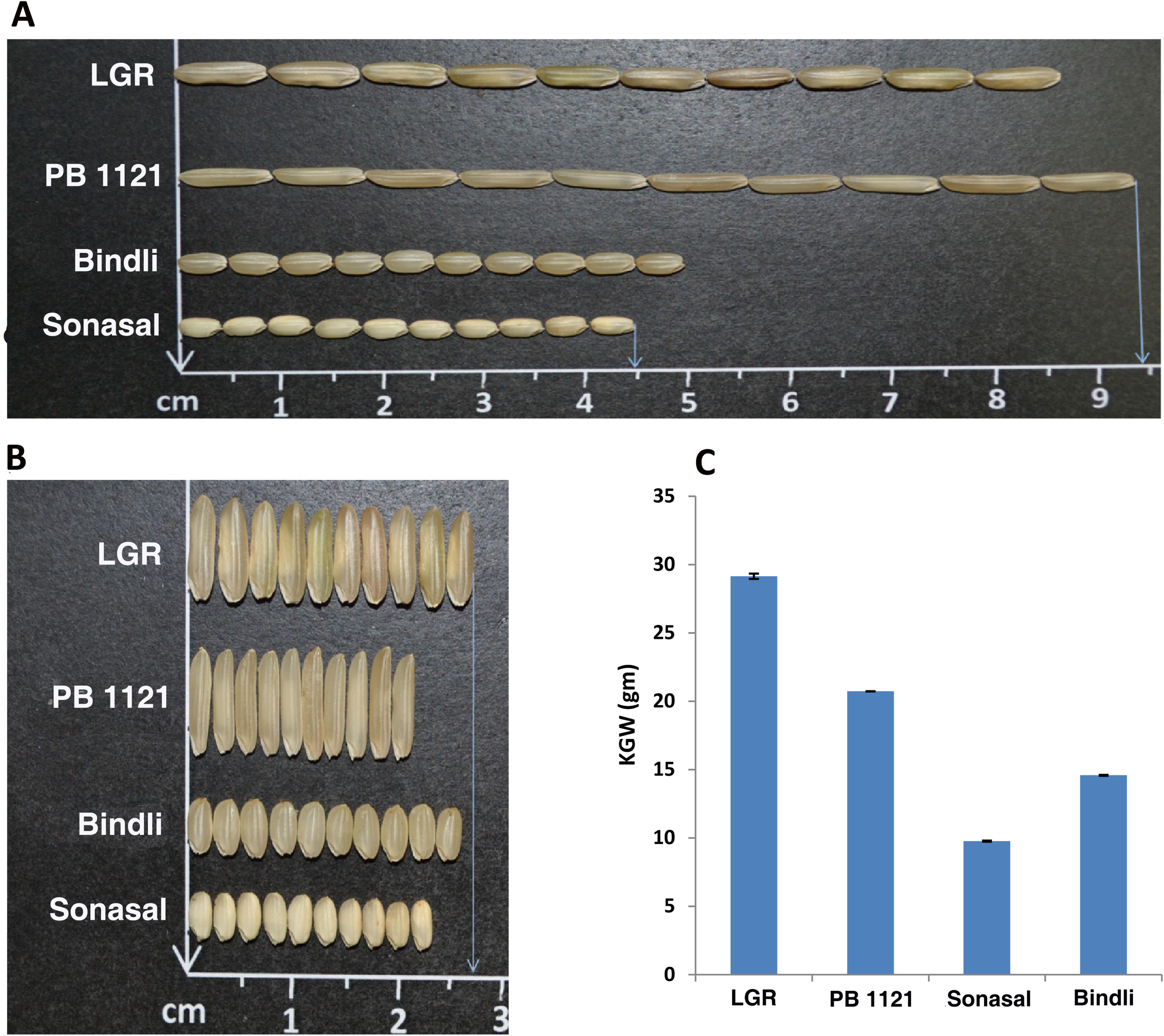
Seed parameter comparative analysis of four rice genotypes. (**A**) Seed length, (**B**) Seed width and (**C**) thousand grain weight (dehusked).

### Genome wide discovery of sequence polymorphism (SNPs and InDels)

A total of 4756763 SNPs and 588088 InDels were identified by comparing sequences from all the four rice genotypes with reference to Nipponbare genome. To minimize the detection of false-positives, sequence variants were quality filtered resulting in identification of 3564117 SNPs and 503654 InDels with overall density of 954.8 and 134.9 per 100 kb, respectively (Supplemental Table 3). Density of insertions and deletions was 65.1 and 69.8per 100 kb, respectively. The read depth of these two polymorphisms varied from 10 to more than 11000 (Supplemental Figure 1). The summary of identified SNP and InDel types is discussed in Supplemental Text, Supplemental Figure 2 & 3.

Next, SNPs and InDels were identified individually in four genotypes. SNPs ranged from 1343234 in Bindli to as high as 2417702 in LGR (Supplemental Figure 4A). However, the homozygous SNPs in LGR, PB 1121, Bindli and Sonasal were 2049708, 1608856, 1093340 and 995545, respectively (Supplemental Table 4). A total of 98 of these SNPs were validated by individually amplifying the randomly selected variant spanning genomic regions and sequencing by Sanger method. Primers used for sequence validation are given in Supplemental Table 5. There was no uniformity in the distribution of SNPs and InDels over the rice chromosomes (Supplemental Figure 4B-C). In all the genotypes, chromosome 1, the largest chromosome in rice, was found to have highest number of SNPs and InDels, whereas chromosome 9 being the smallest accounted for the least variations. The average density of DNA polymorphism was much higher in LGR and PB 1121 in comparison to Sonasal and Bindli. Though several SNPs were shared by all the four genotypes, many SNPs were unique to individual genotypes (Supplemental Figure 4D). Interestingly, LGR accounted for 882808 unique SNPs which was significantly more other than three genotypes, indicating the potential of LGR as a source of novel allelic variations. The average density of homozygous SNPs in LGR, PB 1121, Bindli and Sonasal was 549.1, 431.0, 292.9 and 266.7 per 100 kb, respectively. The average densities of InDels were found to be 79.4 (LGR, 38.0 insertions and 41.35 deletions), 63.3 (PB 1121, 30.36 insertions and 32.94 deletions), 44.67 (Bindli, 21.44 insertions and 23.22 deletions) and 40.0 (Sonasal, 19.26 insertions and 20.77 deletions) per 100 kb. The chromosomal distribution of SNPs and InDels frequency is shown in Supplemental Figure 5A-B.

### Large scale validation of SNPs and identification of novel variations

Recently, a large SNP dataset (3KRG) was generated by the genome sequencing of 3000 diverse rice accessions (Alexandrov et al., 2015; Mansueto et al., 2017). So, in order to validate the SNPs identified in these newly sequenced four indian genotypes, we compared them with 3KRG multi-allelic 32 million SNP dataset. Distribution of the SNPs among both the datasets (common and novel) over different chromosomes is shown in Figure 2A. This comparative analysis revealed that 3KRG dataset represents 97.7% of SNPs identified in our genotypes (Figure 2B). Thus, sharing of ∼98% SNPs authenticate the nucleotide variations identified in the four genotypes. On the other hand, 79369 (2.3%) SNPs discovered in these four genotypes did not match to 3KRG dataset seems to be novel (Figure2B). One recent study has proposed that the genetic variation in rice crop is near saturation with the sequencing of diverse 3000 accessions, and suggested that any newly sequenced genotype may only add up to ∼1000 novel SNPs (Wang et al., 2018). Identification of such large number of novel SNPs in contrast to 3KRG dataset indicates the still unexplored potential of genetic diversity among indian rice germplasm. Among the four genotypes, LGR contributed the most (24%) number of novel SNPs, suggesting it to be very different among the studied genotypes (Figure 2C). Bindli accounted for least number (13%) of novel SNPs. These novel nucleotide variations especially those from LGR, can be employed for more marker generation in crop improvement breeding programs.

**Figure 2:**
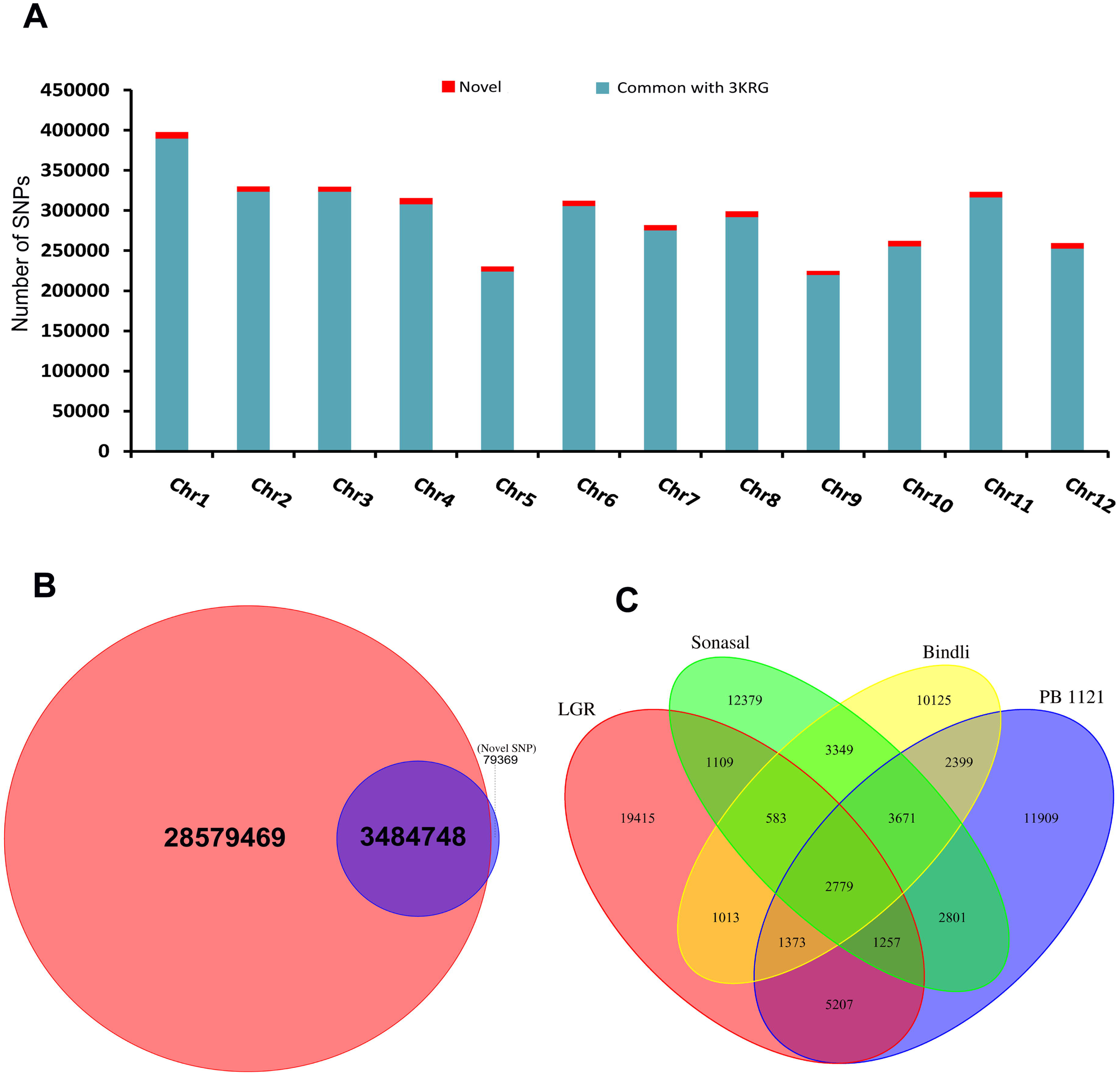
Comparative analysis of SNPs identified among our four genotypes with 3KRG (3000) rice accessions. (**A**) Chromosome wise bar plot distribution of common (our four and 3KRG dataset) and novel SNPs identified in this study. (**B**) Pie chart share among 3004 accessions, red circle; 3kRG dataset, purple circle; total SNPs of four genotypes. (**C**) Venn diagram distribution of novel SNPs (79369) identified among four indian genotypes with comparison to other 3000 accession.

### Pair-wise analysis of sequence polymorphism

Pair-wise analysis was carried out to identify SNPs and InDels between LGR and PB 1121 (LG/PB), LGR and Sonasal (LG/SN), LGR and Bindli (LG/BN), PB 1121 and Sonasal (PB/SN), PB 1121 and Bindli (PB/BN), and Sonasal and Bindli (SN/BN) rice genotypes. Maximum number of polymorphisms was found in LG/BN followed by LG/SN and least variations were found in SN/BN (Supplemental Table 6). Even the density of polymorphisms measured as number of SNPs and InDels per 100 kb was maximum for LG/BN followed by LG/SN and minimum for SN/BN (Supplemental Table 7 & 8). These nucleotide variations were scattered across all the 12 chromosomes (Figure 3A-B). SNPs and InDels in LG/BN and LG/SN were 3.5 and 3.4 folds higher than SN/BN again highlighting the potential of LGR as a source of novel variations for rice improvement (Supplemental Table 6). Among all the four genotypes, genomes of Sonasal and Bindli showed minimum polymorphism and seem to be very similar. This might be because both are short grain rice genotypes grown commonly in India. Alignment of these polymorphisms on different chromosomes revealed several low density (≤ 5 SNP per 100 kb) and high density (≥ 500 SNP per 100 kb) SNP regions (Figure 3C; Supplemental Table 9). Low (≤ 2 per 100 kb) and high (≥ 50 per 100 kb) density regions for InDels were also detected (Figure 3D; Supplemental Table 9). As per above criteria, maximum high density SNP regions (2085) and high density InDel regions (2774) were found in the case of LG/BN whereas most low density SNP regions (1005) were detected for SN/BN. Low density InDel regions (390) were detected for PB/SN followed by SN/BN. When, very stringent criteria were adopted to uncover both polymorphism hotspots (≥ 1000 SNP per 100 kb) and no SNP or InDel regions, we found more number of suppressed polymorphic regions in the case of PB/SN, PB/BN and SN/BN suggesting that PB 1121, Sonasal and Bindli might have common origin. Comparison of LGR genome with these three genotypes decreased the number of low diversity regions substantially, suggesting it to be very different in the pool and might have a different origin.

**Figure 3:**
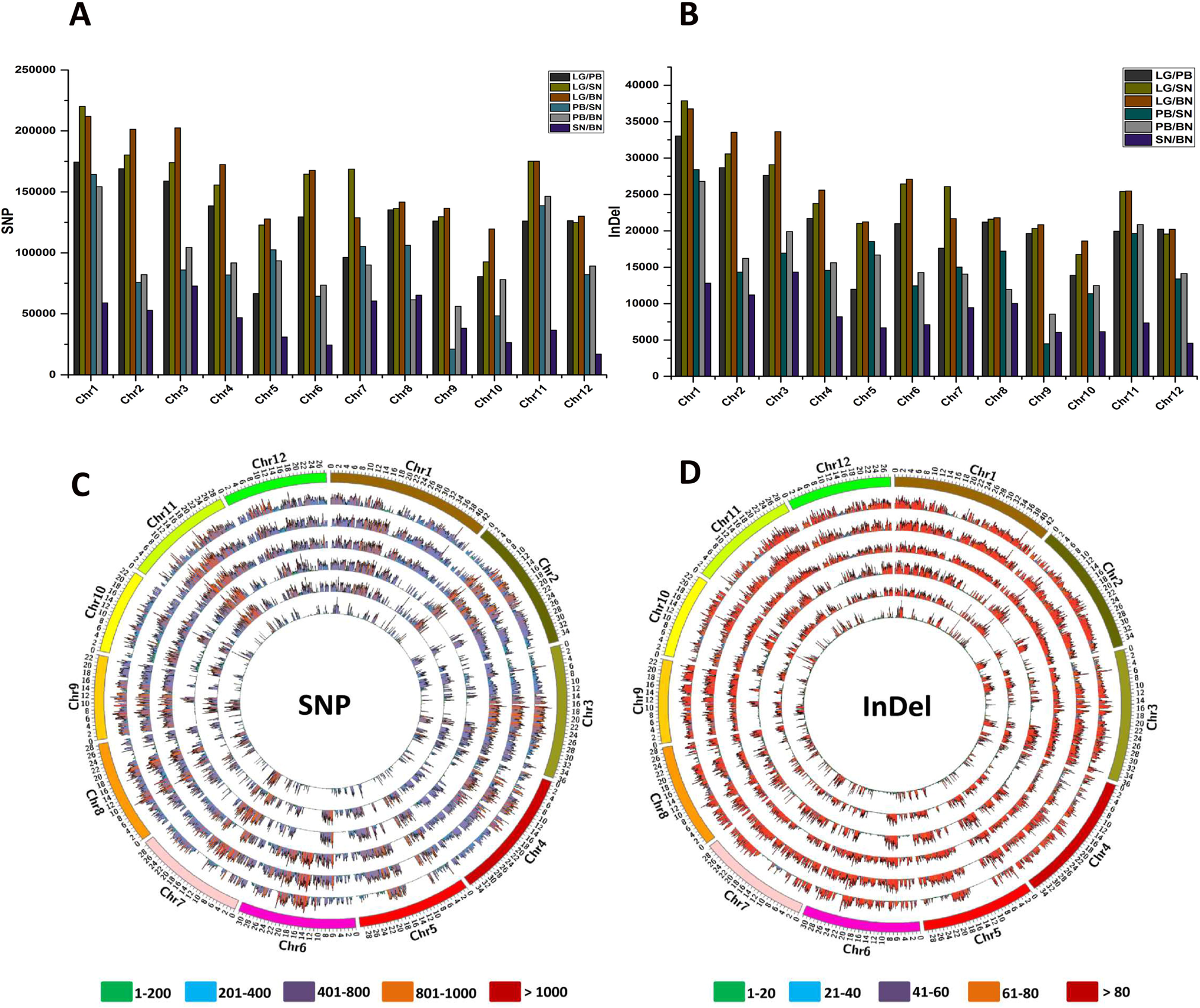
Abundance and distribution of sequence polymorphism identified on the rice chromosomes for pair wise genotypes combinations. Total number of SNPs (**A**) and InDels (**B**) identified on each rice chromosome are represented with bar graphs. LG/PB denotes SNPs/InDels detected between LGR and PB 1121 genotypes, LG/SN denotes SNPs/InDels detected between LGR and Sonasal genotypes, LG/BN denotes SNPs/InDels detected between LGR and Bindli genotypes, PB/SN denotes SNPs/InDels detected between PB 1121 and Sonasal genotypes, PB/BN denotes SNPs/InDels detected between PB 1121 and Bindli genotypes and SN/BN denotes SNPs/InDels detected between Sonasal and Bindli genotypes. Distribution of SNPs (**C**) and InDels (**D**) identified on each rice chromosome (100 kb window size) is represented with the Circos diagram. The 12 rice chromosomes (Chr1-12) are shown on outermost circle, middle to innermost circles represent SNP/InDel distribution in LG/PB, LG/SN, LG/BN, PB/SN, PB/BN and SN/BN comparison. Different range of SNPs and InDels are depicted by different colours as shown at the bottom of the Circos diagram.

### Phylogeny estimation of four sequenced genotypes

As mentioned earlier, the four rice genotypes used in this study were collected from different parts of India and were selected based on difference in their grain size/weight (Figure 1). So, in order to determine the varietal group of these genotypes, large scale phylogenetic analysis was carried out by merging the MAF-filtered genome-wide SNPs (∼4.8 million) of 3KRG dataset (Alexandrov et al., 2015; Mansueto et al., 2017). All the four genotypes fell in the major clade of *japonica* (Figure 4A). PB 1121, Sonasal and Bindli were placed at the edge of aromatic subgroup whereas LGR was found to be placed at the edge of temperate *japonica* clade (Figure 4A). This is in accordance with earlier SSR marker based characterization of PB 1121 as premium basmati, and Sonasal and Bindli as short grain aromatic landraces (Singh et al., 2000; Singh et al., 2018). On the other hand, Nipponbare was placed in the center of the *japonica* clade, indicating that unlike Nipponbare, the four genotypes might be very different from the core members of *japonica* group. This explains the discovery of so many novel SNPs in these four Indian genotypes. So, we suggest that these four genotypes and others present at the edges of different clades of this phylogenetic tree can be used as source of novel polymorphisms for rice improvement programs. To ascertain the above phylogeny, principle component analysis was also employed, which divided the accessions into 5 major groups (Figure 4B). PC 1 and PC 2, the two major variation capturing axes, indeed corroborated the results of maximum likelihood phylogeny.

**Figure 4:**
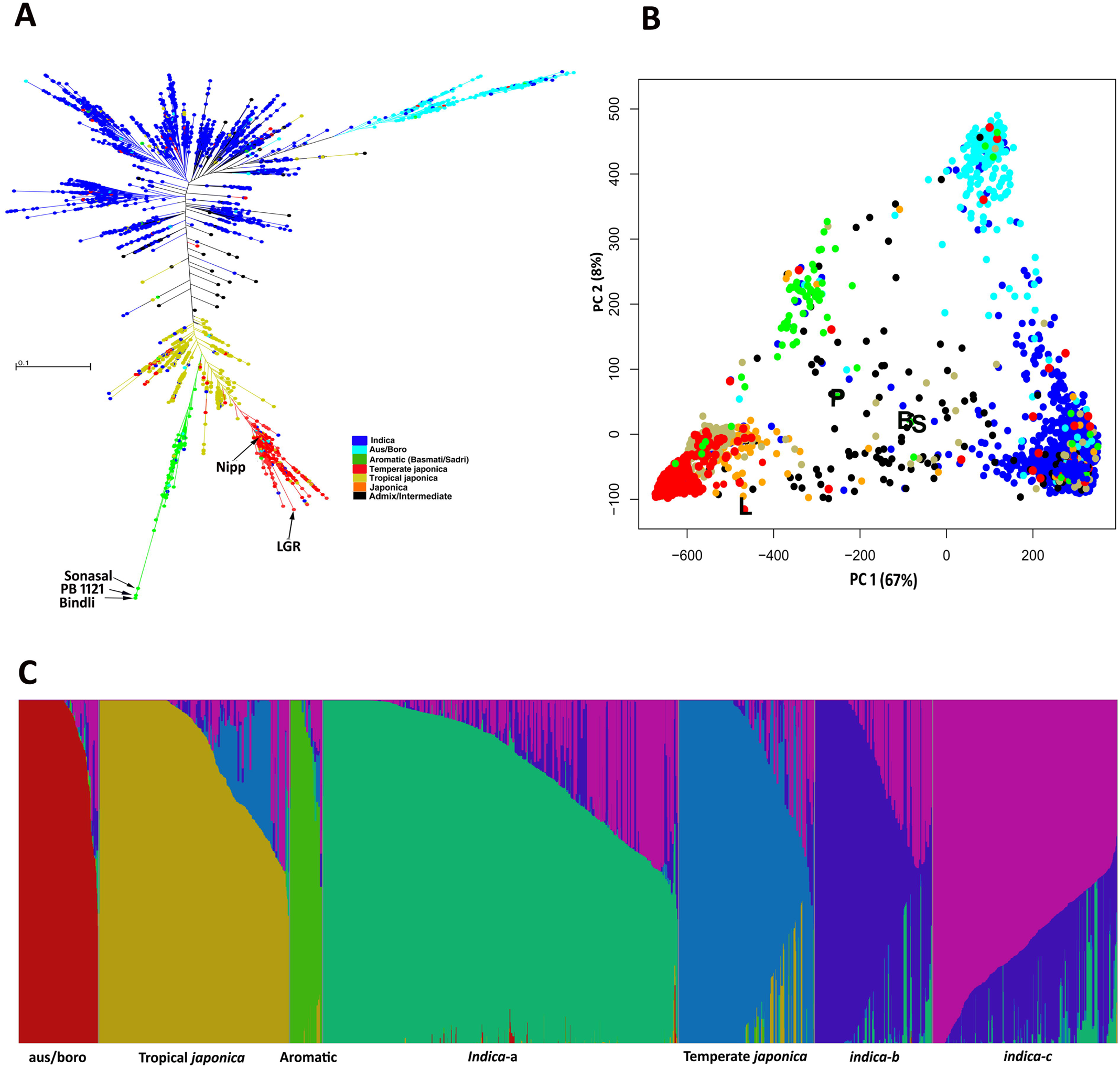
Phylogeny of 3004 rice accessions using genome-wide SNPs. (**A**) Phylogenetic tree analysis (Maximum likelihood estimation) of 3004 rice accession. Rice population groups are shown by different colors. (**B**) Bayesian method structural analysis based on genome-wide SNPs at K=7 for 3004 rice accessions. (**C**) Principal component analysis for 3004 rice accessions. Positions of LGR, PB 1121, Sonasal and Bindli are marked with name initials. Color depiction is same as in phylogenetic tree, in addition black and orange represents admixed and unclassified *japonica*, respectively.

Bayesian based population structuring separated the 3004 rice accessions in 7 (best K) population clusters, supporting the phylogenetic analysis (Figure 4C). LGR was found to carry major fraction (70%) of temperate *japonica* followed by that of *indica*, and found to be least admixed (Supplemental Table 10). Interestingly, LGR genome was found to be devoid of any aromatic genomic fragment corroborating its position away from the aromatic clade. PB 1121, Sonasal and Bindi were found to carry significant portion (mean 20%) of aromatic gene pool and hence clubbed together within the aromatic group. Therefore, we suggest that despite being the members of *japonica,* genomes of aromatic varieties such as PB 1121, Sonasal and Bindli might have accumulated the *indica* gene pool fraction via gene flow/introgression due to common cultivation with *indica* cultivars in the Indian subcontinent (Supplemental Table 10).

Besides this, in order to understand the ancestry of basmati/sadri rice, we performed population structural analysis of all 3004 rice genotypes (1743 *indica*; 840 *japonica*; 215 *aus*; 71 aromatic and 135 admix). Interestingly, with K=3, *japonica* (50%), *aus* (37%) and *indica* (13%) groups emerged as the major contributors, in descending order, of aromatic genome, suggesting that the genomic hybridization/introgression among the *japonica* and *aus* as the probable origin of aromatic rice group (Supplemental Table 11). One recent study also suggested the similar outcome for aromatic rice ancestry (Civan et al., 2019).

### Comparative genome analysis of LGG with SGG

Comparative analysis of the genomes of Long Grain genotypes (LGG; LGR and PB 1121) with that of Short Grain genotypes (SGG; Bindli and Sonasal) identified 504439 SNPs and 78166 InDels. Absolute number of these identified SNPs and InDels over different chromosomes is shown in Figure 5A whereas their distribution on all the chromosomes is shown in Figure 5B. As expected, most of nucleotide variations were found in the intergenic regions, though about 30% SNPs and 26% InDels were also detected in the genic regions (Figure 6A-B). There were 267374 (229525 SNPs and 37849 InDels) variants mapped to the regulatory regions (2 kb upstream or/and downstream) of 20550 genes, in which 169132 variants (145003 SNPs and 24129 InDels) were found within 2 kb upstream regions of 17297 genes. The genic region variations had the density of 89.9 and 12.2 per 100 kb for SNP and InDel, respectively. Within the genic region, introns were found to harbour the maximum number of SNPs (∼54%) and InDels (68.3%) (Figure 6C-D). High fraction of SNPs (∼36%) and InDels (∼11%) were found in the CDS regions, though considerable proportion of SNPs (∼10%) and InDels (∼21%) were also identified in the Untranslated Regions (UTRs). Number of InDels was more in UTR regions (21%) as compared to CDS regions (∼11%) (Figure 6C-D). So, polymorphism in these regions suggests that both divergence in the gene function and expression regulation contribute to grain size/weight in the four genotypes. Within CDS regions, missense SNPs were significantly higher (t test, p-value < 2.2e-16) than nonsense SNPs. In chromosome-wise analysis of genic region variations between LGG and SGG, significantly elevated (t test, p-value = 1.1e-07) frequency (202.5 per 100 kb) of SNPs was found on Chromosome 11 (Supplemental Figure 6A). This is interesting because there are seven other chromosomes, which are bigger than Chromosome 11, and gene density on each rice chromosome is nearly same, ranging from 143 genes per Mb on Chromosome 11 to 153 genes/Mb on Chromosome 3. These SNPs were present on more than 45% genes of Chromosome 11, followed by Chromosome 5 (∼38%) and (∼32%) Chromosome 12 (Supplemental Figure 6B). We noted that more number of genes on Chromosome 9 and Chromosome 2 were found to be resistant to nucleotide variations as only ∼10% and 16% of their genes, respectively, accumulated the SNPs. Thus, it seems that genes on different rice chromosomes vary in their inclination to accumulate SNPs and is independent of chromosome size, gene number and gene density. Moreover, the characterization of nonsynonymous variation and associated KOG enrichment are discussed in Supplemental Text, Supplemental Figure 7 & 8A. Large-effect variations can cause severe aberrations in protein structure and function, and include alteration of splice sites, introduction of premature stop codon, loss of stop codon, disruption of translation initiation codon and frame shift mutation (Supplemental Table 12). Total 4001 genes including 2536 non-T.E ones were found to carry large-effect polymorphisms. The GO analysis revealed that most of these genes were involved in modification of proteins and other macromolecules (Supplemental Figure 8B). At the molecular function level, genes coding for activities like nucleic acid binding, hydrolase activity, pyrophosphatase and protein kinase activity, and other membrane-localised proteins were significantly represented (Supplemental Figure 8C). In addition, domain analysis revealed enrichment of genes encoding for protein kinases, leucine rich repeats, NB-ARC and tyrosine kinases (Supplemental Figure 9).

**Figure 5:**
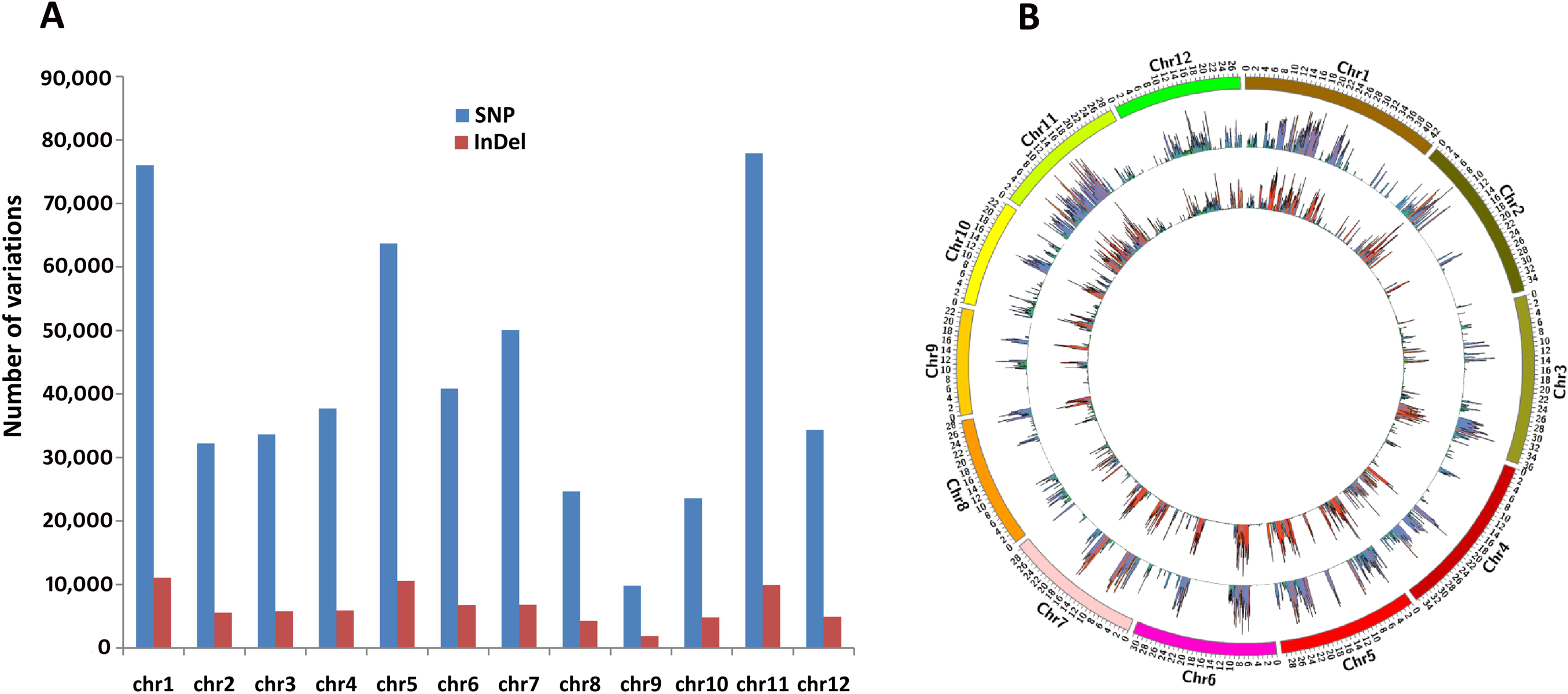
Number and distribution of SNPs and InDels identified between LGG and SGG. (**A**) Total number of SNPs and InDels identified on different rice chromosomes are represented with the bar graphs. (**B**) Abundance of SNPs and InDels and their distribution per 100 kb on different rice chromosome is represented with the Circos diagram. The 12 rice chromosomes (Chr1-12) are shown on outermost circle, middle and innermost circle represents SNP and InDel distribution, respectively. Different range of SNPs and InDels are depicted by different colours as indicated in Figure 3. LGG, Long grain genotype; SGG, Short grain genotype.

**Figure 6:**
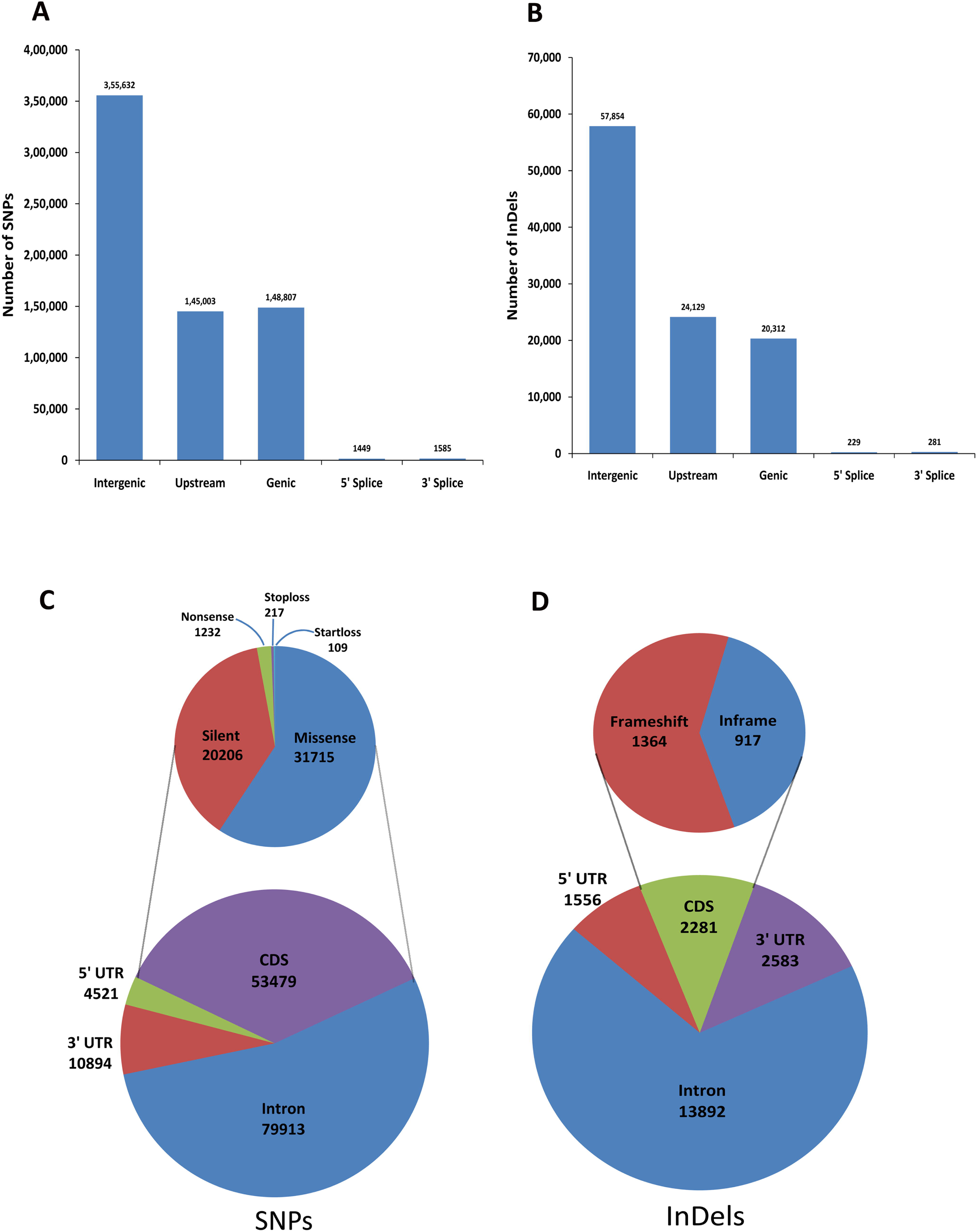
Annotation of SNPs and InDels identified between LGG and SGG comparison. Distribution of SNPs (**A**) and InDels (**B**) in different genomic regions is represented with bar graphs. Distribution of SNPs (**C**) and InDels (**D**) in different genic regions are represented in pie diagrams. LGG, Long grain genotype; SGG, Short grain genotype.

### Polymorphism analysis in known grain size/weight-related genes

There are 52 genes known to be linked with grain size/weight (Wang et al., 2012; Nakagawa et al., 2012; Huang et al., 2013; Folsom et al., 2014; Liu et al., 2015; Che et al., 2015; Si et al., 2016). Among the four genotypes, we found 1642 sequence variations (1385 SNPs and 257 InDels) within 49 of these 52 genes. Interestingly, we did not find any variations in *OsEBS*, *IPA*/*OsSPL14* and *qSW5*/*GW5* genes (Supplemental Table 13). Most of the variations (526) were found in *LRK* gene cluster (*qGY2-1*), which consist of 8 *LRK* genes present in tandem on chromosome 2. On the other side, when nucleotide variations differentiating LGG and SGG were analyzed in these grain size/weight-associated genes, 542 SNPs and 60 InDels were found in 34 genes (Supplemental Table 13). There was a noticeable suppression in the frequency of SNPs (15.9 per gene) or InDels (1.7 per gene) in these grain size/weight-associated genes compared to genome-wide genic SNP (32 per gene) and InDel (4.9 per gene) frequency validating the importance of these genes in LGG and SGG. Importantly, *LRK* gene cluster again emerged as the major site of polymorphism with 356 variations (347 SNPs and 9 InDels). This QTL, which has introgressed from a wild rice accession Dongxiang, consists of *LRK1, LRK2*, *LRK3, LRK4, LRK5, LRK6, LRK7* and *LRK8* genes (LI et al., 2002; He et al., 2006). *LRK* genes are known to affect the yield-related traits (Zha et al., 2009; Kang et al., 2017). Maximum variations (78) was found in *LRK5* and minimum (5) was identified in *LRK3*. Noticeably, in *LRK8* gene, due to occurrence of a nonsense SNP, catalytic domain (cd00180) found truncated in both the long grain genotypes (LGR and PB 1121). Thus, this analysis suggests importance of *LRK* genes in the differentiation between long grain and short grain genotypes.

### GWAS of grain size/weight trait using genome-wide informative SNPs in rice

Comparison of SNPs (504439) differentiating LGG and SGG with 700k SNPs present on a high-density SNP array (McCouch et al., 2016) based on physical positions (bp) identified 34486 common SNPs loci. Further, the GWAS was performed by integration of SNP genotyping (34486 SNPs) and the grain-length phenotyping data for 1150 rice accessions using the methods described previously (Lipka et al., 2012; Kumar et al., 2015). The GWAS identified a total of five significant associations on four different rice chromosomes, including C to A substitution in *GS3* gene, SNP-5.5371749 (G to A) downstream of *GW5* gene and T to G (SNP-11.1644056) substitution on chromosome 11 (Supplemental Table 14). The above fishing of *GS3* and *GW5* validated the outcome of the GWAS analysis, in addition to identification of a novel association [SNP-11.1644056 (T to G)]. Therefore, the SNPs differentiating the low and high grain weight accessions identified in our study have functional significance for large-scale GWAS and thereby, can be utilized for various genomics-assisted breeding applications in rice.

### Validation of grain size/weight-associated genomic loci in RIL mapping population

To validate genomic locus associated with grain length in rice, the SNPs exhibiting parental polymorphism (between PB 1121 vs. Sonasal and PB 1121 vs. Bindli) were genotyped in five of each short and long grain length homozygous mapping individuals of the mapping population [(PB 1121 × Sonasal) and (PB 1121 × Bindli)]. The SNP (SNP-3.16732086) with C to A substitution in *GS3* gene as well as SNP-5.5371749 (G to A) downstream of *GW5* gene and SNP-11.1644056 (T to G) exhibiting strong association with grain length (based on GWAS) were validated in the said mapping population. All short (4.1-5.5 mm) and long (13.1-14.6 mm) grain length parental accessions and homozygous individuals of a RIL mapping population contained the identical short and long grain length-associated SNP alleles identified by GWAS. Henceforth, a stronger marker allele effect of SNP locus with short and long grain length differentiation in rice was evident. A strong trait association potential of C to A SNP in *GS3* gene and G to A SNP downstream of *GW5* gene with the QTL controlling grain length is agreed well with the previous studies (Wan et al., 2008; Anand et al., 2013). The SNP-11.1644056 (T to G) associated with grain length identified by us appears to be novel. However, large-scale validation and genetic mapping of a grain length-associated SNP in the numerous bi- parental mapping populations contrasting for grain length are needed to assure the definitive association potential. Collectively, grain length-associated SNP validated by both GWAS and in RIL mapping population can be considered as target candidate for grain size/weight regulation.

### Identification and Characterization of polymorphic desert around centromeric region of chromosome 5

During the analysis of polymorphism frequency across the genomes of four genotypes, a region between 8.2 to 14.1 Mb on chromosome 5 was found to harbor very less SNPs and InDels. In this region, 5940 nucleotide sequence variations were found (5149 SNPs and 791 InDels) among the four genotypes with reference to Nipponbare. In earlier studies, this 8.97 Mb to 13.5 Mb centromeric spanning region of chromosome 5 was described as polymorphism desert in wild and cultivated Asian rice genotypes (Krishnan S et al., 2014; Rathinasabapathi et al., 2015; Singhabahu et al., 2017; Trinh et al., 2017). So, going by these literatures, we also call this region of 8.2-14.1 Mb on chromosome 5 as polymorphism desert. As this desert region is spans the centromeric/pericentromeric part, the repetitive nature of heterochromatin might hinder efficient generation and assembly of sequencing reads (Skipper, 2007). So, to rule out the possibly of low polymorphism due to lower coverage or biased read generation, an alignment rate analysis was performed. We found 45-61 and 57-81 reads per base for chromosome 5 and ∼ 6 Mb desert region, respectively, suggesting that there was no anomaly in the sequencing (Supplemental Table 15). The pair-wise comparison revealed 40 SNPs per 100 kb in the case of LG/SN (12.2 to 12.3 Mb) and 12 InDels per 100 kb in the case of LG/BN (between 13.9 and 14.0 Mb). In the case of LG/PB, this desert region was found to be extended on both sides from 7.4 to 14.8 Mb in LG/PB (Figs.3C-D). It was postulated in earlier studies that depressed recombinations in centromeric regions could be the reason of reduced SNP and InDel frequencies (Feltus et al., 2004; McMullen et al., 2009). So, the mean SNP and InDel densities at the centromeric regions of all the chromosomes were calculated and compared with rest of chromosomal regions (Supplemental Table 16-17). There was no definite trend in the fluctuation of polymorphism densities over centromeric regions but the centromeric region of chromosome 5 showed significantly suppressed (t test, p-value=2.2e-7) polymorphism in comparison to other chromosomes. The SNP and InDel density varied from 13.5 to 20.5 and 1.5 to 3.5 per 100 kb, respectively. Thus, this finding suggests that such low polymorphic status in and around centromeric region of chromosome 5 is unique among rice chromosomes, and is not related to the recombination ability.

For more insight, this low diversity region was analyzed for gene density and no significant difference within the polymorphism desert (142/Mb) and other regions (152.6/Mb) of chromosome 5 was observed (Supplemental Figure 10). Gene annotation revealed 540 genes as transposons out of 853 loci present within the desert region. Whereas rest of the genes (313) were found to be involved in very fundamental processes like post-translational modification and protein turnover (20.5%), transcription regulation (20.5%), carbohydrate transport and metabolism (20%) (Supplemental Figure 11).

### Molecular mapping of grain size/weight QTL within desert region

Adaption to specific environmental condition is often associated with suppressed mutations, limiting the loss of diversity in specific chromosomal areas harbouring the effective gene(s) (Olsen et al., 2006; Tang et al., 2010; Flowers et al., 2012). The observation of polymorphism suppressed region across chromosome 5 centromeric region led us investigate its possible involvement into seed size/weight regulation. We observed a significant phenotypic variation for grain size/weight (1000-grain weight: 9.3 to 28.7 g with 82% heritability) trait in 190 mapping individuals (PB 1121 x Bindli) and two parental accessions. A normal frequency distribution, including a bi-directional transgressive segregation of grain size/weight trait in mapping individuals and parental accessions was evident. The two years multi-location field phenotyping data of grain size/weight and genotyping information of 18975 SNP markers genetically mapped on 12 rice chromosomes were integrated for molecular mapping of QTLs. This analysis identified four major genomic regions harbouring four QTLs associated with grain size/weight that were mapped on four rice chromosomes (2, 5, 7, 9). The individual major QTLs explained 15.7 to 33.5% phenotypic variation (R^2^) for grain size/weight trait at 4.3 to 4.8 LOD. Of these, the genomic region underlying QTL mapped on 15.76 and 17.91 cM genetic positions of a linkage group 5 of a constructed high-density genetic map (PB 1121 × Bindli) explained maximum phenotypic variation for grain size/weight (33.5% and 4.8 LOD) in rice (Figure 7). The genetic positions (cM) of flanking SNPs, S05_8000770 [T/C] and S05_9091756[C/T] corresponded to 8000770 and 9091756 bp physical positions, respectively, which interestingly reside within the identified desert region on chromosome 5. This grain size/weight QTL identified by us in the desert region of chromosome 5 is novel and robust, and thus have potential to be employed in marker-assisted breeding methods and matter of further exploration in diverse population.

**Figure 7:**
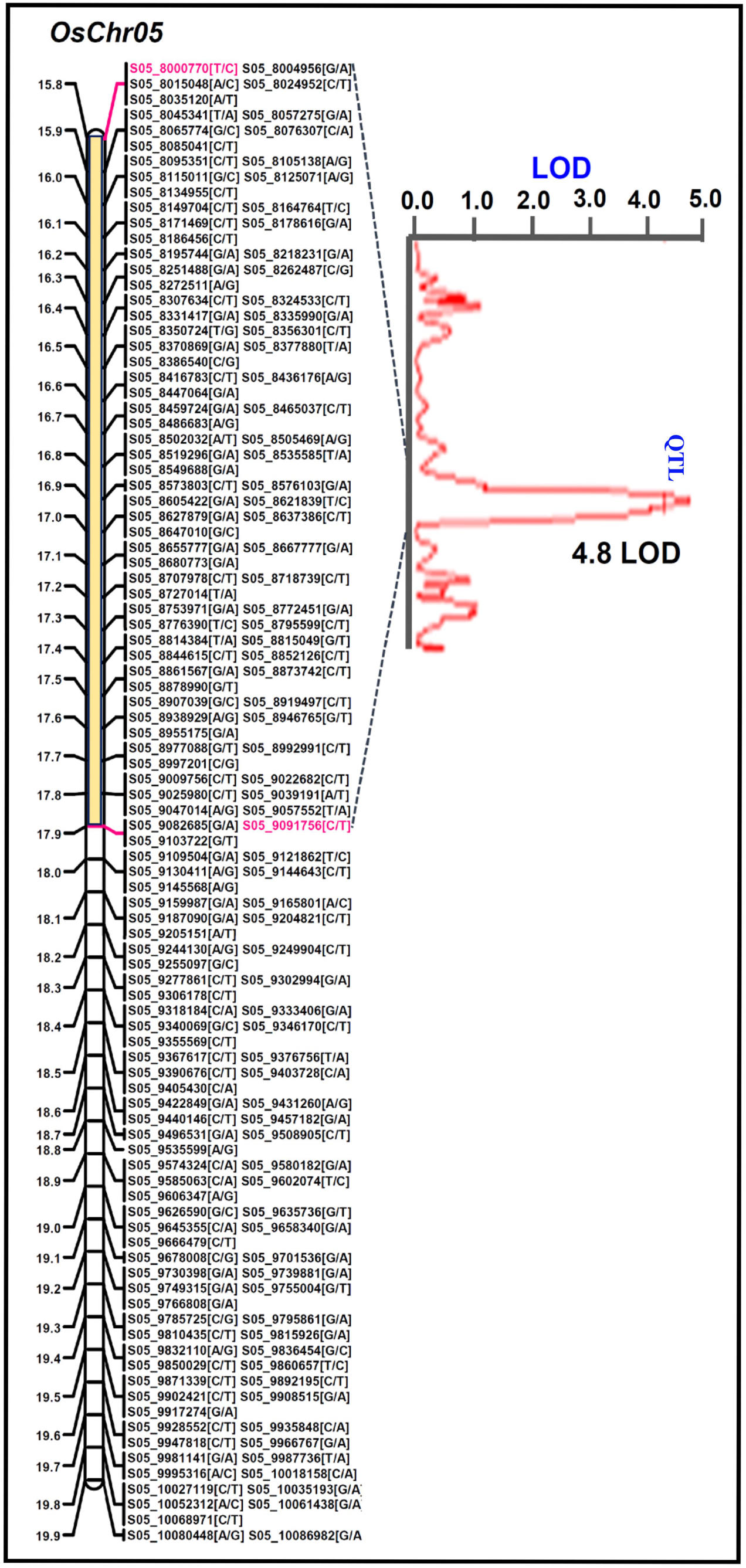
Molecular mapping of a significant major grain size/weight QTL on chromosome 5. The identified QTL flanked by parental polymorphic SNPs, S05_8000770[T/C] and S05_9091756[C/T] mapped on 15.76 and 17.91 cM genetic positions, respectively, of a linkage group 5 of a constructed high-density genetic map (PB 1121 × Bindli) of rice. The genetic positions (cM) of flanking SNPs, S05_8000770 [T/C] and S05_9091756[C/T] correspond to 8000770 and 9091756 bp physical positions, respectively, on chromosome 5.

### Characterization of chromosome 5 desert region across diverse 3000 accessions

Since there is limited information on this desert region and we identified one major QTL region associated with grain size/weight within the desert region in our mapping population (PB 1121 X Bindli), we looked for the polymorphism/diversity status of this region across much diverse 3000 accessions with filtered 3KRG dataset (Alexandrov et al., 2015; Mansueto et al., 2017).

Across 3004 genotypes, we found a total 69286 SNPs within the desert region, varying from 5 to 25946. About 7.4% (5149) of the SNPs were shared by our four genotypes and 60.5% SNPs in four genotypes were common with 3KRG SNP dataset. SNP density was found to vary from 0.08 to 432.4 per 100 kb across 3004 genotypes suggesting that not all genotypes had suppressed polymorphism rate across the centromeric/pericentromeric region of chromosome 5. Next, we calculated the nucleotide diversity (π, 100kb π window) over the chromosome 5 for all 3004 rice accessions. Once again significantly reduced diversity (t test, p-value = 2.0e-6) was found in the 8-14 Mb region, revealing presence of this long low diversity region in several diverse rice accessions (Figure 8A). Next, the boundary for this low diversity region was ascertained by analysing different regions (8-14Mb, 9-14Mb and 10-14Mb) for significant variation in diversity and compared that with entire chromosome 5 mean diversity (Supplemental Table 18).

**Figure 8:**
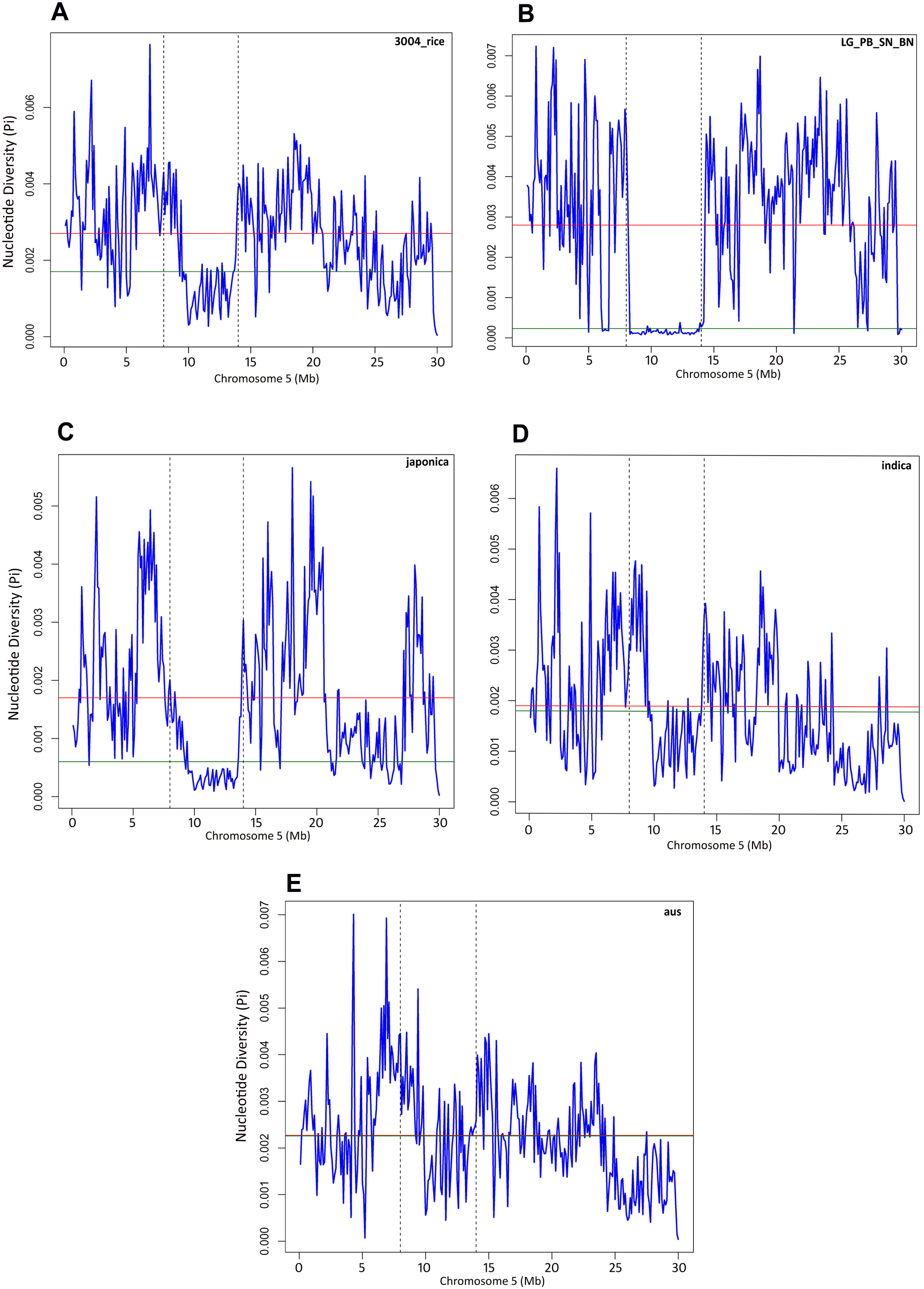
Analysis and distribution of nucleotide diversity (π/Pi) per 100 kb of chromosome 5 for different rice subspecies. **(A)** Pi distribution for all 3004 accessions**. (B)** Pi distribution for our four (LG; LGR, PB; PB 1121, SN; Sonasal and BN; Bindli) indian genotypes**. (C)** Pi distribution for all *japonica* (840) accessions**. (D)** Pi distribution for all *indica* (1743) accessions**. (E)** Pi distribution for all *aus* (215) accessions. All x and y axes represents chromosome window per Mb and Pi value, respectively. Red and green horizontal line represents mean Pi value for entire chromosome and desert (8-14Mb) region, respectively. Vertical dash line marks the boundary (8-14Mb) of desert region.

Next, we looked at the diversity in chromosome 5 of our four indian genotypes, 840 *japonica*, 1743 *indica* and 215 *aus* accessions (Figure 8B-E & Supplemental Table 18). Both *japonica* (t test, p-value = 5.3e-6) and *indica* (t test, p-value = 0.09) groups but not the *aus* group (0.36; NS) showed less nucleotide diversity in the centromeric region. In fact, *indica* accessions showed very less diversity (t test, p-value = 3.2e-4) in the 9-14Mb region, suggesting significance of this region and corroborating the identification of one grain size/weight QTL among our four indian genotypes. Contrastingly, *aus* group showed high nucleotide diversity in this region, suggesting altogether different selection constraint over this rice population. On the similar note, pair-wise genetic differentiation (*F*_ST_) analysis (100kb window) across chromosome 5 have showed sharp reduction in diversity (t test; p-value = 1.1e-10) over the desert region between *japonica* and *indica* [*F*_ST_ (0.12)]. This is in contrast to *F*_ST_ value observed for *japonica* vs *aus* (0.63), and *indica* vs *aus* (0.34) (Figure 9 & Supplemental Table 19). Next, we have performed the Tajima’s D test to determine the selection nature over this desert region and compared with mean D of entire chromosome 5. Across the 3004 genotypes there was 2 fold reduction in D value of this region (Supplemental Figure 12 & Supplemental Table 20). Among the different rice groups, *japonica* (−1.7) seems to experience significant mean positive/purifying selection and *indica* (1.1) was subjected to weaker balancing selection. Interestingly, *indica* observed relatively more positive selection (0.78) over 9-14Mb. This is again in contrast to *aus* rice that showed elevated D value (mean 3.5), suggesting strong balancing selection (Supplemental Figure 12 & Supplemental Table 20). Overall, these result corroborated the results observed for nucleotide diversity and *F*_ST_ analysis.

**Figure 9:**
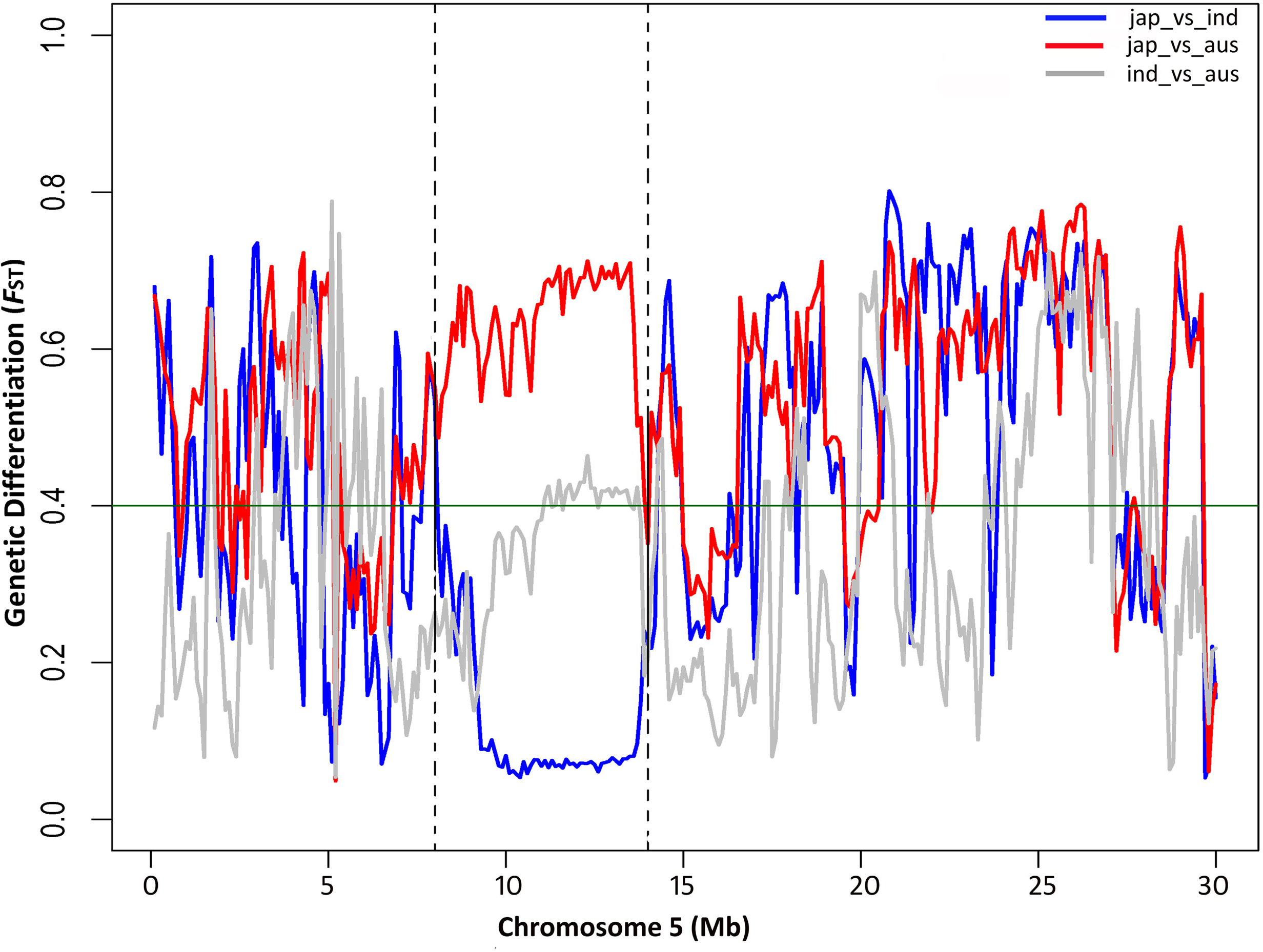
Distribution of genetic differentiation (*F*_ST_) per 100kb across chromosome 5 between different rice subspecies. Scan of genetic differentiation between *indica*, *japonica* and *aus*. The position of chromosome 5 desert region is shown in the region at 8–14.0 Mb (*x* axis) and the level of genetic differentiation (*F*_ST_) on y-axis. In total, 2798 rice accessions were used for analysis, including 1743 *indica*, 840 *japonica* and 215 *aus*. Green line represents the mean *F*_ST_value (0.41) for chromosome 5 among the *aus, indica* and *japonica.* Vertical dash line marks the boundary (8-14Mb) of desert region.

It is considered that selection leaves a genomic signature represented by strong linkage disequilibrium (LD) over selective sweeps as well as adjacent regions (Fijarczyk and Babik, 2015). So, whole chromosome 5 was scanned to understand the dynamics of Linkage disequilibrium (LD) pattern and integrity over this 8-14Mb low diversity polymorphism desert region. Interestingly, we have observed significant elevation in LD over the desert region for all the population groups (Supplemental Figure 13 & Supplemental Table 21). These finding indicates elevated togetherness of various alleles within the desert region and their possibility to co-segregate during recombination and introgression.

In order to ascertain the genetic closeness of *aus* group with wild rice, we have also analysed another large dataset containing 1529 (wild + culticated) accessions including 520 *indica*, 484 *japonica*, 30 *aus* and 446 wild (O. rufipogon) genotypes (Huang et al., 2012). Recently the same dataset was analyzed for genome-wide introgression distribution to show that *japonica*, *indica* and *aus* groups are domesticated independently, where *aus* group experienced the most introgression events (Civáň and Brown, 2018). In order to study the sharing of alleles with wild population, distribution of allele frequency spectrum (AFS) over chromosome 5 was prepared for different rice population (Supplemental Figure 14 & Supplemental Table 22). In case of *japonica*, desert region scored mean allele frequency of 0.99 as against 0.98 for chromosome-wide, suggesting further allele fixation over the region under selection. Interestingly, *indica* rice scored very significant uplift (0.94) for desert region as compared to entire chromosome (0.83). This indicates that alleles are nearly fixed over desert region in *japonica* and *indica.* On the other hand, both *aus* and wild rice populations had same allele frequency (0.84) for the desert region and the whole chromosome (0.83).These results suggest a clear divergence of *aus* rice from *japonica* and *indica* over the desert region and its genetic proximity with wild rice. Recent studies have also indicated that in contrast to modern varieties, *aus* group exhibit the evolutionary proximity with wild rice and is composed of genome prior to most domestication bottleneck events.

Further, in-depth analysis of 3KRG dataset found that *aus* group genotypes have retained much more SNP/allelic variations than *indica* (t test, p-value = 3.4e-06) and *japonica* (t test, p-value = 1.5e-10) genotypes and seems to be have experienced weak positive selection at chromosome 5 desert region (Supplemental Table 23). So, to identify the outliers, tukey method using interquartile range was employed, which determined SNP density of 197.2 per 100 kb as the upper threshold value (Figure10A). However, this cut-off value for categorization for desert presence or absent genotypes, hereafter called Desert and non-Desert genotypes was tested by principle component analysis (PCA) method. The above PCA divided the 3004 rice accessions into two major distinct populations and depicted a mean SNP density (210 SNPs/100 kb) similar to tukey threshold for intermediate/admixed genotypes (falling in between the two separated clusters) (Figure 10B). Thus, considering the above SNP density threshold for this 8-14Mb low diversity region categorization, 2519 rice accessions (84%) with SNP densities under the threshold were classified as Desert genotypes, whereas 485 accessions with SNP density over the threshold were assigned as Non-Desert genotypes. Even the bayesian population structuring for this genomic region divided the 3004 rice accessions into two groups (best K=2), corroborating the findings of tukey and PC methods, and independently validating the separation of Desert and Non-Desert genotypes (Figure 10C). On the same note, the phylogenetic analysis based on SNPs within this desert region too displayed independent clustering of Desert and Non-Desert genotypes (Figure 10D). Such a large population of desert genotypes justifies the importance of suppressed polymorphic status of chromosome 5 centromeric region during rice adaptation and its genome evolution. Interestingly, as we inferred, significant number of the *aus* genotypes (63%) were found to be Non-Desert genotypes (t test, p-value = 5.1e-14; χ^2^, p-value = 6.2e-10). On the other hand, only 17.8% of *indica* and ∼3% of *japonica* belonged to Non-Desert group. We also suggest that most of the *aus* genotypes did not face selection constrain over the course of domestication and evolution, and remained closer to the wild rice accessions. We also believe that selection pressure for specific agronomic requirements constrained the gene flow/introgression from *japonica* and/or *indica* to the traditional *aus* group. This hypothesis is backed by a recent study highlighting introgression spectrum, from *japonica* and *indica* to *aus* rice (Civáň and Brown, 2018). We noted that source of the majority of Non-Desert accessions is Bangladesh (∼23%) and India (∼22%). Interestingly, Indo-bangla geographical region has been already proposed as home ground for adaptation/domestication of *aus* genome (Civán et al., 2015) (Supplemental Table 24).

**Figure 10:**
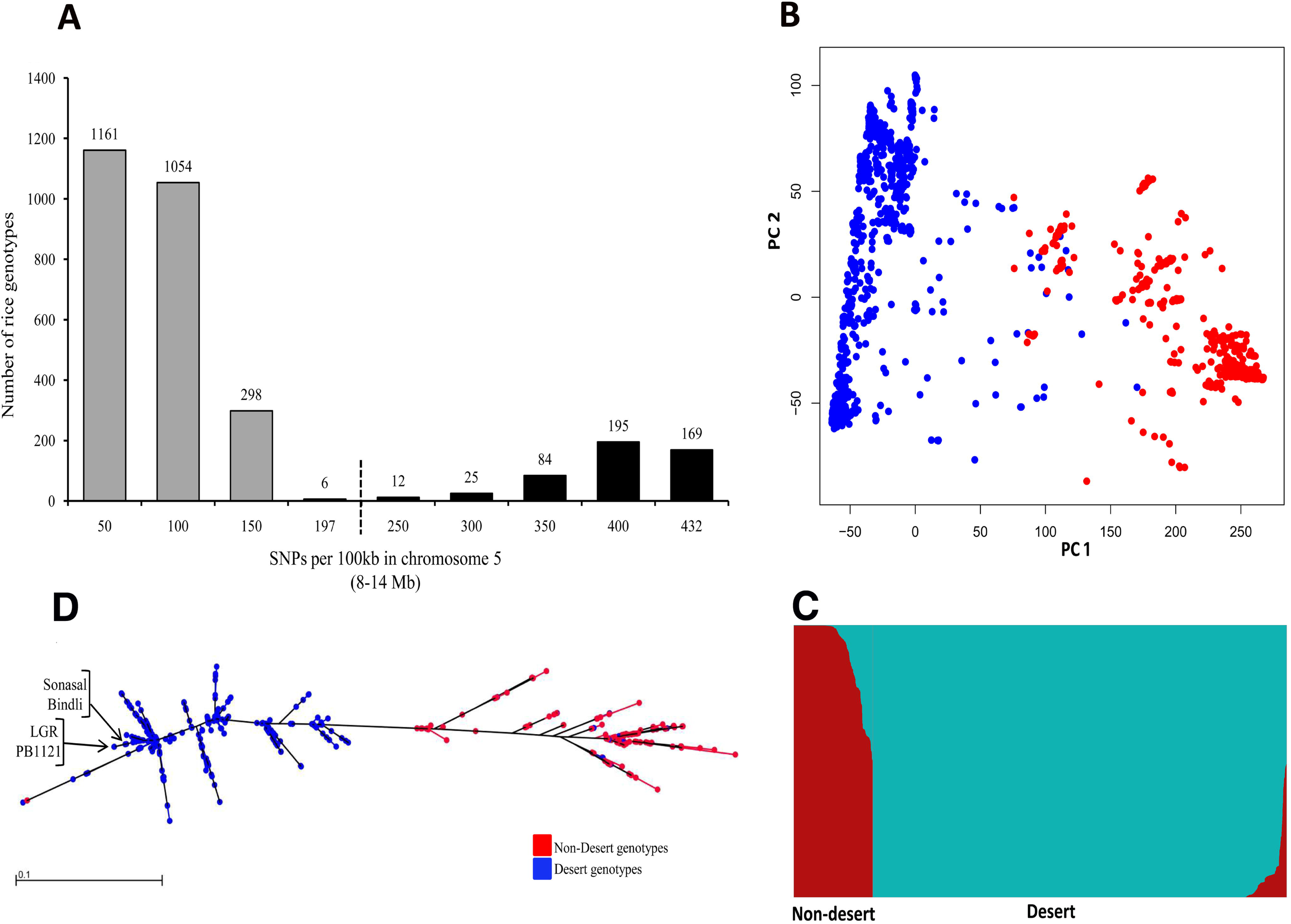
Distribution and phylogenetic divergence of 3004 rice accessions based on SNPs identified within chromosome 5 desert region (8-14 Mb). (**A**) Total number of rice accessions having ranges of SNPs detected in desert region is shown in the bar graphs. The number of rice accessions within expected range of SNP density is indicated with grey bars and the outlier accessions with more than expected SNPs per 100 kb are shown in black bars. The cut-off value for outliers is shown with dotted line. (**B**) Principal component analysis for 3004 rice accessions for the desert region. (**C**) Bayesian method structural analysis over the desert region at K=2 for 3004 rice accessions. (**D**) Phylogenetic distance correlation and divergence based on maximum likelihood estimation. Color depiction for B and D is same, based on desert and non-desert genotype categorization.

### Multiple overlapping selective sweeps within *japonica* and *indica* over chromosome 5 centromere spanning desert region

Selective sweeps are usually observed in and around the genomic regions (spanning few 100 kbs) associated with traits adopted during rice cultivation and domestication (Olsen et al., 2006; Tang et al., 2010). So, we hypothesized that this ∼ 6 Mb polymorphism desert region might have witnessed multiple selective sweeps and harboring loci crucial to grain size/weight and other important traits. We scanned the whole length of chromosome 5 for evidence of selection signal using two different methods: Cross Population Extended Haplotype Homozogysity (XP-EHH) and Cross-Population Composite Likelihood Ratio test (XP-CLR) (Sabeti et al., 2007; Tang et al., 2007; Chen et al., 2010). We found significantly elevated mean XP-EHH score (1.53) for the ∼6 Mb desert region in the Desert genotypes with respect to Non-Desert group. Consistent to this, we found significantly elevated mean (XP-EHH) selective sweep score over the desert region within *japonica* and *indica* genotypes than *aus* rice group (Figure 11A-C). Specifically, in case of *japonica* to Non-Desert group, the desert region had elevated mean XP-EHH score of 2.0 (t test, p-value < 2.2e-16) compared to overall score (0.4) across the chromosome 5. Similarly, *indica* population also showed increased mean XP-EHH score (1.5) for the desert region (t test, p-value < 2.2e-16) in comparison to overall score (0.27) for chromosome 5. However, in *aus* population, mean XP-EHH score (0.26) for desert region was similar to overall score (0.2) for the chromosome 5. The XP-EHH method relies on linkage disequilibrium (LD), which breaks down quickly over time and provides weak power to detect historical sweeps that ended up to several thousand generations ago. To circumvent this concern, another method XP-CLR was employed (Chen et al., 2010). Corroborating the findings of XP-EHH analysis, this method also showed significantly elevated XP-CLR score (51.9) for the desert region in Desert genotypes with respect to Non-Desert group. Consistent to this, we found significantly elevated score over the desert region for *japonica* and *indica* genotypes as well, but not for *aus* group (Figure 11D-F). In the case of *japonica* vs Non-Desert group, the desert region showed elevated mean (119.2) XP-CLR score (p-value=1.4e-13), compared to overall score (32.1) across the chromosome 5. Similarly, *indica* population also witnessed surge in XP-CLR score (66.8) for the desert region (p-value, 4.2e-10) in comparison to overall score (5.3) for chromosome 5. Based on these findings, we can interpret that the strong phenomenon of multiple and overlapping selective sweeps happened over the centromeric/pericentromeric region of chromosome 5. Furthermore, the amplitude of selection was stronger within *japonica* than *indica* population in contrast to *aus* population and thought to be created historically long time ago. This observation is also in accordance with the recent studies suggesting that *aus* rice group might have originated from distinct wild progenitor (Civán et al., 2015; Choi et al., 2017; Civáň and Brown, 2017). We suggest that chromosome 5 desert region is very crucial genomic region that might be harbouring important trait loci created by targeted selection, more specifically in *japonica* and *indica* genotypes owing to their phylogeographical isolation from *aus* group during the course of domestication.

**Figure 11:**
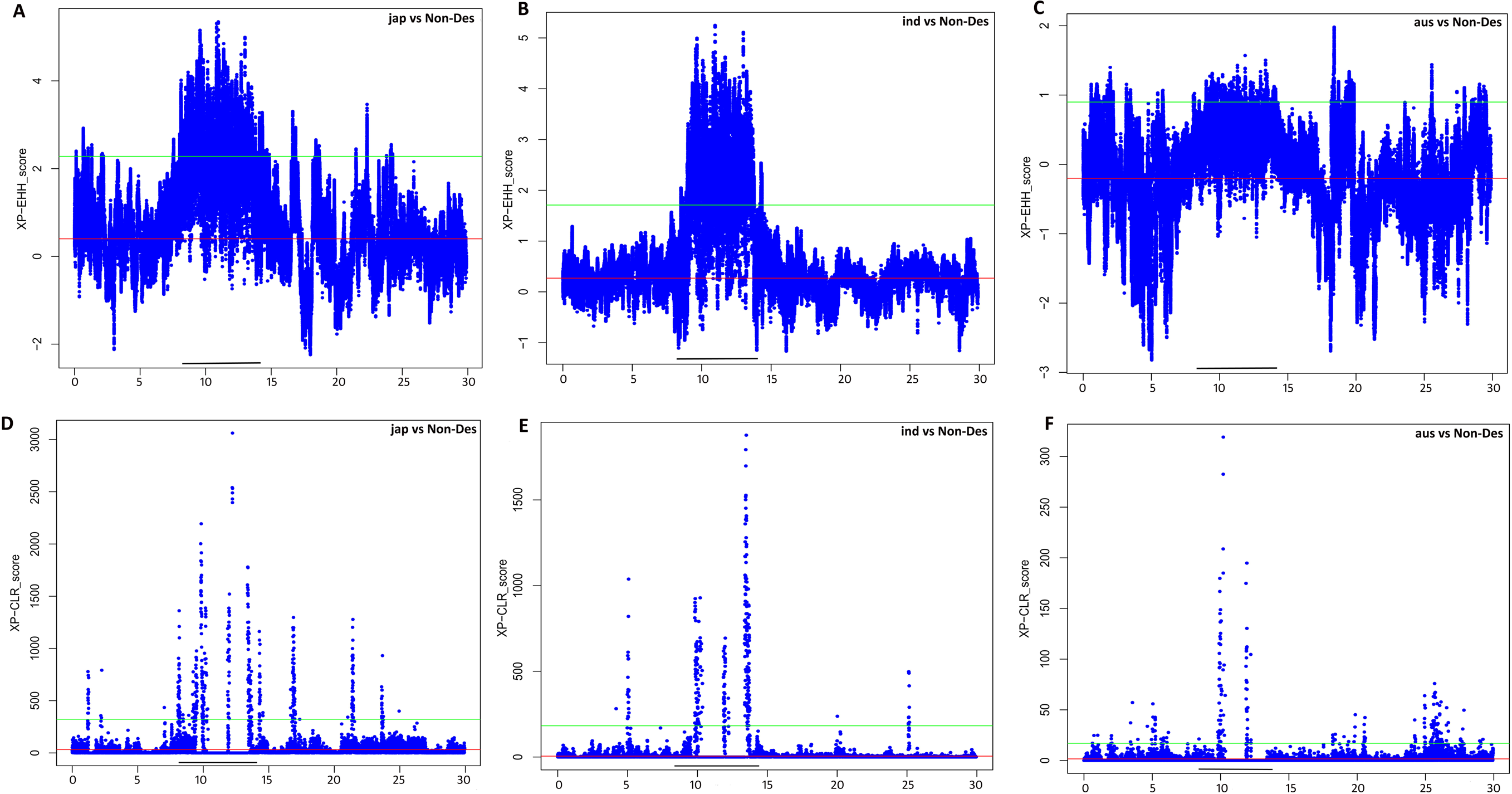
Selective sweep scan (per 100 kb) over chromosome 5 for different rice varietal groups. (**A-C**) Analysis of chromosome-wide XP-EHH score for the detection of selective sweep(s) signal. (**A**) Scan for *japonica* group with reference to categorized non-desert (non-des) accessions. (**B**) Scan for *indica* group with reference to categorized Non-Desert accessions. (**C**) Scan for *aus* group with reference to categorized Non-Desert accessions. (**D-F**) Analysis of chromosome-wide XP-CLR score for the detection of selective sweep(s) signal. (**D**) Scan for *japonica* group with reference to categorized Non-Desert accessions. (**E**) Scan for *indica* group with reference to categorized Non-Desert accessions. (**F**) Scan for *aus* group with reference to categorized Non-Desert accessions. Where x-axis represents the chromosomal length (Mb). Red line represents the mean selection score for chromosome-wide scan for respective varietal group. Green line represents the top 5% cut-off value based on z-score. Black bar depicts the range of desert region (8-14 Mb).

### Association analysis of chromosome 5 desert for grain size/weight and other domestication-related traits

Cultivation of rice started about 10-12 ka (Khush, 1997; Choi et al., 2017) and during its domestication several key traits have been selected generation after generation. Our analyses suggested this 8-14 Mb region on the chromosome 5 had distinct and reduced diversity (*F*_ST_) between two largest rice subspecies *indica* and *japonica* in contrast to *aus* population (Figure 9). This is further backed with strong positive selection signal and LD pattern among the *japonica* and *indica* genotypes (Supplemental Figure 12-13). These observations led us to speculate association of this desert region with domestication events. So, attempt was made to correlate 69286 SNPs identified in this region among the 3000 rice accessions with different domestication-related traits including grain size/weight. Interestingly, significant associations were identified for grain length, width and weight (Figure 12A-C; Supplemental Table 25). Moreover, in support of the QTL region identified within the chromosome 5 desert region in PB 1121 x Bindli mapping population, one SNP of G/A polymorphism at 8027721 of chromosome 5 showed significant association with all the three (length, width and weight) important grain size/weight parameters (Figure 12A-C; Supplemental Table 25). Although, the minor allele frequency was towards lower side, it showed strong association explaining about 15% variation for grain weight, 12% variation for grain length and remarkable 27% variation for grain width. It seems that G/A SNP could cause variation in grain size/weight more effectively in *indica* and tropical *japonica* rice genotypes. This nucleotide variation (G/A) was found to be present just 4 bases away from GTCAT element known as cis-regulatory Skn-1_motif (Lescot et al., 2002), which is required for the expression of genes in the endosperm (Takaiwa et al., 1991; Wu et al., 2000; Jiang et al., 2014). In addition, an ubiquitin conjugating enzyme coding gene (*OsUBC43*, LOC_Os05g14300) having preferential expression during panicle and seed development was found at 6.4 kb downstream of G/A SNP site (Kawahara et al., 2013). GW2, an E3 ligase (LOC_Os02g14720) is known to negatively regulate grain size in rice, mainly the width (Song et al., 2007). Further study is required to explore the probable collaboration between *OsUBC43* and *GW2* in regulation of grain size/weight trait. Interestingly, in our analysis, five other domestication-related traits such as panicle axis, seed shattering, heading date, seed coat color and culm length, were also found to be significantly associated with some other SNPs residing in the chromosome 5 desert region (Supplemental Figure 15 & Supplemental Table 25). Thus, desert region of chromosome 5 seems to be very important genomic region harbouring loci associated with various crucial domesticated-related traits including grain size/weight.

**Figure 12:**
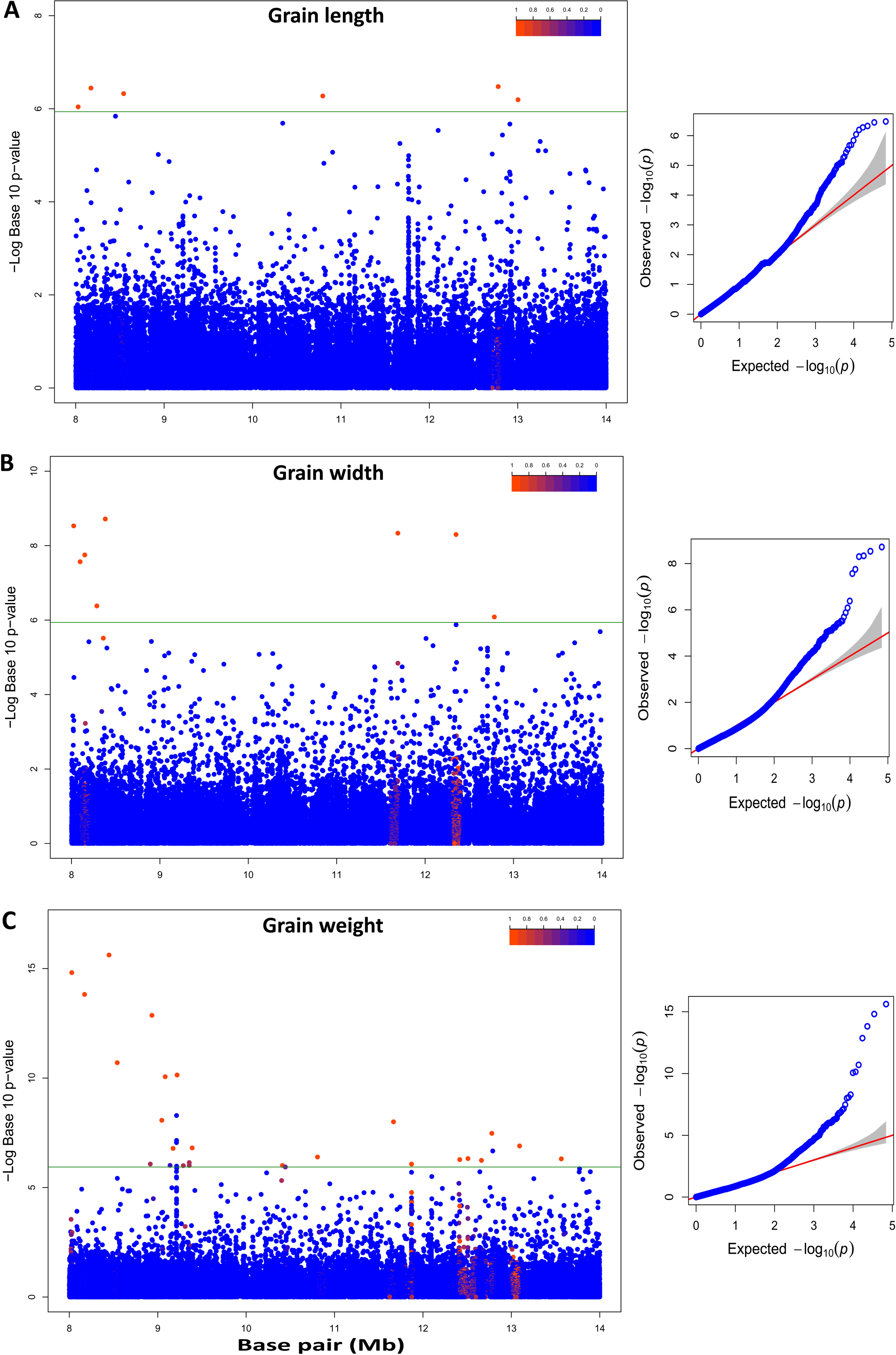
Mapping of associations for various grain size/weight traits across chromosome 5 desert (8-14 Mb) region. **(A)** Manhattan and QQ plots of compressed MLM for grain length. Negative log_10_- transformed *P* values (y axis) values from the compressed mixed linear model are plotted against position (x axis) on chromosome 5 desert region. (**B**) Manhattan and QQ plots of compressed MLM for grain width, as in A. (**C**) Manhattan and QQ plots of compressed MLM for grain weight, as in A. Green line represents the cut-off for significant marker association. Color bar (blue to red) denotes extent of Linkage Disequilibrium.

## Discussion

In the present study, we performed genome resequencing of four indian rice genotypes with contrasting grain size/weight trait and found a significant number of high-quality DNA polymorphisms (SNPs and InDels). Notably, our study revealed a substantial numbers of novel sequence variants (SNPs and InDels), hitherto unreported even among 3000 rice accession analyses. We found that the diverse nature of genotypes (especially of LGR and to some extent Sonasal and Bindli) considered for the study were responsible for larger sequence variations. This highlights the necessity of further studying the polymorphism in many other indian rice genotypes. All the four resquenced genotypes were found to be member of *japonica*. LGR grouped with temperate *japonica* and PB 1121, Sonasal and Bindli turned out to be aromatic varieties. The novel nucleotide variations identified in this study highlights the unexplored potential of indian genotypes that can be useful for marker assisted selection (MAS) in breeding programs.The frequency of SNPs and InDels were higher in the coding and regulatory regions. We believe that the genetic basis of contrasting rice seed size/weight underlying the large-effect SNPs and InDels identified here would provide valuable insights. Further, the functional relevance of SNPs based on the integrated analysis of DNA polymorphisms within seed size/weight QTLs and conserved functional domains led to identification of putative candidate genes. These genes could be possibly involved in the regulation of seed development or seed size/weight determination and can be used as targets for crop improvement after functional validation with forward or reverse genetics approaches. In GWAS, 5 SNPs were found to be significantly associated with grain size/weight trait, and the top three SNPs were validated in two different RIL mapping populations raised by crossing LGG and SGG. Appearance of two previously known QTLs, *GS3* and *GW5*, in this analysis validated the approach adopted in this study. Therefore, the novel polymorphic site SNP-11.1644056 (T to G), which was also validated in two different RIL mapping populations can be a strong candidate for various genomics-assisted breeding applications in rice.

We identifies a ∼6 Mb long polymorphism suppressed region (8-14 Mb), spanning the centromere/pericentromere of chromosome 5, previously named as polymorphism desert, in four sequenced rice genotypes. The nucleotide diversity (Pi) analysis with 3004 accessions revealed presence of this 8-14Mb low nucleotide diversity region in most of *japonica* and *indica* accessions. This region was found to possess positive selection within *japonica* and *indica*, in contrast to balancing selection observed in *aus* rice. Subsequent analysis revealed significant LD elevation over desert region in all the rice groups, maximum in *indica*. Moreover, a reduced genetic differentiation (*F*_ST_) between *indica* and *japonica* in contrast to *aus* was noticed over the desert region. However, to ascertain the direction of this *aus* divergence, its genetic proximity with wild rice was assessed by allele frequency spectrum over chromosome 5. Indeed, we found elevated alleles fixation over this selection region for *japonica* and *indica.* In wild and *aus* groups, the allele frequency was found to be similar and unchanged.

Our study divided 3004 rice accessions into Desert and Non-Desert genotype. Although, most of the rice genotypes (84%) were found to be devoid of frequent allelic variations within desert region, *aus* genotypes were found to retain much elevated allelic variations. Moreover, this low diversity region was identified as the hotspot for multiple and overlapping selection sweeps, more pronounced in *japonica* and *indica* but not in *aus* population. This might be due to domestication, as modern varieties (*japonica* and *indica*) have been under strong selection for specific agronomic traits, constraining the adoption/selection of genomic regions from the standing variation within their progenitors. However, the *aus* genotypes, representing traditional genotypes that are genetically much closer to wild rice varieties, were found to be subjected to a lesser degree of selection. Thus, we believe this ∼ 6 Mb region as an evolutionary long selection site adopted into *japonica* and *indica* from their wild progenitors, whereas observation of intermittent desert region in some *aus* genotypes may be attributed to gene flow/introgression from *japonica* and *indica*. Noticeably, the desert region-targeted association analysis revealed presence of SNPs that were associated with several important domestication-related agronomic traits including grain size/weight. Interestingly, this desert region was found to contain one QTL region significantly associated with grain size/weight in RIL mapping population (PB 1121 x Bindli). Different analysis results such as elevated tajima’s D, enhanced LD, increased allele fixation and long selection sweeps indicate chromosome 5 desert region as a probable domestication footprint. No wonder, a major QTL for grain size/weight was identified in it.

We suggest that this important region can be utilized as valuable resource for the better adaptability and yield-related traits. Interestingly, the genomic hybridization/introgression among the *aus* and *japonica* was indicated as probable origin of aromatic rice group, clearing the speculation regarding the ancestry of basmati/sadri rice. Overall, this comprehensive study can be an important resource for genome-wide genotyping and establishing the relationship between grain traits with genomic regions, which can be very useful in rice improvement programs. All these data and analyses can be retrieved from our RiceSzWtBase.

## Materials & Methods

### Plant materials, extraction and genomic DNA sequencing

All the rice accessions [long grain: LGR (LG) and PB 1121 (PB); short grain: Sonasal (SN) and Bindli (BN)] were grown in the same season of 2015 in NIPGR field. For the analysis of seed parameter, matured and dried seeds of all the genotype were subjected to length, width and weight measurement. Rice seedling of 10 days old were used for the isolation of genomic DNA using Sigma GenElute™ Plant gDNA kit. Integrity of genomic DNA was analyzed using Bioanalyzer 2100 (Agilent Technologies, Singapore). Samples for sequencing were prepared by IlluminaTruSeq DNA sample preparation kit (Illumina, USA). Approximately, one μg high quality genomic DNA was fragmented using Covaris to generate a library of 300 to 400□bp average insert size. Sheared DNA was subjected to end repairing, adenylation of 3′ ends followed by indexed paired end ligation. Ligated products were purified for the size range of 400 to 500−bp with QiagenMinElute gel extraction kit and further enriched by PCR. The generated libraries were further assessed for concentration, desired fragment size and pooled as per manufactures protocols. Sequencing was performed with 90 bases pair-end chemistry on IlluminaHiSeq 2000.

### Read filtering and alignment

Subsequent standard bioinformatics analysis such as, raw reads quality check and removal of low quality bases (< Q30 Phred score) was done. The bases from the 5’ and 3’ end were trimmed for any specific bias observed in base composition. After performing the quality trimming, adapter sequences were also trimmed from the reads. The filtered reads were then mapped over the rice reference genome Nipponbare (IRGSP-1.0 pseudomolecule/MSU7) using BWA program with –q20 setting. Picard program was employed to get rid of duplicate reads and further retained sequence reads were used for the alignment with reference genome.

### Variant calling and annotation

Post sequence reads realignment, single nucleotide polymorphisms (SNPs and InDels) were called via Genome Analysis TKLite-2.3-9 toolkit Unified Genotyper (GATK). The SNPs and InDels with a polymorphism call rate of < 90% were eliminated. After calling, in order to eliminate the low quality variants, a stringent criteria of read depth ≥10 and quality score □≥□30 was adopted to filter total variants and only good quality variants were considered for further analysis. Further, all the SNPs consecutive and adjacent to InDels were also eliminated. Genome-wide distribution of SNPs and InDels per 100 kb was graphically depicted using Circos program (Krzywinski et al., 2009). Plant Ensembl database was used to obtain gene model for annotation. All the identified SNPs and InDels were annotated using customized VariMAT (SciGenome, India) programme for intergenic, genic, intron, exon, splice-site, 5′ UTR, 3′ UTR and coding-region annotation. The same language programme was used for the mapping of variants to all transcript forms and prediction of different variant classes such as silent, missense, non-sense, start-loss, stop-loss, inframe and frameshift.

### Phylogenetic analysis

3KRGP filtered (∼4.8 million) SNP dataset was retrieved from SNP seek web resources (Alexandrov et al., 2015) and filtered to final 3000 rice accessions (due non availability of varietal and geographical data). Phylogenetic tree (maximum likelihood) generation for both complete genome and chromosome 5 desert region was generated using SNPhylo programme (Lee et al., 2014). Common SNPs among 3000 and our four rice genotypes were considered for genome-wide phylogeny. SNP data was filtered by LD information, as it improves interpretation of phylogenetic relationships from genomic data. LD threshold of 0.5 was applied for genome-wide phylogenetic tree generation due low depth data of 3KRG dataset as this allowed more SNP sites to remain in the analysis. Similarly, LD threshold of 0.9 was used for desert region phylogeny estimation. The Newick format was used to export the consensus trees and imported into Archaeopteryx (version 0.9901 beta) for topology visualization. In case of genome-wide analysis, prior information of one of the five groups – *indica*, *aus*/boro, Aromatic (basmati/sadri) and *japonica* (tropical or temperate) for 3000 rice accessions was used for assignment of varietal group for our rice genotypes.

### SNP validation

Sequencing and SNP dataset of previous reports were used for validation of identified SNPs. The Full 3K RG SNP Dataset (∼ 32 million, multi-allelic) was downloaded on 20/12/2018 from Rice SNP-Seek Database (http://snp-seek.irri.org/) and compared (Alexandrov et al., 2015; Mansueto et al., 2017). For experimental validation, 98 randomly selected SNPs distributed across the genome were amplified with their flanking sequences using customised primers. The flanking amplicons were obtained *via* PCR, using genomic DNA of four rice genotypes as template. The sequencing of PCR products were performed using BigDye^®^ Terminator v3.1 Cycle Sequencing Kit in an automated ABI 3730xl DNA Analyzer as per manufacturer’s instructions (Applied Biosystems, California, USA). Sequence variations were detected after alignment using ClustalW software.

### Gene function categorization and conserved domain prediction

Functional class categorization of genes was performed using KOG prediction method, where eggNOG version 4.5 database was queried with the protein sequences for KOG classes estimation (Huerta-Cepas et al., 2016). Gene ontology (GO) enrichment analysis was performed using agriGO online tool with p-value cut-off of ≤ 0.05 (Du et al., 2010). For prediction of conserved domain(s) in the proteins, their sequences were queried against the latest repository of predicted PFAM domains under the rice genome annotation project (Kawahara et al., 2013).

### Identification of grain size/weight QTL regions

Rice QTLs for grain size/weight and yield-related traits were downloaded from the Gramene QTL Database (http://www.gramene.org/qtl/). The corresponding start and end coordinates for each QTL as per the latest version (MSU7) of the rice genome annotation were determined via BLASTN search. The presence of genes and their overlap with the nucleotide variants within the QTL region were determined using latest GFF of rice genome annotation.

### Nucleotide diversity, Tajima’s D, LD, Genetic differentiation (*F*_ST_), Allele frequency and Selection sweep analyses

In total, 406569 SNPs identified on chromosome 5 among the 3004 rice accessions was utilised for this analysis. The R package “adegenet” and “hierfstat” was used for the pair-wise *F*_ST_ estimation between different rice population groups. All above analyses were performed with 3004 rice accessions using different modules of vcftools software with default parameters over 100kb window (Danecek et al., 2011). Analysis of selection sweep over the chromosome 5 was performed using default setting of XP-EHH (Sabeti et al., 2007; Tang et al., 2007), and XP-CLR with grid size 2500bp and 200 SNP size (Chen et al., 2010). Allele frequency spectrum for wild and cultivated rice was estimated using SNP data downloaded from RiceHap3 database (http://server.ncgr.ac.cn/RiceHap3) on 20/02/19. All subsequent plots were generated with R programme.

### Genome wide association analysis (GWAS) for grain size/weight trait

The SNPs differentiating low (Sonasal and Bindli) and high (PB 1121 and LGR) grain weight Indian accessions were identified by comparing the whole genome resequencing data of all four accessions. The identified SNPs were then matched with SNPs available on a high-density (700 K) genotyping array described in reference (McCouch et al., 2016), based on their respective physical positions. The genotyping data for identified common SNPs, available for 1146 rice accessions along with phenotypic data for respective accessions were obtained. Further the SNP genotype (filtered with minor allele frequency of > 0.05) and grain length phenotype data for 1150 rice accessions (including four accessions sequenced in the current study) were combined with their relative kinship matrix (K) and PCA (principal component analysis) information using P3D/compressed mixed linear model (CMLM) as per mentioned in the references (Lipka et al., 2012; Upadhyaya et al., 2015; Kumar et al., 2015). The inflation factor (λ) and test statistics were evaluated using the quantile-quantile (Q-Q) plot. A significance p-value (corrected for false discovery rate) threshold of 0.05 was used during the analysis. Finally, the 100 kb genomic region (based on accepted LD decay in different rice populations) on either side of most significantly associated SNPs were identified as QTL region.

### Validation of associated SNP in a RIL mapping population

To assure the potential of SNP for grain size/weight association, the trait-associated SNP was selected to validate in traditional bi-parental mapping population. For this, five of each short (Sonasal and Bindli with grain length: 5.5-6 mm) and long (PB 1121: 13.1 mm) grain length homozygous individuals derived from the two mapping population, (PB 1121 x Sonasal) and (PB 1121 × Bindli) along with parental accessions were selected for DNA isolation. The grain length-associated SNP exhibiting polymorphism between the mapping parents was genotyped in the selected five homozygous short and long grain length mapping individuals using IlluminaGoldenGate SNP genotyping assay as followed in reference (Bajaj et al., 2015). The correspondence of short and long grain length-associated SNPs with their presence in the short and long grain length homozygous mapping individuals was determined to validate the grain length trait association potential of SNPs in rice.

### Trait association analysis for chromosome 5 desert region

The genotyping data of 2262 rice accessions along with the phenotypic data of 18 different yield-related traits were obtained from SNP seek database. In this, the SNP and phenotype data of our four resequenced genotypes were added. Association analysis for chromosome 5 desert region was performed with same association model and parameters as mentioned in the earlier GWAS section.

## Data availability

Raw read sequences of LGR, PB 1121, Sonasal and Bindli are required to be deposited. The datasets generated and analysed during the current study are publically available on RiceSzWtBase database (www.nipgr.res.in/jkt/home.php).

## Authors’ contributions

AnK performed all the experiments and analyses, and drafted the manuscript. AnK, AD and SKP performed GWAS experiment. AnK, VK and ArK grew the rice varieties and performed genomic DNA isolation. GKS and AKS characterized Sonasal, Bindli and PB 1121 varieties. BCP characterized LGR variety. This collaborative project has been conceived and designed by JKT, AKT, SKP, AKS and GKS. JKT conceived this particular study and planned the experiments and analyses. AnK and JKT wrote the manuscript. All the authors read and approved the final manuscript.

## Acknowledgements

This work was financially supported by the grant BT/AB/NIPGR/SEED BIOLOGY/2012 from Department of Biotechnology (DBT), Government of India, and core grant from the National Institute of Plant Genome Research. AnK and VK acknowledge University Grant Commission for the Senior Research Fellowship. AD acknowledges DBT for Senior Research Fellowship and ArK acknowledges Short-term Research Fellowship from NIPGR. Computational facility of SubDIC at NIPGR is also acknowledged. The authors are thankful to DBT-eLibrary Consortium (DeLCON) for providing access to e-resources. We thank Dr. Manoj Prasad, NIPGR for his valuable suggestions.

## References

Alexandrov, N., Tai, S., Wang, W., Mansueto, L., Palis, K., Fuentes, R. R., Ulat, V. J., Chebotarov, D., Zhang, G., Li, Z., et al. (2015). SNP-Seek database of SNPs derived from 3000 rice genomes. Nucleic Acids Res. 43:D1023–D1027.

Anand, D., Baunthiyal, M., Krishnan, S. G., Singh, N. K., Prabhu, K. V., **and** Singh, A. K. (2013). Novel InDel variation in GS3 locus and development of InDel based marker for marker assisted breeding of short grain aromatic rices. J. Plant Biochem. Biotechnol. 24:120–127.

Bajaj, D., Upadhyaya, H. D., Khan, Y., Das, S., Badoni, S., Shree, T., Kumar, V., Tripathi, S., Gowda, C. L. L., Singh, S., et al. (2015). A combinatorial approach of comprehensive QTL-based comparative genome mapping and transcript profiling identified a seed weight-regulating candidate gene in chickpea. Sci. Rep. 5:9264.

Che, R., Tong, H., Shi, B., Liu, Y., Fang, S., Liu, D., Xiao, Y., Hu, B., Liu, L., Wang, H., et al. (2015). Control of grain size and rice yield by GL2-mediated brassinosteroid responses. Nat. Plants 2:15195.

Chen, H., Patterson, N., **and** Reich, D. (2010). Population differentiation as a test for selective sweeps. Genome Res. 20:393–402.

Choi, J. Y., Platts, A. E., Fuller, D. Q., Hsing, Y. I., Wing, R. A., Purugganan, M. D., **and** Kim, Y. (2017). The rice paradox: Multiple origins but single domestication in Asian Rice. Mol. Biol. Evol. 34:969–979.

Civan, P., Ali, S., Batista-Navarro, R., Drosou, K., Ihejieto, C., Chakraborty, D., Ray, A., Gladieux, P., Brown, T. A., **and** Niimura, Y. (2019). Origin of the Aromatic Group of Cultivated Rice (Oryza sativa L.) Traced to the Indian Subcontinent. Genome Biol. Evol. 11:832–843.

Civán, P., Craig, H., Cox, C. J., and Brown, T. A. (2015). Three geographically separate domestications of Asian rice. Nat. Plants 1.

Civáň, P., and Brown, T. A. (2017). Origin of rice (Oryza sativa L.) domestication genes. Genet. Resour. Crop Evol. 64:1125–1132.

Civáň, P., and Brown, T. A. (2018). Role of genetic introgression during the evolution of cultivated ice (Oryza sativa L.). BMC Evol. Biol. 18.

Danecek, P., Auton, A., Abecasis, G., Albers, C. A., Banks, E., DePristo, M. A., Handsaker, R. E., Lunter, G., Marth, G. T., Sherry, S. T., et al. (2011). The variant call format and VCFtools. Bioinformatics 27:2156–2158.

Du, Z., Zhou, X., Ling, Y., Zhang, Z., **and** Su, Z. (2010). agriGO: A GO analysis toolkit for the agricultural community. Nucleic Acids Res. 38:W64–W70.

Duitama, J., Silva, A., Sanabria, Y., Cruz, D. F., Quintero, C., Ballen, C., Lorieux, M., Scheffler, B., Farmer, A., Torres, E., et al. (2015). Whole genome sequencing of elite rice cultivars as a comprehensive information resource for marker assisted selection. PLoS One 10.

Feltus, F. A., Wan, J., Schulze, S. R., Estill, J. C., Jiang, N., **and** Paterson, A. H. (2004). An SNP resource for rice genetics and breeding based on subspecies Indica and Japonica genome alignments. Genome Res. 14:1812–1819.

Fijarczyk, A., and Babik, W. (2015). Detecting balancing selection in genomes: Limits and prospects. Mol. Ecol. 24:3529–3545.

Fitzgerald, M. A., McCouch, S. R., **and** Hall, R. D. (2009). Not just a grain of rice: the quest for quality. Trends Plant Sci. 14:133–139.

Flowers, J. M., Molina, J., Rubinstein, S., Huang, P., Schaal, B. A., **and** Purugganan, M. D. (2012). Natural selection in gene-dense regions shapes the genomic pattern of polymorphism in wild and domesticated rice. Mol. Biol. Evol. 29:675–687.

Folsom, J. J., Begcy, K., Hao, X., Wang, D., **and** Walia, H. (2014). Rice Fertilization-Independent Endosperm1 Regulates Seed Size under Heat Stress by Controlling Early Endosperm Development. PLANT Physiol. 165:238–248.

Gao, Z.-Y., Zhao, S.-C., He, W.-M., Guo, L.-B., Peng, Y.-L., Wang, J.-J., Guo, X.-S., Zhang, X.-M., Rao, Y.-C., Zhang, C., et al. (2013). Dissecting yield-associated loci in super hybrid rice by resequencing recombinant inbred lines and improving parental genome sequences. Proc. Natl. Acad. Sci. U. S. A. 110:14492–7.

He, G., Luo, X., Tian, F., Li, K., Zhu, Z., Su, W., Qian, X., Fu, Y., Wang, X., Sun, C., et al. (2006) Haplotype variation in structure and expression of a gene cluster associated with a quantitative trait locus for improved yield in rice. Genome Res. 16:618–626.

Huang, X., Zhao, Y., Wei, X., Li, C., Wang, A., Zhao, Q., Li, W., Guo, Y., Deng, L., Zhu, C., et al. (2011). Genome-wide association study of flowering time and grain yield traits in a worldwide collection of rice germplasm. Nat. Genet. 44:32–39.

Huang, X., Kurata, N., Wei, X., Wang, Z. X., Wang, A., Zhao, Q., Zhao, Y., Liu, K., Lu, H., Li, W., et al. (2012). A map of rice genome variation reveals the origin of cultivated rice. Nature 490:497–501.

Huang, R., Jiang, L., Zheng, J., Wang, T., Wang, H., Huang, Y., **and** Hong, Z. (2013). Genetic bases of rice grain shape: So many genes, so little known. Trends Plant Sci. 18:218–226.

Huerta-Cepas, J., Szklarczyk, D., Forslund, K., Cook, H., Heller, D., Walter, M. C., Rattei, T., Mende, D. R., Sunagawa, S., Kuhn, M., et al. (2016). EGGNOG 4.5: A hierarchical orthology framework with improved functional annotations for eukaryotic, prokaryotic and viral sequences. Nucleic Acids Res. 44:D286–D293.

Jain, M., Moharana, K. C., Shankar, R., Kumari, R., **and** Garg, R. (2014). Genomewide discovery of DNA polymorphisms in rice cultivars with contrasting drought and salinity stress response and their functional relevance. Plant Biotechnol. J. 12:253–264.

Jiang, S.-Y., Vanitha, J., Bai, Y., **and** Ramachandran, S. (2014). Identification and molecular characterization of tissue-preferred rice genes and their upstream regularly sequences on a genome-wide level. BMC Plant Biol. 14:1–14.

Jianlong, X., Qingzhong, X., Lijun, L., **and** Zhikang, L. (2002). Genetic dissection of grain weight and its related traits in rice (Oryza sativa L.). Zhongguo shuidao kexue 16:6—10.

Kanamori, H., Fujisawa, M., Katagiri, S., Oono, Y., Fujisawa, H., Karasawa, W., Kurita, K., Sasaki, H., Mori, S., Hamada, M., et al. (2013). A BAC physical map of *aus* rice cultivar “Kasalath”, and the map-based genomic sequence of “Kasalath” chromosome 1. Plant J. 76:699–708.

Kang, J., Li, J., Gao, S., Tian, C., **and** Zha, X. (2017). Overexpression of the leucine-rich receptor-like kinase gene LRK2 increases drought tolerance and tiller number in rice. Plant Biotechnol. J. 15:1175–1185.

Kawahara, Y., de la Bastide, M., Hamilton, J. P., Kanamori, H., McCombie, W. R., Ouyang, S., Schwartz, D. C., Tanaka, T., Wu, J., Zhou, S., et al. (2013). Improvement of the Oryza sativa Nipponbare reference genome using next generation sequence and optical map data. Rice (N. Y). 6:4.

Khush, G. S. (1997). Origin, dispersal, cultivation and variation of rice. Plant Mol. Biol. 35:25–34.

Krishnan S, G., Waters, D. L. E., **and** Henry, R. J. (2014). Australian wild rice reveals pre-domestication origin of polymorphism deserts in rice genome. PLoS One 9:e98843.

Krzywinski, M., Schein, J., Birol, I., Connors, J., Gascoyne, R., Horsman, D., Jones, S. J., **and** Marra, M. A. (2009). Circos: An information aesthetic for comparative genomics. Genome Res. 19:1639–1645.

Kumar, V., Singh, A., Mithra, S. V. A., Krishnamurthy, S. L., Parida, S. K., Jain, S., Tiwari, K. K., Kumar, P., Rao, A. R., Sharma, S. K., et al. (2015). Genome-wide association mapping of salinity tolerance in rice (Oryza sativa). DNA Res. 22:133–145.

Lee, T.-H., Guo, H., Wang, X., Kim, C., **and** Paterson, A. H. (2014). SNPhylo: a pipeline to construct a phylogenetic tree from huge SNP data. BMC Genomics 15:162.

Lescot, M., Déhais, P., Thijs, G., Marchal, K., Moreau, Y., Van De Peer, Y., Rouzé, P., and Rombauts, S. (2002). PlantCARE, a database of plant cis-acting regulatory elements and a portal to tools for in silico analysis of promoter sequences. Nucleic Acids Res. 30:325–327.

Li, D., Sun, C., Fu, Y., Li, C., Zhu, Z., Chen, L., Cai, H., **and** Wang, X. (2002). Identification and mapping of genes for improving yield from Chinese common wild rice (O. rufipogon Griff.) using ad-vanced backcross QTL analysis. Chinese Sci. Bull. 47:1533.

Lipka, A. E., Tian, F., Wang, Q., Peiffer, J., Li, M., Bradbury, P. J., Gore, M. A., Buckler, E. S., **and** Zhang, Z. (2012). GAPIT: genome association and prediction integrated tool. Bioinformatics 28:2397–2399.

Liu, S., Hua, L., Dong, S., Chen, H., Zhu, X., Jiang, J., Zhang, F., Li, Y., Fang, X., **and** Chen, F. (2015). OsMAPK6, a mitogen-activated protein kinase, influences rice grain size and biomass production. Plant J. 84:672–681.

Mansueto, L., Fuentes, R. R., Borja, F. N., Detras, J., Abrio-Santos, J. M., Chebotarov, D., Sanciangco, M., Palis, K., Copetti, D., Poliakov, A., et al. (2017). Rice SNP-seek database update: New SNPs, indels, and queries. Nucleic Acids Res. 45:D1075–D1081.

McCouch, S. R., Wright, M. H., Tung, C.-W., Maron, L. G., McNally, K. L., Fitzgerald, M., Singh, N., DeClerck, G., Agosto-Perez, F., Korniliev, P., et al. (2016). Open access resources for genome-wide association mapping in rice. Nat. Commun. 7:10532.

McMullen, M. D., Kresovich, S., Villeda, H. S., Bradbury, P., Li, H., Sun, Q., Flint-Garcia, S. A., Thornsberry, J., Acharya, C., Bottoms, C., et al. (2009). Genetic properties of the maize nested association mapping population. Science 325:737–40.

McNally, K. L., Childs, K. L., Bohnert, R., Davidson, R. M., Zhao, K., Ulat, V. J., Zeller, G., Clark, R. M., Hoen, D. R., Bureau, T. E., et al. (2009). Genomewide SNP variation reveals relationships among landraces and modern varieties of rice. Proc. Natl. Acad. Sci. 106:12273–12278.

Nakagawa, H., Tanaka, A., Tanabata, T., Ohtake, M., Fujioka, S., Nakamura, H., Ichikawa, H., **and** Mori, M. (2012). Short grain1 decreases organ elongation and brassinosteroid response in rice. Plant Physiol. 158:1208–19.

Olsen, K. M., Caicedo, A. L., Polato, N., McClung, A., McCouch, S., **and** Purugganan, M. D. (2006). Selection under domestication: Evidence for a sweep in the rice waxy genomic region. Genetics 173:975–983.

Rathinasabapathi, P., Purushothaman, N.,VL, R., **and** Parani, M. (2015). Whole genome sequencing and analysis of Swarna, a widely cultivated indica rice variety with low glycemic index. Sci. Rep. 5:11303.

Sabeti, P. C., Varilly, P., Fry, B., Lohmueller, J., Hostetter, E., Cotsapas, C., Xie, X., Byrne, E. H., McCarroll, S. A., Gaudet, R., et al. (2007). Genome-wide detection and characterization of positive selection in human populations. Nature 449:913–918.

Si, L., Chen, J., Huang, X., Gong, H., Luo, J., Hou, Q., Zhou, T., Lu, T., Zhu, J., Shangguan, Y., et al. (2016). OsSPL13 controls grain size in cultivated rice. Nat. Genet. 48:447–456.

Singh, R. K., Singh, U. S., and Khush, G. S. (2000). Aromatic Rices. New Delhi: Oxford & IBH Publishing Co. Pvt. Ltd.

Singh, V., Singh, A. K., Mohapatra, T., S, G. K., and Ellur, R. K. (2018). Pusa Basmati 1121 – a rice variety with exceptional kernel elongation and volume expansion after cooking. Rice 11:19.

Singhabahu, S., Wijesinghe, C., Gunawardana, D., Senarath Yapa, M. D., Kannangara, M., Edirisinghe, R., and Dissanayake, V. H. W. (2017). Whole Genome Sequencing and Analysis of Godawee, a Salt Tolerant Indica Rice Variety. Rice Res. Open Access 5.

Skipper, M. (2007). Mysteries of heterochromatic sequences unravelled. Nat. Rev. Genet. 8:567.

Song, X.-J., Huang, W., Shi, M., Zhu, M.-Z., **and** Lin, H.-X. (2007). A QTL for rice grain width and weight encodes a previously unknown RING-type E3 ubiquitin ligase. Nat. Genet. 39:623–630.

Subbaiyan, G. K., Waters, D. L. E., Katiyar, S. K., Sadananda, A. R., Vaddadi, S., **and** Henry, R. J. (2012). Genome-wide DNA polymorphisms in elite indica rice inbreds discovered by whole-genome sequencing. Plant Biotechnol. J. 10:623–634.

Takaiwa, F., Oono, K., Wing, D., **and** Kato, A. (1991). Sequence of three members and expression of a new major subfamily of glutelin genes from rice. Plant Mol. Biol. 17:875–885.

Tan, Y. F., Xing, Y. Z., Li, J. X., Yu, S. B., Xu, C. G., **and** Zhang, Q. (2000). Genetic bases of appearance quality of rice grains in Shanyou 63, an elite rice hybrid. TAG Theor. Appl. Genet. 101:823–829.

Tang, K., Thornton, K. R., **and** Stoneking, M. (2007). A new approach for using genome scans to detect recent positive selection in the human genome. PLoS Biol. 5:1587–1602.

Tang, H., Sezen, U., **and** Paterson, A. H. (2010). Domestication and plant genomes. Curr. Opin. Plant Biol. 13:160–166.

Trinh, H., Nguyen, K. T., Nguyen, L. Van, Pham, H. Q., Huong, C. T., Xuan, T. D., Anh, L. H., Caccamo, M., Ayling, S., Diep, N. T., et al. (2017). Whole-Genome Characteristics and Polymorphic Analysis of Vietnamese Rice Landraces as a Comprehensive Information Resource for Marker-Assisted Selection. Int. J. Genomics 2017:9272363.

Upadhyaya, H. D., Bajaj, D., Das, S., Saxena, M. S., Badoni, S., Kumar, V., Tripathi, S., Gowda, C. L. L., Sharma, S., Tyagi, A. K., et al. (2015). A genome-scale integrated approach aids in genetic dissection of complex flowering time trait in chickpea. Plant Mol. Biol. 89:403–420.

Wan, X., Weng, J., Zhai, H., Wang, J., Lei, C., Liu, X., Guo, T., Jiang, L., Su, N., **and** Wan, J. (2008). Quantitative trait loci (QTL) analysis for rice grain width and fine mapping of an identified QTL allele gw-5 in a recombination hotspot region on chromosome 5. Genetics 179:2239–2252.

Wang, S., Wu, K., Yuan, Q., Liu, X., Liu, Z., Lin, X., Zeng, R., Zhu, H., Dong, G., Qian, Q., et al. (2012). Control of grain size, shape and quality by OsSPL16 in rice. Nat. Genet. 44:950–4.

Wang, W., Mauleon, R., Hu, Z., Chebotarov, D., Tai, S., Wu, Z., Li, M., Zheng, T., Fuentes, R. R., Zhang, F., et al. (2018). Genomic variation in 3,010 diverse accessions of Asian cultivated rice. Nature 557:43–49.

Wu, C. Y., Washida, H., Onodera, Y., Harada, K., **and** Takaiwa, F. (2000). Quantitative nature of the prolamin-box, ACGT and AACA motifs in a rice glutelin gene promoter: Minimal cis-element requirements for endosperm-specific gene expression. Plant J. 23:415–421.

Zha, X., Luo, X., Qian, X., He, G., Yang, M., Li, Y., **and** Yang, J. (2009). Over-expression of the rice *LRK1* gene improves quantitative yield components. Plant Biotechnol. J. 7:611–620.

