## Supplemental Tables and Figures for "Genome-wide analysis of polymorphisms identified domestication-associated polymorphism desert carrying important rice grain size/weight QTL"

### Supplemental Information (SI)

#### Supplemental Text

##### Analysis of SNPs and InDels

Among all the SNPs identified in the four genotypes with reference to Nipponbare, 2531339 were found to be transitions (Ts; A/G and C/T) and 1032778 were transversions (Tv; A/C, A/T, G/C and G/T). Number of Ts detected in Sonasal, Bindli, PB 1121 and LGR were 711327, 778456, 1149764, and 1468919, respectively, whereas Tv in the same order were found to be 284218, 314884, 459092 and 580789 (Supplemental Figure 2). Here, in all the four genotypes, frequency of Ts was about 2.5 fold higher than Tv which is similar to what has been reported earlier for rice and other plants (Agarwal et al. 2012; Batley et al. 2003; Subbaiyan et al. 2012). Elevated level of Ts/Tv ratio suggests that transitions are tolerated more during mutation and selection, as they are more likely synonymous in protein-coding sequences and also less likely to disrupt RNA secondary structure than transversions (Kimura 1977). The length distribution of InDels detected in Sonasal, Bindli, PB 1121 and LGR varied from 1 to 24 bp (insertion) and up to 39 bp for deletions. (Supplemental Figure 3). Most of the InDels (~55.1%) were of single base whereas InDels of 2 to 4 bp varied from 16.1 to 5.0%. InDels longer than 4 bp accounted up to 16.1% which can be attributed to large genetic divergence of these rice accessions.

##### Analysis of loci carrying nonsynonymous variation between LGG and SGG

The ratio of nonsynonymous and synonymous SNPs for LGG/SGG was found to be 1.57. The dissemination of nonsynonymous SNP per kb per gene was found to be in the range of 0.043-25.5. However, 622 genes were found to have unusual high density (t test, p-value $< 2.2e-16$) of nonsynonymous SNPs (> 4.04 SNPs per kb, Tukey method) than the expected range (Supplemental Figure 7). Though most of these outlier genes could not be assigned any function, 20% of the genes were found to be responsible for signal transduction followed by 7% associated with intracellular trafficking, secretion and vesicular transport, suggesting importance of signal transduction processes in differentiating LGG and SGG. The number gene loci carrying nonsynonymous variation were estimated. Where the nonsynonymous SNPs were present in 8801 genes including 6010 non-TE genes, which as per eggNOG and KOG predictions were found to be involved in signal transduction mechanisms, post-translational modification/protein turnover, transcription and intracellular trafficking/secretion (Supplemental Figure 8A).

### Supplemental Tables

**Supplemental Table 1**: Grain parameters analysis of rice genotypes.

| | LGR | PB 1121 | Sonasal | Bindli |
|---|---|---|---|---|
| Grain length (mm) | 8.77 | 9.26 | 4.4 | 5.06 |
| Grain width (mm) | 2.72 | 2.18 | 2.32 | 2.57 |
| Grain L/W ratio | 3.23 | 4.25 | 1.9 | 1.97 |
| 1000 Grain weight dehusked (gm) | 29.2 | 20.7 | 9.8 | 14.5 |

L/W, length-to-width

**Supplemental Table 2:** Raw read and alignment summary of rice genotypes

| Feature/Genotype | LGR | PB 1121 | SONASAL | BINDLI |
|---|---|---|---|---|
| **Total raw reads** | 243,460,282 | 245,148,640 | 182,713,442 | 257,078,610 |
| **Total read aligned (%)** | 233,801,702 (96.04%) | 225,862,728 (92.15%) | 174,568,682 (95.6%) | 249,992,579 (97.26%) |
| **Average mapping quality (MAPQ)** | 40.34 | 40.76 | 42.28 | 44.58 |
| **Average insert size (bp)** | ~302 | ~276 | ~276 | ~285 |
| **Coverage (%)** | 91.41 | 91.96 | 93.44 | 94.31 |
| **Depth (X)** | 59.43 | 57.61 | 43.94 | 64.98 |

**Supplemental Table 3**: Distribution and filtering of identified sequence polymorphism with reference to rice genotype Nipponbare.

| **Chromosome** | **Raw** | | **Filtered** | |
|---|---|---|---|---|
| | **SNP** | **InDel** | **SNP** | **InDel** |
| Chr1 | 510274 | 73459 | 397486 | 63129 |
| Chr2 | 422125 | 57776 | 329787 | 50059 |
| Chr3 | 411193 | 59109 | 329517 | 51229 |
| Chr4 | 439352 | 49601 | 315581 | 42131 |
| Chr5 | 295470 | 39370 | 230274 | 34068 |
| Chr6 | 421029 | 50666 | 311958 | 43535 |
| Chr7 | 387570 | 44796 | 281705 | 38408 |
| Chr8 | 399321 | 47526 | 298898 | 40382 |
| Chr9 | 303438 | 34928 | 224564 | 30130 |
| Chr10 | 355074 | 39824 | 261946 | 33595 |
| Chr11 | 440848 | 49777 | 323103 | 42234 |
| Chr12 | 371069 | 41256 | 259298 | 34754 |
| Overall | 4756763 | 588088 | 3564117 | 503654 |

**Supplemental Table 4**: Abundance of sequence polymorphism identified among four rice genotypes with respect to the rice genotype Nipponbare.

| | **SNPs** | | | **InDels** | | |
|---|---|---|---|---|---|---|
| | Total | Homozygous | Heterozygous | Total | Homozygous | Heterozygous |
| **LGR** | 2332496 | 2049708 | 282788 | 312226 | 296218 | 16008 |
| **PB 1121** | 1837517 | 1608856 | 228661 | 249458 | 236311 | 13147 |
| **SONASAL** | 1215613 | 995545 | 220068 | 162924 | 149466 | 13458 |
| **BINDLI** | 1270617 | 1093340 | 177277 | 176094 | 166714 | 9380 |

**Supplemental Table 5**: List of primer used for the validation of identified SNPs among the four sequenced genotypes.

| Chromosome | Sequence | Start | End | Amplicon size |
|---|---|---|---|---|
| Chr1 | GCTATAGACATCGCTCTTTGTTGTTGC | 1357057 | 1357083 | 743 |
| | GGAAACACGGAGAAAGAGAAGATTGAG | 1357799 | 1357773 | |
| Chr2 | GGGTTGTTCTGAATTGAGGTTGG | 2935006 | 2935028 | 844 |
| | TTGTTCATTGTCTGGGAAAATACCTC | 2935849 | 2935824 | |
| Chr4 | GAGATCGCAGATCCAAAGGTTGTT | 1004165 | 1004188 | 730 |
| | CAACTTTACCAAATTCTCCGCGAT | 1004894 | 1004871 | |
| Chr6 | CCTGTTGAGTAAACAAAATTCTCCTC | 1348745 | 1348770 | 632 |
| | AGATTAGATGGAGTAGCTAATGCATG | 1349376 | 1349351 | |
| Chr8 | ACTGGGATCTGACTGGACTGTACAT | 2002562 | 2002586 | 781 |
| | TGCAGGATGATAATTGTATTTGCCC | 2003342 | 2003318 | |
| Chr10 | GATGTCGGTTCTGTCACCACT | 998092 | 998112 | 662 |
| | AAACATTGAGCAGCGCGGTA | 998753 | 998734 | |

**Supplemental Table 6**: Abundance of sequence polymorphisms identified in pair-wise rice genotypes.

| Genotype pair | Total | SNP | InDel |
|---|---|---|---|
| **LG/PB** | 1783948 | 1527437 | 256511 |
| **LG/SN** | 2142984 | 1844574 | 298410 |
| **LG/BN** | 2221783 | 1915382 | 306401 |
| **PB/SN** | 1262800 | 1076505 | 186295 |
| **PB/BN** | 1312230 | 1120711 | 191519 |
| **SN/BN** | 634306 | 530468 | 103838 |

BN, Bindli; LG, LGR; PB, PB 1121; SN, Sonasal

**Supplemental Table 7**: Distribution of SNP frequency per 100 kb over different chromosomes in pair-wise comparison.

| Chromosome | LG/PB | LG/SN | LG/BN | PB/SN | PB/BN | SN/BN | Average |
|---|---|---|---|---|---|---|---|
| Chr1 | 403.0 | 508.6 | 489.9 | 379.6 | 356.5 | 136.2 | 379.0 |
| Chr2 | 469.9 | 501.6 | 560.3 | 210.8 | 228.6 | 147.1 | 353.1 |
| Chr3 | 436.2 | 477.8 | 556.2 | 236.2 | 286.7 | 199.8 | 365.5 |
| Chr4 | 390.3 | 438.5 | 485.7 | 230.3 | 258.4 | 131.7 | 322.5 |
| Chr5 | 222.2 | 410.1 | 426.4 | 342.0 | 311.8 | 103.2 | 302.6 |
| Chr6 | 414.4 | 526.4 | 536.4 | 206.4 | 235.3 | 78.3 | 332.9 |
| Chr7 | 324.3 | 567.9 | 433.5 | 354.4 | 303.3 | 203.7 | 364.5 |
| Chr8 | 475.4 | 479.4 | 497.8 | 373.5 | 216.0 | 229.4 | 378.6 |
| Chr9 | 547.9 | 563.6 | 593.4 | 91.6 | 244.1 | 165.8 | 367.7 |
| Chr10 | 347.2 | 398.6 | 515.6 | 208.5 | 336.0 | 113.7 | 319.9 |
| Chr11 | 434.6 | 603.7 | 603.5 | 478.0 | 504.2 | 126.1 | 458.4 |
| Chr12 | 459.0 | 453.3 | 472.3 | 298.2 | 324.1 | 61.4 | 344.7 |
| Overall | 410.4 | 494.1 | 514.2 | 284.1 | 300.4 | 141.3 | 357.4 |

BN, Bindli; LG, LGR; PB, PB 1121; SN, Sonasal

**Supplemental Table 8**: Distribution of InDel frequency per 100 kb over different chromosomes in pair-wise comparison

| Chromosome | LG/PB | LG/SN | LG/BN | PB/SN | PB/BN | SN/BN | Average |
|---|---|---|---|---|---|---|---|
| Chr1 | 76.3 | 87.5 | 84.9 | 65.6 | 61.9 | 29.6 | 67.6 |
| Chr2 | 79.8 | 85.1 | 93.3 | 39.8 | 45.2 | 31.2 | 62.4 |
| Chr3 | 75.8 | 79.9 | 92.4 | 46.6 | 54.6 | 39.3 | 64.8 |
| Chr4 | 61.2 | 66.9 | 72.1 | 41.0 | 44.0 | 23.1 | 51.4 |
| Chr5 | 39.9 | 70.1 | 70.9 | 61.8 | 55.7 | 22.3 | 53.4 |
| Chr6 | 67.2 | 84.6 | 86.7 | 39.8 | 45.7 | 22.7 | 57.8 |
| Chr7 | 59.3 | 87.8 | 73.0 | 50.6 | 47.3 | 31.8 | 58.3 |
| Chr8 | 74.5 | 75.9 | 76.7 | 60.5 | 42.0 | 35.2 | 60.8 |
| Chr9 | 85.3 | 88.3 | 90.5 | 19.5 | 37.1 | 26.3 | 57.8 |
| Chr10 | 59.9 | 72.2 | 80.2 | 49.0 | 53.9 | 26.5 | 56.9 |
| Chr11 | 68.7 | 87.6 | 87.8 | 67.7 | 71.9 | 25.3 | 68.2 |
| Chr12 | 73.5 | 71.0 | 73.4 | 48.6 | 51.3 | 16.6 | 55.7 |
| Overall | 68.5 | 79.7 | 81.8 | 49.2 | 50.9 | 27.5 | 59.6 |

BN, Bindli; LG, LGR; PB, PB 1121; SN, Sonasal

**Supplemental Table 9**: Differential polymorphism density regions identified in pair-wise analysis of four rice genotypes.

| | SNPs per 100 kb | | | | | InDels per 100 kb | | |
|---|---|---|---|---|---|---|---|---|
| **Genotype pair** | **Low Density region (≤5)** | **High Density region (≥500)** | **SNP Hotspot (≥1000)** | **No SNP regions** | | **Low Density region (≤2)** | **High Density region (≥50)** | **No InDel region** |
| LG/PB | 203 | 1517 | 105 | 14 | | 140 | 2330 | 31 |
| LG/SN | 34 | 1966 | 141 | 1 | | 62 | 2725 | 12 |
| LG/BN | 25 | 2085 | 140 | 2 | | 60 | 2774 | 11 |
| PB/SN | 708 | 1092 | 67 | 185 | | 390 | 1602 | 95 |
| PB/BN | 467 | 1152 | 84 | 33 | | 238 | 1640 | 42 |
| SN/BN | 1005 | 471 | 36 | 65 | | 372 | 763 | 57 |

BN, Bindli; LG, LGR; PB, PB 1121; SN, Sonasal

**Supplemental Table 12**: Large effect SNPs and InDels detected between long grain genotypes (LGG) and short grain genotypes (SGG).

| Variant class | SNPs | InDels |
|---|---|---|
| 5_splice_site | 1449 | 229 |
| 3_splice_site | 1585 | 281 |
| Nonsense | 1232 | 0 |
| Startloss | 109 | 0 |
| Stoploss | 217 | 0 |
| Frameshift | 0 | 1364 |
| Total | 4592 | 1874 |

**Supplemental Table 13**: Distribution and annotation of sequence variations identified in grain trait-related genes among four rice genotypes and LGG vs SGG.

| | #Gene loci | SNP | InDel | Silent | Missense | Nonsense | StartLoss | StopLoss | Frameshift |
|---|---|---|---|---|---|---|---|---|---|
| Overall | 49 | 1385 | 257 | 274 | 316 | 3 | 1 | 2 | 5 |
| LGG/SGG | 34 | 542 | 60 | 147 | 173 | 2 | 0 | 1 | 0 |

**Supplemental Table 14:** Details of genome wide marker trait associations for grain length in rice.

| SNP-ID | Chromosome | Position (IRGSP 1.0) | SGG allele | LGG allele | p Value | Allele effect | $R^2$ | Annotation |
|---|---|---|---|---|---|---|---|---|
| *SNP-3.16732086 | 3 | 16733441 | C | A | 1.49E-48 | 0.41 | 0.600 | *GS3* |
| *SNP-5.5371749 | 5 | 5371772 | G | A | 7.92E-10 | -0.20 | 0.508 | *GW5* |
| *SNP-11.1644056 | 11 | 1645056 | T | G | 1.62E-05 | 0.61 | 0.497 | LOC_Os11g04060 |
| SNP-6.8401944 | 6 | 8402944 | A | C | 5.90E-05 | -0.80 | 0.496 | Intergenic (LOC_Os06g14800-LOC_Os06g14810) |
| SNP-6.24237618 | 6 | 24238616 | C | T | 0.00032205 | 0.42 | 0.494 | Intergenic (LOC_Os06g40650-LOC_Os06g14810) |

SGG (Short Grain Genotype) - Sonasal and Bindli; LGG (Long Grain Genotype) - LGR and PB 1121

*Validated in the bi-parental mapping population

**Supplemental Table 15**: Analysis and distribution of reads alignment rate over the desert region among four rice genotypes.

| | LGR | PB 1121 | Sonasal | Bindli |
|---|---|---|---|---|
| Reads mapped per Mb genome | 580307 | 565563 | 431570 | 637679 |
| Genome-wide reads count (Depth) | 58.0 | 56.5 | 43.1 | 63.7 |
| Reads mapped to desert region (Chr5: 8-14 Mb) | 4887934 | 4461145 | 3457518 | 4485587 |
| Reads count over desert region (Depth) | 81.4 | 74.3 | 57.6 | 74.7 |

**Supplemental Table 16**: Distribution of SNP frequency per 100 kb over centromeric region of 12 rice chromosomes.

| Chromosome | Coordinates (Mb) | LG/PB | LG/SN | LG/BN | PB/SN | PB/BN | SN/BN | Average |
|---|---|---|---|---|---|---|---|---|
| Chr1 | 16.3-17.1 | 104.5 | 563.0 | 563.8 | 538.8 | 540.4 | 21.3 | 388.6 |
| Chr2 | 13.3-13.9 | 456.7 | 457.2 | 467.8 | 0.2 | 3.3 | 3.5 | 231.4 |
| Chr3 | 19.2-19.6 | 664.5 | 659.8 | 666.0 | 9.8 | 11.0 | 4.8 | 336.0 |
| Chr4 | 9.5-9.9 | 569.5 | 557.8 | 537.8 | 9.0 | 8.0 | 9.0 | 281.8 |
| Chr5 | 12.2-12.6 | 13.5 | 19.3 | 20.5 | 15.0 | 15.3 | 16.8 | 16.7 |
| Chr6 | 15.1-15.4 | 847.0 | 856.3 | 859.3 | 4.3 | 6.3 | 3.0 | 429.4 |
| Chr7 | 11.8-12.4 | 312.3 | 634.2 | 14.0 | 556.7 | 317.2 | 648.8 | 413.9 |
| Chr8 | 12.3-13.5 | 486.3 | 332.3 | 495.7 | 469.3 | 6.7 | 475.5 | 377.6 |
| Chr9 | 2.5-3.0 | 529.4 | 534.4 | 534.4 | 4.6 | 9.8 | 8.6 | 270.2 |
| Chr10 | 7.6-8.8 | 378.1 | 365.1 | 385.0 | 45.1 | 81.6 | 4.3 | 209.8 |
| Chr11 | 11.9-12.1 | 352.0 | 499.0 | 498.5 | 357.0 | 354.5 | 22.5 | 347.3 |
| Chr12 | 11.6-12.2 | 286.3 | 279.2 | 287.7 | 198.3 | 206.3 | 8.5 | 211.1 |

BN, Bindli; LG, LGR; PB, PB 1121; SN, Sonasal

**Supplemental Table 17**: Distribution of InDel frequency per 100 kb over centromeric region of 12 rice chromosomes.

| Chromosome | Coordinates (Mb) | LG/PB | LG/SN | LG/BN | PB/SN | PB/BN | SN/BN | Average |
|---|---|---|---|---|---|---|---|---|
| Chr1 | 16.3-17.1 | 17.6 | 47.4 | 48.0 | 42.6 | 43.3 | 3.5 | 33.7 |
| Chr2 | 13.3-13.9 | 45.3 | 46.2 | 45.2 | 0.5 | 1.7 | 2.0 | 23.5 |
| Chr3 | 19.2-19.6 | 39.8 | 40.8 | 39.5 | 2.5 | 2.3 | 1.8 | 21.1 |
| Chr4 | 9.5-9.9 | 38.3 | 37.3 | 35.8 | 5.3 | 7.5 | 6.8 | 21.8 |
| Chr5 | 12.2-12.6 | 1.5 | 2.3 | 3.3 | 2.3 | 3.3 | 3.5 | 2.7 |
| Chr6 | 15.1-15.4 | 66.3 | 67.3 | 67.0 | 2.7 | 2.7 | 3.0 | 34.8 |
| Chr7 | 11.8-12.4 | 20.8 | 44.8 | 4.5 | 40.5 | 20.8 | 44.8 | 29.4 |
| Chr8 | 12.3-13.5 | 37.9 | 31.3 | 36.6 | 39.3 | 3.4 | 38.6 | 31.2 |
| Chr9 | 2.5-3.0 | 42.4 | 41.4 | 43.0 | 1.4 | 1.8 | 3.4 | 22.2 |
| Chr10 | 7.6-8.8 | 34.8 | 33.3 | 35.5 | 8.6 | 10.9 | 3.3 | 21.1 |
| Chr11 | 11.9-12.1 | 27.5 | 29.0 | 29.5 | 31.0 | 31.0 | 6.5 | 25.8 |
| Chr12 | 11.6-12.2 | 25.8 | 26.3 | 26.7 | 20.8 | 20.7 | 3.3 | 20.6 |

BN, Bindli; LG, LGR; PB, PB 1121; SN, Sonasal

**Supplemental Table 18:** Distribution of nucleotide diversity (100kb window) over chromosome 5 among different rice population.

| | Mean Nucleotide Diversity ( π) | | | |
|---|---|---|---|---|
| Genotypes | Chromosome 5 | 8-14 Mb | 9-14 Mb | 10-14 Mb |
| *japonica* | 0.00168 | 0.000620307 | 0.000484748 | 0.000480832 |
| *indica* | 0.00188288 | 0.00185248 | 0.00145144 | 0.00130437 |
| *aus* | 0.00227407 | 0.00226275 | 0.00203694 | 0.00191577 |
| Our_four | 0.0028 | 0.000238923 | 0.000150479 | 0.000150688 |
| All_3004 | 0.0027 | 0.00176355 | 0.00136891 | 0.00126627 |

**Supplemental Table 19:** Distribution of genetic differentiation (100kb window) over chromosome 5 among different rice population.

| | Mean $F_{ST}$ | | | |
|---|---|---|---|---|
| **Genotypes** | **Chromosome 5** | **8-14 Mb** | **9-14 Mb** | **10-14 Mb** |
| *japonica*_vs_*indica* | 0.4067197 | 0.121689 | 0.0857836 | 0.0811114 |
| *japonica*_vs_*aus* | 0.5367887 | 0.631003 | 0.636291 | 0.645917 |
| *indica*_vs_*aus* | 0.3436793 | 0.342931 | 0.364444 | 0.388686 |

**Supplemental Table 20:** Distribution of Tajima's D estimation (100kb window) over chromosome 5 among different rice population.

| | **Mean Tajima's D** | | | |
|---|---|---|---|---|
| **Genotypes** | **Chromosome 5** | **8-14 Mb** | **9-14 Mb** | **10-14 Mb** |
| *japonica* | 0.112514 | -1.70457 | -1.83241 | -1.8177 |
| *indica* | 1.23784 | 1.19813 | 0.781445 | 0.616101 |
| *aus* | 2.08648 | 3.54369 | 3.80695 | 4.07429 |
| All_3004 | 2.30587 | 1.0662 | 0.643976 | 0.543008 |

**Supplemental Table 21:** Distribution of Linkage Disequilibrium estimation (100kb window) over chromosome 5 among different rice population.

| | **Mean Linkage Disequilibrium** | |
|---|---|---|
| **Genotypes** | **Chromosome 5** | **Desert region (8-14 Mb)** |
| *japonica* | 0.213276 | 0.2255637 |
| *indica* | 0.1573135 | 0.2204522 |
| *aus* | 0.1596137 | 0.203914 |
| All_3004 | 0.2028565 | 0.2256317 |

**Supplemental Table 22**: Distribution of allele frequency estimation (100kb window) over chromosome 5 among different rice population.

| | **Mean Allele frequency** | |
|---|---|---|
| **Genotypes** | **Chromosome 5** | **Desert region (8-14 Mb)** |
| *japonica* (484) | 0.98 | 0.99 |
| *indica* (520) | 0.83 | 0.94 |
| *aus* (30) | 0.82 | 0.84 |
| Wild, O. rufipogon (446) | 0.83 | 0.84 |

Values in parenthesis denote number of accessions.

**Supplemental Table 23**: Distribution of 3004 rice genotypes based on outlier SNP analysis in chromosome 5 desert region (8-14 Mb) as per varietal group.

| Rice varietal group | Total genotypes | Number of outlier genotypes | % genotypes in outlier range | Avg. SNP count |
|---|---|---|---|---|
| *indica* | 1743 | 311 | 17.8 | 7406 |
| Tropical *japonica* | 388 | 12 | 3.1 | 3251 |
| Temperate *japonica* | 320 | 9 | 2.8 | 3745 |
| *aus/boro* | 215 | 136 | 63.3 | 16387 |
| Intermediate/admixed | 135 | 8 | 5.9 | 3734 |
| *japonica* | 132 | 4 | 3.0 | 3256 |
| Aromatic (basmati/sadri) | 71 | 5 | 7.4 | 3503 |

### Supplemental Figures

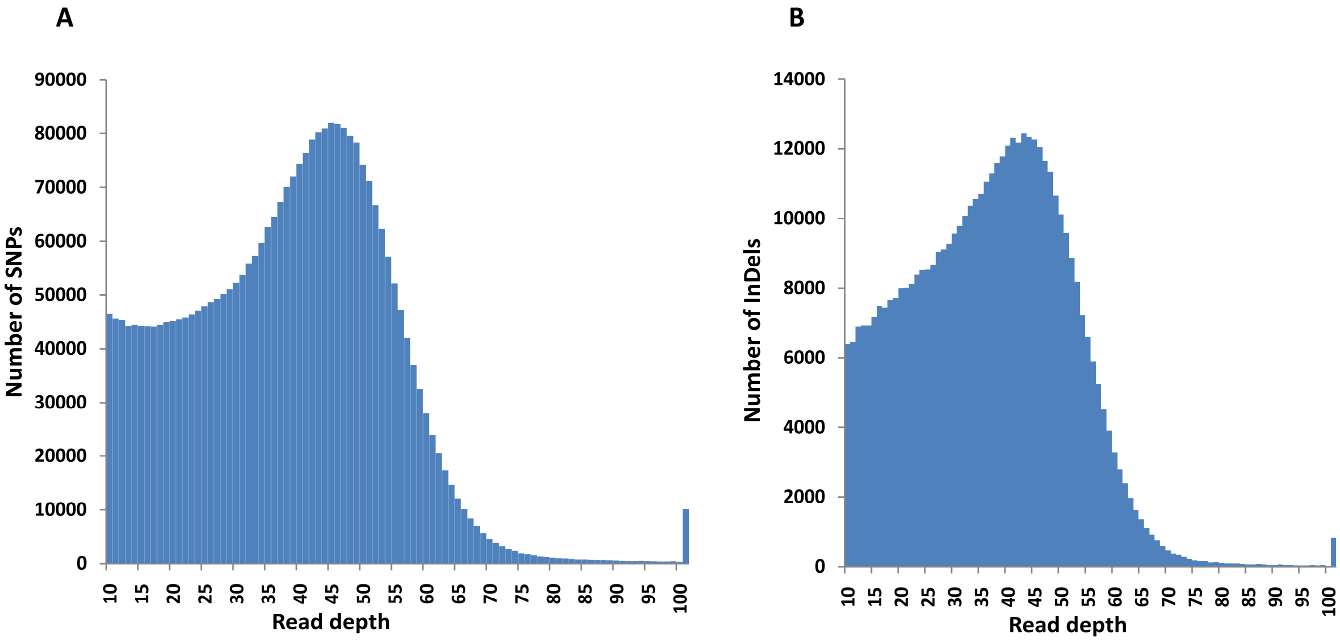


**Supplemental Figure 1**: **Read depth distribution of identified SNPs and InDels.** The number of SNPs (**A**) and InDels (**B**) identified with different read depths are shown.

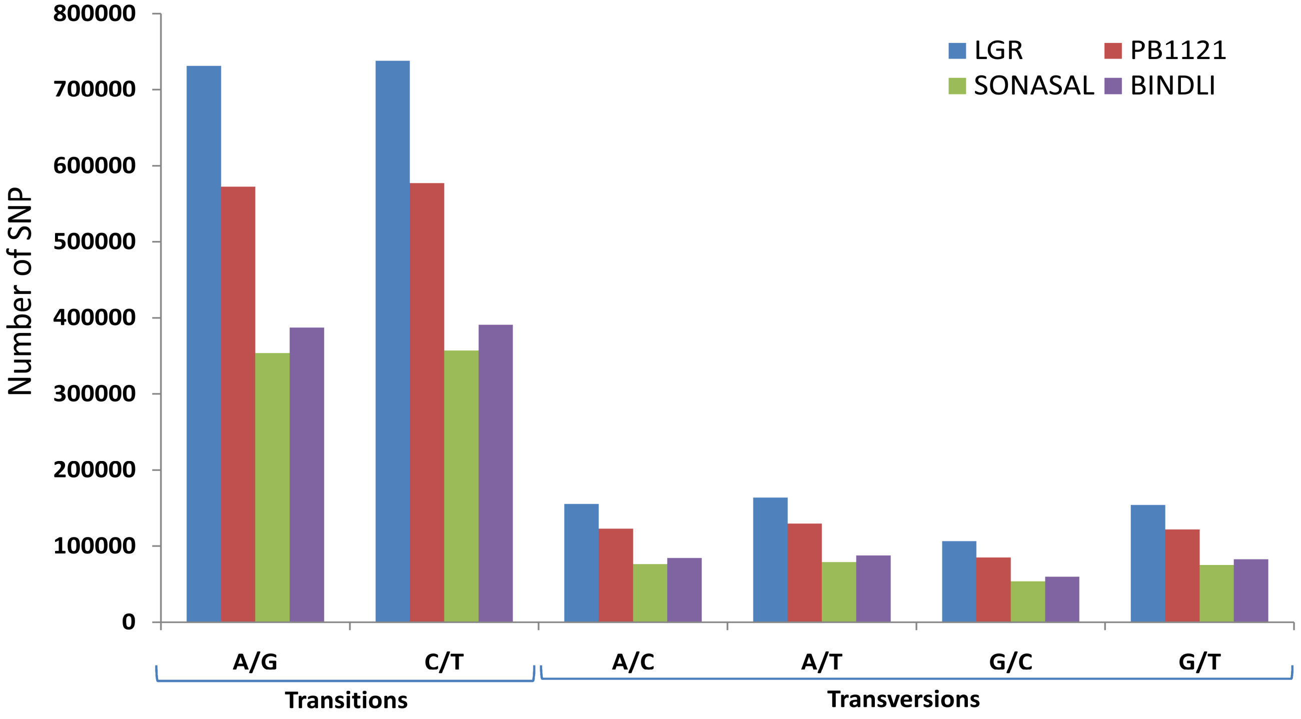


**Supplemental Figure 2**: Distribution and number of different substitution types of single nucleotide polymorphism identified in four rice genotypes with respect to Nipponbare.

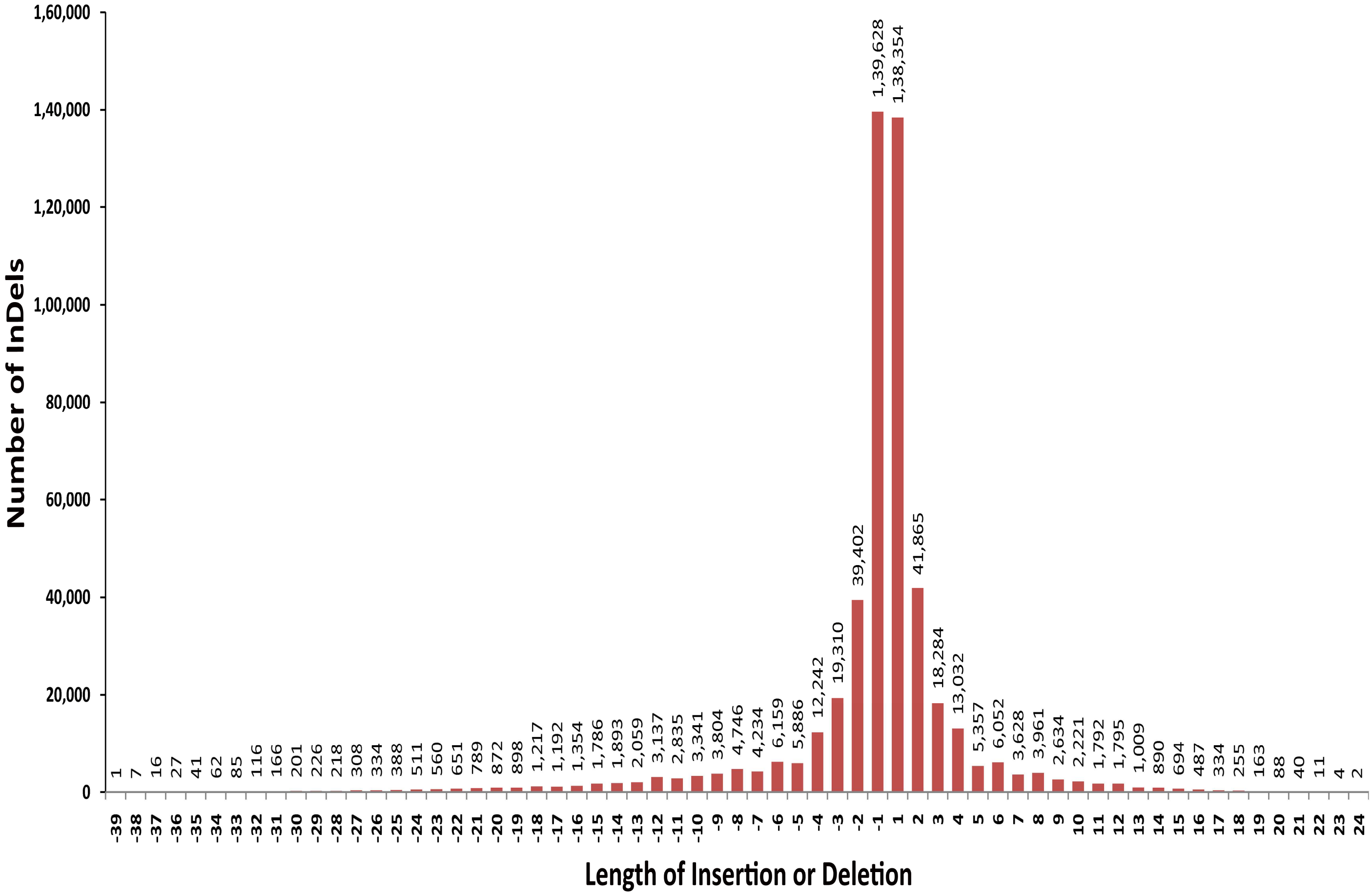


**Supplemental Figure 3**: **Length distribution of InDels.** Number of insertions and deletions (*y*-axis) of various lengths (*x*-axis, in bp) identified across the genomes of all four rice genotypes with reference to Nipponbare. On x-axis, - denotes deletion whereas + is insertion.

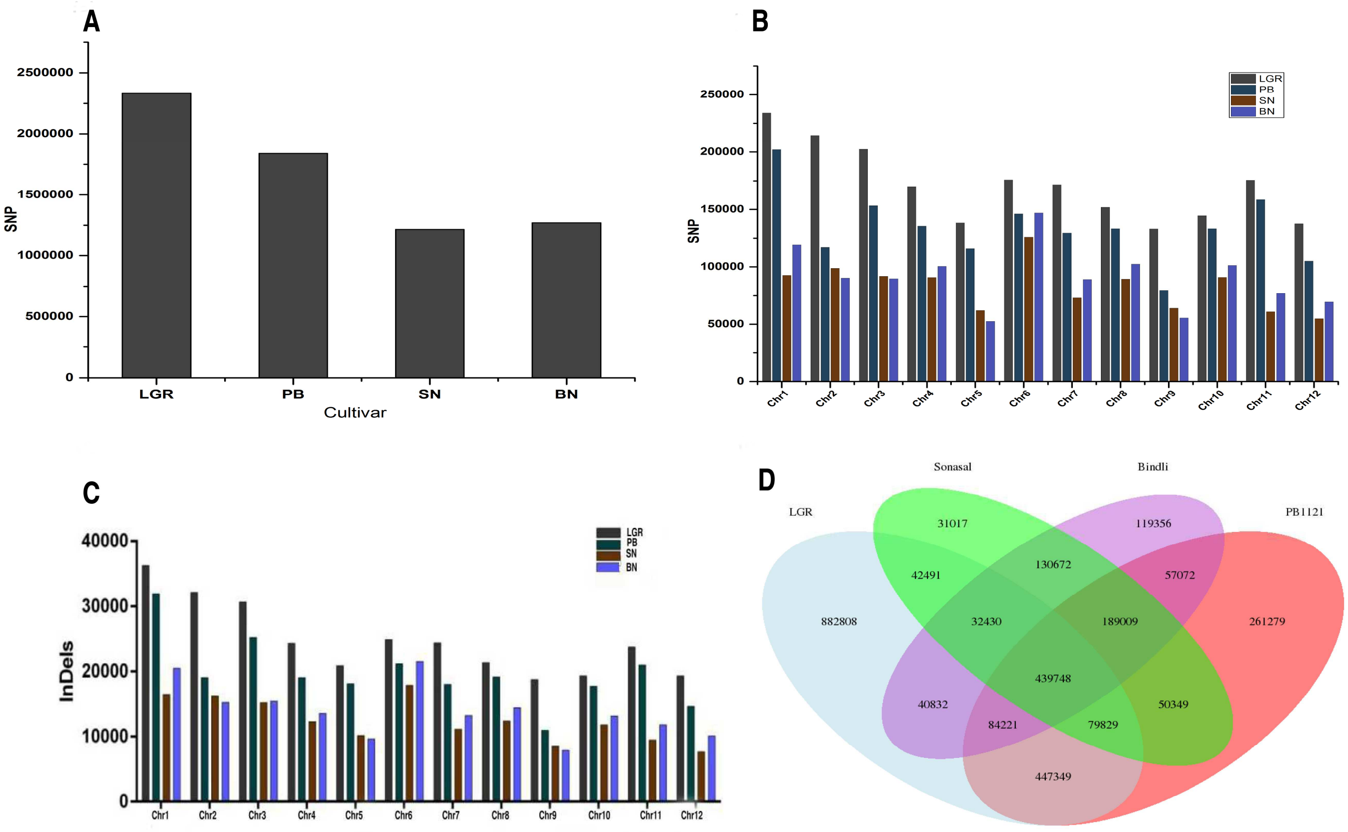


**Supplemental Figure 4**: **Abundance of SNPs among four rice genotypes with reference to Nipponbare.** (**A**) Total number of SNPs. (**B**) Distribution of homozygous SNPs over different chromosomes. (**C**) Distribution of homozygous InDels over different chromosomes. (**D**) Venn diagram representation for shared homozygous single nucleotide polymorphisms identified with respect to Nipponbare among the four rice genotypes. PB, PB 1121; SN, Sonasal; BN, Bindli

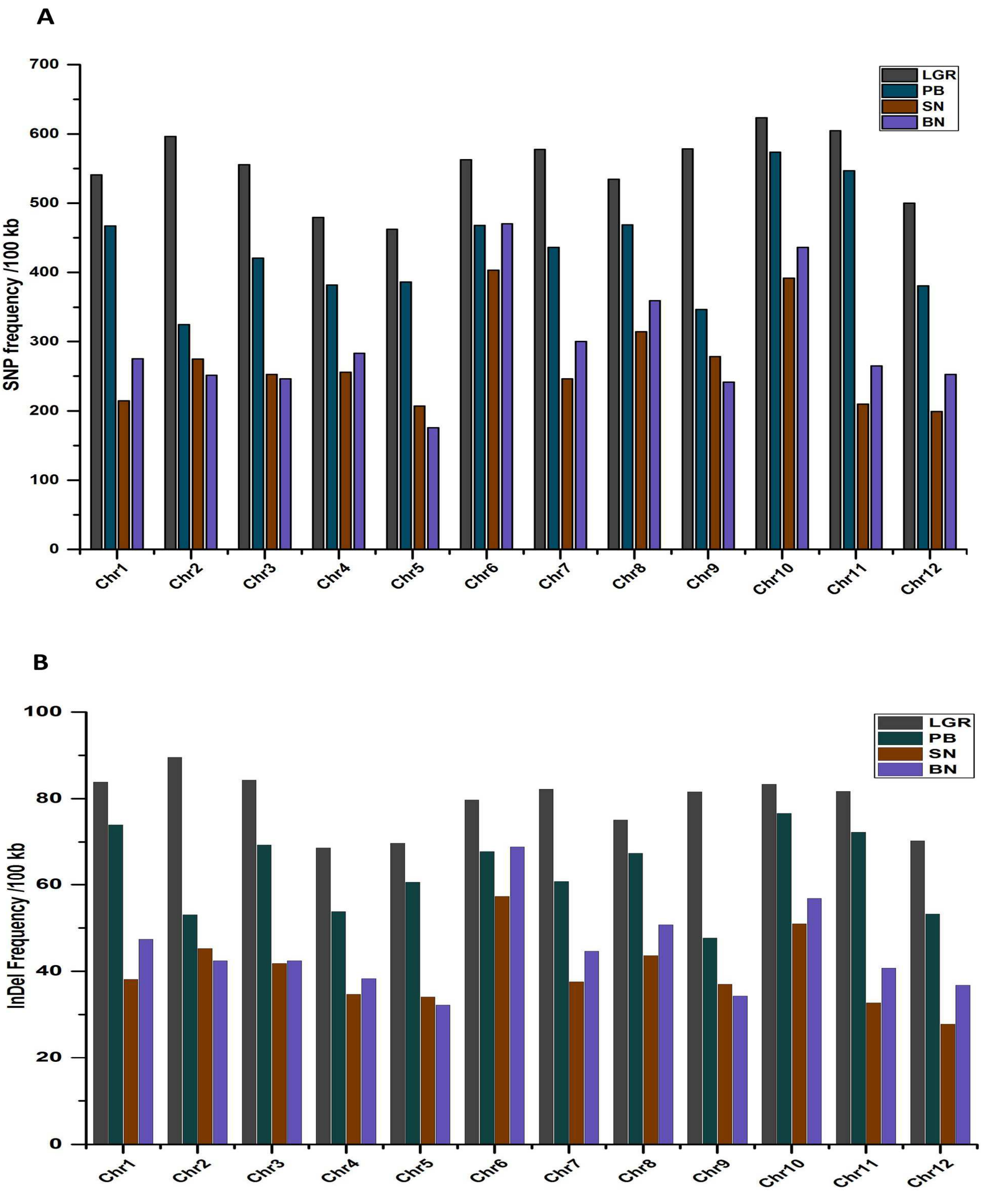


**Supplemental Figure 5**: **Polymorphism frequency distribution.** Chromosome wise per 100 kb SNPs (**A**) and InDels (**B**) among four rice genotypes with reference to Nipponbare. PB, PB 1121; SN, Sonasal; BN, Bindli

A

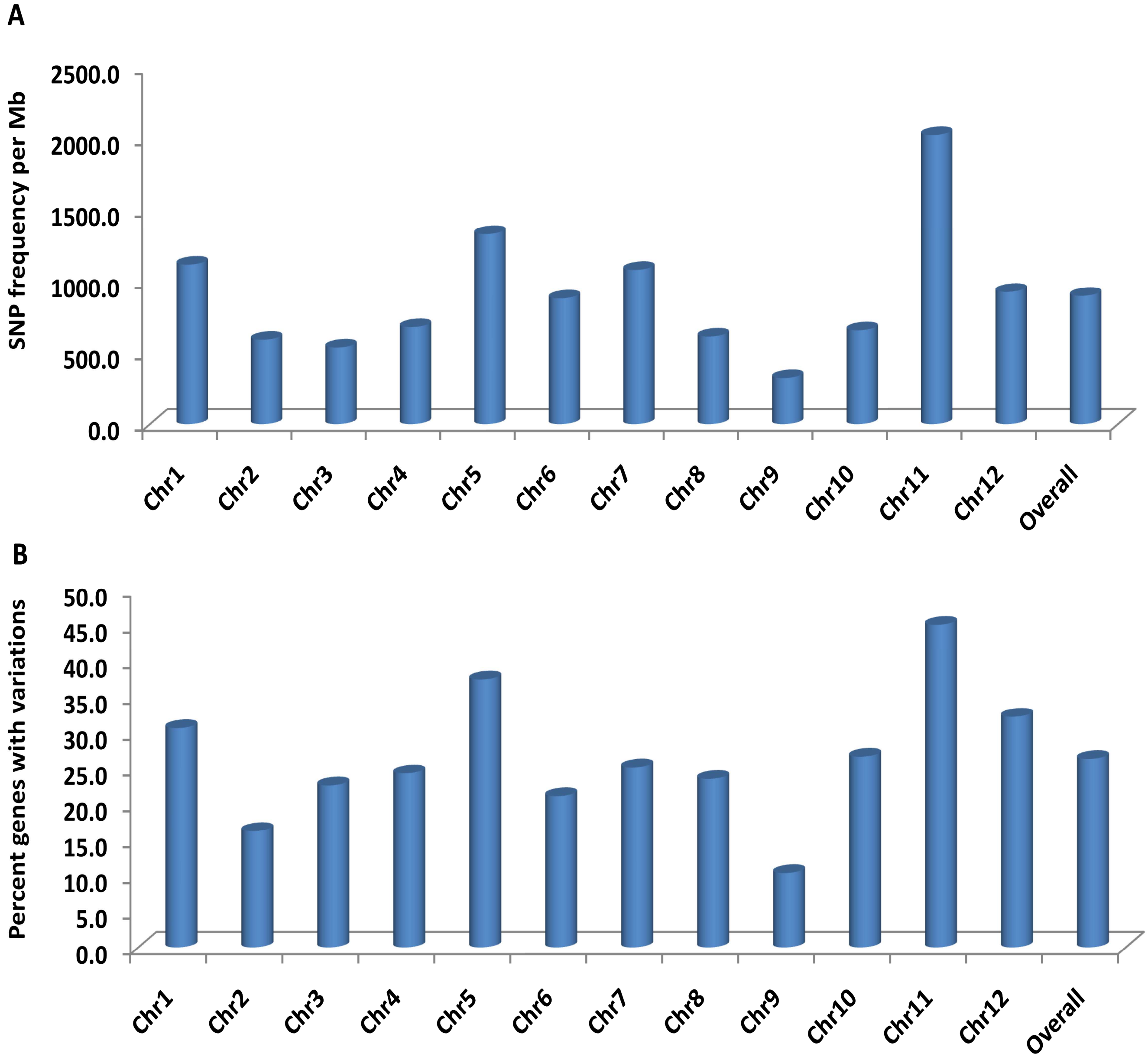


**Supplemental Figure 6**: **Distribution of SNP density on genes across the chromosomes in LGG and SGG comparison**. (**A**) Density of SNP per Mb. (**B**) Percentage of genes carrying the SNPs on all the twelve chromosome. LGG, Long grain genotype; SGG, Short grain genotype.

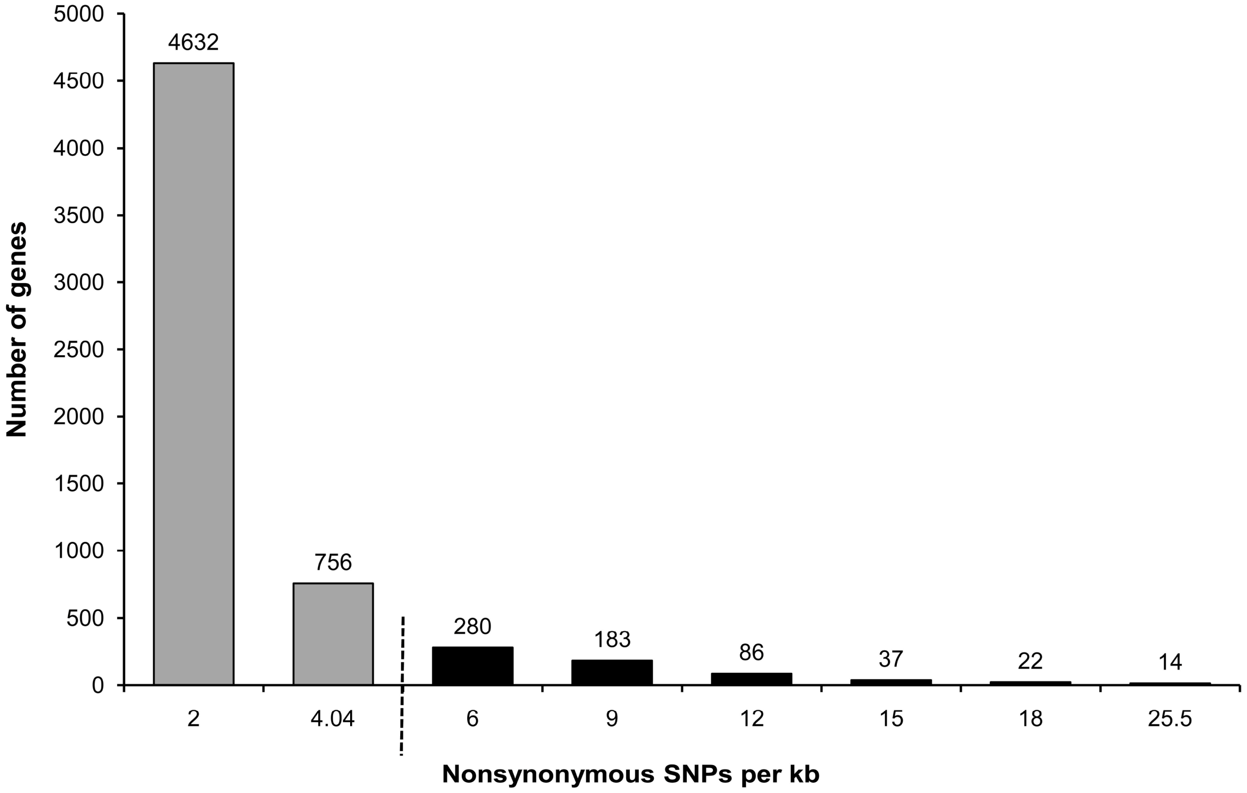


**Supplemental Figure 7**: **Distribution and non-uniformity in abundance of nonsynonymous SNPs per kb in genes identified between LGG and SGG.** Fraction of genes with expected frequency of nonsynonymous SNPs per kb is depicted with grey bars and the set of genes with outlier values showing more than expected number of nonsynonymous SNPs per kb is represented with black bars. The cut-off value for outliers is shown with dotted line.

A

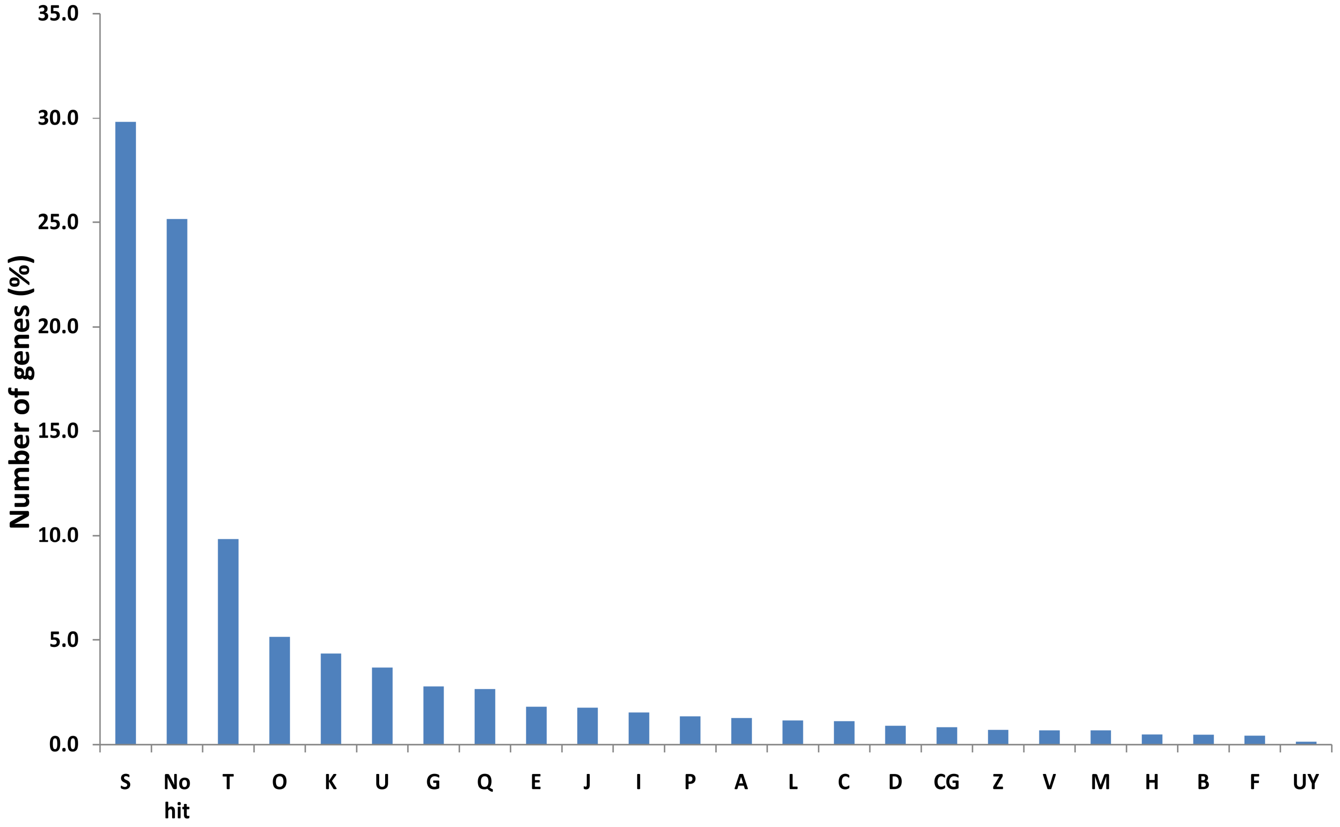
35.0
30.0
25.0
20.0
15.0
10.0
5.0
0.0
Number of genes (%)
S
No hit
T
O
K
U
G
Q
E
J
I
P
A
L
C
D
CG
Z
V
M
H
B
F
UY

B

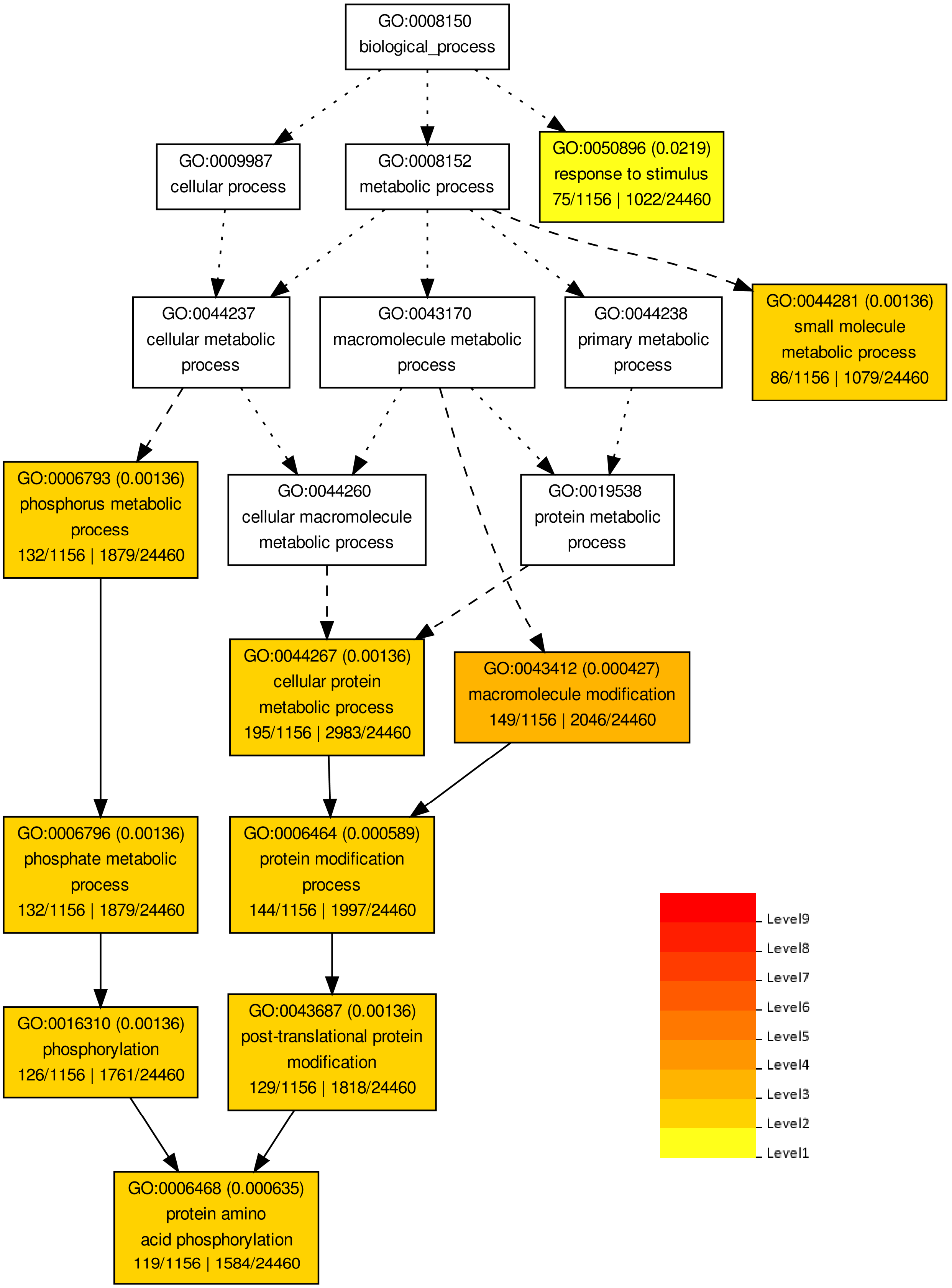
GO:0008150
biological_process
GO:0009987
cellular process
GO:0008152
metabolic process
GO:0050896 (0.0219)
response to stimulus
75/1156 | 1022/24460
GO:0044237
cellular metabolic process
GO:0043170
macromolecule metabolic process
GO:0044238
primary metabolic process
GO:0044281 (0.00136)
small molecule metabolic process
86/1156 | 1079/24460
GO:0006793 (0.00136)
phosphorus metabolic process
132/1156 | 1879/24460
GO:0044260
cellular macromolecule metabolic process
GO:0019538
protein metabolic process
GO:0044267 (0.00136)
cellular protein metabolic process
195/1156 | 2983/24460
GO:0043412 (0.000427)
macromolecule modification
149/1156 | 2046/24460
GO:0006796 (0.00136)
phosphate metabolic process
132/1156 | 1879/24460
GO:0006464 (0.000589)
protein modification process
144/1156 | 1997/24460
GO:0016310 (0.00136)
phosphorylation
126/1156 | 1761/24460
GO:0043687 (0.00136)
post-translational protein modification
129/1156 | 1818/24460
GO:0006468 (0.000635)
protein amino acid phosphorylation
119/1156 | 1584/24460
Level9
Level8
Level7
Level6
Level5
Level4
Level3
Level2
Level1

C

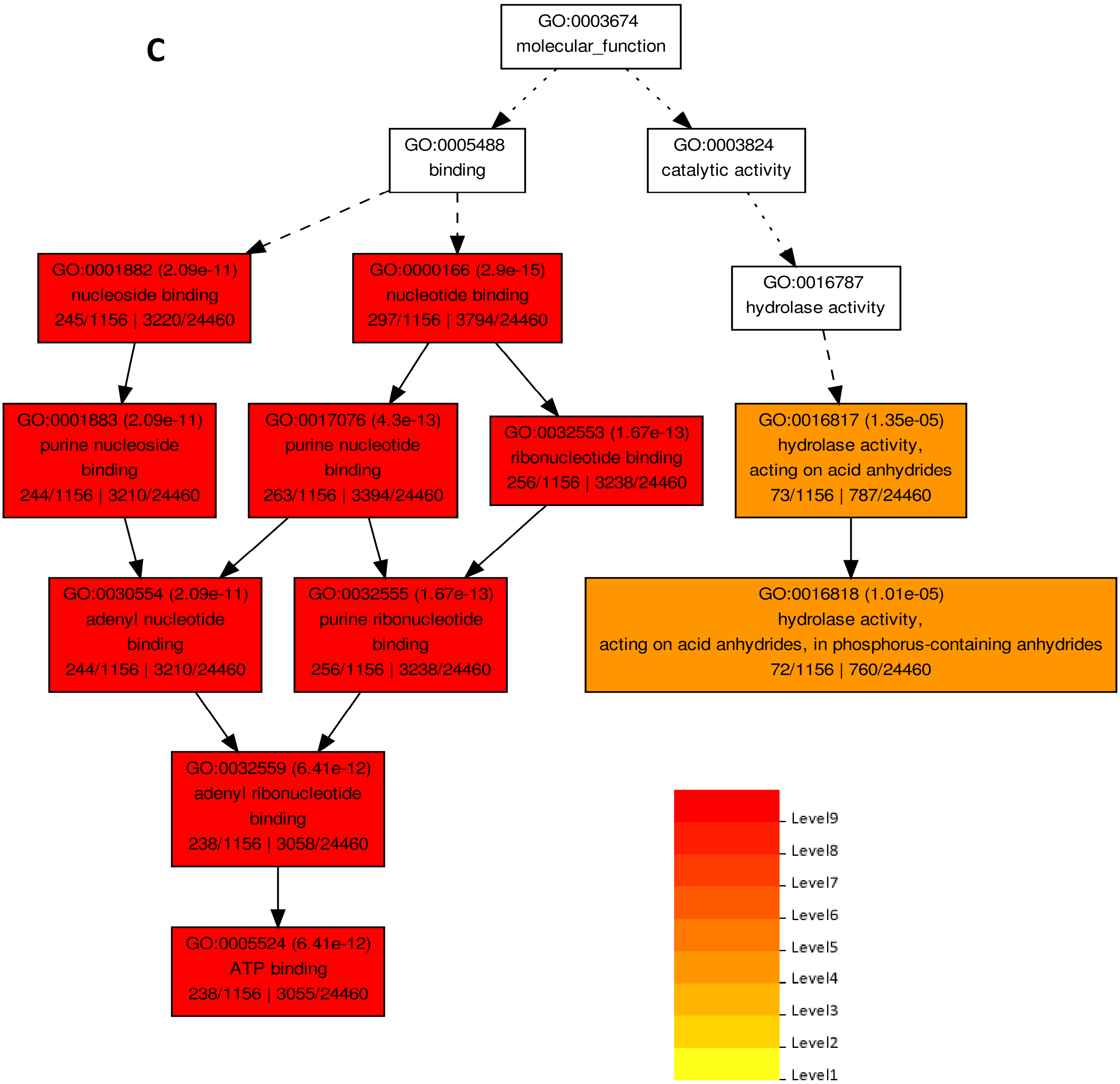


**Supplemental Figure 8**: **Functional categorization of genes identified with nonsynonymous/large-effect SNPs and InDels in LGG and SGG comparison.** (**A**) Distribution of the eukaryotic orthologous group (KOG) classes in the genes identified with nonsynonymous SNPs between LGG and SGG. A; RNA processing and modification, B; Chromatin structure and dynamics, C; Energy production and conversion, D; Cell cycle control, cell division and chromosome partitioning, E; Amino acid transport and metabolism, F; Nucleotide transport and metabolism, G; Carbohydrate transport and metabolism, H; Coenzyme transport and metabolism, I; Lipid transport and metabolism, J; Translation, ribosomal structure and biogenesis, K; Transcription, L; Replication, recombination and repair, M; Cell wall/membrane/envelope biogenesis, N; Cell motility, O; Posttranslational modification, protein turnover, chaperones, P; Inorganic ion transport and metabolism, Q; Secondary metabolites biosynthesis, transport and catabolism, R; General function prediction only, S; Function unknown, T; Signal transduction mechanisms, U; Intracellular trafficking, secretion, and vesicular transport, V; Defense mechanisms, W; Extracellular structures, Y; Nuclear structure, Z; Cytoskeleton, CG; UDP-Glycosyltransferase, UY; nuclear pore complex protein. Gene ontology term (**B**) Biological process, (**C**) Molecular process enrichment in the genes detected with large-effect SNPs/InDels in LGG/SGG. Colour shading is according to the significance level (white box– no significant difference).

.

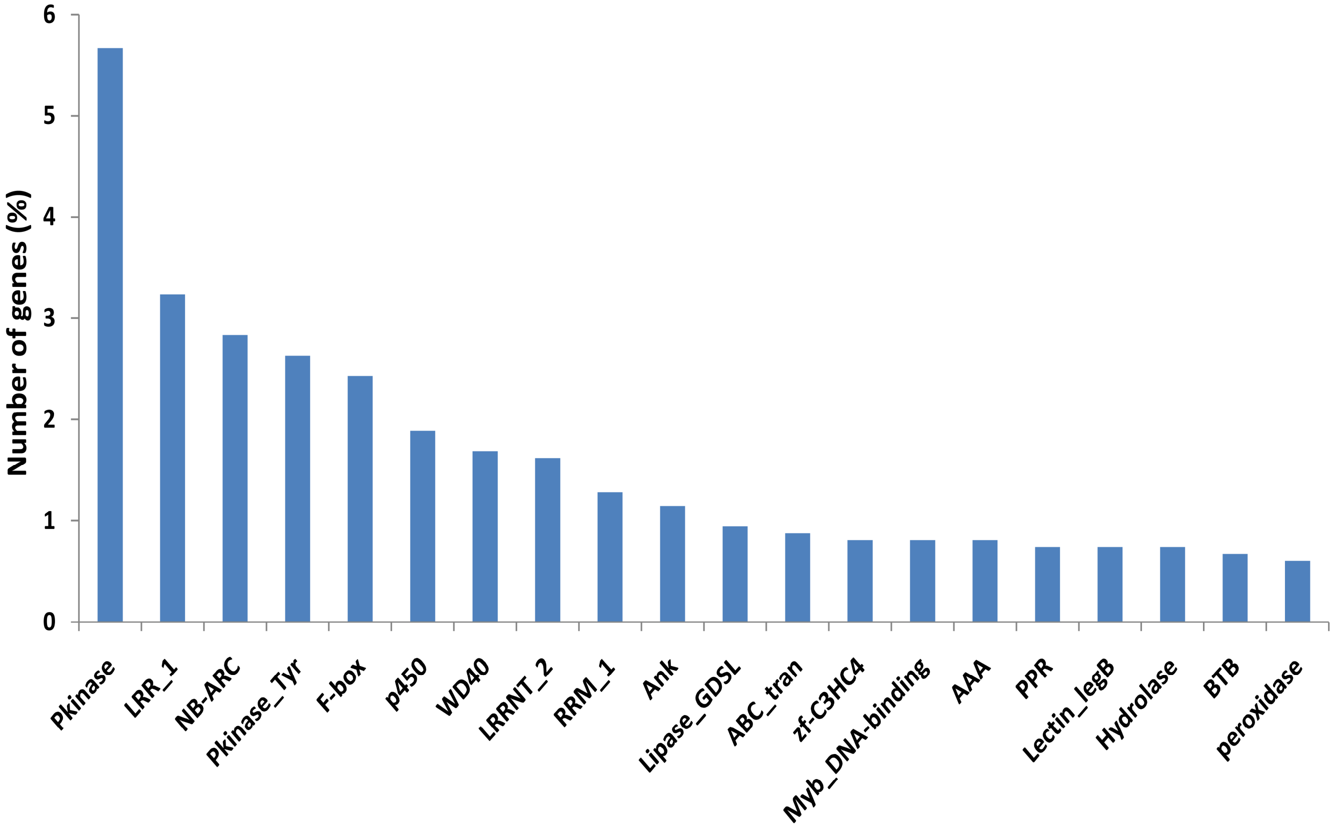


**Supplemental Figure 9**: Frequency of top 20 Pfam domains represented in the genes containing large-effect SNPs and/or InDels in LGG and SGG comparison.

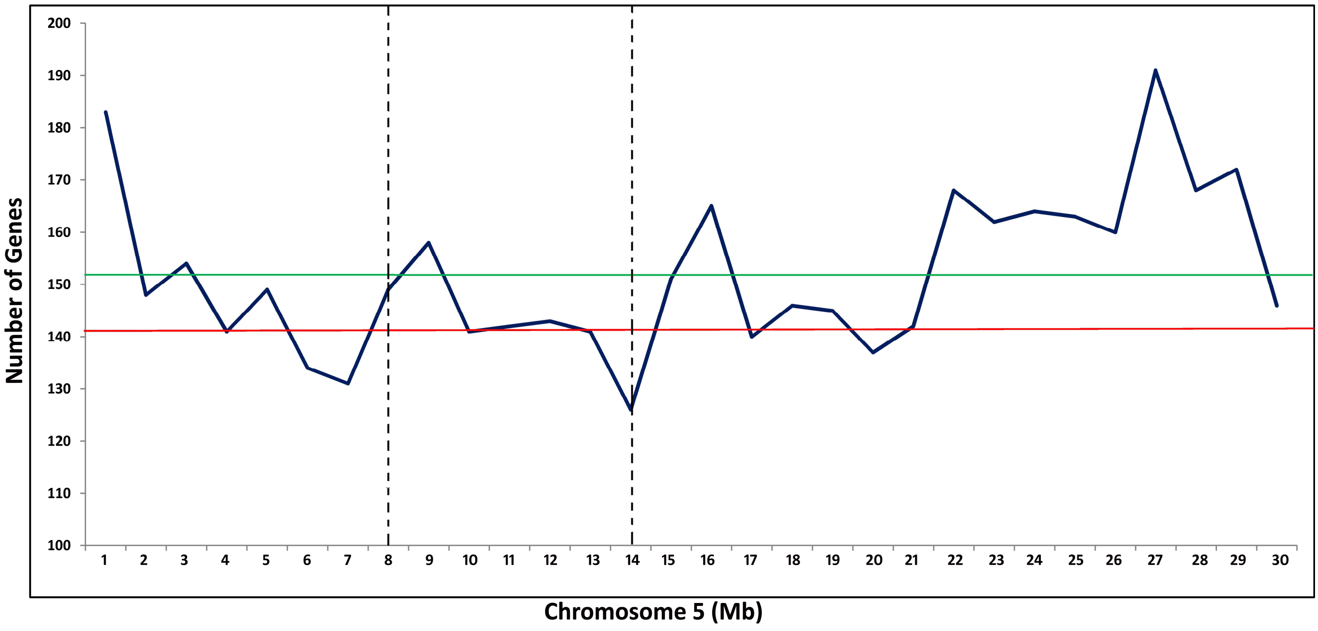


**Supplemental Figure 10: Distribution of gene density per Mb over the chromosome 5**. Red and green line represents the mean gene density (per 100 kb) for desert (8-14 Mb) region and full chromosome, respectively. Vertical dashed line marks the boundary of desert region.

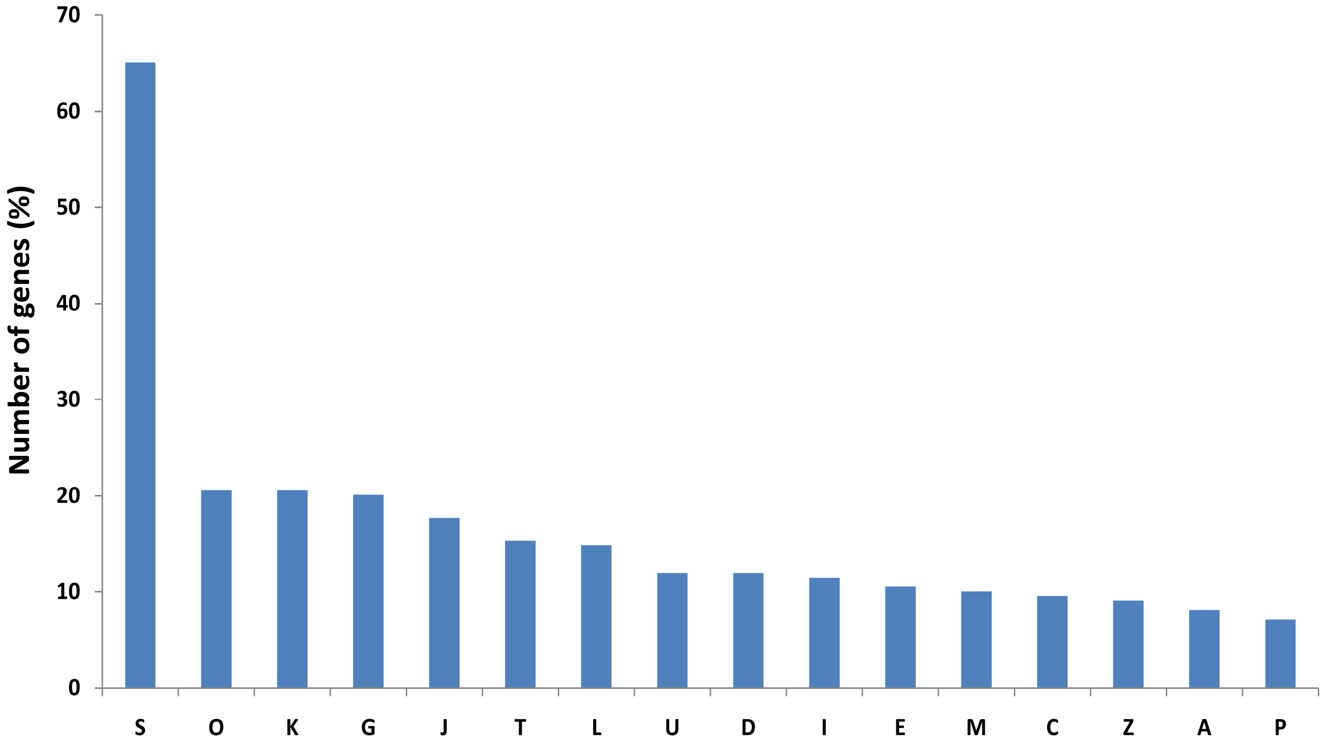


**Supplemental Figure 11**: **Functional categorization and distribution of the eukaryotic orthologous group (KOG) classes of genes residing in chromosome 5 desert region**. A; RNA processing and modification, C; Energy production and conversion, D; Cell cycle control, cell division and chromosome partitioning, E; Amino acid transport and metabolism, G; Carbohydrate transport and metabolism, I; Lipid transport and metabolism, J; Translation, ribosomal structure and biogenesis, K; Transcription, L; Replication, recombination and repair, M; Cell wall/membrane/envelope biogenesis, O; Posttranslational modification, protein turnover, chaperones, P; Inorganic ion transport and metabolism, S; Function unknown, T; Signal transduction mechanisms, U; Intracellular trafficking, secretion, and vesicular transport, Z; Cytoskeleton.

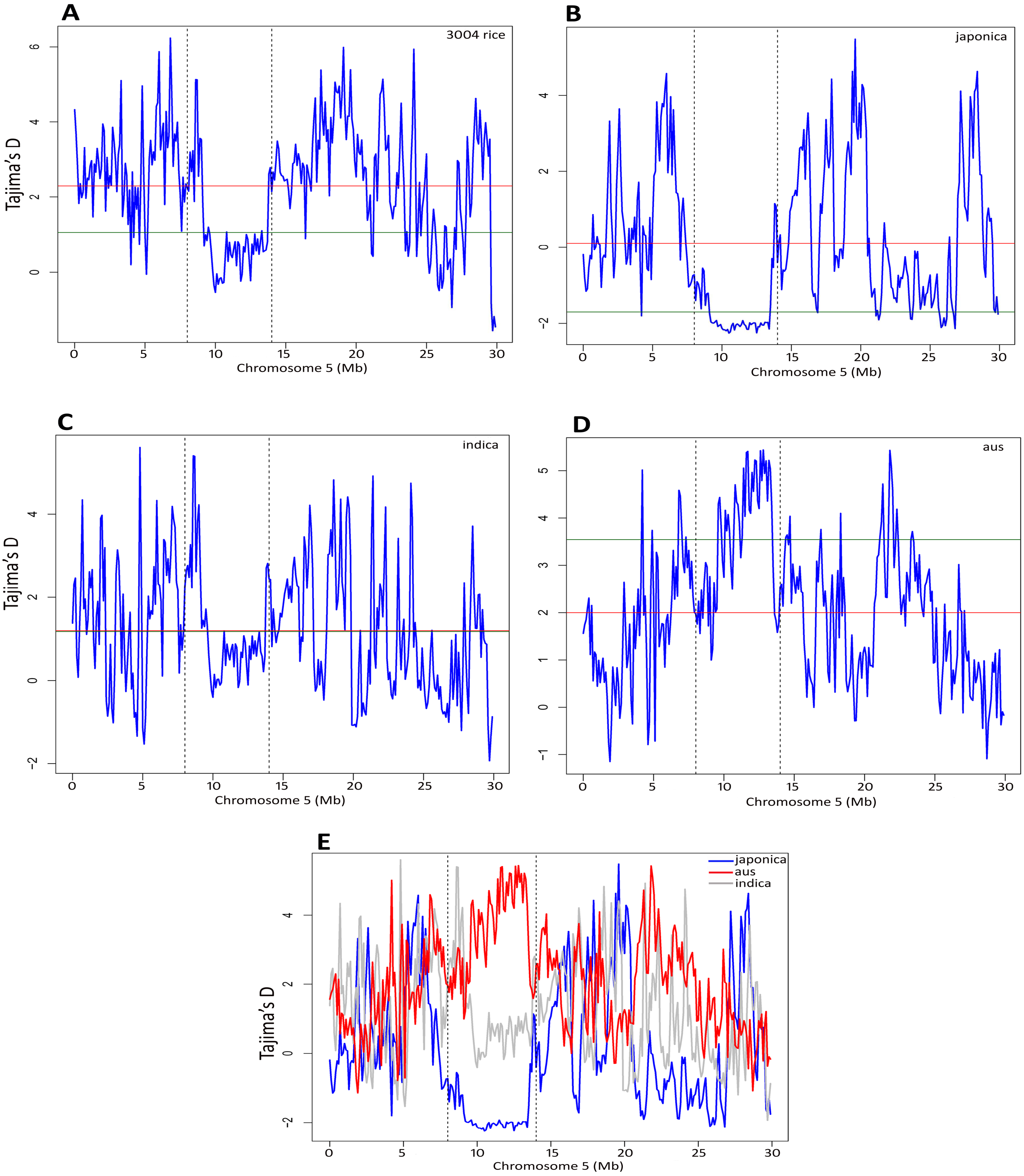


**Supplemental Figure 12**: **Analysis and distribution of Tajima's D estimation per 100 kb of chromosome 5 for different rice groups. (A)** Tajima's D distribution for all 3004 accessions**. (B)** Tajima's D distribution for all *japonica* (840) accessions**. (C)** Tajima's D distribution for all *indica* (1743) accessions**. (D)** Tajima's D distribution for all *aus* (215) accessions. **(E)** Joint panel representation for *japonica*, *indica* and *aus* rice population. All x and y axes represents chromosome window per Mb and Tajima's D value, respectively. Red and green horizontal line represents mean Tajima's D value for entire chromosome and desert (8-14Mb) region, respectively. Vertical dash line marks the boundary (8-14Mb) of desert region.

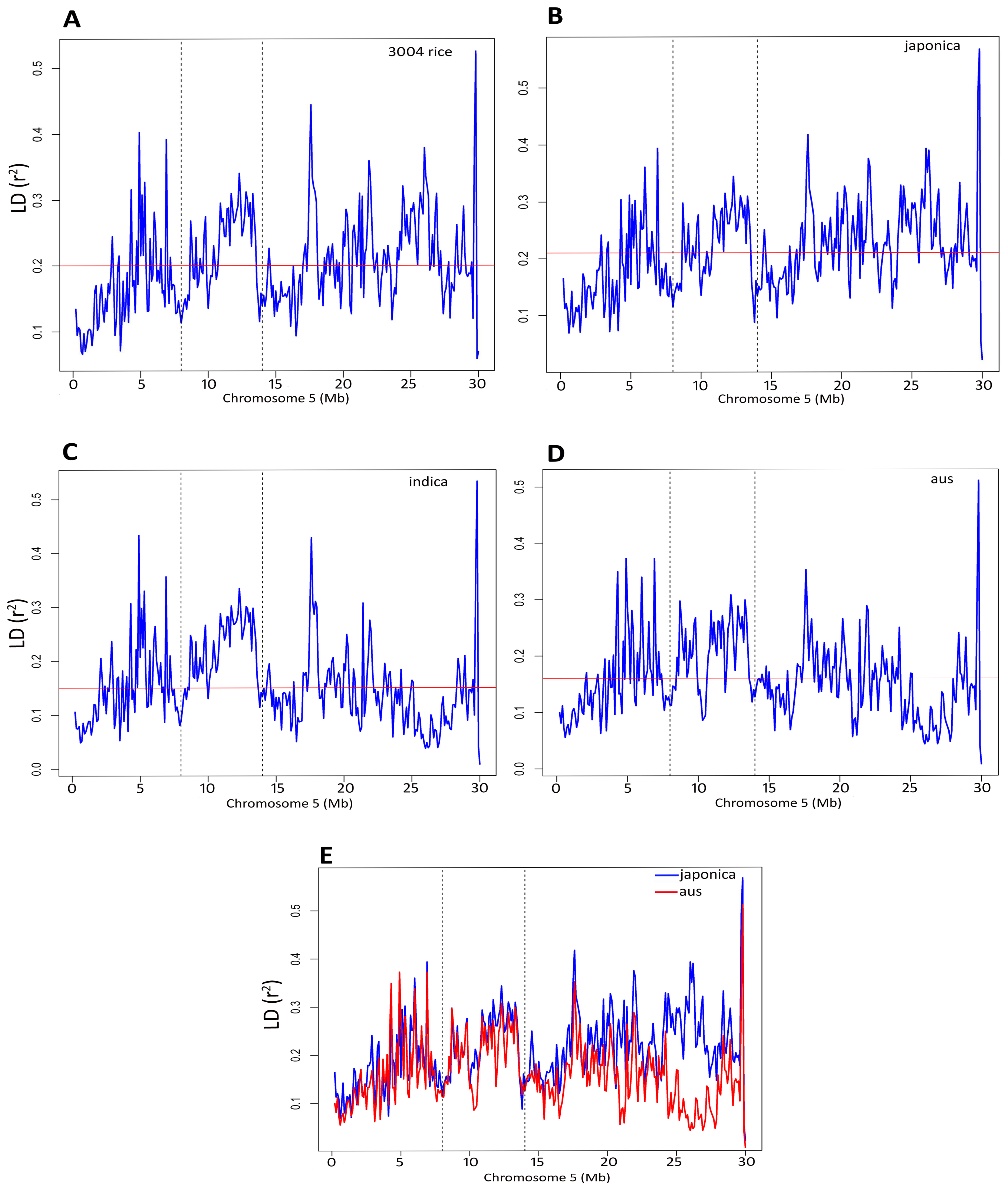


**Supplemental Figure 13**: **Analysis and distribution of Linkage Disequilibrium ($r^2$) estimation per 100 kb of chromosome 5 for different rice groups. (A)** LD **($r^2$)** distribution for all 3004 accessions**. (B)** LD **($r^2$)** distribution for all *japonica* (840) accessions**. (C)** LD **($r^2$)** distribution for all *indica* (1743) accessions**. (D)** LD **($r^2$)** distribution for all *aus* (215) accessions. **(E)** Joint panel representation for *japonica*, *indica* and *aus* rice population. All x and y axes represents chromosome window per Mb and LD **($r^2$)** value, respectively. Red and horizontal line represents mean Tajima's D value for entire chromosome 5. Vertical dash line marks the boundary (8-14Mb) of desert region.

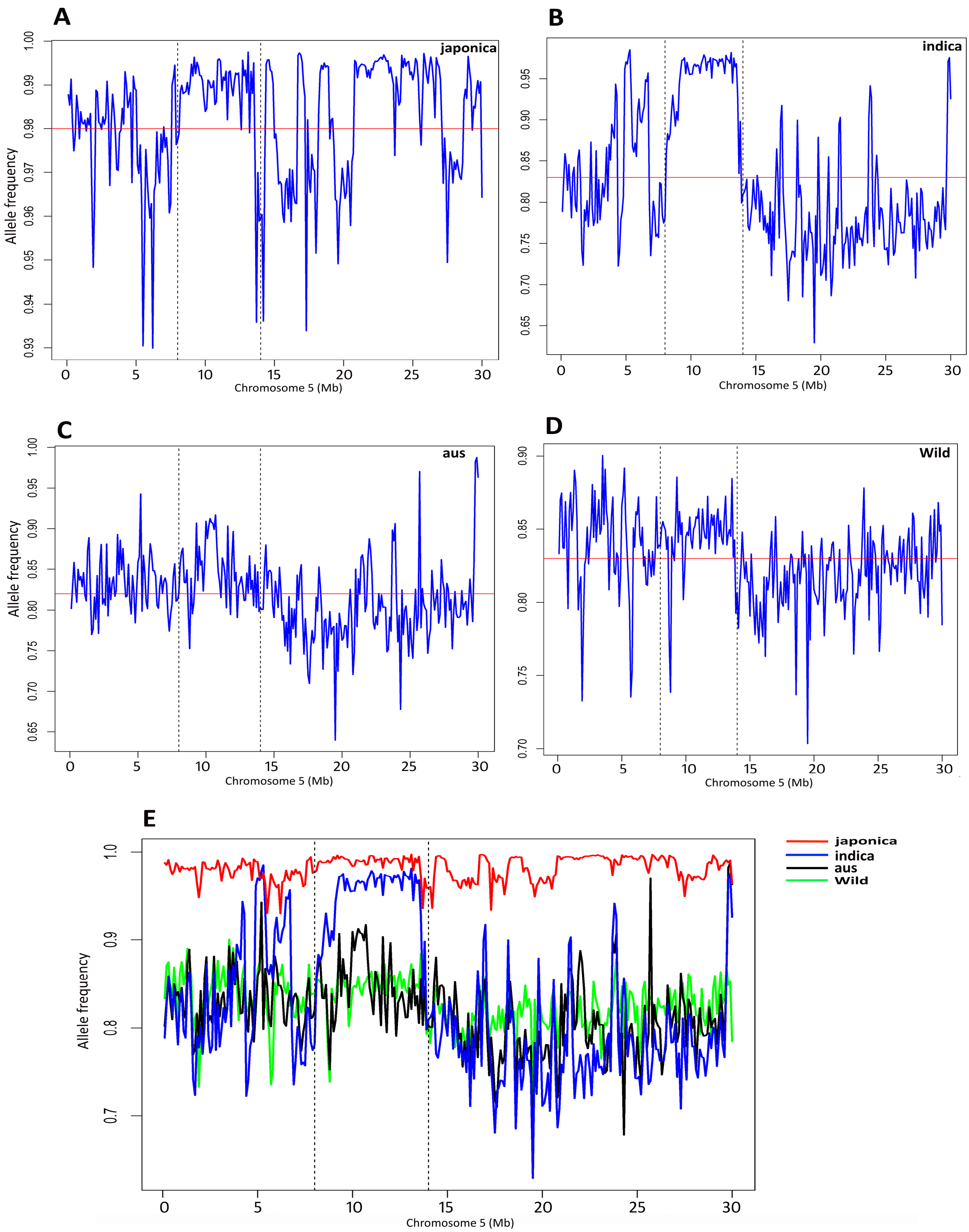


**Supplemental Figure 14**: **Analysis of allele frequency spectrum per 100 kb of chromosome 5 for different rice groups. (A)** Allele frequency distribution for all *japonica* (840) accessions**. (B)** Allele frequency distribution for all *indica* (1743) accessions**. (C)** Allele frequency distribution for all *aus* (215) accessions. **(D)** Allele frequency distribution for wild (446) accessions. **(E)** Joint panel representation for *japonica*, *indica*, *aus* and wild rice population. All x and y axes represents chromosome window per Mb and Allele frequency, respectively. Red and horizontal line represents mean Allele frequency for entire chromosome 5. Vertical dash line marks the boundary (8-14Mb) of desert region.

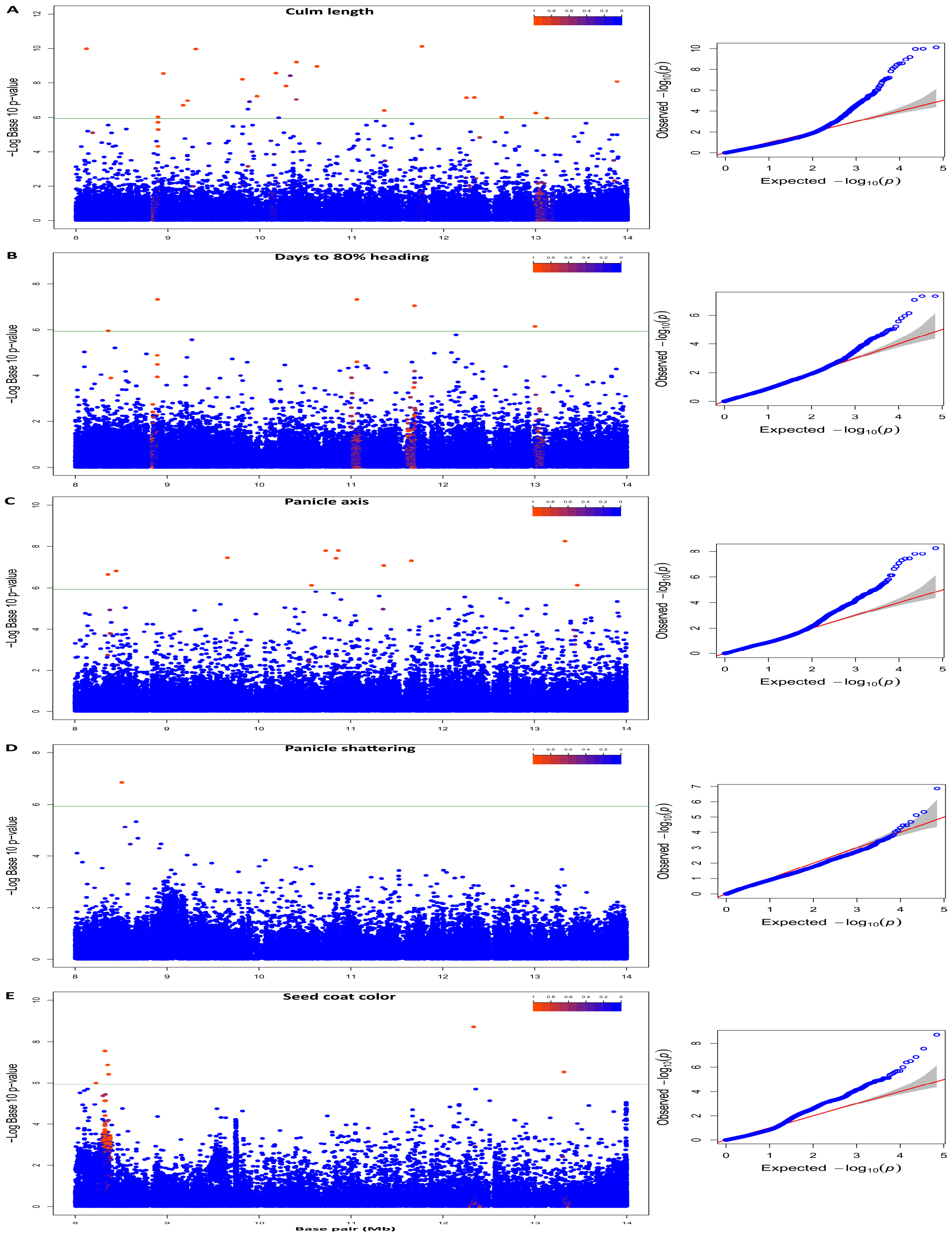


**Supplemental Figure 15**: **Mapping of associations for various yield-related domesticated traits across chromosome 5 desert (8-14 Mb) region**. (**A**) Manhattan and QQ plots of compressed MLM for culm length. Negative $\log_{10}$-transformed *P* values (y axis) values from the compressed mixed linear model are plotted against position (x axis) on chromosome 5 desert region. (**B**) Manhattan plots of compressed MLM for 80% heading date, as in A. (**C**) Manhattan plots of compressed MLM for panicle axis, as in A. (**D**) Manhattan plots of compressed MLM for panicle shattering, as in A. (**E**) Manhattan plots of compressed MLM for seed coat color, as in A. Color bar (blue to red) denotes LD ($r^2$) value, extent of Linkage Disequilibrium. Green horizontal line indicates the cut-off for significant marker association.
