## Supplemental Table 10 for "Genome-wide analysis of polymorphisms identified domestication-associated polymorphism desert carrying important rice grain size/weight QTL"

**Supplemental Table 10: Genome fraction estimation of 3004 rice genotypes through fastStructure (K=7) meanQ analysis**

| Accession | Population_1 | Population_2 | Population_3 | Population_4 | Population_5 | Population_6 | Population_7 |
|---|---|---|---|---|---|---|---|
| B001 | 0.000 | 0.000 | 0.000 | 0.000 | 0.984 | 0.016 | 0.000 |
| B002 | 0.000 | 0.000 | 0.000 | 0.000 | 1.000 | 0.000 | 0.000 |
| B003 | 0.000 | 0.402 | 0.000 | 0.000 | 0.598 | 0.000 | 0.000 |
| B004 | 0.000 | 0.000 | 0.000 | 0.000 | 1.000 | 0.000 | 0.000 |
| B005 | 0.000 | 0.000 | 0.000 | 0.000 | 1.000 | 0.000 | 0.000 |
| B006 | 0.000 | 0.000 | 0.000 | 0.581 | 0.000 | 0.419 | 0.000 |
| B007 | 0.000 | 0.000 | 0.000 | 0.733 | 0.005 | 0.262 | 0.000 |
| B008 | 0.000 | 0.020 | 0.000 | 0.000 | 0.980 | 0.000 | 0.000 |
| B009 | 0.000 | 0.000 | 0.000 | 0.138 | 0.000 | 0.862 | 0.000 |
| B010 | 0.000 | 0.000 | 0.000 | 0.000 | 0.013 | 0.987 | 0.000 |
| B011 | 0.078 | 0.000 | 0.000 | 0.413 | 0.005 | 0.505 | 0.000 |
| B012 | 0.078 | 0.000 | 0.004 | 0.552 | 0.045 | 0.322 | 0.000 |
| B013 | 0.000 | 0.000 | 0.000 | 0.158 | 0.156 | 0.242 | 0.444 |
| B014 | 0.000 | 0.000 | 0.000 | 0.000 | 0.964 | 0.036 | 0.000 |
| B015 | 0.000 | 0.000 | 0.000 | 0.000 | 0.000 | 1.000 | 0.000 |
| B016 | 0.000 | 0.075 | 0.000 | 0.000 | 0.925 | 0.000 | 0.000 |
| B017 | 0.000 | 0.000 | 0.000 | 0.000 | 1.000 | 0.000 | 0.000 |
| B018 | 0.000 | 0.957 | 0.000 | 0.000 | 0.043 | 0.000 | 0.000 |
| B019 | 0.014 | 0.022 | 0.000 | 0.000 | 0.000 | 0.093 | 0.870 |
| B020 | 0.000 | 0.398 | 0.000 | 0.000 | 0.000 | 0.000 | 0.602 |
| B021 | 0.000 | 0.000 | 0.000 | 0.581 | 0.033 | 0.385 | 0.001 |
| B023 | 0.000 | 0.000 | 0.000 | 0.000 | 0.960 | 0.040 | 0.000 |
| B024 | 0.000 | 0.000 | 0.000 | 0.225 | 0.019 | 0.756 | 0.000 |
| B025 | 0.000 | 0.933 | 0.000 | 0.000 | 0.067 | 0.000 | 0.000 |
| B026 | 0.078 | 0.000 | 0.000 | 0.411 | 0.005 | 0.506 | 0.000 |
| B027 | 0.000 | 0.000 | 0.000 | 0.221 | 0.000 | 0.000 | 0.779 |
| B028 | 0.035 | 0.000 | 0.000 | 0.289 | 0.041 | 0.636 | 0.000 |
| B029 | 0.000 | 0.000 | 0.000 | 0.000 | 0.098 | 0.385 | 0.517 |
| B030 | 0.017 | 0.000 | 0.000 | 0.969 | 0.000 | 0.014 | 0.000 |
| B031 | 0.046 | 0.000 | 0.000 | 0.954 | 0.000 | 0.000 | 0.000 |
| B032 | 0.018 | 0.000 | 0.000 | 0.044 | 0.000 | 0.302 | 0.637 |
| B033 | 0.017 | 0.000 | 0.000 | 0.000 | 0.000 | 0.257 | 0.726 |
| B034 | 0.000 | 0.000 | 0.000 | 0.000 | 1.000 | 0.000 | 0.000 |
| B035 | 0.002 | 0.030 | 0.015 | 0.134 | 0.010 | 0.194 | 0.615 |
| B036 | 0.000 | 0.536 | 0.000 | 0.000 | 0.464 | 0.000 | 0.000 |
| B037 | 0.000 | 0.695 | 0.000 | 0.004 | 0.285 | 0.012 | 0.003 |
| B038 | 0.000 | 0.139 | 0.000 | 0.000 | 0.861 | 0.000 | 0.000 |
| B039 | 0.000 | 0.000 | 0.000 | 0.331 | 0.357 | 0.311 | 0.000 |
| B040 | 0.000 | 0.000 | 0.000 | 0.000 | 0.000 | 0.680 | 0.320 |
| B043 | 0.000 | 0.857 | 0.000 | 0.000 | 0.143 | 0.000 | 0.000 |
| B044 | 0.000 | 0.038 | 0.183 | 0.375 | 0.000 | 0.404 | 0.000 |
| B045 | 0.000 | 0.000 | 0.000 | 0.000 | 1.000 | 0.000 | 0.000 |
| B046 | 0.000 | 0.000 | 0.000 | 0.000 | 1.000 | 0.000 | 0.000 |
| B047 | 0.000 | 0.000 | 0.000 | 0.141 | 0.734 | 0.125 | 0.000 |
| B048 | 0.048 | 0.000 | 0.000 | 0.263 | 0.000 | 0.001 | 0.688 |
| B049 | 0.847 | 0.000 | 0.000 | 0.000 | 0.082 | 0.066 | 0.005 |
| B051 | 0.000 | 0.033 | 0.000 | 0.109 | 0.000 | 0.275 | 0.583 |
| B052 | 0.310 | 0.000 | 0.000 | 0.690 | 0.000 | 0.000 | 0.000 |
| B053 | 0.000 | 0.747 | 0.000 | 0.000 | 0.242 | 0.011 | 0.000 |
| B054 | 0.000 | 0.834 | 0.000 | 0.000 | 0.166 | 0.000 | 0.000 |
| B055 | 0.000 | 0.169 | 0.000 | 0.000 | 0.711 | 0.121 | 0.000 |
| B056 | 0.000 | 0.000 | 0.000 | 0.003 | 0.980 | 0.017 | 0.000 |
| B057 | 0.000 | 0.000 | 0.000 | 0.000 | 0.975 | 0.008 | 0.016 |
| B058 | 0.000 | 0.000 | 0.000 | 0.099 | 0.000 | 0.221 | 0.680 |
| B059 | 0.000 | 0.000 | 0.000 | 0.000 | 0.000 | 1.000 | 0.000 |
| B060 | 0.000 | 0.000 | 0.000 | 0.000 | 0.000 | 1.000 | 0.000 |
| B061 | 0.000 | 0.000 | 0.000 | 0.000 | 0.000 | 1.000 | 0.000 |
| B062 | 0.000 | 0.000 | 0.000 | 0.000 | 0.009 | 0.978 | 0.013 |
| B063 | 0.000 | 0.000 | 0.000 | 0.000 | 0.000 | 0.772 | 0.228 |
| B064 | 0.021 | 0.000 | 0.000 | 0.250 | 0.008 | 0.721 | 0.000 |
| B065 | 0.000 | 0.000 | 0.000 | 0.000 | 0.000 | 0.497 | 0.503 |
| B066 | 0.000 | 0.048 | 0.000 | 0.000 | 0.952 | 0.000 | 0.000 |
| B067 | 0.000 | 0.000 | 0.000 | 0.000 | 0.000 | 0.928 | 0.072 |
| B068 | 0.000 | 0.000 | 0.000 | 0.000 | 1.000 | 0.000 | 0.000 |

| | | | | | | | |
|---|---|---|---|---|---|---|---|
| B069 | 0.000 | 0.390 | 0.000 | 0.000 | 0.610 | 0.000 | 0.000 |
| B070 | 0.000 | 0.368 | 0.000 | 0.000 | 0.632 | 0.000 | 0.000 |
| B071 | 0.000 | 0.000 | 0.000 | 0.000 | 1.000 | 0.000 | 0.000 |
| B072 | 0.000 | 0.000 | 0.000 | 0.089 | 0.159 | 0.752 | 0.000 |
| B073 | 0.000 | 0.000 | 0.034 | 0.382 | 0.000 | 0.585 | 0.000 |
| B074 | 0.000 | 0.000 | 0.000 | 0.000 | 0.032 | 0.968 | 0.000 |
| B075 | 0.000 | 0.007 | 0.000 | 0.292 | 0.018 | 0.631 | 0.052 |
| B076 | 0.000 | 0.000 | 0.001 | 0.374 | 0.018 | 0.607 | 0.000 |
| B077 | 0.000 | 0.000 | 0.000 | 0.000 | 1.000 | 0.000 | 0.000 |
| B079 | 0.000 | 0.000 | 0.000 | 0.151 | 0.000 | 0.849 | 0.000 |
| B081 | 0.000 | 0.004 | 0.000 | 0.135 | 0.002 | 0.681 | 0.177 |
| B082 | 0.000 | 0.000 | 0.000 | 0.479 | 0.043 | 0.478 | 0.000 |
| B083 | 0.000 | 0.000 | 0.000 | 0.000 | 0.000 | 0.955 | 0.045 |
| B084 | 0.000 | 0.000 | 0.000 | 0.000 | 0.603 | 0.000 | 0.397 |
| B085 | 0.000 | 0.000 | 0.000 | 0.000 | 0.000 | 0.857 | 0.143 |
| B086 | 0.000 | 0.000 | 0.000 | 0.102 | 0.538 | 0.360 | 0.000 |
| B087 | 0.038 | 0.000 | 0.000 | 0.949 | 0.013 | 0.000 | 0.000 |
| B088 | 0.003 | 0.030 | 0.015 | 0.133 | 0.011 | 0.195 | 0.614 |
| B089 | 0.000 | 0.024 | 0.000 | 0.399 | 0.000 | 0.577 | 0.000 |
| B090 | 0.055 | 0.041 | 0.000 | 0.341 | 0.000 | 0.563 | 0.000 |
| B091 | 0.072 | 0.000 | 0.000 | 0.327 | 0.000 | 0.599 | 0.001 |
| B092 | 0.000 | 0.000 | 0.000 | 0.000 | 0.010 | 0.990 | 0.000 |
| B093 | 0.000 | 0.048 | 0.000 | 0.306 | 0.005 | 0.642 | 0.000 |
| B094 | 0.000 | 0.000 | 0.000 | 0.182 | 0.014 | 0.804 | 0.000 |
| B095 | 0.000 | 0.000 | 0.000 | 0.713 | 0.000 | 0.287 | 0.000 |
| B096 | 0.000 | 0.007 | 0.000 | 0.504 | 0.000 | 0.489 | 0.000 |
| B097 | 0.000 | 0.000 | 0.000 | 0.552 | 0.000 | 0.448 | 0.000 |
| B100 | 0.000 | 0.061 | 0.000 | 0.000 | 0.939 | 0.000 | 0.000 |
| B101 | 0.000 | 0.000 | 0.000 | 0.000 | 0.930 | 0.000 | 0.070 |
| B102 | 0.000 | 0.105 | 0.000 | 0.000 | 0.895 | 0.000 | 0.000 |
| B103 | 0.000 | 0.000 | 0.016 | 0.000 | 0.977 | 0.006 | 0.001 |
| B104 | 0.000 | 0.000 | 0.000 | 0.000 | 0.000 | 0.834 | 0.166 |
| B105 | 0.000 | 0.000 | 0.000 | 0.000 | 0.026 | 0.642 | 0.332 |
| B106 | 0.000 | 0.000 | 0.005 | 0.591 | 0.000 | 0.404 | 0.000 |
| B107 | 0.000 | 0.000 | 0.000 | 0.637 | 0.000 | 0.363 | 0.000 |
| B108 | 0.054 | 0.000 | 0.000 | 0.232 | 0.007 | 0.707 | 0.000 |
| B109 | 0.000 | 0.224 | 0.000 | 0.062 | 0.713 | 0.000 | 0.001 |
| B110 | 0.000 | 0.000 | 0.022 | 0.000 | 0.978 | 0.000 | 0.000 |
| B111 | 0.000 | 0.081 | 0.000 | 0.000 | 0.919 | 0.000 | 0.000 |
| B112 | 0.000 | 0.000 | 0.000 | 0.000 | 0.000 | 1.000 | 0.000 |
| B113 | 0.000 | 0.000 | 0.000 | 0.346 | 0.000 | 0.644 | 0.009 |
| B114 | 0.000 | 0.000 | 0.000 | 0.000 | 0.000 | 1.000 | 0.000 |
| B115 | 0.000 | 0.000 | 0.000 | 0.000 | 0.000 | 0.972 | 0.028 |
| B116 | 0.000 | 0.000 | 0.000 | 0.416 | 0.000 | 0.584 | 0.000 |
| B117 | 0.000 | 0.000 | 0.000 | 0.000 | 1.000 | 0.000 | 0.000 |
| B118 | 0.000 | 0.000 | 0.000 | 0.000 | 0.027 | 0.694 | 0.279 |
| B119 | 0.000 | 0.000 | 0.000 | 0.000 | 0.016 | 0.984 | 0.000 |
| B120 | 0.000 | 0.000 | 0.000 | 0.000 | 0.005 | 0.651 | 0.344 |
| B121 | 0.000 | 0.000 | 0.000 | 0.000 | 0.000 | 1.000 | 0.000 |
| B122 | 0.000 | 0.000 | 0.000 | 0.000 | 1.000 | 0.000 | 0.000 |
| B123 | 0.000 | 0.000 | 0.000 | 0.326 | 0.074 | 0.426 | 0.173 |
| B124 | 0.000 | 0.000 | 0.000 | 0.000 | 0.813 | 0.091 | 0.096 |
| B125 | 0.000 | 0.000 | 0.000 | 0.000 | 0.041 | 0.495 | 0.465 |
| B126 | 0.000 | 0.000 | 0.000 | 0.000 | 0.034 | 0.966 | 0.000 |
| B127 | 0.000 | 0.000 | 0.000 | 0.000 | 0.031 | 0.969 | 0.000 |
| B128 | 0.000 | 0.000 | 0.000 | 0.000 | 0.169 | 0.221 | 0.611 |
| B129 | 0.000 | 0.000 | 0.000 | 0.458 | 0.036 | 0.506 | 0.000 |
| B130 | 0.000 | 0.000 | 0.000 | 0.000 | 0.077 | 0.923 | 0.000 |
| B131 | 0.008 | 0.000 | 0.035 | 0.478 | 0.000 | 0.479 | 0.000 |
| B132 | 0.000 | 0.000 | 0.000 | 0.464 | 0.131 | 0.405 | 0.000 |
| B133 | 0.000 | 0.000 | 0.001 | 0.531 | 0.047 | 0.421 | 0.000 |
| B134 | 0.000 | 0.410 | 0.000 | 0.000 | 0.590 | 0.000 | 0.000 |
| B135 | 0.000 | 0.000 | 0.006 | 0.602 | 0.038 | 0.345 | 0.009 |
| B136 | 0.000 | 0.130 | 0.000 | 0.000 | 0.870 | 0.000 | 0.000 |
| B137 | 0.000 | 0.000 | 0.000 | 0.186 | 0.045 | 0.406 | 0.363 |
| B138 | 0.012 | 0.044 | 0.000 | 0.159 | 0.000 | 0.216 | 0.570 |
| B139 | 0.000 | 0.004 | 0.000 | 0.000 | 0.084 | 0.547 | 0.366 |

| | | | | | | | |
|---|---|---|---|---|---|---|---|
| B140 | 0.000 | 0.000 | 0.000 | 0.028 | 0.000 | 0.972 | 0.000 |
| B141 | 0.000 | 0.000 | 0.000 | 0.000 | 0.408 | 0.000 | 0.592 |
| B142 | 0.000 | 0.000 | 0.000 | 0.000 | 0.620 | 0.000 | 0.380 |
| B143 | 0.000 | 0.000 | 0.000 | 0.000 | 0.683 | 0.101 | 0.216 |
| B144 | 0.000 | 0.308 | 0.000 | 0.000 | 0.612 | 0.079 | 0.000 |
| B145 | 0.035 | 0.000 | 0.000 | 0.000 | 0.081 | 0.205 | 0.678 |
| B146 | 0.000 | 0.000 | 0.000 | 0.000 | 0.000 | 0.223 | 0.777 |
| B147 | 0.000 | 0.000 | 0.000 | 0.070 | 0.079 | 0.851 | 0.000 |
| B148 | 0.000 | 0.126 | 0.000 | 0.000 | 0.874 | 0.000 | 0.000 |
| B149 | 0.000 | 0.000 | 0.000 | 0.000 | 0.000 | 1.000 | 0.000 |
| B150 | 0.000 | 0.000 | 0.000 | 0.094 | 0.068 | 0.838 | 0.000 |
| B151 | 0.000 | 0.000 | 0.000 | 0.145 | 0.000 | 0.855 | 0.000 |
| B152 | 0.011 | 0.000 | 0.000 | 0.000 | 0.823 | 0.116 | 0.050 |
| B153 | 0.000 | 0.001 | 0.000 | 0.000 | 0.050 | 0.440 | 0.509 |
| B154 | 0.000 | 0.000 | 0.000 | 0.000 | 1.000 | 0.000 | 0.000 |
| B155 | 0.012 | 0.119 | 0.000 | 0.141 | 0.000 | 0.459 | 0.269 |
| B156 | 0.000 | 0.000 | 0.000 | 0.099 | 0.000 | 0.901 | 0.000 |
| B157 | 0.000 | 0.000 | 0.000 | 0.000 | 0.000 | 0.642 | 0.358 |
| B158 | 0.000 | 0.000 | 0.000 | 0.000 | 0.472 | 0.372 | 0.157 |
| B159 | 0.000 | 0.013 | 0.000 | 0.000 | 0.000 | 0.681 | 0.307 |
| B160 | 0.000 | 0.000 | 0.000 | 0.000 | 1.000 | 0.000 | 0.000 |
| B161 | 0.000 | 0.014 | 0.000 | 0.000 | 0.986 | 0.000 | 0.000 |
| B162 | 0.000 | 0.000 | 0.000 | 0.000 | 1.000 | 0.000 | 0.000 |
| B163 | 0.011 | 0.025 | 0.000 | 0.124 | 0.000 | 0.840 | 0.000 |
| B164 | 0.917 | 0.000 | 0.000 | 0.083 | 0.000 | 0.000 | 0.000 |
| B165 | 0.000 | 0.000 | 0.000 | 0.571 | 0.000 | 0.429 | 0.000 |
| B166 | 0.000 | 0.000 | 0.000 | 0.000 | 1.000 | 0.000 | 0.000 |
| B167 | 0.000 | 0.000 | 0.000 | 0.000 | 1.000 | 0.000 | 0.000 |
| B168 | 0.000 | 0.275 | 0.000 | 0.000 | 0.725 | 0.000 | 0.000 |
| B169 | 0.000 | 0.000 | 0.000 | 0.000 | 1.000 | 0.000 | 0.000 |
| B170 | 0.000 | 0.000 | 0.000 | 0.000 | 1.000 | 0.000 | 0.000 |
| B171 | 0.000 | 0.000 | 0.000 | 0.000 | 1.000 | 0.000 | 0.000 |
| B173 | 0.067 | 0.245 | 0.103 | 0.560 | 0.024 | 0.000 | 0.000 |
| B176 | 0.000 | 0.180 | 0.000 | 0.000 | 0.000 | 0.075 | 0.745 |
| B179 | 0.000 | 0.000 | 0.000 | 0.000 | 1.000 | 0.000 | 0.000 |
| B180 | 0.018 | 0.000 | 0.002 | 0.188 | 0.000 | 0.000 | 0.792 |
| B181 | 0.000 | 0.000 | 0.000 | 0.000 | 0.005 | 0.995 | 0.000 |
| B182 | 0.000 | 0.000 | 0.000 | 0.000 | 1.000 | 0.000 | 0.000 |
| B183 | 0.000 | 0.000 | 0.000 | 0.000 | 1.000 | 0.000 | 0.000 |
| B184 | 0.007 | 0.000 | 0.024 | 0.539 | 0.049 | 0.381 | 0.000 |
| B185 | 0.000 | 0.000 | 0.000 | 0.001 | 0.000 | 0.574 | 0.425 |
| B187 | 0.116 | 0.009 | 0.000 | 0.364 | 0.000 | 0.419 | 0.091 |
| B188 | 0.000 | 1.000 | 0.000 | 0.000 | 0.000 | 0.000 | 0.000 |
| B189 | 0.000 | 0.964 | 0.000 | 0.000 | 0.036 | 0.000 | 0.000 |
| B190 | 0.000 | 1.000 | 0.000 | 0.000 | 0.000 | 0.000 | 0.000 |
| B191 | 0.008 | 0.687 | 0.000 | 0.000 | 0.306 | 0.000 | 0.000 |
| B192 | 0.089 | 0.438 | 0.000 | 0.030 | 0.000 | 0.000 | 0.444 |
| B193 | 0.000 | 0.008 | 0.000 | 0.000 | 0.008 | 0.000 | 0.984 |
| B194 | 0.000 | 0.000 | 0.000 | 0.109 | 0.008 | 0.147 | 0.737 |
| B195 | 0.002 | 0.000 | 0.000 | 0.162 | 0.007 | 0.214 | 0.616 |
| B196 | 0.000 | 0.697 | 0.000 | 0.000 | 0.240 | 0.000 | 0.063 |
| B197 | 0.000 | 0.000 | 0.000 | 0.413 | 0.000 | 0.587 | 0.000 |
| B198 | 0.000 | 0.000 | 0.000 | 0.000 | 0.000 | 1.000 | 0.000 |
| B199 | 0.000 | 0.000 | 0.000 | 0.000 | 0.982 | 0.018 | 0.000 |
| B200 | 0.004 | 0.000 | 0.000 | 0.000 | 0.000 | 0.130 | 0.867 |
| B201 | 0.000 | 0.018 | 0.000 | 0.000 | 0.000 | 0.132 | 0.850 |
| B202 | 0.032 | 0.032 | 0.000 | 0.000 | 0.038 | 0.446 | 0.452 |
| B203 | 0.019 | 0.000 | 0.000 | 0.431 | 0.000 | 0.551 | 0.000 |
| B204 | 0.000 | 0.002 | 0.000 | 0.000 | 0.998 | 0.000 | 0.000 |
| B205 | 0.000 | 0.263 | 0.000 | 0.000 | 0.737 | 0.000 | 0.000 |
| B207 | 0.000 | 0.000 | 0.030 | 0.458 | 0.000 | 0.512 | 0.000 |
| B208 | 0.000 | 0.000 | 0.000 | 0.000 | 0.000 | 1.000 | 0.000 |
| B210 | 0.000 | 0.011 | 0.000 | 0.258 | 0.000 | 0.731 | 0.000 |
| B212 | 0.018 | 0.000 | 0.030 | 0.044 | 0.869 | 0.039 | 0.000 |
| B213 | 0.000 | 0.000 | 0.000 | 0.000 | 0.000 | 1.000 | 0.000 |
| B214 | 0.000 | 0.000 | 0.000 | 0.000 | 0.000 | 0.000 | 1.000 |
| B215 | 0.000 | 0.000 | 0.000 | 0.000 | 1.000 | 0.000 | 0.000 |

| | | | | | | | |
|---|---|---|---|---|---|---|---|
| B216 | 0.000 | 0.000 | 0.000 | 0.277 | 0.022 | 0.700 | 0.000 |
| B217 | 0.000 | 0.000 | 0.000 | 0.000 | 0.000 | 0.000 | 1.000 |
| B218 | 0.000 | 0.000 | 0.000 | 0.000 | 1.000 | 0.000 | 0.000 |
| B219 | 0.000 | 0.000 | 0.000 | 0.571 | 0.000 | 0.429 | 0.000 |
| B221 | 0.413 | 0.000 | 0.000 | 0.000 | 0.000 | 0.587 | 0.000 |
| B222 | 0.000 | 0.004 | 0.000 | 0.184 | 0.003 | 0.809 | 0.000 |
| B223 | 0.000 | 0.145 | 0.000 | 0.000 | 0.855 | 0.000 | 0.000 |
| B224 | 0.045 | 0.000 | 0.000 | 0.253 | 0.000 | 0.702 | 0.000 |
| B225 | 0.000 | 0.000 | 0.024 | 0.000 | 0.976 | 0.000 | 0.000 |
| B226 | 0.000 | 0.000 | 0.000 | 0.000 | 1.000 | 0.000 | 0.000 |
| B227 | 0.000 | 0.000 | 0.000 | 0.473 | 0.121 | 0.406 | 0.000 |
| B228 | 0.003 | 0.332 | 0.000 | 0.010 | 0.648 | 0.005 | 0.001 |
| B229 | 0.000 | 0.017 | 0.000 | 0.418 | 0.000 | 0.566 | 0.000 |
| B230 | 0.000 | 0.190 | 0.000 | 0.000 | 0.810 | 0.000 | 0.000 |
| B232 | 0.000 | 0.000 | 0.000 | 0.000 | 0.000 | 0.000 | 1.000 |
| B233 | 0.000 | 0.000 | 0.000 | 0.000 | 0.000 | 0.517 | 0.483 |
| B234 | 0.000 | 0.000 | 0.000 | 0.000 | 0.000 | 1.000 | 0.000 |
| B235 | 0.000 | 0.000 | 0.000 | 0.039 | 0.961 | 0.000 | 0.000 |
| B236 | 0.000 | 0.000 | 0.000 | 0.000 | 1.000 | 0.000 | 0.000 |
| B238 | 0.000 | 0.000 | 0.000 | 0.416 | 0.044 | 0.540 | 0.000 |
| B239 | 0.000 | 0.000 | 0.000 | 0.176 | 0.012 | 0.813 | 0.000 |
| B240 | 0.000 | 0.000 | 0.000 | 0.000 | 1.000 | 0.000 | 0.000 |
| B241 | 0.000 | 0.565 | 0.000 | 0.000 | 0.427 | 0.008 | 0.000 |
| B242 | 0.000 | 0.071 | 0.000 | 0.234 | 0.000 | 0.000 | 0.696 |
| B243 | 0.940 | 0.000 | 0.000 | 0.060 | 0.000 | 0.000 | 0.000 |
| B244 | 0.000 | 0.000 | 0.000 | 0.075 | 0.029 | 0.896 | 0.000 |
| B245 | 0.000 | 0.338 | 0.000 | 0.000 | 0.662 | 0.000 | 0.000 |
| B246 | 0.000 | 0.000 | 0.000 | 0.695 | 0.030 | 0.275 | 0.000 |
| B247 | 0.003 | 0.018 | 0.000 | 0.497 | 0.000 | 0.482 | 0.000 |
| B248 | 0.000 | 0.000 | 0.000 | 0.067 | 0.000 | 0.933 | 0.000 |
| B249 | 0.000 | 0.000 | 0.000 | 0.000 | 0.004 | 0.996 | 0.000 |
| B250 | 0.000 | 0.000 | 0.000 | 0.000 | 1.000 | 0.000 | 0.000 |
| B252 | 0.000 | 0.009 | 0.000 | 0.000 | 0.021 | 0.516 | 0.454 |
| B253 | 0.000 | 0.000 | 0.000 | 0.114 | 0.034 | 0.508 | 0.344 |
| B254 | 0.000 | 0.000 | 0.000 | 0.111 | 0.028 | 0.512 | 0.349 |
| B255 | 0.000 | 0.021 | 0.000 | 0.000 | 0.000 | 0.501 | 0.477 |
| B258 | 0.000 | 0.044 | 0.000 | 0.000 | 0.956 | 0.000 | 0.000 |
| B259 | 0.000 | 0.000 | 0.000 | 0.000 | 0.009 | 0.991 | 0.000 |
| B260 | 0.000 | 0.000 | 0.000 | 0.001 | 0.000 | 0.999 | 0.000 |
| B261 | 0.000 | 0.000 | 0.000 | 0.030 | 0.000 | 0.970 | 0.000 |
| B263 | 0.000 | 0.000 | 0.000 | 0.000 | 1.000 | 0.000 | 0.000 |
| B264 | 0.000 | 0.000 | 0.000 | 0.027 | 0.000 | 0.973 | 0.000 |
| B265 | 0.000 | 0.005 | 0.022 | 0.354 | 0.000 | 0.618 | 0.000 |
| B266 | 0.000 | 0.624 | 0.000 | 0.000 | 0.376 | 0.000 | 0.000 |
| B267 | 0.000 | 0.000 | 0.000 | 0.005 | 0.000 | 0.353 | 0.642 |
| B268 | 0.070 | 0.000 | 0.000 | 0.274 | 0.004 | 0.652 | 0.000 |
| B269 | 0.000 | 0.000 | 0.000 | 0.000 | 1.000 | 0.000 | 0.000 |
| CX2 | 0.000 | 0.000 | 0.000 | 0.005 | 0.024 | 0.507 | 0.464 |
| CX3 | 0.000 | 0.000 | 0.000 | 0.397 | 0.000 | 0.603 | 0.000 |
| CX4 | 0.000 | 0.000 | 0.000 | 0.000 | 0.442 | 0.261 | 0.297 |
| CX5 | 0.000 | 0.000 | 0.000 | 0.018 | 0.100 | 0.227 | 0.655 |
| CX6 | 0.000 | 0.000 | 0.000 | 0.000 | 0.545 | 0.000 | 0.455 |
| CX8 | 0.000 | 0.049 | 0.000 | 0.000 | 0.000 | 0.191 | 0.760 |
| CX9 | 0.000 | 0.000 | 0.000 | 0.000 | 0.000 | 0.374 | 0.626 |
| CX10 | 0.000 | 0.000 | 0.000 | 0.000 | 0.030 | 0.970 | 0.000 |
| CX11 | 0.000 | 0.297 | 0.000 | 0.000 | 0.435 | 0.268 | 0.000 |
| CX12 | 0.038 | 0.056 | 0.000 | 0.000 | 0.000 | 0.308 | 0.598 |
| CX13 | 0.045 | 0.000 | 0.000 | 0.000 | 0.257 | 0.126 | 0.572 |
| CX14 | 0.000 | 0.000 | 0.000 | 0.000 | 1.000 | 0.000 | 0.000 |
| CX15 | 0.000 | 0.095 | 0.000 | 0.000 | 0.067 | 0.095 | 0.744 |
| CX16 | 0.000 | 0.000 | 0.000 | 0.051 | 0.948 | 0.000 | 0.000 |
| CX17 | 0.000 | 0.000 | 0.000 | 0.000 | 0.000 | 0.488 | 0.512 |
| CX18 | 0.000 | 0.000 | 0.000 | 0.003 | 0.024 | 0.780 | 0.192 |
| CX19 | 0.000 | 0.000 | 0.000 | 0.000 | 0.000 | 0.513 | 0.487 |
| CX20 | 0.000 | 0.030 | 0.000 | 0.000 | 0.000 | 0.498 | 0.472 |
| CX21 | 0.000 | 0.000 | 0.000 | 0.000 | 0.000 | 0.000 | 1.000 |
| CX22 | 0.000 | 0.000 | 0.000 | 0.000 | 0.000 | 0.000 | 1.000 |

| | | | | | | | |
|---|---|---|---|---|---|---|---|
| CX23 | 0.000 | 0.000 | 0.000 | 0.414 | 0.000 | 0.000 | 0.586 |
| CX24 | 0.084 | 0.000 | 0.000 | 0.314 | 0.000 | 0.203 | 0.399 |
| CX25 | 0.000 | 0.000 | 0.000 | 0.000 | 0.000 | 0.000 | 1.000 |
| CX26 | 0.000 | 0.000 | 0.000 | 0.000 | 0.000 | 0.166 | 0.834 |
| CX27 | 0.000 | 0.000 | 0.000 | 0.183 | 0.000 | 0.126 | 0.691 |
| CX28 | 0.000 | 0.000 | 0.000 | 0.000 | 0.105 | 0.000 | 0.895 |
| CX29 | 0.000 | 0.000 | 0.000 | 0.000 | 0.514 | 0.257 | 0.229 |
| CX30 | 0.000 | 0.000 | 0.000 | 0.000 | 0.021 | 0.000 | 0.979 |
| CX31 | 0.000 | 0.000 | 0.000 | 0.000 | 0.000 | 0.146 | 0.854 |
| CX32 | 0.000 | 0.570 | 0.106 | 0.000 | 0.168 | 0.155 | 0.000 |
| CX33 | 0.000 | 0.000 | 0.000 | 0.000 | 0.000 | 0.377 | 0.623 |
| CX34 | 0.037 | 0.000 | 0.000 | 0.430 | 0.000 | 0.166 | 0.367 |
| CX35 | 0.000 | 0.000 | 0.000 | 0.000 | 0.000 | 0.303 | 0.697 |
| CX37 | 0.000 | 0.000 | 0.000 | 0.000 | 0.000 | 0.147 | 0.853 |
| CX42 | 0.000 | 0.000 | 0.000 | 0.000 | 0.000 | 0.511 | 0.489 |
| CX43 | 0.000 | 0.005 | 0.000 | 0.000 | 0.003 | 0.224 | 0.768 |
| CX44 | 0.000 | 0.000 | 0.000 | 0.128 | 0.000 | 0.140 | 0.732 |
| CX45 | 0.000 | 0.000 | 0.000 | 0.000 | 0.000 | 0.000 | 1.000 |
| CX46 | 0.000 | 0.000 | 0.456 | 0.000 | 0.000 | 0.097 | 0.447 |
| CX47 | 0.000 | 0.101 | 0.000 | 0.000 | 0.899 | 0.000 | 0.000 |
| CX48 | 0.000 | 0.003 | 0.000 | 0.000 | 0.007 | 0.130 | 0.861 |
| CX49 | 0.000 | 0.000 | 0.000 | 0.000 | 0.757 | 0.000 | 0.243 |
| CX50 | 0.000 | 0.000 | 0.000 | 0.000 | 0.000 | 0.378 | 0.622 |
| CX51 | 0.000 | 0.000 | 0.000 | 0.380 | 0.122 | 0.498 | 0.000 |
| CX52 | 0.000 | 0.000 | 0.000 | 0.693 | 0.000 | 0.307 | 0.000 |
| CX53 | 0.000 | 0.000 | 0.000 | 0.344 | 0.000 | 0.344 | 0.312 |
| CX54 | 0.000 | 0.000 | 0.000 | 0.549 | 0.000 | 0.443 | 0.008 |
| CX55 | 0.000 | 0.000 | 0.000 | 0.000 | 0.000 | 1.000 | 0.000 |
| CX56 | 0.000 | 0.000 | 0.000 | 0.000 | 1.000 | 0.000 | 0.000 |
| CX57 | 0.000 | 0.000 | 0.000 | 0.027 | 0.827 | 0.000 | 0.147 |
| CX58 | 0.000 | 0.113 | 0.000 | 0.000 | 0.887 | 0.000 | 0.000 |
| CX59 | 0.000 | 0.000 | 1.000 | 0.000 | 0.000 | 0.000 | 0.000 |
| CX60 | 0.000 | 0.000 | 0.022 | 0.000 | 0.085 | 0.161 | 0.732 |
| CX61 | 0.000 | 0.001 | 0.011 | 0.295 | 0.016 | 0.676 | 0.000 |
| CX63 | 1.000 | 0.000 | 0.000 | 0.000 | 0.000 | 0.000 | 0.000 |
| CX64 | 0.000 | 0.038 | 0.000 | 0.171 | 0.000 | 0.000 | 0.791 |
| CX65 | 0.000 | 0.000 | 1.000 | 0.000 | 0.000 | 0.000 | 0.000 |
| CX66 | 0.000 | 0.000 | 1.000 | 0.000 | 0.000 | 0.000 | 0.000 |
| CX67 | 0.000 | 0.000 | 1.000 | 0.000 | 0.000 | 0.000 | 0.000 |
| CX68 | 0.000 | 0.000 | 0.000 | 0.147 | 0.000 | 0.000 | 0.853 |
| CX69 | 0.000 | 0.000 | 0.328 | 0.000 | 0.000 | 0.182 | 0.490 |
| CX70 | 0.000 | 0.000 | 0.000 | 0.171 | 0.000 | 0.000 | 0.829 |
| CX71 | 0.000 | 0.000 | 0.000 | 0.970 | 0.000 | 0.030 | 0.000 |
| CX72 | 0.172 | 0.000 | 0.649 | 0.000 | 0.000 | 0.058 | 0.121 |
| CX73 | 0.000 | 0.000 | 0.000 | 0.000 | 0.000 | 0.000 | 1.000 |
| CX74 | 0.000 | 0.000 | 0.000 | 0.000 | 1.000 | 0.000 | 0.000 |
| CX75 | 0.000 | 0.000 | 0.002 | 0.082 | 0.000 | 0.287 | 0.628 |
| CX76 | 0.000 | 0.007 | 0.000 | 0.133 | 0.000 | 0.204 | 0.657 |
| CX77 | 0.000 | 0.776 | 0.000 | 0.000 | 0.206 | 0.018 | 0.000 |
| CX78 | 0.000 | 0.212 | 0.000 | 0.000 | 0.788 | 0.000 | 0.000 |
| CX79 | 0.000 | 0.000 | 0.000 | 0.000 | 0.000 | 0.284 | 0.716 |
| CX80 | 0.000 | 0.000 | 0.000 | 0.227 | 0.000 | 0.182 | 0.591 |
| CX82 | 0.000 | 0.000 | 0.000 | 0.069 | 0.000 | 0.252 | 0.678 |
| CX83 | 0.000 | 0.000 | 0.000 | 0.105 | 0.000 | 0.144 | 0.751 |
| CX84 | 0.000 | 0.000 | 0.000 | 0.000 | 0.000 | 0.342 | 0.658 |
| CX85 | 0.004 | 0.000 | 0.000 | 0.000 | 0.000 | 0.336 | 0.660 |
| CX86 | 0.000 | 0.000 | 0.000 | 0.000 | 0.021 | 0.404 | 0.576 |
| CX87 | 0.000 | 0.000 | 0.000 | 0.000 | 0.000 | 0.514 | 0.486 |
| CX88 | 0.000 | 0.000 | 0.000 | 0.000 | 0.000 | 1.000 | 0.000 |
| CX89 | 0.000 | 0.051 | 0.000 | 0.000 | 0.000 | 0.415 | 0.534 |
| CX90 | 0.000 | 0.161 | 0.000 | 0.000 | 0.000 | 0.039 | 0.800 |
| CX91 | 0.000 | 0.000 | 0.000 | 0.000 | 0.000 | 0.157 | 0.843 |
| CX92 | 0.000 | 0.000 | 0.000 | 0.000 | 0.000 | 0.001 | 0.999 |
| CX93 | 0.000 | 0.022 | 0.000 | 0.000 | 0.000 | 0.480 | 0.498 |
| CX94 | 0.000 | 0.000 | 0.000 | 0.617 | 0.000 | 0.005 | 0.378 |
| CX96 | 0.017 | 0.000 | 0.000 | 0.052 | 0.000 | 0.221 | 0.710 |
| CX97 | 0.000 | 0.000 | 0.000 | 0.000 | 0.000 | 0.360 | 0.640 |

| | | | | | | | |
|---|---|---|---|---|---|---|---|
| CX98 | 0.180 | 0.000 | 0.000 | 0.820 | 0.000 | 0.000 | 0.000 |
| CX99 | 0.131 | 0.000 | 0.012 | 0.681 | 0.000 | 0.176 | 0.000 |
| CX100 | 0.000 | 0.169 | 0.000 | 0.000 | 0.000 | 0.038 | 0.793 |
| CX101 | 0.000 | 0.002 | 0.000 | 0.000 | 0.005 | 0.928 | 0.066 |
| CX102 | 0.000 | 0.000 | 0.000 | 0.319 | 0.000 | 0.681 | 0.000 |
| CX103 | 0.100 | 0.000 | 0.000 | 0.451 | 0.264 | 0.185 | 0.000 |
| CX104 | 0.000 | 0.000 | 1.000 | 0.000 | 0.000 | 0.000 | 0.000 |
| CX106 | 0.000 | 1.000 | 0.000 | 0.000 | 0.000 | 0.000 | 0.000 |
| CX107 | 0.000 | 0.010 | 0.000 | 0.947 | 0.000 | 0.044 | 0.000 |
| CX108 | 0.539 | 0.000 | 0.000 | 0.000 | 0.038 | 0.109 | 0.314 |
| CX109 | 0.018 | 0.360 | 0.000 | 0.084 | 0.538 | 0.000 | 0.000 |
| CX110 | 0.000 | 0.000 | 0.964 | 0.000 | 0.036 | 0.000 | 0.000 |
| CX111 | 0.000 | 1.000 | 0.000 | 0.000 | 0.000 | 0.000 | 0.000 |
| CX112 | 0.206 | 0.256 | 0.538 | 0.000 | 0.000 | 0.000 | 0.000 |
| CX113 | 0.000 | 1.000 | 0.000 | 0.000 | 0.000 | 0.000 | 0.000 |
| CX114 | 0.000 | 0.000 | 0.000 | 0.000 | 0.000 | 1.000 | 0.000 |
| CX115 | 0.061 | 0.000 | 0.000 | 0.344 | 0.095 | 0.501 | 0.000 |
| CX116 | 0.000 | 0.000 | 0.000 | 0.000 | 1.000 | 0.000 | 0.000 |
| CX117 | 0.000 | 0.000 | 0.000 | 0.000 | 0.000 | 0.623 | 0.377 |
| CX118 | 0.025 | 0.000 | 0.000 | 0.000 | 0.000 | 0.614 | 0.362 |
| CX119 | 0.000 | 0.000 | 0.000 | 0.000 | 0.000 | 0.573 | 0.427 |
| CX120 | 0.000 | 0.098 | 0.000 | 0.170 | 0.000 | 0.000 | 0.731 |
| CX121 | 0.000 | 0.000 | 0.000 | 0.014 | 0.000 | 0.219 | 0.767 |
| CX122 | 0.000 | 0.020 | 0.002 | 0.000 | 0.000 | 0.353 | 0.626 |
| CX123 | 0.000 | 0.000 | 0.000 | 0.000 | 0.000 | 1.000 | 0.000 |
| CX124 | 0.000 | 0.122 | 0.000 | 0.301 | 0.000 | 0.101 | 0.476 |
| CX125 | 0.000 | 0.000 | 0.000 | 0.000 | 0.000 | 0.002 | 0.998 |
| CX126 | 0.000 | 0.000 | 0.000 | 0.000 | 0.000 | 0.004 | 0.996 |
| CX128 | 0.000 | 0.000 | 0.000 | 0.434 | 0.005 | 0.306 | 0.254 |
| CX129 | 0.000 | 0.844 | 0.000 | 0.000 | 0.108 | 0.047 | 0.000 |
| CX130 | 0.071 | 0.000 | 0.000 | 0.793 | 0.064 | 0.072 | 0.000 |
| CX131 | 0.000 | 0.000 | 0.000 | 0.000 | 0.000 | 1.000 | 0.000 |
| CX132 | 0.000 | 1.000 | 0.000 | 0.000 | 0.000 | 0.000 | 0.000 |
| CX133 | 0.000 | 0.000 | 0.000 | 0.000 | 0.030 | 0.970 | 0.000 |
| CX134 | 0.000 | 0.000 | 0.000 | 0.000 | 0.003 | 0.000 | 0.997 |
| CX138 | 0.000 | 0.000 | 0.000 | 0.000 | 0.820 | 0.180 | 0.000 |
| CX139 | 0.017 | 0.356 | 0.000 | 0.083 | 0.544 | 0.000 | 0.000 |
| CX140 | 0.000 | 0.000 | 0.000 | 0.000 | 1.000 | 0.000 | 0.000 |
| CX141 | 0.000 | 0.000 | 0.000 | 0.449 | 0.003 | 0.000 | 0.549 |
| CX142 | 0.000 | 0.322 | 0.000 | 0.000 | 0.678 | 0.000 | 0.000 |
| CX143 | 0.000 | 0.000 | 1.000 | 0.000 | 0.000 | 0.000 | 0.000 |
| CX144 | 0.000 | 0.000 | 0.000 | 0.284 | 0.004 | 0.287 | 0.425 |
| CX145 | 0.000 | 0.000 | 0.000 | 0.196 | 0.000 | 0.000 | 0.804 |
| CX146 | 0.000 | 0.000 | 0.000 | 0.445 | 0.000 | 0.000 | 0.555 |
| CX147 | 0.000 | 0.000 | 0.000 | 0.204 | 0.000 | 0.000 | 0.796 |
| CX148 | 0.000 | 0.000 | 0.000 | 0.332 | 0.000 | 0.000 | 0.668 |
| CX149 | 0.117 | 0.000 | 0.883 | 0.000 | 0.000 | 0.000 | 0.000 |
| CX150 | 0.000 | 0.000 | 0.000 | 0.000 | 0.000 | 0.017 | 0.983 |
| CX151 | 0.000 | 1.000 | 0.000 | 0.000 | 0.000 | 0.000 | 0.000 |
| CX152 | 0.000 | 0.000 | 0.000 | 0.000 | 0.000 | 0.000 | 1.000 |
| CX153 | 0.047 | 0.000 | 0.000 | 0.898 | 0.054 | 0.000 | 0.000 |
| CX154 | 0.000 | 0.000 | 0.000 | 0.981 | 0.017 | 0.000 | 0.002 |
| CX155 | 0.156 | 0.000 | 0.000 | 0.844 | 0.000 | 0.000 | 0.000 |
| CX156 | 0.000 | 0.000 | 0.000 | 0.169 | 0.000 | 0.000 | 0.831 |
| CX157 | 0.150 | 0.006 | 0.018 | 0.825 | 0.002 | 0.000 | 0.000 |
| CX158 | 0.000 | 0.000 | 0.000 | 0.656 | 0.000 | 0.000 | 0.344 |
| CX160 | 0.244 | 0.007 | 0.003 | 0.565 | 0.005 | 0.176 | 0.000 |
| CX161 | 0.042 | 0.000 | 0.000 | 0.000 | 0.000 | 0.184 | 0.774 |
| CX162 | 0.000 | 0.000 | 0.000 | 0.000 | 0.000 | 1.000 | 0.000 |
| CX165 | 0.000 | 0.000 | 0.000 | 0.000 | 1.000 | 0.000 | 0.000 |
| CX182 | 0.000 | 0.000 | 0.000 | 0.689 | 0.002 | 0.309 | 0.000 |
| CX205 | 0.000 | 0.505 | 0.000 | 0.000 | 0.425 | 0.070 | 0.000 |
| CX206 | 0.000 | 0.000 | 0.000 | 0.000 | 0.000 | 0.113 | 0.888 |
| CX207 | 0.000 | 0.000 | 0.000 | 0.138 | 0.025 | 0.486 | 0.351 |
| CX210 | 0.000 | 0.000 | 0.000 | 0.000 | 0.402 | 0.236 | 0.362 |
| CX211 | 0.000 | 0.000 | 0.000 | 0.000 | 1.000 | 0.000 | 0.000 |
| CX212 | 0.000 | 0.000 | 0.000 | 0.000 | 1.000 | 0.000 | 0.000 |

| | | | | | | | |
|---|---|---|---|---|---|---|---|
| CX213 | 0.000 | 0.147 | 0.000 | 0.000 | 0.811 | 0.000 | 0.042 |
| CX214 | 0.000 | 1.000 | 0.000 | 0.000 | 0.000 | 0.000 | 0.000 |
| CX218 | 0.000 | 0.000 | 0.000 | 0.000 | 0.023 | 0.000 | 0.977 |
| CX219 | 0.029 | 0.000 | 0.000 | 0.194 | 0.000 | 0.264 | 0.513 |
| CX220 | 0.000 | 0.906 | 0.035 | 0.000 | 0.055 | 0.000 | 0.004 |
| CX221 | 0.000 | 0.000 | 0.000 | 0.014 | 0.000 | 0.725 | 0.261 |
| CX225 | 0.027 | 0.000 | 0.024 | 0.000 | 0.000 | 0.045 | 0.904 |
| CX226 | 0.000 | 0.010 | 0.000 | 0.000 | 0.000 | 0.045 | 0.945 |
| CX227 | 0.649 | 0.000 | 0.000 | 0.000 | 0.000 | 0.000 | 0.351 |
| CX228 | 0.447 | 0.000 | 0.000 | 0.000 | 0.000 | 0.553 | 0.000 |
| CX230 | 0.000 | 0.000 | 0.000 | 0.000 | 0.000 | 0.161 | 0.839 |
| CX231 | 0.000 | 0.000 | 0.000 | 0.369 | 0.000 | 0.042 | 0.590 |
| CX232 | 0.000 | 0.016 | 0.000 | 0.000 | 0.000 | 0.000 | 0.984 |
| CX233 | 0.000 | 0.000 | 0.000 | 0.241 | 0.000 | 0.088 | 0.671 |
| CX234 | 0.051 | 0.000 | 0.000 | 0.000 | 0.000 | 0.077 | 0.872 |
| CX235 | 0.000 | 0.000 | 0.000 | 0.000 | 0.000 | 0.000 | 1.000 |
| CX236 | 0.245 | 0.026 | 0.000 | 0.000 | 0.000 | 0.099 | 0.630 |
| CX237 | 0.025 | 0.000 | 0.031 | 0.106 | 0.000 | 0.000 | 0.838 |
| CX238 | 0.000 | 0.000 | 0.030 | 0.049 | 0.004 | 0.122 | 0.794 |
| CX240 | 0.000 | 0.000 | 0.000 | 0.073 | 0.000 | 0.000 | 0.927 |
| CX241 | 0.000 | 1.000 | 0.000 | 0.000 | 0.000 | 0.000 | 0.000 |
| CX242 | 0.000 | 0.514 | 0.000 | 0.019 | 0.000 | 0.000 | 0.467 |
| CX243 | 0.000 | 0.873 | 0.000 | 0.000 | 0.000 | 0.000 | 0.127 |
| CX247 | 0.000 | 0.045 | 0.000 | 0.000 | 0.000 | 0.366 | 0.589 |
| CX248 | 0.000 | 0.986 | 0.000 | 0.000 | 0.000 | 0.000 | 0.014 |
| CX249 | 0.000 | 0.000 | 0.000 | 0.000 | 0.130 | 0.112 | 0.759 |
| CX250 | 0.000 | 0.000 | 0.000 | 0.000 | 0.005 | 0.174 | 0.822 |
| CX251 | 0.000 | 0.000 | 0.000 | 0.000 | 1.000 | 0.000 | 0.000 |
| CX262 | 0.000 | 1.000 | 0.000 | 0.000 | 0.000 | 0.000 | 0.000 |
| CX263 | 0.000 | 0.040 | 0.000 | 0.000 | 0.001 | 0.463 | 0.496 |
| CX265 | 0.000 | 0.022 | 0.000 | 0.000 | 0.027 | 0.579 | 0.372 |
| CX266 | 0.000 | 1.000 | 0.000 | 0.000 | 0.000 | 0.000 | 0.000 |
| CX267 | 0.103 | 0.000 | 0.000 | 0.090 | 0.000 | 0.555 | 0.252 |
| CX268 | 0.000 | 0.000 | 0.000 | 0.023 | 0.042 | 0.445 | 0.491 |
| CX269 | 0.000 | 1.000 | 0.000 | 0.000 | 0.000 | 0.000 | 0.000 |
| CX270 | 0.000 | 0.082 | 0.000 | 0.000 | 0.000 | 0.813 | 0.105 |
| CX273 | 0.000 | 0.000 | 0.000 | 0.377 | 0.000 | 0.035 | 0.588 |
| CX274 | 0.000 | 0.000 | 0.000 | 0.375 | 0.000 | 0.038 | 0.588 |
| CX275 | 0.000 | 0.000 | 0.000 | 0.355 | 0.000 | 0.059 | 0.586 |
| CX276 | 0.000 | 0.000 | 0.000 | 0.201 | 0.000 | 0.112 | 0.687 |
| CX277 | 0.000 | 0.000 | 0.000 | 0.000 | 0.852 | 0.148 | 0.000 |
| CX278 | 0.008 | 0.000 | 0.211 | 0.000 | 0.022 | 0.316 | 0.442 |
| CX280 | 0.000 | 0.561 | 0.000 | 0.000 | 0.000 | 0.081 | 0.358 |
| CX281 | 0.000 | 0.000 | 0.000 | 0.000 | 0.000 | 0.224 | 0.776 |
| CX282 | 0.000 | 0.000 | 0.000 | 0.000 | 0.665 | 0.000 | 0.335 |
| CX284 | 0.000 | 0.000 | 0.000 | 0.000 | 0.695 | 0.192 | 0.113 |
| CX285 | 0.000 | 0.693 | 0.000 | 0.000 | 0.256 | 0.000 | 0.051 |
| CX286 | 0.000 | 0.000 | 0.000 | 0.000 | 0.659 | 0.000 | 0.341 |
| CX287 | 0.000 | 0.000 | 0.000 | 0.000 | 0.875 | 0.000 | 0.125 |
| CX288 | 0.000 | 0.046 | 0.000 | 0.000 | 0.027 | 0.568 | 0.359 |
| CX290 | 0.000 | 0.000 | 0.000 | 0.000 | 0.000 | 0.151 | 0.849 |
| CX291 | 0.000 | 0.053 | 0.000 | 0.173 | 0.000 | 0.000 | 0.774 |
| CX296 | 0.014 | 0.000 | 0.000 | 0.000 | 0.000 | 0.301 | 0.685 |
| CX303 | 0.017 | 0.000 | 0.000 | 0.000 | 0.052 | 0.247 | 0.684 |
| CX304 | 0.000 | 0.000 | 0.000 | 0.000 | 0.000 | 0.000 | 1.000 |
| CX305 | 0.000 | 0.000 | 0.000 | 0.000 | 0.000 | 0.154 | 0.846 |
| CX306 | 0.000 | 0.000 | 0.000 | 0.000 | 1.000 | 0.000 | 0.000 |
| CX307 | 0.005 | 0.264 | 0.000 | 0.000 | 0.626 | 0.033 | 0.073 |
| CX313 | 0.000 | 0.065 | 0.000 | 0.127 | 0.000 | 0.154 | 0.654 |
| CX314 | 0.000 | 0.027 | 0.000 | 0.000 | 0.000 | 0.306 | 0.667 |
| CX315 | 0.000 | 0.000 | 0.000 | 0.000 | 0.955 | 0.045 | 0.001 |
| CX316 | 0.000 | 0.000 | 0.000 | 0.000 | 0.571 | 0.000 | 0.429 |
| CX317 | 0.000 | 0.000 | 0.000 | 0.000 | 0.785 | 0.000 | 0.215 |
| CX318 | 0.000 | 0.000 | 0.000 | 0.000 | 0.000 | 0.522 | 0.478 |
| CX319 | 0.017 | 0.000 | 0.000 | 0.684 | 0.000 | 0.299 | 0.000 |
| CX328 | 0.000 | 0.000 | 0.000 | 0.000 | 0.009 | 0.812 | 0.179 |
| CX329 | 0.000 | 0.000 | 0.000 | 0.000 | 1.000 | 0.000 | 0.000 |

| | | | | | | | |
|---|---|---|---|---|---|---|---|
| CX330 | 0.000 | 0.000 | 0.000 | 0.000 | 1.000 | 0.000 | 0.000 |
| CX340 | 0.000 | 0.000 | 0.000 | 0.000 | 0.039 | 0.182 | 0.779 |
| CX341 | 0.000 | 0.000 | 0.000 | 0.025 | 0.000 | 0.392 | 0.582 |
| CX342 | 0.000 | 0.000 | 0.000 | 0.000 | 0.044 | 0.206 | 0.750 |
| CX343 | 0.000 | 0.133 | 0.000 | 0.000 | 0.029 | 0.186 | 0.653 |
| CX344 | 0.000 | 0.026 | 0.000 | 0.000 | 0.958 | 0.000 | 0.016 |
| CX345 | 0.000 | 0.059 | 0.000 | 0.000 | 0.941 | 0.000 | 0.000 |
| CX346 | 0.000 | 0.000 | 0.000 | 0.000 | 0.000 | 0.485 | 0.515 |
| CX347 | 0.000 | 0.000 | 0.000 | 0.000 | 0.075 | 0.319 | 0.606 |
| CX348 | 0.000 | 0.000 | 0.000 | 0.000 | 0.000 | 0.299 | 0.701 |
| CX349 | 0.000 | 0.000 | 0.000 | 0.122 | 0.000 | 0.466 | 0.412 |
| CX350 | 0.000 | 0.000 | 0.000 | 0.000 | 0.974 | 0.026 | 0.000 |
| CX351 | 0.000 | 0.000 | 0.000 | 0.119 | 0.881 | 0.000 | 0.000 |
| CX352 | 0.000 | 0.811 | 0.000 | 0.000 | 0.089 | 0.000 | 0.100 |
| CX353 | 0.000 | 0.365 | 0.000 | 0.070 | 0.503 | 0.004 | 0.058 |
| CX354 | 0.000 | 0.255 | 0.000 | 0.000 | 0.731 | 0.015 | 0.000 |
| CX355 | 0.000 | 0.896 | 0.000 | 0.104 | 0.000 | 0.000 | 0.000 |
| CX356 | 0.000 | 0.000 | 0.000 | 0.000 | 1.000 | 0.000 | 0.000 |
| CX357 | 0.000 | 0.027 | 0.000 | 0.000 | 0.001 | 0.283 | 0.689 |
| CX358 | 0.000 | 0.000 | 0.000 | 0.000 | 0.000 | 0.126 | 0.874 |
| CX359 | 0.000 | 0.948 | 0.000 | 0.000 | 0.000 | 0.000 | 0.052 |
| CX360 | 0.000 | 0.000 | 0.000 | 0.000 | 0.000 | 0.352 | 0.648 |
| CX361 | 0.000 | 0.000 | 0.000 | 0.000 | 0.011 | 0.329 | 0.659 |
| CX362 | 0.000 | 0.051 | 0.000 | 0.000 | 0.000 | 0.119 | 0.830 |
| CX363 | 0.000 | 0.042 | 0.000 | 0.000 | 0.000 | 0.145 | 0.813 |
| CX364 | 0.016 | 0.000 | 0.000 | 0.037 | 0.002 | 0.155 | 0.789 |
| CX365 | 0.000 | 0.000 | 0.042 | 0.000 | 0.338 | 0.085 | 0.534 |
| CX366 | 0.000 | 0.000 | 0.000 | 0.000 | 0.010 | 0.330 | 0.661 |
| CX367 | 0.250 | 0.750 | 0.000 | 0.000 | 0.000 | 0.000 | 0.000 |
| CX368 | 1.000 | 0.000 | 0.000 | 0.000 | 0.000 | 0.000 | 0.000 |
| CX369 | 0.000 | 0.006 | 0.000 | 0.000 | 0.000 | 0.000 | 0.994 |
| CX370 | 0.000 | 0.000 | 0.000 | 0.000 | 0.027 | 0.146 | 0.827 |
| CX371 | 0.000 | 0.915 | 0.000 | 0.001 | 0.001 | 0.033 | 0.049 |
| CX372 | 0.013 | 0.853 | 0.014 | 0.000 | 0.040 | 0.000 | 0.080 |
| CX373 | 0.000 | 1.000 | 0.000 | 0.000 | 0.000 | 0.000 | 0.000 |
| CX374 | 0.000 | 1.000 | 0.000 | 0.000 | 0.000 | 0.000 | 0.000 |
| CX375 | 0.000 | 0.000 | 0.000 | 0.000 | 0.000 | 0.087 | 0.913 |
| CX376 | 0.023 | 0.000 | 0.000 | 0.000 | 0.000 | 0.152 | 0.825 |
| CX377 | 0.000 | 0.000 | 0.000 | 0.000 | 0.000 | 0.179 | 0.821 |
| CX378 | 0.000 | 0.000 | 0.000 | 0.000 | 0.000 | 0.340 | 0.660 |
| CX379 | 0.000 | 0.000 | 0.000 | 0.000 | 0.000 | 0.209 | 0.791 |
| CX380 | 0.000 | 0.000 | 0.000 | 0.000 | 0.987 | 0.013 | 0.000 |
| CX381 | 0.000 | 0.000 | 0.000 | 0.000 | 0.000 | 0.234 | 0.766 |
| CX382 | 0.000 | 0.000 | 0.000 | 0.000 | 0.000 | 0.000 | 1.000 |
| CX383 | 0.000 | 0.032 | 0.000 | 0.000 | 0.968 | 0.000 | 0.000 |
| CX384 | 0.000 | 0.000 | 0.000 | 0.000 | 1.000 | 0.000 | 0.000 |
| CX385 | 0.000 | 0.000 | 0.000 | 0.000 | 0.000 | 0.259 | 0.741 |
| CX386 | 0.000 | 0.028 | 0.000 | 0.000 | 0.000 | 0.478 | 0.494 |
| CX387 | 0.000 | 0.000 | 0.000 | 0.030 | 0.000 | 0.152 | 0.818 |
| CX388 | 0.000 | 0.013 | 0.000 | 0.065 | 0.000 | 0.000 | 0.922 |
| CX389 | 0.000 | 0.000 | 0.000 | 0.000 | 0.830 | 0.000 | 0.170 |
| CX390 | 0.000 | 0.005 | 0.000 | 0.330 | 0.010 | 0.654 | 0.000 |
| CX391 | 0.000 | 0.060 | 0.000 | 0.000 | 0.940 | 0.000 | 0.000 |
| CX392 | 0.000 | 0.000 | 0.000 | 0.191 | 0.000 | 0.000 | 0.809 |
| CX393 | 0.000 | 0.000 | 0.000 | 0.000 | 0.000 | 0.424 | 0.576 |
| CX394 | 0.000 | 0.000 | 0.000 | 0.009 | 0.000 | 0.360 | 0.630 |
| CX395 | 0.014 | 0.388 | 0.000 | 0.000 | 0.100 | 0.103 | 0.396 |
| CX396 | 0.000 | 0.000 | 0.000 | 0.000 | 1.000 | 0.000 | 0.000 |
| CX397 | 0.000 | 0.000 | 0.000 | 0.000 | 1.000 | 0.000 | 0.000 |
| CX399 | 0.035 | 0.000 | 0.000 | 0.082 | 0.000 | 0.227 | 0.657 |
| CX400 | 0.081 | 0.174 | 0.028 | 0.186 | 0.236 | 0.000 | 0.295 |
| CX401 | 0.112 | 0.041 | 0.044 | 0.363 | 0.117 | 0.322 | 0.000 |
| CX402 | 0.300 | 0.000 | 0.120 | 0.247 | 0.333 | 0.000 | 0.000 |
| CX403 | 0.000 | 0.000 | 0.000 | 0.000 | 0.000 | 0.171 | 0.829 |
| CX431 | 0.000 | 0.000 | 0.000 | 0.000 | 0.009 | 0.169 | 0.822 |
| CX534 | 0.000 | 0.000 | 0.010 | 0.000 | 0.935 | 0.055 | 0.000 |
| CX542 | 0.173 | 0.000 | 0.000 | 0.000 | 0.072 | 0.166 | 0.588 |

| | | | | | | | |
|---|---|---|---|---|---|---|---|
| CX548 | 0.000 | 0.000 | 0.000 | 0.000 | 0.009 | 0.521 | 0.469 |
| IRIS_313-7620 | 0.000 | 0.000 | 0.000 | 1.000 | 0.000 | 0.000 | 0.000 |
| IRIS_313-7635 | 0.000 | 0.000 | 0.000 | 0.000 | 0.000 | 0.221 | 0.779 |
| IRIS_313-7636 | 0.000 | 0.007 | 0.000 | 0.000 | 0.000 | 0.000 | 0.993 |
| IRIS_313-7638 | 0.140 | 0.041 | 0.000 | 0.693 | 0.000 | 0.126 | 0.000 |
| IRIS_313-7641 | 0.000 | 0.000 | 0.000 | 0.000 | 0.000 | 0.000 | 1.000 |
| IRIS_313-7646 | 0.215 | 0.370 | 0.000 | 0.415 | 0.000 | 0.000 | 0.000 |
| IRIS_313-7650 | 0.371 | 0.000 | 0.000 | 0.572 | 0.058 | 0.000 | 0.000 |
| IRIS_313-7651 | 0.299 | 0.060 | 0.000 | 0.641 | 0.000 | 0.000 | 0.000 |
| IRIS_313-7654 | 0.000 | 0.000 | 0.000 | 0.000 | 0.000 | 0.000 | 1.000 |
| IRIS_313-7664 | 0.000 | 0.000 | 0.000 | 0.011 | 0.000 | 0.246 | 0.743 |
| IRIS_313-7665 | 0.000 | 0.010 | 0.000 | 0.000 | 0.000 | 0.063 | 0.927 |
| IRIS_313-7668 | 0.000 | 0.020 | 0.000 | 0.000 | 0.000 | 0.025 | 0.955 |
| IRIS_313-7681 | 0.001 | 0.000 | 0.000 | 0.999 | 0.000 | 0.000 | 0.000 |
| IRIS_313-7684 | 0.043 | 0.000 | 0.000 | 0.047 | 0.000 | 0.340 | 0.570 |
| IRIS_313-7685 | 0.046 | 0.000 | 0.000 | 0.000 | 0.000 | 0.226 | 0.728 |
| IRIS_313-7688 | 0.000 | 0.000 | 0.000 | 0.000 | 0.000 | 0.185 | 0.815 |
| IRIS_313-7689 | 0.046 | 0.000 | 0.000 | 0.025 | 0.000 | 0.178 | 0.751 |
| IRIS_313-7690 | 0.059 | 0.011 | 0.008 | 0.485 | 0.011 | 0.301 | 0.126 |
| IRIS_313-7691 | 0.000 | 0.000 | 0.000 | 0.000 | 0.000 | 0.146 | 0.854 |
| IRIS_313-7696 | 0.000 | 0.000 | 0.000 | 0.142 | 0.000 | 0.000 | 0.858 |
| IRIS_313-7698 | 0.000 | 0.000 | 0.000 | 0.000 | 0.000 | 0.049 | 0.951 |
| IRIS_313-7699 | 0.000 | 0.024 | 0.000 | 0.000 | 0.000 | 0.000 | 0.976 |
| IRIS_313-7719 | 0.000 | 0.000 | 0.000 | 0.680 | 0.000 | 0.000 | 0.320 |
| IRIS_313-7720 | 0.235 | 0.000 | 0.000 | 0.765 | 0.000 | 0.000 | 0.000 |
| IRIS_313-7722 | 0.136 | 0.448 | 0.000 | 0.416 | 0.000 | 0.000 | 0.000 |
| IRIS_313-7725 | 0.000 | 0.132 | 0.000 | 0.868 | 0.000 | 0.000 | 0.000 |
| IRIS_313-7728 | 0.255 | 0.000 | 0.000 | 0.745 | 0.000 | 0.000 | 0.000 |
| IRIS_313-7736 | 0.076 | 0.000 | 0.010 | 0.914 | 0.000 | 0.000 | 0.000 |
| IRIS_313-7758 | 0.377 | 0.000 | 0.000 | 0.553 | 0.069 | 0.000 | 0.000 |
| IRIS_313-7766 | 0.000 | 0.000 | 0.000 | 0.000 | 0.000 | 0.079 | 0.921 |
| IRIS_313-7769 | 0.000 | 0.023 | 0.000 | 0.714 | 0.000 | 0.262 | 0.000 |
| IRIS_313-7770 | 0.000 | 0.000 | 0.000 | 0.000 | 0.000 | 0.413 | 0.587 |
| IRIS_313-7773 | 0.000 | 0.000 | 0.000 | 0.054 | 0.000 | 0.125 | 0.821 |
| IRIS_313-7778 | 0.000 | 0.000 | 0.000 | 0.684 | 0.000 | 0.316 | 0.000 |
| IRIS_313-7780 | 0.000 | 0.000 | 0.000 | 0.178 | 0.000 | 0.203 | 0.619 |
| IRIS_313-7792 | 0.000 | 0.000 | 0.000 | 0.186 | 0.000 | 0.224 | 0.589 |
| IRIS_313-7793 | 0.027 | 0.000 | 0.000 | 0.973 | 0.000 | 0.000 | 0.000 |
| IRIS_313-7795 | 0.000 | 0.000 | 0.000 | 1.000 | 0.000 | 0.000 | 0.000 |
| IRIS_313-7797 | 0.000 | 0.000 | 0.000 | 0.050 | 0.000 | 0.000 | 0.950 |
| IRIS_313-7799 | 0.316 | 0.027 | 0.000 | 0.657 | 0.000 | 0.000 | 0.000 |
| IRIS_313-7807 | 0.000 | 0.000 | 0.000 | 0.041 | 0.000 | 0.229 | 0.730 |
| IRIS_313-7808 | 0.000 | 0.013 | 0.000 | 0.095 | 0.000 | 0.035 | 0.857 |
| IRIS_313-7809 | 0.014 | 0.000 | 0.030 | 0.000 | 0.000 | 0.142 | 0.814 |
| IRIS_313-7815 | 0.000 | 0.000 | 0.000 | 0.000 | 0.000 | 0.075 | 0.925 |
| IRIS_313-7816 | 0.000 | 0.000 | 0.000 | 0.069 | 0.000 | 0.204 | 0.727 |
| IRIS_313-7819 | 0.000 | 0.000 | 0.000 | 0.000 | 0.000 | 0.041 | 0.959 |
| IRIS_313-7820 | 0.000 | 0.000 | 0.000 | 0.000 | 0.000 | 0.015 | 0.985 |
| IRIS_313-7824 | 0.000 | 0.000 | 0.000 | 0.000 | 0.000 | 0.249 | 0.751 |
| IRIS_313-7826 | 0.000 | 0.000 | 0.000 | 0.000 | 0.000 | 0.218 | 0.782 |
| IRIS_313-7832 | 0.000 | 0.000 | 0.000 | 0.000 | 0.000 | 0.179 | 0.821 |
| IRIS_313-7838 | 0.000 | 1.000 | 0.000 | 0.000 | 0.000 | 0.000 | 0.000 |
| IRIS_313-7850 | 0.004 | 0.690 | 0.000 | 0.000 | 0.295 | 0.010 | 0.000 |
| IRIS_313-7856 | 0.000 | 0.798 | 0.000 | 0.000 | 0.188 | 0.015 | 0.000 |
| IRIS_313-7859 | 0.000 | 0.814 | 0.000 | 0.000 | 0.186 | 0.000 | 0.000 |
| IRIS_313-7863 | 0.000 | 0.929 | 0.000 | 0.000 | 0.031 | 0.028 | 0.012 |
| IRIS_313-7866 | 0.000 | 0.695 | 0.014 | 0.000 | 0.051 | 0.041 | 0.198 |
| IRIS_313-7868 | 0.000 | 0.934 | 0.000 | 0.000 | 0.000 | 0.000 | 0.066 |
| IRIS_313-7870 | 0.000 | 0.886 | 0.000 | 0.000 | 0.114 | 0.000 | 0.000 |
| IRIS_313-7876 | 0.000 | 1.000 | 0.000 | 0.000 | 0.000 | 0.000 | 0.000 |
| IRIS_313-7883 | 0.000 | 0.936 | 0.000 | 0.000 | 0.064 | 0.000 | 0.000 |
| IRIS_313-7885 | 0.000 | 0.906 | 0.000 | 0.000 | 0.079 | 0.015 | 0.000 |
| IRIS_313-7902 | 0.000 | 0.962 | 0.000 | 0.026 | 0.000 | 0.007 | 0.005 |
| IRIS_313-7907 | 0.000 | 0.830 | 0.000 | 0.000 | 0.047 | 0.000 | 0.123 |
| IRIS_313-7909 | 0.000 | 0.859 | 0.000 | 0.137 | 0.004 | 0.000 | 0.000 |
| IRIS_313-7911 | 0.000 | 0.000 | 0.000 | 0.000 | 0.000 | 0.000 | 1.000 |
| IRIS_313-7912 | 0.000 | 1.000 | 0.000 | 0.000 | 0.000 | 0.000 | 0.000 |

| | | | | | | | |
|---|---|---|---|---|---|---|---|
| IRIS_313-7914 | 0.000 | 1.000 | 0.000 | 0.000 | 0.000 | 0.000 | 0.000 |
| IRIS_313-7922 | 0.000 | 1.000 | 0.000 | 0.000 | 0.000 | 0.000 | 0.000 |
| IRIS_313-7924 | 0.026 | 0.781 | 0.000 | 0.193 | 0.000 | 0.000 | 0.000 |
| IRIS_313-7933 | 0.045 | 0.310 | 0.365 | 0.000 | 0.280 | 0.000 | 0.000 |
| IRIS_313-7959 | 0.030 | 0.940 | 0.000 | 0.029 | 0.000 | 0.000 | 0.000 |
| IRIS_313-7992 | 0.006 | 0.994 | 0.000 | 0.000 | 0.000 | 0.000 | 0.000 |
| IRIS_313-7993 | 0.055 | 0.903 | 0.000 | 0.000 | 0.000 | 0.000 | 0.043 |
| IRIS_313-7994 | 0.000 | 0.997 | 0.000 | 0.000 | 0.003 | 0.000 | 0.000 |
| IRIS_313-8003 | 0.000 | 1.000 | 0.000 | 0.000 | 0.000 | 0.000 | 0.000 |
| IRIS_313-8010 | 0.000 | 0.966 | 0.000 | 0.031 | 0.000 | 0.002 | 0.000 |
| IRIS_313-8011 | 0.000 | 0.477 | 0.000 | 0.000 | 0.523 | 0.000 | 0.000 |
| IRIS_313-8023 | 0.000 | 0.449 | 0.000 | 0.000 | 0.551 | 0.000 | 0.000 |
| IRIS_313-8024 | 0.000 | 0.295 | 0.000 | 0.000 | 0.705 | 0.000 | 0.000 |
| IRIS_313-8025 | 0.000 | 0.318 | 0.000 | 0.000 | 0.682 | 0.000 | 0.000 |
| IRIS_313-8026 | 0.000 | 0.296 | 0.000 | 0.000 | 0.704 | 0.000 | 0.000 |
| IRIS_313-8027 | 0.000 | 0.497 | 0.000 | 0.000 | 0.503 | 0.000 | 0.000 |
| IRIS_313-8029 | 0.000 | 0.529 | 0.000 | 0.000 | 0.471 | 0.000 | 0.000 |
| IRIS_313-8031 | 0.000 | 0.339 | 0.000 | 0.000 | 0.661 | 0.000 | 0.000 |
| IRIS_313-8032 | 0.000 | 0.297 | 0.000 | 0.000 | 0.703 | 0.000 | 0.000 |
| IRIS_313-8033 | 0.000 | 0.115 | 0.000 | 0.000 | 0.885 | 0.000 | 0.000 |
| IRIS_313-8037 | 0.000 | 0.493 | 0.000 | 0.000 | 0.507 | 0.000 | 0.000 |
| IRIS_313-8039 | 0.000 | 0.322 | 0.000 | 0.000 | 0.678 | 0.000 | 0.000 |
| IRIS_313-8041 | 0.000 | 0.597 | 0.000 | 0.000 | 0.403 | 0.000 | 0.000 |
| IRIS_313-8044 | 0.000 | 0.921 | 0.000 | 0.000 | 0.053 | 0.025 | 0.001 |
| IRIS_313-8046 | 0.037 | 0.933 | 0.000 | 0.000 | 0.030 | 0.000 | 0.000 |
| IRIS_313-8048 | 0.000 | 0.336 | 0.000 | 0.000 | 0.664 | 0.000 | 0.000 |
| IRIS_313-8049 | 0.000 | 0.423 | 0.000 | 0.000 | 0.577 | 0.000 | 0.000 |
| IRIS_313-8050 | 0.000 | 0.382 | 0.000 | 0.000 | 0.618 | 0.000 | 0.000 |
| IRIS_313-8052 | 0.000 | 0.293 | 0.000 | 0.000 | 0.707 | 0.000 | 0.000 |
| IRIS_313-8053 | 0.000 | 0.418 | 0.000 | 0.000 | 0.582 | 0.000 | 0.000 |
| IRIS_313-8057 | 0.007 | 0.784 | 0.000 | 0.064 | 0.105 | 0.029 | 0.013 |
| IRIS_313-8058 | 0.000 | 0.840 | 0.000 | 0.000 | 0.058 | 0.000 | 0.102 |
| IRIS_313-8060 | 0.010 | 0.704 | 0.000 | 0.000 | 0.195 | 0.000 | 0.091 |
| IRIS_313-8061 | 0.000 | 0.747 | 0.000 | 0.000 | 0.208 | 0.045 | 0.000 |
| IRIS_313-8062 | 0.005 | 0.775 | 0.000 | 0.027 | 0.174 | 0.010 | 0.010 |
| IRIS_313-8063 | 0.000 | 0.751 | 0.000 | 0.000 | 0.249 | 0.000 | 0.000 |
| IRIS_313-8064 | 0.000 | 0.702 | 0.000 | 0.000 | 0.265 | 0.000 | 0.033 |
| IRIS_313-8065 | 0.000 | 0.683 | 0.000 | 0.000 | 0.317 | 0.000 | 0.000 |
| IRIS_313-8066 | 0.000 | 0.109 | 0.000 | 0.000 | 0.891 | 0.000 | 0.000 |
| IRIS_313-8067 | 0.000 | 0.212 | 0.000 | 0.000 | 0.030 | 0.029 | 0.730 |
| IRIS_313-8068 | 0.000 | 0.001 | 0.000 | 0.015 | 0.000 | 0.049 | 0.935 |
| IRIS_313-8069 | 0.000 | 0.022 | 0.000 | 0.092 | 0.000 | 0.000 | 0.886 |
| IRIS_313-8072 | 0.000 | 1.000 | 0.000 | 0.000 | 0.000 | 0.000 | 0.000 |
| IRIS_313-8073 | 1.000 | 0.000 | 0.000 | 0.000 | 0.000 | 0.000 | 0.000 |
| IRIS_313-8074 | 0.000 | 0.828 | 0.000 | 0.000 | 0.172 | 0.000 | 0.000 |
| IRIS_313-8075 | 0.000 | 0.205 | 0.000 | 0.000 | 0.795 | 0.000 | 0.000 |
| IRIS_313-8076 | 0.005 | 0.646 | 0.000 | 0.000 | 0.348 | 0.000 | 0.000 |
| IRIS_313-8084 | 0.000 | 0.147 | 0.000 | 0.000 | 0.853 | 0.000 | 0.000 |
| IRIS_313-8085 | 0.000 | 0.116 | 0.000 | 0.000 | 0.884 | 0.000 | 0.000 |
| IRIS_313-8087 | 0.000 | 0.000 | 0.000 | 0.000 | 1.000 | 0.000 | 0.000 |
| IRIS_313-8090 | 0.000 | 0.221 | 0.000 | 0.000 | 0.779 | 0.000 | 0.000 |
| IRIS_313-8095 | 0.000 | 0.226 | 0.000 | 0.000 | 0.774 | 0.000 | 0.000 |
| IRIS_313-8096 | 0.000 | 0.196 | 0.000 | 0.000 | 0.804 | 0.000 | 0.000 |
| IRIS_313-8097 | 0.000 | 0.144 | 0.000 | 0.000 | 0.812 | 0.044 | 0.000 |
| IRIS_313-8099 | 0.000 | 0.239 | 0.000 | 0.000 | 0.761 | 0.000 | 0.000 |
| IRIS_313-8102 | 0.000 | 1.000 | 0.000 | 0.000 | 0.000 | 0.000 | 0.000 |
| IRIS_313-8105 | 0.000 | 0.117 | 0.000 | 0.000 | 0.883 | 0.000 | 0.000 |
| IRIS_313-8109 | 0.000 | 0.000 | 0.000 | 0.000 | 1.000 | 0.000 | 0.000 |
| IRIS_313-8111 | 0.000 | 0.000 | 0.000 | 0.000 | 1.000 | 0.000 | 0.000 |
| IRIS_313-8112 | 0.000 | 0.206 | 0.000 | 0.000 | 0.794 | 0.000 | 0.000 |
| IRIS_313-8113 | 0.000 | 0.000 | 0.000 | 0.000 | 1.000 | 0.000 | 0.000 |
| IRIS_313-8114 | 0.002 | 0.362 | 0.000 | 0.005 | 0.631 | 0.000 | 0.000 |
| IRIS_313-8115 | 0.000 | 0.428 | 0.000 | 0.000 | 0.572 | 0.000 | 0.000 |
| IRIS_313-8116 | 0.000 | 0.353 | 0.000 | 0.000 | 0.631 | 0.017 | 0.000 |
| IRIS_313-8118 | 0.000 | 0.336 | 0.000 | 0.000 | 0.647 | 0.000 | 0.017 |
| IRIS_313-8119 | 0.000 | 0.000 | 0.000 | 0.000 | 1.000 | 0.000 | 0.000 |
| IRIS_313-8121 | 0.003 | 0.499 | 0.000 | 0.000 | 0.477 | 0.021 | 0.000 |

| | | | | | | | |
|---|---|---|---|---|---|---|---|
| IRIS_313-8123 | 0.000 | 0.209 | 0.000 | 0.000 | 0.791 | 0.000 | 0.000 |
| IRIS_313-8124 | 0.000 | 0.312 | 0.000 | 0.000 | 0.688 | 0.000 | 0.000 |
| IRIS_313-8125 | 0.000 | 0.194 | 0.000 | 0.000 | 0.806 | 0.000 | 0.000 |
| IRIS_313-8126 | 0.000 | 0.166 | 0.000 | 0.000 | 0.834 | 0.000 | 0.000 |
| IRIS_313-8127 | 0.000 | 0.268 | 0.000 | 0.000 | 0.732 | 0.000 | 0.000 |
| IRIS_313-8128 | 0.000 | 0.000 | 0.000 | 0.000 | 1.000 | 0.000 | 0.000 |
| IRIS_313-8129 | 0.000 | 0.064 | 0.000 | 0.000 | 0.936 | 0.000 | 0.000 |
| IRIS_313-8132 | 0.000 | 0.000 | 0.000 | 0.000 | 1.000 | 0.000 | 0.000 |
| IRIS_313-8134 | 0.001 | 0.098 | 0.000 | 0.000 | 0.901 | 0.000 | 0.000 |
| IRIS_313-8135 | 0.000 | 0.082 | 0.000 | 0.000 | 0.918 | 0.000 | 0.000 |
| IRIS_313-8136 | 0.000 | 0.217 | 0.000 | 0.000 | 0.783 | 0.000 | 0.000 |
| IRIS_313-8137 | 0.003 | 0.324 | 0.000 | 0.008 | 0.659 | 0.005 | 0.000 |
| IRIS_313-8138 | 0.000 | 0.086 | 0.000 | 0.000 | 0.914 | 0.000 | 0.000 |
| IRIS_313-8139 | 0.000 | 0.360 | 0.000 | 0.000 | 0.610 | 0.000 | 0.030 |
| IRIS_313-8140 | 0.000 | 0.000 | 0.000 | 0.000 | 1.000 | 0.000 | 0.000 |
| IRIS_313-8141 | 0.000 | 0.166 | 0.000 | 0.000 | 0.834 | 0.000 | 0.000 |
| IRIS_313-8142 | 0.000 | 0.184 | 0.000 | 0.000 | 0.816 | 0.000 | 0.000 |
| IRIS_313-8143 | 0.000 | 0.424 | 0.000 | 0.000 | 0.576 | 0.000 | 0.000 |
| IRIS_313-8145 | 0.001 | 0.283 | 0.000 | 0.007 | 0.709 | 0.000 | 0.000 |
| IRIS_313-8147 | 0.000 | 0.151 | 0.000 | 0.000 | 0.849 | 0.000 | 0.000 |
| IRIS_313-8148 | 0.000 | 0.000 | 0.000 | 0.000 | 1.000 | 0.000 | 0.000 |
| IRIS_313-8149 | 0.000 | 0.062 | 0.000 | 0.000 | 0.938 | 0.000 | 0.000 |
| IRIS_313-8151 | 0.000 | 0.219 | 0.000 | 0.006 | 0.773 | 0.001 | 0.000 |
| IRIS_313-8154 | 0.000 | 0.416 | 0.000 | 0.000 | 0.584 | 0.000 | 0.000 |
| IRIS_313-8155 | 0.000 | 0.313 | 0.000 | 0.000 | 0.687 | 0.000 | 0.000 |
| IRIS_313-8158 | 0.000 | 0.448 | 0.000 | 0.000 | 0.552 | 0.000 | 0.000 |
| IRIS_313-8159 | 0.000 | 0.448 | 0.000 | 0.000 | 0.552 | 0.000 | 0.000 |
| IRIS_313-8160 | 0.000 | 0.274 | 0.000 | 0.000 | 0.700 | 0.013 | 0.013 |
| IRIS_313-8161 | 0.000 | 0.000 | 0.000 | 0.000 | 0.000 | 0.250 | 0.750 |
| IRIS_313-8162 | 0.000 | 0.000 | 0.000 | 0.000 | 1.000 | 0.000 | 0.000 |
| IRIS_313-8164 | 0.000 | 0.186 | 0.000 | 0.000 | 0.764 | 0.050 | 0.000 |
| IRIS_313-8165 | 0.000 | 0.000 | 0.000 | 0.000 | 1.000 | 0.000 | 0.000 |
| IRIS_313-8166 | 0.000 | 0.337 | 0.000 | 0.000 | 0.663 | 0.000 | 0.000 |
| IRIS_313-8167 | 0.000 | 0.280 | 0.000 | 0.000 | 0.720 | 0.000 | 0.000 |
| IRIS_313-8168 | 0.000 | 0.322 | 0.000 | 0.000 | 0.678 | 0.000 | 0.000 |
| IRIS_313-8170 | 0.000 | 0.456 | 0.000 | 0.000 | 0.544 | 0.000 | 0.000 |
| IRIS_313-8171 | 0.000 | 0.000 | 0.000 | 0.000 | 1.000 | 0.000 | 0.000 |
| IRIS_313-8172 | 0.048 | 0.353 | 0.098 | 0.000 | 0.352 | 0.002 | 0.147 |
| IRIS_313-8173 | 0.046 | 0.784 | 0.054 | 0.000 | 0.054 | 0.061 | 0.000 |
| IRIS_313-8177 | 0.000 | 0.305 | 0.000 | 0.000 | 0.695 | 0.000 | 0.000 |
| IRIS_313-8180 | 0.000 | 0.921 | 0.000 | 0.000 | 0.052 | 0.025 | 0.001 |
| IRIS_313-8183 | 0.000 | 0.924 | 0.006 | 0.000 | 0.044 | 0.025 | 0.000 |
| IRIS_313-8184 | 0.102 | 0.320 | 0.261 | 0.000 | 0.061 | 0.000 | 0.255 |
| IRIS_313-8185 | 0.000 | 0.654 | 0.001 | 0.000 | 0.300 | 0.045 | 0.000 |
| IRIS_313-8186 | 0.000 | 0.785 | 0.000 | 0.000 | 0.192 | 0.023 | 0.000 |
| IRIS_313-8192 | 0.000 | 0.854 | 0.000 | 0.000 | 0.060 | 0.078 | 0.007 |
| IRIS_313-8193 | 0.000 | 0.870 | 0.000 | 0.000 | 0.059 | 0.071 | 0.000 |
| IRIS_313-8195 | 0.000 | 0.359 | 0.000 | 0.000 | 0.641 | 0.000 | 0.000 |
| IRIS_313-8200 | 0.000 | 0.290 | 0.000 | 0.000 | 0.701 | 0.009 | 0.000 |
| IRIS_313-8202 | 0.000 | 0.892 | 0.016 | 0.000 | 0.000 | 0.091 | 0.000 |
| IRIS_313-8204 | 0.000 | 0.044 | 0.000 | 0.000 | 0.956 | 0.000 | 0.000 |
| IRIS_313-8205 | 0.000 | 0.523 | 0.000 | 0.000 | 0.450 | 0.008 | 0.019 |
| IRIS_313-8208 | 0.000 | 0.292 | 0.006 | 0.000 | 0.702 | 0.000 | 0.000 |
| IRIS_313-8209 | 0.010 | 0.775 | 0.024 | 0.000 | 0.157 | 0.011 | 0.023 |
| IRIS_313-8212 | 0.038 | 0.000 | 0.020 | 0.000 | 0.000 | 0.318 | 0.623 |
| IRIS_313-8213 | 0.000 | 0.693 | 0.000 | 0.000 | 0.307 | 0.000 | 0.000 |
| IRIS_313-8214 | 0.000 | 0.369 | 0.000 | 0.000 | 0.631 | 0.000 | 0.000 |
| IRIS_313-8215 | 0.000 | 0.000 | 0.000 | 0.000 | 0.026 | 0.626 | 0.347 |
| IRIS_313-8216 | 0.000 | 0.194 | 0.000 | 0.000 | 0.806 | 0.000 | 0.000 |
| IRIS_313-8217 | 0.000 | 0.000 | 0.000 | 0.000 | 1.000 | 0.000 | 0.000 |
| IRIS_313-8218 | 0.000 | 0.000 | 0.000 | 0.000 | 1.000 | 0.000 | 0.000 |
| IRIS_313-8232 | 0.000 | 0.811 | 0.000 | 0.189 | 0.000 | 0.000 | 0.000 |
| IRIS_313-8244 | 0.117 | 0.000 | 0.000 | 0.712 | 0.000 | 0.170 | 0.000 |
| IRIS_313-8252 | 0.983 | 0.000 | 0.017 | 0.000 | 0.000 | 0.000 | 0.000 |
| IRIS_313-8253 | 0.000 | 0.000 | 0.000 | 1.000 | 0.000 | 0.000 | 0.000 |
| IRIS_313-8256 | 0.000 | 0.000 | 0.328 | 0.000 | 0.672 | 0.000 | 0.000 |
| IRIS_313-8265 | 0.030 | 0.000 | 0.000 | 0.372 | 0.000 | 0.389 | 0.209 |

| | | | | | | | |
|---|---|---|---|---|---|---|---|
| IRIS_313-8268 | 0.000 | 0.000 | 1.000 | 0.000 | 0.000 | 0.000 | 0.000 |
| IRIS_313-8277 | 0.000 | 1.000 | 0.000 | 0.000 | 0.000 | 0.000 | 0.000 |
| IRIS_313-8279 | 0.000 | 1.000 | 0.000 | 0.000 | 0.000 | 0.000 | 0.000 |
| IRIS_313-8283 | 0.875 | 0.000 | 0.023 | 0.080 | 0.000 | 0.000 | 0.021 |
| IRIS_313-8285 | 0.000 | 0.907 | 0.000 | 0.000 | 0.000 | 0.035 | 0.058 |
| IRIS_313-8288 | 0.153 | 0.068 | 0.000 | 0.363 | 0.000 | 0.091 | 0.325 |
| IRIS_313-8291 | 0.000 | 0.000 | 0.000 | 0.461 | 0.000 | 0.000 | 0.539 |
| IRIS_313-8292 | 0.000 | 0.008 | 0.000 | 0.830 | 0.000 | 0.162 | 0.000 |
| IRIS_313-8293 | 0.016 | 0.000 | 0.004 | 0.980 | 0.000 | 0.001 | 0.000 |
| IRIS_313-8302 | 0.437 | 0.319 | 0.011 | 0.000 | 0.233 | 0.000 | 0.000 |
| IRIS_313-8303 | 0.049 | 0.000 | 0.000 | 0.951 | 0.000 | 0.000 | 0.000 |
| IRIS_313-8305 | 0.096 | 0.000 | 0.000 | 0.904 | 0.000 | 0.000 | 0.000 |
| IRIS_313-8306 | 0.094 | 0.000 | 0.000 | 0.906 | 0.000 | 0.000 | 0.000 |
| IRIS_313-8312 | 0.000 | 0.000 | 0.000 | 0.443 | 0.000 | 0.000 | 0.557 |
| IRIS_313-8314 | 0.000 | 1.000 | 0.000 | 0.000 | 0.000 | 0.000 | 0.000 |
| IRIS_313-8315 | 0.000 | 0.000 | 0.000 | 0.809 | 0.000 | 0.191 | 0.000 |
| IRIS_313-8316 | 0.000 | 0.001 | 0.000 | 0.468 | 0.001 | 0.000 | 0.529 |
| IRIS_313-8321 | 0.922 | 0.000 | 0.016 | 0.061 | 0.000 | 0.000 | 0.001 |
| IRIS_313-8323 | 0.000 | 1.000 | 0.000 | 0.000 | 0.000 | 0.000 | 0.000 |
| IRIS_313-8324 | 0.000 | 0.283 | 0.012 | 0.000 | 0.087 | 0.232 | 0.386 |
| IRIS_313-8326 | 0.000 | 0.111 | 0.889 | 0.000 | 0.000 | 0.000 | 0.000 |
| IRIS_313-8332 | 0.051 | 0.000 | 0.000 | 0.579 | 0.000 | 0.370 | 0.000 |
| IRIS_313-8339 | 0.000 | 0.981 | 0.000 | 0.000 | 0.019 | 0.000 | 0.000 |
| IRIS_313-8341 | 0.000 | 0.000 | 0.000 | 0.555 | 0.000 | 0.445 | 0.000 |
| IRIS_313-8342 | 1.000 | 0.000 | 0.000 | 0.000 | 0.000 | 0.000 | 0.000 |
| IRIS_313-8349 | 0.066 | 0.000 | 0.008 | 0.926 | 0.000 | 0.000 | 0.000 |
| IRIS_313-8356 | 0.000 | 1.000 | 0.000 | 0.000 | 0.000 | 0.000 | 0.000 |
| IRIS_313-8368 | 0.000 | 0.000 | 0.000 | 0.129 | 0.000 | 0.000 | 0.871 |
| IRIS_313-8380 | 0.000 | 0.000 | 0.000 | 0.116 | 0.000 | 0.882 | 0.002 |
| IRIS_313-8381 | 0.000 | 0.999 | 0.000 | 0.000 | 0.000 | 0.001 | 0.000 |
| IRIS_313-8382 | 0.498 | 0.065 | 0.000 | 0.000 | 0.000 | 0.115 | 0.323 |
| IRIS_313-8383 | 0.000 | 0.000 | 0.000 | 0.751 | 0.000 | 0.249 | 0.000 |
| IRIS_313-8385 | 0.000 | 0.000 | 0.922 | 0.000 | 0.078 | 0.000 | 0.000 |
| IRIS_313-8386 | 0.000 | 0.000 | 0.000 | 0.676 | 0.000 | 0.324 | 0.000 |
| IRIS_313-8387 | 0.000 | 0.000 | 0.000 | 0.000 | 1.000 | 0.000 | 0.000 |
| IRIS_313-8390 | 1.000 | 0.000 | 0.000 | 0.000 | 0.000 | 0.000 | 0.000 |
| IRIS_313-8391 | 0.034 | 0.000 | 0.000 | 0.966 | 0.000 | 0.000 | 0.000 |
| IRIS_313-8392 | 0.000 | 0.049 | 0.000 | 0.203 | 0.000 | 0.116 | 0.633 |
| IRIS_313-8398 | 1.000 | 0.000 | 0.000 | 0.000 | 0.000 | 0.000 | 0.000 |
| IRIS_313-8399 | 0.000 | 0.056 | 0.000 | 0.000 | 0.944 | 0.000 | 0.000 |
| IRIS_313-8400 | 0.000 | 0.689 | 0.000 | 0.000 | 0.262 | 0.000 | 0.049 |
| IRIS_313-8401 | 0.000 | 0.000 | 0.000 | 0.774 | 0.000 | 0.226 | 0.000 |
| IRIS_313-8405 | 0.000 | 0.000 | 0.001 | 0.517 | 0.068 | 0.414 | 0.000 |
| IRIS_313-8407 | 0.000 | 0.000 | 0.000 | 0.492 | 0.000 | 0.000 | 0.508 |
| IRIS_313-8409 | 0.004 | 0.000 | 0.000 | 0.477 | 0.012 | 0.507 | 0.000 |
| IRIS_313-8410 | 1.000 | 0.000 | 0.000 | 0.000 | 0.000 | 0.000 | 0.000 |
| IRIS_313-8412 | 0.000 | 0.000 | 0.000 | 0.254 | 0.006 | 0.456 | 0.285 |
| IRIS_313-8414 | 0.048 | 0.000 | 0.000 | 0.667 | 0.090 | 0.194 | 0.000 |
| IRIS_313-8431 | 0.000 | 1.000 | 0.000 | 0.000 | 0.000 | 0.000 | 0.000 |
| IRIS_313-8433 | 0.000 | 0.000 | 0.000 | 0.000 | 0.000 | 1.000 | 0.000 |
| IRIS_313-8434 | 0.006 | 0.941 | 0.000 | 0.001 | 0.039 | 0.000 | 0.013 |
| IRIS_313-8435 | 0.000 | 0.000 | 0.000 | 0.142 | 0.000 | 0.858 | 0.000 |
| IRIS_313-8436 | 0.000 | 0.873 | 0.000 | 0.000 | 0.052 | 0.076 | 0.000 |
| IRIS_313-8437 | 0.000 | 0.000 | 0.000 | 1.000 | 0.000 | 0.000 | 0.000 |
| IRIS_313-8439 | 0.000 | 1.000 | 0.000 | 0.000 | 0.000 | 0.000 | 0.000 |
| IRIS_313-8444 | 0.000 | 0.271 | 0.000 | 0.000 | 0.729 | 0.000 | 0.000 |
| IRIS_313-8450 | 0.126 | 0.000 | 0.000 | 0.874 | 0.000 | 0.000 | 0.000 |
| IRIS_313-8453 | 0.007 | 0.000 | 0.000 | 0.780 | 0.010 | 0.203 | 0.000 |
| IRIS_313-8454 | 0.000 | 0.000 | 0.000 | 0.000 | 0.000 | 1.000 | 0.000 |
| IRIS_313-8457 | 0.000 | 0.000 | 0.000 | 0.551 | 0.000 | 0.000 | 0.449 |
| IRIS_313-8458 | 0.061 | 0.000 | 0.000 | 0.557 | 0.000 | 0.000 | 0.382 |
| IRIS_313-8466 | 0.000 | 0.000 | 0.000 | 0.619 | 0.000 | 0.000 | 0.381 |
| IRIS_313-8468 | 0.000 | 0.000 | 0.000 | 0.438 | 0.000 | 0.000 | 0.562 |
| IRIS_313-8474 | 0.000 | 0.000 | 0.000 | 0.962 | 0.000 | 0.038 | 0.000 |
| IRIS_313-8481 | 0.000 | 0.000 | 0.000 | 0.000 | 1.000 | 0.000 | 0.000 |
| IRIS_313-8485 | 0.000 | 0.000 | 0.000 | 1.000 | 0.000 | 0.000 | 0.000 |
| IRIS_313-8486 | 0.000 | 0.516 | 0.066 | 0.000 | 0.419 | 0.000 | 0.000 |

| | | | | | | | |
|---|---|---|---|---|---|---|---|
| IRIS_313-8492 | 0.000 | 0.000 | 0.000 | 0.278 | 0.000 | 0.000 | 0.722 |
| IRIS_313-8493 | 0.000 | 0.000 | 0.000 | 0.275 | 0.000 | 0.000 | 0.725 |
| IRIS_313-8498 | 0.000 | 0.053 | 0.849 | 0.094 | 0.000 | 0.005 | 0.000 |
| IRIS_313-8502 | 0.000 | 0.000 | 0.000 | 0.000 | 0.914 | 0.000 | 0.086 |
| IRIS_313-8509 | 0.106 | 0.000 | 0.000 | 0.842 | 0.000 | 0.052 | 0.000 |
| IRIS_313-8514 | 0.000 | 0.000 | 0.000 | 0.000 | 0.000 | 0.000 | 1.000 |
| IRIS_313-8519 | 0.000 | 0.529 | 0.041 | 0.000 | 0.429 | 0.000 | 0.000 |
| IRIS_313-8523 | 0.000 | 1.000 | 0.000 | 0.000 | 0.000 | 0.000 | 0.000 |
| IRIS_313-8530 | 0.096 | 0.000 | 0.000 | 0.904 | 0.000 | 0.000 | 0.000 |
| IRIS_313-8536 | 0.000 | 0.000 | 0.000 | 0.482 | 0.000 | 0.000 | 0.518 |
| IRIS_313-8554 | 0.968 | 0.032 | 0.000 | 0.000 | 0.000 | 0.000 | 0.000 |
| IRIS_313-8557 | 0.000 | 0.000 | 0.000 | 0.492 | 0.000 | 0.000 | 0.508 |
| IRIS_313-8559 | 0.086 | 0.000 | 0.000 | 0.621 | 0.000 | 0.293 | 0.000 |
| IRIS_313-8565 | 0.000 | 0.627 | 0.011 | 0.029 | 0.332 | 0.000 | 0.000 |
| IRIS_313-8567 | 0.000 | 0.000 | 0.000 | 0.933 | 0.009 | 0.000 | 0.058 |
| IRIS_313-8568 | 0.141 | 0.000 | 0.000 | 0.859 | 0.000 | 0.000 | 0.000 |
| IRIS_313-8571 | 0.000 | 0.000 | 0.000 | 1.000 | 0.000 | 0.000 | 0.000 |
| IRIS_313-8572 | 0.000 | 1.000 | 0.000 | 0.000 | 0.000 | 0.000 | 0.000 |
| IRIS_313-8578 | 0.000 | 0.654 | 0.000 | 0.000 | 0.346 | 0.000 | 0.000 |
| IRIS_313-8580 | 0.000 | 0.615 | 0.000 | 0.000 | 0.385 | 0.000 | 0.000 |
| IRIS_313-8585 | 0.000 | 0.000 | 0.000 | 0.761 | 0.000 | 0.239 | 0.000 |
| IRIS_313-8586 | 0.000 | 0.000 | 0.000 | 1.000 | 0.000 | 0.000 | 0.000 |
| IRIS_313-8591 | 0.000 | 0.000 | 0.000 | 0.358 | 0.000 | 0.000 | 0.642 |
| IRIS_313-8594 | 0.000 | 1.000 | 0.000 | 0.000 | 0.000 | 0.000 | 0.000 |
| IRIS_313-8595 | 0.259 | 0.099 | 0.000 | 0.642 | 0.000 | 0.000 | 0.000 |
| IRIS_313-8599 | 0.000 | 0.681 | 0.000 | 0.000 | 0.319 | 0.000 | 0.000 |
| IRIS_313-8603 | 0.000 | 0.000 | 0.000 | 0.552 | 0.000 | 0.448 | 0.000 |
| IRIS_313-8606 | 0.000 | 0.000 | 0.000 | 0.286 | 0.000 | 0.000 | 0.714 |
| IRIS_313-8608 | 0.000 | 0.000 | 0.000 | 0.390 | 0.000 | 0.000 | 0.610 |
| IRIS_313-8614 | 0.042 | 0.000 | 0.000 | 0.958 | 0.000 | 0.000 | 0.000 |
| IRIS_313-8616 | 0.000 | 0.000 | 0.000 | 0.000 | 0.000 | 0.185 | 0.815 |
| IRIS_313-8621 | 0.000 | 0.000 | 0.000 | 0.240 | 0.000 | 0.154 | 0.606 |
| IRIS_313-8622 | 0.165 | 0.004 | 0.000 | 0.621 | 0.000 | 0.210 | 0.000 |
| IRIS_313-8626 | 0.000 | 1.000 | 0.000 | 0.000 | 0.000 | 0.000 | 0.000 |
| IRIS_313-8627 | 0.000 | 0.781 | 0.000 | 0.000 | 0.219 | 0.000 | 0.000 |
| IRIS_313-8631 | 0.048 | 0.000 | 0.000 | 0.634 | 0.000 | 0.318 | 0.000 |
| IRIS_313-8632 | 0.083 | 0.000 | 0.026 | 0.891 | 0.000 | 0.000 | 0.000 |
| IRIS_313-8637 | 0.000 | 0.589 | 0.042 | 0.000 | 0.369 | 0.000 | 0.000 |
| IRIS_313-8638 | 0.000 | 0.000 | 0.000 | 0.972 | 0.000 | 0.028 | 0.000 |
| IRIS_313-8641 | 1.000 | 0.000 | 0.000 | 0.000 | 0.000 | 0.000 | 0.000 |
| IRIS_313-8643 | 0.000 | 0.000 | 0.000 | 0.298 | 0.000 | 0.000 | 0.702 |
| IRIS_313-8645 | 0.000 | 0.000 | 0.000 | 0.000 | 0.000 | 1.000 | 0.000 |
| IRIS_313-8647 | 0.091 | 0.000 | 0.000 | 0.658 | 0.000 | 0.251 | 0.000 |
| IRIS_313-8655 | 0.989 | 0.000 | 0.011 | 0.000 | 0.000 | 0.000 | 0.000 |
| IRIS_313-8656 | 0.108 | 0.000 | 0.892 | 0.000 | 0.000 | 0.000 | 0.000 |
| IRIS_313-8657 | 0.113 | 0.000 | 0.000 | 0.742 | 0.000 | 0.000 | 0.146 |
| IRIS_313-8658 | 0.000 | 0.566 | 0.000 | 0.000 | 0.434 | 0.000 | 0.000 |
| IRIS_313-8659 | 0.000 | 0.000 | 0.000 | 0.842 | 0.000 | 0.158 | 0.000 |
| IRIS_313-8660 | 0.126 | 0.000 | 0.000 | 0.670 | 0.000 | 0.204 | 0.000 |
| IRIS_313-8664 | 0.000 | 0.985 | 0.015 | 0.000 | 0.000 | 0.000 | 0.000 |
| IRIS_313-8665 | 0.000 | 0.195 | 0.000 | 0.000 | 0.805 | 0.000 | 0.000 |
| IRIS_313-8669 | 0.005 | 0.783 | 0.000 | 0.000 | 0.212 | 0.000 | 0.000 |
| IRIS_313-8674 | 0.000 | 0.000 | 0.000 | 1.000 | 0.000 | 0.000 | 0.000 |
| IRIS_313-8679 | 0.000 | 0.000 | 0.000 | 1.000 | 0.000 | 0.000 | 0.000 |
| IRIS_313-8681 | 0.000 | 0.000 | 0.000 | 0.228 | 0.000 | 0.000 | 0.772 |
| IRIS_313-8683 | 0.000 | 0.000 | 0.000 | 0.945 | 0.000 | 0.055 | 0.000 |
| IRIS_313-8687 | 0.000 | 1.000 | 0.000 | 0.000 | 0.000 | 0.000 | 0.000 |
| IRIS_313-8690 | 0.000 | 0.000 | 0.047 | 0.000 | 0.897 | 0.055 | 0.000 |
| IRIS_313-8694 | 0.000 | 1.000 | 0.000 | 0.000 | 0.000 | 0.000 | 0.000 |
| IRIS_313-8697 | 0.000 | 0.000 | 0.000 | 0.983 | 0.000 | 0.017 | 0.000 |
| IRIS_313-8699 | 0.088 | 0.015 | 0.000 | 0.645 | 0.000 | 0.057 | 0.195 |
| IRIS_313-8702 | 0.000 | 0.000 | 0.000 | 0.999 | 0.000 | 0.000 | 0.001 |
| IRIS_313-8703 | 0.021 | 0.000 | 0.000 | 0.979 | 0.000 | 0.000 | 0.000 |
| IRIS_313-8704 | 0.169 | 0.000 | 0.000 | 0.831 | 0.000 | 0.000 | 0.000 |
| IRIS_313-8712 | 0.000 | 0.083 | 0.917 | 0.000 | 0.000 | 0.000 | 0.000 |
| IRIS_313-8713 | 0.000 | 0.000 | 0.000 | 0.446 | 0.000 | 0.000 | 0.554 |
| IRIS_313-8717 | 0.043 | 0.000 | 0.000 | 0.957 | 0.000 | 0.000 | 0.000 |

| | | | | | | | |
|---|---|---|---|---|---|---|---|
| IRIS_313-8721 | 0.976 | 0.000 | 0.024 | 0.000 | 0.000 | 0.000 | 0.000 |
| IRIS_313-8722 | 0.000 | 0.000 | 0.000 | 1.000 | 0.000 | 0.000 | 0.000 |
| IRIS_313-8723 | 0.000 | 0.000 | 0.000 | 0.569 | 0.000 | 0.000 | 0.431 |
| IRIS_313-8725 | 0.000 | 0.000 | 0.000 | 0.624 | 0.000 | 0.000 | 0.376 |
| IRIS_313-8727 | 0.105 | 0.000 | 0.000 | 0.711 | 0.003 | 0.181 | 0.000 |
| IRIS_313-8731 | 0.062 | 0.000 | 0.000 | 0.859 | 0.000 | 0.079 | 0.000 |
| IRIS_313-8732 | 0.000 | 0.000 | 0.000 | 0.004 | 0.000 | 0.199 | 0.797 |
| IRIS_313-8733 | 0.073 | 0.000 | 0.007 | 0.920 | 0.000 | 0.000 | 0.000 |
| IRIS_313-8735 | 0.000 | 0.000 | 0.079 | 0.000 | 0.885 | 0.036 | 0.000 |
| IRIS_313-8737 | 0.072 | 0.000 | 0.000 | 0.900 | 0.000 | 0.028 | 0.000 |
| IRIS_313-8739 | 0.298 | 0.150 | 0.000 | 0.479 | 0.000 | 0.073 | 0.000 |
| IRIS_313-8743 | 0.000 | 0.000 | 0.000 | 0.568 | 0.010 | 0.422 | 0.000 |
| IRIS_313-8744 | 0.000 | 0.000 | 0.000 | 0.509 | 0.000 | 0.009 | 0.482 |
| IRIS_313-8745 | 0.000 | 1.000 | 0.000 | 0.000 | 0.000 | 0.000 | 0.000 |
| IRIS_313-8747 | 0.075 | 0.011 | 0.812 | 0.000 | 0.102 | 0.000 | 0.000 |
| IRIS_313-8751 | 0.000 | 0.000 | 0.000 | 0.996 | 0.000 | 0.000 | 0.004 |
| IRIS_313-8754 | 0.058 | 0.000 | 0.000 | 0.815 | 0.000 | 0.126 | 0.000 |
| IRIS_313-8755 | 0.000 | 0.000 | 0.000 | 0.000 | 1.000 | 0.000 | 0.000 |
| IRIS_313-8757 | 0.068 | 0.000 | 0.000 | 0.932 | 0.000 | 0.000 | 0.000 |
| IRIS_313-8765 | 0.000 | 0.000 | 1.000 | 0.000 | 0.000 | 0.000 | 0.000 |
| IRIS_313-8767 | 0.000 | 0.000 | 0.000 | 0.545 | 0.000 | 0.000 | 0.455 |
| IRIS_313-8768 | 0.000 | 1.000 | 0.000 | 0.000 | 0.000 | 0.000 | 0.000 |
| IRIS_313-8769 | 0.000 | 1.000 | 0.000 | 0.000 | 0.000 | 0.000 | 0.000 |
| IRIS_313-8771 | 1.000 | 0.000 | 0.000 | 0.000 | 0.000 | 0.000 | 0.000 |
| IRIS_313-8778 | 0.000 | 1.000 | 0.000 | 0.000 | 0.000 | 0.000 | 0.000 |
| IRIS_313-8781 | 0.000 | 0.082 | 0.000 | 0.396 | 0.000 | 0.000 | 0.522 |
| IRIS_313-8783 | 0.452 | 0.000 | 0.000 | 0.128 | 0.000 | 0.420 | 0.000 |
| IRIS_313-8785 | 0.000 | 0.000 | 0.000 | 1.000 | 0.000 | 0.000 | 0.000 |
| IRIS_313-8789 | 1.000 | 0.000 | 0.000 | 0.000 | 0.000 | 0.000 | 0.000 |
| IRIS_313-8791 | 0.000 | 0.000 | 0.000 | 0.434 | 0.000 | 0.000 | 0.566 |
| IRIS_313-8793 | 0.000 | 0.000 | 0.000 | 0.999 | 0.000 | 0.000 | 0.001 |
| IRIS_313-8796 | 0.044 | 0.000 | 0.000 | 0.910 | 0.000 | 0.046 | 0.000 |
| IRIS_313-8803 | 0.000 | 1.000 | 0.000 | 0.000 | 0.000 | 0.000 | 0.000 |
| IRIS_313-8811 | 0.000 | 0.000 | 0.000 | 0.473 | 0.000 | 0.000 | 0.527 |
| IRIS_313-8812 | 0.000 | 0.013 | 0.000 | 0.273 | 0.000 | 0.000 | 0.714 |
| IRIS_313-8813 | 0.000 | 0.000 | 1.000 | 0.000 | 0.000 | 0.000 | 0.000 |
| IRIS_313-8814 | 0.000 | 0.000 | 1.000 | 0.000 | 0.000 | 0.000 | 0.000 |
| IRIS_313-8815 | 0.000 | 0.507 | 0.049 | 0.000 | 0.444 | 0.000 | 0.000 |
| IRIS_313-8822 | 0.871 | 0.049 | 0.000 | 0.000 | 0.000 | 0.080 | 0.000 |
| IRIS_313-8831 | 0.000 | 0.008 | 0.000 | 0.512 | 0.000 | 0.000 | 0.480 |
| IRIS_313-8833 | 0.000 | 0.000 | 0.000 | 0.503 | 0.000 | 0.000 | 0.497 |
| IRIS_313-8844 | 0.031 | 0.000 | 0.000 | 0.858 | 0.000 | 0.111 | 0.000 |
| IRIS_313-8846 | 0.041 | 0.000 | 0.000 | 0.959 | 0.000 | 0.000 | 0.000 |
| IRIS_313-8850 | 0.151 | 0.000 | 0.000 | 0.803 | 0.000 | 0.046 | 0.000 |
| IRIS_313-8854 | 0.000 | 0.000 | 0.000 | 0.962 | 0.000 | 0.038 | 0.000 |
| IRIS_313-8856 | 0.000 | 0.128 | 0.000 | 0.000 | 0.872 | 0.000 | 0.000 |
| IRIS_313-8857 | 0.000 | 0.529 | 0.045 | 0.000 | 0.426 | 0.000 | 0.000 |
| IRIS_313-8859 | 0.000 | 0.000 | 0.000 | 0.000 | 0.000 | 1.000 | 0.000 |
| IRIS_313-8864 | 1.000 | 0.000 | 0.000 | 0.000 | 0.000 | 0.000 | 0.000 |
| IRIS_313-8865 | 0.000 | 0.989 | 0.000 | 0.011 | 0.000 | 0.000 | 0.000 |
| IRIS_313-8870 | 0.000 | 0.000 | 0.000 | 1.000 | 0.000 | 0.000 | 0.000 |
| IRIS_313-8872 | 0.000 | 0.521 | 0.000 | 0.030 | 0.245 | 0.204 | 0.000 |
| IRIS_313-8873 | 0.000 | 0.000 | 0.000 | 0.352 | 0.613 | 0.000 | 0.035 |
| IRIS_313-8876 | 0.000 | 1.000 | 0.000 | 0.000 | 0.000 | 0.000 | 0.000 |
| IRIS_313-8880 | 0.007 | 0.154 | 0.000 | 0.000 | 0.155 | 0.530 | 0.154 |
| IRIS_313-8883 | 0.000 | 0.000 | 0.072 | 0.000 | 0.870 | 0.058 | 0.000 |
| IRIS_313-8884 | 0.000 | 0.789 | 0.000 | 0.000 | 0.211 | 0.000 | 0.000 |
| IRIS_313-8889 | 0.000 | 0.010 | 0.030 | 0.428 | 0.000 | 0.532 | 0.000 |
| IRIS_313-8890 | 0.000 | 0.000 | 0.007 | 0.000 | 0.993 | 0.000 | 0.000 |
| IRIS_313-8894 | 0.000 | 1.000 | 0.000 | 0.000 | 0.000 | 0.000 | 0.000 |
| IRIS_313-8895 | 0.000 | 0.000 | 0.000 | 1.000 | 0.000 | 0.000 | 0.000 |
| IRIS_313-8900 | 0.054 | 0.000 | 0.000 | 0.946 | 0.000 | 0.000 | 0.000 |
| IRIS_313-8903 | 0.000 | 0.028 | 0.000 | 0.000 | 0.000 | 0.000 | 0.972 |
| IRIS_313-8909 | 0.000 | 0.000 | 0.000 | 1.000 | 0.000 | 0.000 | 0.000 |
| IRIS_313-8911 | 0.053 | 0.458 | 0.000 | 0.396 | 0.000 | 0.004 | 0.089 |
| IRIS_313-8914 | 0.000 | 0.003 | 0.000 | 0.001 | 0.034 | 0.631 | 0.331 |
| IRIS_313-8920 | 0.055 | 0.000 | 0.000 | 0.945 | 0.000 | 0.000 | 0.000 |

| | | | | | | | |
|---|---|---|---|---|---|---|---|
| IRIS_313-8921 | 0.000 | 0.000 | 0.000 | 0.308 | 0.688 | 0.004 | 0.000 |
| IRIS_313-8922 | 0.027 | 0.000 | 0.000 | 0.821 | 0.000 | 0.000 | 0.152 |
| IRIS_313-8923 | 0.000 | 0.792 | 0.000 | 0.000 | 0.208 | 0.000 | 0.000 |
| IRIS_313-8924 | 0.084 | 0.000 | 0.010 | 0.905 | 0.000 | 0.000 | 0.000 |
| IRIS_313-8925 | 0.119 | 0.000 | 0.000 | 0.662 | 0.000 | 0.218 | 0.001 |
| IRIS_313-8927 | 0.000 | 0.396 | 0.000 | 0.000 | 0.604 | 0.000 | 0.000 |
| IRIS_313-8930 | 0.000 | 0.000 | 0.000 | 0.092 | 0.000 | 0.115 | 0.793 |
| IRIS_313-8932 | 0.000 | 0.000 | 0.001 | 0.477 | 0.010 | 0.000 | 0.513 |
| IRIS_313-8935 | 0.000 | 0.130 | 0.000 | 0.000 | 0.036 | 0.000 | 0.834 |
| IRIS_313-8940 | 0.000 | 0.000 | 0.000 | 0.000 | 0.000 | 1.000 | 0.000 |
| IRIS_313-8946 | 0.000 | 0.000 | 0.004 | 0.776 | 0.000 | 0.220 | 0.000 |
| IRIS_313-8948 | 0.000 | 0.000 | 0.000 | 0.840 | 0.000 | 0.124 | 0.036 |
| IRIS_313-8955 | 0.000 | 1.000 | 0.000 | 0.000 | 0.000 | 0.000 | 0.000 |
| IRIS_313-8956 | 0.000 | 0.000 | 0.000 | 0.553 | 0.032 | 0.000 | 0.415 |
| IRIS_313-8957 | 0.025 | 0.000 | 0.000 | 0.975 | 0.000 | 0.000 | 0.000 |
| IRIS_313-8960 | 0.000 | 1.000 | 0.000 | 0.000 | 0.000 | 0.000 | 0.000 |
| IRIS_313-8963 | 1.000 | 0.000 | 0.000 | 0.000 | 0.000 | 0.000 | 0.000 |
| IRIS_313-8967 | 0.000 | 0.000 | 0.000 | 0.308 | 0.000 | 0.228 | 0.465 |
| IRIS_313-8968 | 0.114 | 0.000 | 0.000 | 0.695 | 0.000 | 0.191 | 0.000 |
| IRIS_313-8976 | 0.000 | 1.000 | 0.000 | 0.000 | 0.000 | 0.000 | 0.000 |
| IRIS_313-8978 | 0.000 | 0.000 | 0.000 | 0.590 | 0.000 | 0.000 | 0.410 |
| IRIS_313-8980 | 0.000 | 0.000 | 0.000 | 1.000 | 0.000 | 0.000 | 0.000 |
| IRIS_313-8982 | 0.144 | 0.000 | 0.000 | 0.500 | 0.000 | 0.356 | 0.000 |
| IRIS_313-8985 | 0.000 | 0.000 | 0.000 | 0.992 | 0.000 | 0.008 | 0.000 |
| IRIS_313-8986 | 0.238 | 0.000 | 0.000 | 0.412 | 0.000 | 0.350 | 0.000 |
| IRIS_313-8987 | 0.101 | 0.006 | 0.000 | 0.494 | 0.000 | 0.000 | 0.399 |
| IRIS_313-8988 | 0.069 | 0.000 | 0.000 | 0.931 | 0.000 | 0.000 | 0.000 |
| IRIS_313-8994 | 0.000 | 0.000 | 0.000 | 0.871 | 0.000 | 0.127 | 0.003 |
| IRIS_313-8996 | 0.000 | 0.000 | 0.000 | 0.997 | 0.000 | 0.003 | 0.000 |
| IRIS_313-8998 | 0.000 | 1.000 | 0.000 | 0.000 | 0.000 | 0.000 | 0.000 |
| IRIS_313-8999 | 0.000 | 0.635 | 0.059 | 0.000 | 0.306 | 0.000 | 0.000 |
| IRIS_313-9002 | 0.000 | 0.000 | 0.000 | 0.000 | 1.000 | 0.000 | 0.000 |
| IRIS_313-9005 | 0.000 | 0.000 | 0.000 | 0.504 | 0.000 | 0.000 | 0.496 |
| IRIS_313-9006 | 0.000 | 0.000 | 0.000 | 0.928 | 0.000 | 0.000 | 0.072 |
| IRIS_313-9017 | 0.000 | 0.000 | 0.000 | 0.103 | 0.000 | 0.000 | 0.897 |
| IRIS_313-9019 | 0.000 | 0.000 | 0.000 | 1.000 | 0.000 | 0.000 | 0.000 |
| IRIS_313-9020 | 0.000 | 0.000 | 0.000 | 1.000 | 0.000 | 0.000 | 0.000 |
| IRIS_313-9023 | 0.071 | 0.000 | 0.002 | 0.437 | 0.000 | 0.000 | 0.490 |
| IRIS_313-9032 | 0.000 | 0.029 | 0.000 | 0.000 | 0.000 | 0.000 | 0.971 |
| IRIS_313-9039 | 0.097 | 0.000 | 0.019 | 0.789 | 0.000 | 0.096 | 0.000 |
| IRIS_313-9044 | 0.042 | 0.000 | 0.000 | 0.000 | 0.000 | 0.195 | 0.762 |
| IRIS_313-9048 | 0.000 | 0.598 | 0.048 | 0.000 | 0.354 | 0.000 | 0.000 |
| IRIS_313-9049 | 0.099 | 0.000 | 0.017 | 0.884 | 0.000 | 0.000 | 0.000 |
| IRIS_313-9050 | 0.000 | 1.000 | 0.000 | 0.000 | 0.000 | 0.000 | 0.000 |
| IRIS_313-9053 | 0.000 | 0.371 | 0.523 | 0.000 | 0.106 | 0.000 | 0.000 |
| IRIS_313-9054 | 0.000 | 0.000 | 0.000 | 0.497 | 0.000 | 0.000 | 0.503 |
| IRIS_313-9062 | 0.000 | 0.000 | 0.000 | 0.990 | 0.000 | 0.010 | 0.000 |
| IRIS_313-9065 | 0.000 | 0.000 | 0.000 | 0.000 | 0.000 | 1.000 | 0.000 |
| IRIS_313-9066 | 0.000 | 0.214 | 0.000 | 0.000 | 0.000 | 0.000 | 0.786 |
| IRIS_313-9067 | 0.080 | 0.000 | 0.005 | 0.914 | 0.000 | 0.000 | 0.000 |
| IRIS_313-9070 | 0.000 | 0.000 | 0.000 | 0.512 | 0.000 | 0.000 | 0.488 |
| IRIS_313-9072 | 0.122 | 0.000 | 0.000 | 0.878 | 0.000 | 0.000 | 0.000 |
| IRIS_313-9081 | 0.000 | 0.000 | 0.000 | 0.000 | 1.000 | 0.000 | 0.000 |
| IRIS_313-9083 | 0.000 | 0.000 | 1.000 | 0.000 | 0.000 | 0.000 | 0.000 |
| IRIS_313-9091 | 0.077 | 0.003 | 0.007 | 0.791 | 0.011 | 0.024 | 0.086 |
| IRIS_313-9097 | 0.000 | 0.000 | 0.000 | 0.587 | 0.000 | 0.000 | 0.413 |
| IRIS_313-9098 | 0.000 | 0.000 | 0.000 | 0.456 | 0.000 | 0.000 | 0.544 |
| IRIS_313-9101 | 0.000 | 0.574 | 0.000 | 0.075 | 0.000 | 0.000 | 0.351 |
| IRIS_313-9102 | 0.000 | 0.000 | 0.000 | 0.410 | 0.000 | 0.000 | 0.590 |
| IRIS_313-9108 | 0.042 | 0.011 | 0.049 | 0.300 | 0.013 | 0.584 | 0.000 |
| IRIS_313-9111 | 0.000 | 0.000 | 0.000 | 0.113 | 0.000 | 0.887 | 0.000 |
| IRIS_313-9112 | 0.000 | 0.000 | 0.000 | 0.545 | 0.000 | 0.000 | 0.455 |
| IRIS_313-9114 | 0.000 | 0.000 | 0.000 | 0.679 | 0.000 | 0.000 | 0.321 |
| IRIS_313-9115 | 0.000 | 0.000 | 0.000 | 1.000 | 0.000 | 0.000 | 0.000 |
| IRIS_313-9116 | 0.000 | 0.000 | 0.000 | 1.000 | 0.000 | 0.000 | 0.000 |
| IRIS_313-9117 | 0.000 | 0.000 | 0.000 | 0.415 | 0.000 | 0.000 | 0.585 |
| IRIS_313-9119 | 0.000 | 0.000 | 0.000 | 1.000 | 0.000 | 0.000 | 0.000 |

| | | | | | | | |
|---|---|---|---|---|---|---|---|
| IRIS_313-9120 | 0.000 | 0.000 | 0.480 | 0.502 | 0.000 | 0.000 | 0.017 |
| IRIS_313-9121 | 0.072 | 0.000 | 0.000 | 0.928 | 0.000 | 0.000 | 0.000 |
| IRIS_313-9123 | 0.000 | 0.789 | 0.000 | 0.000 | 0.200 | 0.011 | 0.000 |
| IRIS_313-9129 | 0.000 | 1.000 | 0.000 | 0.000 | 0.000 | 0.000 | 0.000 |
| IRIS_313-9131 | 0.000 | 0.000 | 0.000 | 1.000 | 0.000 | 0.000 | 0.000 |
| IRIS_313-9137 | 0.820 | 0.024 | 0.000 | 0.000 | 0.000 | 0.130 | 0.026 |
| IRIS_313-9139 | 0.097 | 0.000 | 0.000 | 0.903 | 0.000 | 0.000 | 0.000 |
| IRIS_313-9140 | 0.000 | 0.000 | 0.000 | 0.000 | 0.881 | 0.119 | 0.000 |
| IRIS_313-9148 | 0.142 | 0.000 | 0.006 | 0.851 | 0.000 | 0.000 | 0.000 |
| IRIS_313-9156 | 0.051 | 0.000 | 0.000 | 0.949 | 0.000 | 0.000 | 0.000 |
| IRIS_313-9160 | 0.000 | 0.000 | 0.000 | 1.000 | 0.000 | 0.000 | 0.000 |
| IRIS_313-9170 | 0.000 | 0.000 | 0.891 | 0.000 | 0.096 | 0.014 | 0.000 |
| IRIS_313-9172 | 0.000 | 0.066 | 0.934 | 0.000 | 0.000 | 0.000 | 0.000 |
| IRIS_313-9174 | 0.000 | 0.000 | 0.000 | 0.949 | 0.000 | 0.051 | 0.000 |
| IRIS_313-9176 | 0.000 | 0.773 | 0.000 | 0.000 | 0.227 | 0.000 | 0.000 |
| IRIS_313-9182 | 0.000 | 0.020 | 0.000 | 0.751 | 0.000 | 0.228 | 0.000 |
| IRIS_313-9184 | 0.000 | 0.000 | 0.010 | 0.478 | 0.023 | 0.487 | 0.002 |
| IRIS_313-9187 | 0.046 | 0.305 | 0.000 | 0.650 | 0.000 | 0.000 | 0.000 |
| IRIS_313-9188 | 0.000 | 0.000 | 0.000 | 0.450 | 0.000 | 0.000 | 0.550 |
| IRIS_313-9190 | 0.016 | 0.000 | 0.000 | 0.933 | 0.000 | 0.000 | 0.052 |
| IRIS_313-9193 | 0.000 | 0.409 | 0.000 | 0.000 | 0.591 | 0.000 | 0.000 |
| IRIS_313-9197 | 0.000 | 0.025 | 0.000 | 0.813 | 0.000 | 0.035 | 0.128 |
| IRIS_313-9198 | 0.000 | 0.000 | 0.000 | 0.791 | 0.000 | 0.162 | 0.047 |
| IRIS_313-9201 | 0.000 | 0.693 | 0.000 | 0.000 | 0.282 | 0.025 | 0.000 |
| IRIS_313-9204 | 0.000 | 0.000 | 0.000 | 0.090 | 0.000 | 0.910 | 0.000 |
| IRIS_313-9208 | 0.000 | 0.013 | 0.000 | 0.467 | 0.000 | 0.000 | 0.520 |
| IRIS_313-9209 | 0.000 | 0.000 | 0.000 | 0.995 | 0.000 | 0.000 | 0.005 |
| IRIS_313-9210 | 0.000 | 0.000 | 0.000 | 0.389 | 0.000 | 0.000 | 0.611 |
| IRIS_313-9218 | 0.000 | 0.000 | 0.000 | 1.000 | 0.000 | 0.000 | 0.000 |
| IRIS_313-9227 | 0.000 | 0.000 | 0.000 | 0.004 | 0.000 | 0.223 | 0.773 |
| IRIS_313-9228 | 0.000 | 0.000 | 0.000 | 0.000 | 1.000 | 0.000 | 0.000 |
| IRIS_313-9233 | 0.000 | 0.000 | 0.000 | 0.000 | 1.000 | 0.000 | 0.000 |
| IRIS_313-9239 | 0.000 | 0.999 | 0.000 | 0.000 | 0.001 | 0.000 | 0.000 |
| IRIS_313-9242 | 0.000 | 0.471 | 0.000 | 0.529 | 0.000 | 0.000 | 0.000 |
| IRIS_313-9243 | 0.000 | 0.010 | 0.000 | 0.685 | 0.010 | 0.295 | 0.000 |
| IRIS_313-9249 | 0.000 | 0.000 | 0.000 | 0.001 | 0.982 | 0.017 | 0.000 |
| IRIS_313-9251 | 0.000 | 0.000 | 0.000 | 0.425 | 0.000 | 0.000 | 0.575 |
| IRIS_313-9253 | 0.000 | 0.000 | 0.000 | 0.000 | 0.051 | 0.949 | 0.000 |
| IRIS_313-9256 | 0.189 | 0.015 | 0.015 | 0.624 | 0.000 | 0.138 | 0.020 |
| IRIS_313-9258 | 0.074 | 0.000 | 0.000 | 0.926 | 0.000 | 0.000 | 0.000 |
| IRIS_313-9259 | 0.051 | 0.000 | 0.000 | 0.905 | 0.044 | 0.000 | 0.000 |
| IRIS_313-9262 | 0.036 | 0.000 | 0.000 | 0.964 | 0.000 | 0.000 | 0.000 |
| IRIS_313-9267 | 0.000 | 1.000 | 0.000 | 0.000 | 0.000 | 0.000 | 0.000 |
| IRIS_313-9271 | 0.044 | 0.000 | 0.000 | 0.955 | 0.000 | 0.002 | 0.000 |
| IRIS_313-9273 | 0.000 | 0.014 | 0.000 | 0.411 | 0.000 | 0.000 | 0.575 |
| IRIS_313-9281 | 0.000 | 0.000 | 0.000 | 0.991 | 0.000 | 0.009 | 0.000 |
| IRIS_313-9283 | 1.000 | 0.000 | 0.000 | 0.000 | 0.000 | 0.000 | 0.000 |
| IRIS_313-9285 | 0.000 | 0.000 | 0.000 | 0.810 | 0.000 | 0.186 | 0.004 |
| IRIS_313-9286 | 0.000 | 0.000 | 0.000 | 0.977 | 0.000 | 0.023 | 0.000 |
| IRIS_313-9287 | 0.000 | 0.000 | 0.000 | 0.999 | 0.000 | 0.001 | 0.000 |
| IRIS_313-9288 | 0.000 | 0.000 | 0.000 | 0.960 | 0.000 | 0.000 | 0.040 |
| IRIS_313-9294 | 0.023 | 0.000 | 0.000 | 0.977 | 0.000 | 0.000 | 0.000 |
| IRIS_313-9297 | 0.000 | 1.000 | 0.000 | 0.000 | 0.000 | 0.000 | 0.000 |
| IRIS_313-9301 | 0.000 | 0.924 | 0.000 | 0.000 | 0.000 | 0.076 | 0.000 |
| IRIS_313-9302 | 0.000 | 0.000 | 0.000 | 1.000 | 0.000 | 0.000 | 0.000 |
| IRIS_313-9310 | 0.000 | 0.000 | 0.000 | 0.465 | 0.000 | 0.000 | 0.535 |
| IRIS_313-9313 | 0.282 | 0.000 | 0.000 | 0.379 | 0.000 | 0.338 | 0.000 |
| IRIS_313-9314 | 0.000 | 0.000 | 0.000 | 0.337 | 0.000 | 0.000 | 0.663 |
| IRIS_313-9317 | 0.000 | 0.000 | 0.000 | 0.656 | 0.000 | 0.339 | 0.005 |
| IRIS_313-9320 | 0.000 | 0.000 | 0.000 | 0.646 | 0.000 | 0.000 | 0.354 |
| IRIS_313-9324 | 0.000 | 0.000 | 0.000 | 0.057 | 0.000 | 0.943 | 0.000 |
| IRIS_313-9325 | 0.000 | 0.000 | 0.000 | 0.241 | 0.000 | 0.000 | 0.759 |
| IRIS_313-9329 | 0.000 | 0.000 | 0.000 | 0.481 | 0.000 | 0.003 | 0.516 |
| IRIS_313-9342 | 0.000 | 0.000 | 0.000 | 1.000 | 0.000 | 0.000 | 0.000 |
| IRIS_313-9346 | 0.000 | 0.000 | 0.000 | 0.000 | 1.000 | 0.000 | 0.000 |
| IRIS_313-9347 | 0.019 | 0.000 | 0.000 | 0.777 | 0.000 | 0.204 | 0.000 |
| IRIS_313-9348 | 0.061 | 0.000 | 0.085 | 0.786 | 0.015 | 0.001 | 0.052 |

| | | | | | | | |
|---|---|---|---|---|---|---|---|
| IRIS_313-9351 | 0.140 | 0.010 | 0.003 | 0.672 | 0.000 | 0.176 | 0.000 |
| IRIS_313-9357 | 0.000 | 0.000 | 0.000 | 1.000 | 0.000 | 0.000 | 0.000 |
| IRIS_313-9360 | 0.000 | 0.000 | 0.002 | 0.998 | 0.000 | 0.000 | 0.000 |
| IRIS_313-9363 | 0.000 | 0.976 | 0.000 | 0.000 | 0.000 | 0.024 | 0.000 |
| IRIS_313-9366 | 0.000 | 0.561 | 0.000 | 0.000 | 0.439 | 0.000 | 0.000 |
| IRIS_313-9368 | 1.000 | 0.000 | 0.000 | 0.000 | 0.000 | 0.000 | 0.000 |
| IRIS_313-9372 | 0.000 | 0.000 | 0.000 | 0.031 | 0.000 | 0.969 | 0.000 |
| IRIS_313-9375 | 0.000 | 1.000 | 0.000 | 0.000 | 0.000 | 0.000 | 0.000 |
| IRIS_313-9379 | 0.000 | 0.000 | 0.000 | 0.000 | 1.000 | 0.000 | 0.000 |
| IRIS_313-9384 | 0.165 | 0.000 | 0.000 | 0.835 | 0.000 | 0.000 | 0.000 |
| IRIS_313-9388 | 0.409 | 0.030 | 0.000 | 0.562 | 0.000 | 0.000 | 0.000 |
| IRIS_313-9389 | 0.000 | 1.000 | 0.000 | 0.000 | 0.000 | 0.000 | 0.000 |
| IRIS_313-9391 | 0.000 | 0.000 | 0.000 | 0.979 | 0.000 | 0.021 | 0.000 |
| IRIS_313-9392 | 0.000 | 0.387 | 0.000 | 0.033 | 0.186 | 0.393 | 0.000 |
| IRIS_313-9397 | 0.082 | 0.000 | 0.000 | 0.918 | 0.000 | 0.000 | 0.000 |
| IRIS_313-9400 | 0.000 | 0.000 | 0.333 | 0.000 | 0.000 | 0.000 | 0.667 |
| IRIS_313-9402 | 0.000 | 0.007 | 0.000 | 0.991 | 0.002 | 0.000 | 0.000 |
| IRIS_313-9403 | 0.166 | 0.000 | 0.000 | 0.834 | 0.000 | 0.000 | 0.000 |
| IRIS_313-9404 | 0.000 | 0.289 | 0.030 | 0.000 | 0.196 | 0.000 | 0.484 |
| IRIS_313-9405 | 0.000 | 0.884 | 0.000 | 0.000 | 0.116 | 0.000 | 0.000 |
| IRIS_313-9406 | 0.002 | 0.003 | 0.000 | 0.995 | 0.000 | 0.000 | 0.000 |
| IRIS_313-9409 | 0.000 | 0.000 | 0.000 | 0.511 | 0.000 | 0.000 | 0.489 |
| IRIS_313-9410 | 0.000 | 0.000 | 0.000 | 0.000 | 1.000 | 0.000 | 0.000 |
| IRIS_313-9415 | 0.000 | 0.000 | 0.000 | 0.981 | 0.000 | 0.000 | 0.019 |
| IRIS_313-9422 | 1.000 | 0.000 | 0.000 | 0.000 | 0.000 | 0.000 | 0.000 |
| IRIS_313-9423 | 0.000 | 1.000 | 0.000 | 0.000 | 0.000 | 0.000 | 0.000 |
| IRIS_313-9424 | 0.000 | 0.000 | 0.000 | 1.000 | 0.000 | 0.000 | 0.000 |
| IRIS_313-9427 | 0.000 | 0.000 | 0.000 | 0.883 | 0.000 | 0.117 | 0.000 |
| IRIS_313-9429 | 0.000 | 0.000 | 0.033 | 0.544 | 0.000 | 0.424 | 0.000 |
| IRIS_313-9433 | 0.000 | 0.000 | 0.003 | 0.918 | 0.001 | 0.077 | 0.000 |
| IRIS_313-9438 | 0.000 | 0.000 | 0.000 | 0.000 | 0.919 | 0.081 | 0.000 |
| IRIS_313-9445 | 0.000 | 0.161 | 0.588 | 0.000 | 0.114 | 0.000 | 0.137 |
| IRIS_313-9449 | 1.000 | 0.000 | 0.000 | 0.000 | 0.000 | 0.000 | 0.000 |
| IRIS_313-9451 | 0.000 | 0.000 | 0.000 | 0.681 | 0.000 | 0.309 | 0.010 |
| IRIS_313-9452 | 0.000 | 0.898 | 0.000 | 0.000 | 0.102 | 0.000 | 0.000 |
| IRIS_313-9461 | 0.033 | 0.000 | 0.000 | 0.797 | 0.000 | 0.000 | 0.170 |
| IRIS_313-9463 | 0.000 | 0.000 | 0.000 | 0.000 | 1.000 | 0.000 | 0.000 |
| IRIS_313-9464 | 0.000 | 0.429 | 0.000 | 0.312 | 0.000 | 0.000 | 0.259 |
| IRIS_313-9468 | 0.000 | 0.000 | 0.000 | 0.000 | 1.000 | 0.000 | 0.000 |
| IRIS_313-9469 | 0.000 | 0.000 | 0.000 | 0.000 | 0.000 | 1.000 | 0.000 |
| IRIS_313-9470 | 0.000 | 1.000 | 0.000 | 0.000 | 0.000 | 0.000 | 0.000 |
| IRIS_313-9472 | 0.043 | 0.000 | 0.000 | 0.545 | 0.000 | 0.075 | 0.337 |
| IRIS_313-9482 | 0.000 | 0.000 | 0.000 | 0.000 | 0.000 | 0.159 | 0.841 |
| IRIS_313-9484 | 0.054 | 0.000 | 0.000 | 0.946 | 0.000 | 0.000 | 0.000 |
| IRIS_313-9491 | 0.000 | 1.000 | 0.000 | 0.000 | 0.000 | 0.000 | 0.000 |
| IRIS_313-9492 | 0.084 | 0.000 | 0.000 | 0.916 | 0.000 | 0.000 | 0.000 |
| IRIS_313-9503 | 0.000 | 0.000 | 0.000 | 0.000 | 0.000 | 0.058 | 0.942 |
| IRIS_313-9505 | 0.000 | 0.000 | 0.000 | 0.000 | 0.000 | 1.000 | 0.000 |
| IRIS_313-9506 | 0.045 | 0.173 | 0.124 | 0.000 | 0.101 | 0.557 | 0.000 |
| IRIS_313-9516 | 0.224 | 0.000 | 0.000 | 0.724 | 0.000 | 0.000 | 0.052 |
| IRIS_313-9519 | 0.000 | 1.000 | 0.000 | 0.000 | 0.000 | 0.000 | 0.000 |
| IRIS_313-9522 | 0.000 | 0.000 | 0.052 | 0.238 | 0.021 | 0.000 | 0.689 |
| IRIS_313-9523 | 0.000 | 0.259 | 0.000 | 0.000 | 0.741 | 0.000 | 0.000 |
| IRIS_313-9529 | 0.000 | 0.537 | 0.057 | 0.000 | 0.406 | 0.000 | 0.000 |
| IRIS_313-9533 | 0.000 | 0.000 | 0.000 | 0.980 | 0.000 | 0.020 | 0.000 |
| IRIS_313-9539 | 0.000 | 0.168 | 0.000 | 0.000 | 0.832 | 0.000 | 0.000 |
| IRIS_313-9547 | 0.040 | 0.000 | 0.000 | 0.960 | 0.000 | 0.000 | 0.000 |
| IRIS_313-9550 | 0.000 | 1.000 | 0.000 | 0.000 | 0.000 | 0.000 | 0.000 |
| IRIS_313-9551 | 0.025 | 0.000 | 0.000 | 0.975 | 0.000 | 0.000 | 0.000 |
| IRIS_313-9555 | 0.000 | 0.000 | 0.000 | 0.600 | 0.011 | 0.389 | 0.000 |
| IRIS_313-9557 | 0.120 | 0.000 | 0.010 | 0.713 | 0.000 | 0.157 | 0.000 |
| IRIS_313-9558 | 0.000 | 1.000 | 0.000 | 0.000 | 0.000 | 0.000 | 0.000 |
| IRIS_313-9560 | 0.000 | 0.000 | 0.000 | 0.795 | 0.000 | 0.205 | 0.000 |
| IRIS_313-9566 | 0.055 | 0.000 | 0.000 | 0.001 | 0.000 | 0.222 | 0.722 |
| IRIS_313-9567 | 0.000 | 0.000 | 0.000 | 1.000 | 0.000 | 0.000 | 0.000 |
| IRIS_313-9568 | 0.000 | 1.000 | 0.000 | 0.000 | 0.000 | 0.000 | 0.000 |
| IRIS_313-9570 | 0.000 | 0.058 | 0.000 | 0.000 | 0.000 | 0.452 | 0.490 |

| | | | | | | | |
|---|---|---|---|---|---|---|---|
| IRIS_313-9572 | 0.299 | 0.000 | 0.000 | 0.434 | 0.000 | 0.267 | 0.000 |
| IRIS_313-9574 | 0.000 | 0.000 | 0.000 | 0.351 | 0.000 | 0.000 | 0.649 |
| IRIS_313-9575 | 0.000 | 0.004 | 0.000 | 0.994 | 0.001 | 0.000 | 0.000 |
| IRIS_313-9582 | 0.000 | 0.000 | 0.000 | 0.428 | 0.000 | 0.000 | 0.572 |
| IRIS_313-9590 | 0.000 | 0.016 | 0.000 | 0.668 | 0.001 | 0.034 | 0.281 |
| IRIS_313-9593 | 0.000 | 0.000 | 0.000 | 0.802 | 0.000 | 0.198 | 0.000 |
| IRIS_313-9594 | 0.000 | 0.000 | 0.000 | 1.000 | 0.000 | 0.000 | 0.000 |
| IRIS_313-9600 | 0.000 | 0.372 | 0.000 | 0.314 | 0.000 | 0.000 | 0.314 |
| IRIS_313-9601 | 0.053 | 0.000 | 0.947 | 0.000 | 0.000 | 0.000 | 0.000 |
| IRIS_313-9602 | 0.000 | 0.000 | 0.000 | 0.311 | 0.000 | 0.000 | 0.689 |
| IRIS_313-9604 | 0.075 | 0.000 | 0.000 | 0.359 | 0.027 | 0.539 | 0.000 |
| IRIS_313-9605 | 0.000 | 0.000 | 0.000 | 0.000 | 0.000 | 1.000 | 0.000 |
| IRIS_313-9606 | 0.000 | 0.000 | 0.000 | 1.000 | 0.000 | 0.000 | 0.000 |
| IRIS_313-9609 | 0.030 | 0.005 | 0.000 | 0.473 | 0.000 | 0.492 | 0.000 |
| IRIS_313-9610 | 1.000 | 0.000 | 0.000 | 0.000 | 0.000 | 0.000 | 0.000 |
| IRIS_313-9611 | 0.054 | 0.000 | 0.000 | 0.946 | 0.000 | 0.000 | 0.000 |
| IRIS_313-9616 | 0.012 | 0.988 | 0.000 | 0.000 | 0.000 | 0.000 | 0.000 |
| IRIS_313-9617 | 0.036 | 0.000 | 0.000 | 0.964 | 0.000 | 0.000 | 0.000 |
| IRIS_313-9619 | 0.000 | 0.000 | 0.002 | 0.998 | 0.000 | 0.000 | 0.000 |
| IRIS_313-9623 | 0.000 | 0.000 | 0.000 | 0.123 | 0.000 | 0.202 | 0.675 |
| IRIS_313-9626 | 1.000 | 0.000 | 0.000 | 0.000 | 0.000 | 0.000 | 0.000 |
| IRIS_313-9629 | 0.000 | 0.000 | 1.000 | 0.000 | 0.000 | 0.000 | 0.000 |
| IRIS_313-9634 | 0.040 | 0.019 | 0.000 | 0.752 | 0.003 | 0.186 | 0.000 |
| IRIS_313-9636 | 0.880 | 0.000 | 0.080 | 0.000 | 0.039 | 0.000 | 0.000 |
| IRIS_313-9637 | 0.040 | 0.036 | 0.000 | 0.817 | 0.020 | 0.087 | 0.000 |
| IRIS_313-9641 | 1.000 | 0.000 | 0.000 | 0.000 | 0.000 | 0.000 | 0.000 |
| IRIS_313-9648 | 0.170 | 0.394 | 0.000 | 0.416 | 0.000 | 0.019 | 0.000 |
| IRIS_313-9661 | 0.993 | 0.000 | 0.000 | 0.000 | 0.000 | 0.000 | 0.007 |
| IRIS_313-9669 | 0.000 | 0.000 | 0.000 | 0.962 | 0.000 | 0.038 | 0.000 |
| IRIS_313-9682 | 0.000 | 0.000 | 0.868 | 0.000 | 0.100 | 0.000 | 0.031 |
| IRIS_313-9685 | 0.000 | 0.000 | 0.000 | 0.673 | 0.046 | 0.000 | 0.281 |
| IRIS_313-9687 | 0.000 | 0.000 | 0.000 | 0.995 | 0.000 | 0.005 | 0.000 |
| IRIS_313-9691 | 0.000 | 1.000 | 0.000 | 0.000 | 0.000 | 0.000 | 0.000 |
| IRIS_313-9694 | 0.000 | 0.000 | 0.000 | 0.437 | 0.000 | 0.000 | 0.563 |
| IRIS_313-9695 | 0.605 | 0.000 | 0.000 | 0.000 | 0.000 | 0.264 | 0.130 |
| IRIS_313-9696 | 0.000 | 0.000 | 0.000 | 0.380 | 0.000 | 0.012 | 0.608 |
| IRIS_313-9697 | 0.000 | 0.000 | 0.000 | 0.710 | 0.000 | 0.000 | 0.290 |
| IRIS_313-9698 | 0.000 | 0.000 | 0.000 | 0.000 | 1.000 | 0.000 | 0.000 |
| IRIS_313-9699 | 0.000 | 0.000 | 0.000 | 0.446 | 0.000 | 0.000 | 0.554 |
| IRIS_313-9701 | 0.000 | 0.000 | 0.000 | 0.000 | 1.000 | 0.000 | 0.000 |
| IRIS_313-9702 | 0.000 | 0.000 | 0.000 | 0.000 | 0.988 | 0.012 | 0.000 |
| IRIS_313-9703 | 0.000 | 0.000 | 0.000 | 0.000 | 1.000 | 0.000 | 0.000 |
| IRIS_313-9705 | 0.000 | 0.000 | 0.000 | 0.025 | 0.000 | 0.975 | 0.000 |
| IRIS_313-9706 | 0.000 | 0.000 | 0.000 | 0.116 | 0.000 | 0.884 | 0.000 |
| IRIS_313-9708 | 0.000 | 0.000 | 0.000 | 0.009 | 0.000 | 0.991 | 0.000 |
| IRIS_313-9709 | 0.029 | 0.000 | 0.000 | 0.971 | 0.000 | 0.000 | 0.000 |
| IRIS_313-9723 | 0.000 | 0.000 | 0.000 | 0.000 | 0.000 | 1.000 | 0.000 |
| IRIS_313-9724 | 0.000 | 0.000 | 0.000 | 0.000 | 1.000 | 0.000 | 0.000 |
| IRIS_313-9725 | 0.000 | 0.000 | 0.000 | 0.002 | 0.981 | 0.017 | 0.000 |
| IRIS_313-9727 | 0.359 | 0.000 | 0.000 | 0.000 | 0.000 | 0.641 | 0.000 |
| IRIS_313-9730 | 0.000 | 0.000 | 0.000 | 0.000 | 0.000 | 1.000 | 0.000 |
| IRIS_313-9732 | 0.259 | 0.000 | 0.000 | 0.741 | 0.000 | 0.000 | 0.000 |
| IRIS_313-9740 | 0.293 | 0.147 | 0.000 | 0.560 | 0.000 | 0.000 | 0.000 |
| IRIS_313-9742 | 0.000 | 1.000 | 0.000 | 0.000 | 0.000 | 0.000 | 0.000 |
| IRIS_313-9745 | 0.000 | 1.000 | 0.000 | 0.000 | 0.000 | 0.000 | 0.000 |
| IRIS_313-9758 | 0.000 | 0.000 | 0.000 | 0.000 | 0.000 | 1.000 | 0.000 |
| IRIS_313-9759 | 0.000 | 0.048 | 0.000 | 0.000 | 0.952 | 0.000 | 0.000 |
| IRIS_313-9767 | 0.000 | 0.232 | 0.000 | 0.000 | 0.000 | 0.000 | 0.768 |
| IRIS_313-9769 | 0.000 | 0.130 | 0.000 | 0.000 | 0.870 | 0.000 | 0.000 |
| IRIS_313-9770 | 0.000 | 0.095 | 0.000 | 0.000 | 0.905 | 0.000 | 0.000 |
| IRIS_313-9771 | 0.000 | 0.000 | 0.000 | 0.000 | 1.000 | 0.000 | 0.000 |
| IRIS_313-9774 | 0.000 | 0.361 | 0.000 | 0.000 | 0.639 | 0.000 | 0.000 |
| IRIS_313-9778 | 0.544 | 0.000 | 0.000 | 0.436 | 0.004 | 0.016 | 0.000 |
| IRIS_313-9782 | 0.000 | 0.000 | 0.000 | 0.000 | 1.000 | 0.000 | 0.000 |
| IRIS_313-9783 | 0.541 | 0.057 | 0.401 | 0.000 | 0.000 | 0.000 | 0.000 |
| IRIS_313-9789 | 0.000 | 0.740 | 0.000 | 0.000 | 0.189 | 0.000 | 0.071 |
| IRIS_313-9790 | 0.000 | 0.108 | 0.000 | 0.000 | 0.892 | 0.000 | 0.000 |

| | | | | | | | |
|---|---|---|---|---|---|---|---|
| IRIS_313-9791 | 0.037 | 0.000 | 0.000 | 0.406 | 0.000 | 0.557 | 0.000 |
| IRIS_313-9795 | 0.053 | 0.000 | 0.000 | 0.771 | 0.120 | 0.000 | 0.056 |
| IRIS_313-9800 | 0.000 | 0.758 | 0.000 | 0.000 | 0.184 | 0.058 | 0.000 |
| IRIS_313-9809 | 0.000 | 0.000 | 0.000 | 0.990 | 0.000 | 0.000 | 0.010 |
| IRIS_313-9811 | 0.000 | 0.000 | 0.000 | 0.000 | 1.000 | 0.000 | 0.000 |
| IRIS_313-9813 | 0.000 | 0.000 | 0.000 | 0.000 | 1.000 | 0.000 | 0.000 |
| IRIS_313-9814 | 0.000 | 0.092 | 0.000 | 0.000 | 0.908 | 0.000 | 0.000 |
| IRIS_313-9817 | 0.125 | 0.309 | 0.000 | 0.566 | 0.000 | 0.000 | 0.000 |
| IRIS_313-9818 | 0.000 | 0.355 | 0.000 | 0.000 | 0.000 | 0.000 | 0.645 |
| IRIS_313-9822 | 0.000 | 0.000 | 0.000 | 0.143 | 0.000 | 0.857 | 0.000 |
| IRIS_313-9825 | 0.024 | 0.000 | 0.000 | 0.976 | 0.000 | 0.000 | 0.000 |
| IRIS_313-9831 | 0.093 | 0.000 | 0.000 | 0.479 | 0.000 | 0.428 | 0.000 |
| IRIS_313-9832 | 0.116 | 0.000 | 0.010 | 0.744 | 0.000 | 0.129 | 0.000 |
| IRIS_313-9838 | 0.000 | 0.000 | 0.000 | 0.000 | 1.000 | 0.000 | 0.000 |
| IRIS_313-9839 | 0.000 | 0.000 | 0.000 | 0.000 | 1.000 | 0.000 | 0.000 |
| IRIS_313-9841 | 0.051 | 0.219 | 0.000 | 0.477 | 0.032 | 0.061 | 0.160 |
| IRIS_313-9851 | 0.000 | 1.000 | 0.000 | 0.000 | 0.000 | 0.000 | 0.000 |
| IRIS_313-9861 | 0.951 | 0.000 | 0.000 | 0.043 | 0.006 | 0.000 | 0.000 |
| IRIS_313-9862 | 0.159 | 0.000 | 0.000 | 0.806 | 0.000 | 0.000 | 0.035 |
| IRIS_313-9867 | 0.000 | 0.000 | 0.000 | 0.076 | 0.000 | 0.000 | 0.924 |
| IRIS_313-9880 | 0.000 | 0.000 | 0.000 | 0.000 | 1.000 | 0.000 | 0.000 |
| IRIS_313-9882 | 0.000 | 0.000 | 0.012 | 0.635 | 0.000 | 0.349 | 0.004 |
| IRIS_313-9884 | 0.000 | 0.000 | 0.000 | 0.000 | 1.000 | 0.000 | 0.000 |
| IRIS_313-9886 | 0.000 | 0.000 | 0.000 | 0.000 | 1.000 | 0.000 | 0.000 |
| IRIS_313-9887 | 0.000 | 0.000 | 0.000 | 0.000 | 1.000 | 0.000 | 0.000 |
| IRIS_313-9890 | 0.000 | 0.069 | 0.000 | 0.000 | 0.931 | 0.000 | 0.000 |
| IRIS_313-9891 | 0.000 | 0.000 | 0.000 | 0.000 | 1.000 | 0.000 | 0.000 |
| IRIS_313-9897 | 0.000 | 1.000 | 0.000 | 0.000 | 0.000 | 0.000 | 0.000 |
| IRIS_313-9898 | 0.000 | 0.000 | 0.000 | 0.995 | 0.000 | 0.000 | 0.005 |
| IRIS_313-9917 | 0.000 | 0.000 | 0.000 | 0.178 | 0.000 | 0.078 | 0.745 |
| IRIS_313-9922 | 0.000 | 0.000 | 0.000 | 0.000 | 0.029 | 0.090 | 0.880 |
| IRIS_313-9924 | 0.000 | 0.000 | 0.000 | 0.000 | 0.000 | 0.305 | 0.695 |
| IRIS_313-9925 | 0.020 | 0.000 | 0.000 | 0.050 | 0.028 | 0.000 | 0.902 |
| IRIS_313-9926 | 0.057 | 0.000 | 0.013 | 0.930 | 0.000 | 0.000 | 0.000 |
| IRIS_313-9928 | 0.000 | 0.965 | 0.000 | 0.001 | 0.000 | 0.034 | 0.000 |
| IRIS_313-9929 | 0.000 | 1.000 | 0.000 | 0.000 | 0.000 | 0.000 | 0.000 |
| IRIS_313-9935 | 0.068 | 0.000 | 0.014 | 0.918 | 0.000 | 0.000 | 0.000 |
| IRIS_313-9936 | 0.000 | 0.000 | 0.000 | 0.976 | 0.000 | 0.024 | 0.000 |
| IRIS_313-9937 | 0.000 | 0.262 | 0.000 | 0.000 | 0.738 | 0.000 | 0.000 |
| IRIS_313-9939 | 0.000 | 0.445 | 0.000 | 0.303 | 0.000 | 0.000 | 0.252 |
| IRIS_313-9940 | 0.000 | 0.000 | 0.000 | 0.061 | 0.000 | 0.000 | 0.939 |
| IRIS_313-9944 | 0.148 | 0.005 | 0.000 | 0.848 | 0.000 | 0.000 | 0.000 |
| IRIS_313-9949 | 0.065 | 0.773 | 0.136 | 0.000 | 0.026 | 0.000 | 0.000 |
| IRIS_313-9953 | 0.000 | 0.000 | 0.000 | 0.492 | 0.000 | 0.000 | 0.508 |
| IRIS_313-9961 | 0.000 | 0.000 | 0.000 | 0.000 | 1.000 | 0.000 | 0.000 |
| IRIS_313-9963 | 1.000 | 0.000 | 0.000 | 0.000 | 0.000 | 0.000 | 0.000 |
| IRIS_313-9964 | 0.000 | 0.000 | 0.000 | 0.000 | 1.000 | 0.000 | 0.000 |
| IRIS_313-9966 | 0.000 | 0.000 | 0.000 | 0.000 | 0.000 | 0.000 | 1.000 |
| IRIS_313-9968 | 0.000 | 0.000 | 0.000 | 0.401 | 0.000 | 0.599 | 0.000 |
| IRIS_313-9969 | 0.038 | 0.000 | 0.000 | 0.875 | 0.000 | 0.086 | 0.000 |
| IRIS_313-9970 | 0.107 | 0.000 | 0.000 | 0.703 | 0.000 | 0.190 | 0.000 |
| IRIS_313-9974 | 0.000 | 0.000 | 0.000 | 0.000 | 1.000 | 0.000 | 0.000 |
| IRIS_313-9976 | 0.000 | 0.000 | 0.000 | 0.000 | 0.000 | 1.000 | 0.000 |
| IRIS_313-9978 | 0.000 | 0.000 | 1.000 | 0.000 | 0.000 | 0.000 | 0.000 |
| IRIS_313-9980 | 0.000 | 0.556 | 0.000 | 0.000 | 0.444 | 0.000 | 0.000 |
| IRIS_313-9986 | 0.005 | 0.000 | 0.000 | 0.995 | 0.000 | 0.000 | 0.000 |
| IRIS_313-9989 | 0.000 | 0.000 | 0.000 | 1.000 | 0.000 | 0.000 | 0.000 |
| IRIS_313-9995 | 0.000 | 0.730 | 0.000 | 0.000 | 0.197 | 0.000 | 0.073 |
| IRIS_313-9996 | 0.000 | 0.000 | 0.000 | 0.000 | 1.000 | 0.000 | 0.000 |
| IRIS_313-10000 | 0.000 | 0.000 | 0.000 | 0.000 | 0.042 | 0.113 | 0.846 |
| IRIS_313-10001 | 0.000 | 0.000 | 0.000 | 0.139 | 0.000 | 0.095 | 0.766 |
| IRIS_313-10002 | 0.000 | 0.000 | 0.000 | 0.000 | 0.000 | 0.244 | 0.756 |
| IRIS_313-10007 | 0.000 | 0.000 | 0.000 | 0.907 | 0.000 | 0.093 | 0.000 |
| IRIS_313-10010 | 0.066 | 0.000 | 0.000 | 0.934 | 0.000 | 0.000 | 0.000 |
| IRIS_313-10014 | 0.000 | 0.215 | 0.000 | 0.000 | 0.785 | 0.000 | 0.000 |
| IRIS_313-10016 | 0.000 | 0.000 | 0.000 | 0.067 | 0.000 | 0.376 | 0.557 |
| IRIS_313-10020 | 0.870 | 0.000 | 0.083 | 0.000 | 0.047 | 0.000 | 0.000 |

| | | | | | | | |
|---|---|---|---|---|---|---|---|
| IRIS_313-10025 | 0.042 | 0.340 | 0.266 | 0.000 | 0.353 | 0.000 | 0.000 |
| IRIS_313-10026 | 0.224 | 0.000 | 0.000 | 0.574 | 0.060 | 0.142 | 0.000 |
| IRIS_313-10030 | 0.277 | 0.051 | 0.000 | 0.672 | 0.000 | 0.000 | 0.000 |
| IRIS_313-10032 | 0.000 | 0.000 | 1.000 | 0.000 | 0.000 | 0.000 | 0.000 |
| IRIS_313-10034 | 0.000 | 0.000 | 0.000 | 0.559 | 0.000 | 0.435 | 0.006 |
| IRIS_313-10035 | 0.000 | 0.080 | 0.049 | 0.769 | 0.102 | 0.000 | 0.000 |
| IRIS_313-10040 | 0.000 | 0.000 | 0.000 | 0.000 | 0.030 | 0.000 | 0.970 |
| IRIS_313-10041 | 0.000 | 1.000 | 0.000 | 0.000 | 0.000 | 0.000 | 0.000 |
| IRIS_313-10045 | 0.022 | 0.000 | 0.000 | 0.977 | 0.000 | 0.001 | 0.000 |
| IRIS_313-10046 | 0.000 | 0.000 | 0.000 | 0.166 | 0.000 | 0.000 | 0.834 |
| IRIS_313-10047 | 0.000 | 0.000 | 0.000 | 0.000 | 0.000 | 1.000 | 0.000 |
| IRIS_313-10048 | 0.224 | 0.019 | 0.000 | 0.708 | 0.000 | 0.049 | 0.000 |
| IRIS_313-10050 | 0.108 | 0.000 | 0.000 | 0.719 | 0.000 | 0.000 | 0.172 |
| IRIS_313-10051 | 0.000 | 1.000 | 0.000 | 0.000 | 0.000 | 0.000 | 0.000 |
| IRIS_313-10054 | 0.000 | 0.053 | 0.000 | 0.221 | 0.042 | 0.146 | 0.538 |
| IRIS_313-10056 | 0.000 | 0.680 | 0.000 | 0.000 | 0.320 | 0.000 | 0.000 |
| IRIS_313-10057 | 0.000 | 0.000 | 0.000 | 0.000 | 1.000 | 0.000 | 0.000 |
| IRIS_313-10059 | 0.000 | 0.000 | 0.000 | 0.000 | 1.000 | 0.000 | 0.000 |
| IRIS_313-10061 | 0.052 | 0.500 | 0.000 | 0.000 | 0.362 | 0.000 | 0.086 |
| IRIS_313-10062 | 0.000 | 0.634 | 0.000 | 0.000 | 0.366 | 0.000 | 0.000 |
| IRIS_313-10065 | 0.000 | 0.699 | 0.000 | 0.000 | 0.239 | 0.000 | 0.062 |
| IRIS_313-10067 | 0.000 | 0.698 | 0.000 | 0.000 | 0.230 | 0.073 | 0.000 |
| IRIS_313-10071 | 0.000 | 0.751 | 0.000 | 0.000 | 0.192 | 0.057 | 0.000 |
| IRIS_313-10073 | 0.000 | 0.774 | 0.000 | 0.000 | 0.151 | 0.000 | 0.075 |
| IRIS_313-10074 | 0.000 | 0.716 | 0.000 | 0.000 | 0.209 | 0.075 | 0.000 |
| IRIS_313-10075 | 0.000 | 0.714 | 0.000 | 0.000 | 0.227 | 0.060 | 0.000 |
| IRIS_313-10076 | 0.000 | 0.705 | 0.000 | 0.000 | 0.223 | 0.000 | 0.072 |
| IRIS_313-10077 | 0.017 | 0.000 | 0.000 | 0.955 | 0.000 | 0.028 | 0.000 |
| IRIS_313-10078 | 0.000 | 0.701 | 0.000 | 0.000 | 0.225 | 0.074 | 0.000 |
| IRIS_313-10079 | 0.000 | 0.697 | 0.000 | 0.000 | 0.231 | 0.072 | 0.000 |
| IRIS_313-10080 | 0.000 | 0.775 | 0.000 | 0.000 | 0.151 | 0.073 | 0.000 |
| IRIS_313-10082 | 0.000 | 0.729 | 0.000 | 0.000 | 0.227 | 0.000 | 0.044 |
| IRIS_313-10083 | 0.000 | 0.261 | 0.000 | 0.000 | 0.739 | 0.000 | 0.000 |
| IRIS_313-10084 | 0.000 | 0.000 | 0.000 | 0.000 | 1.000 | 0.000 | 0.000 |
| IRIS_313-10089 | 0.000 | 0.115 | 0.000 | 0.000 | 0.885 | 0.000 | 0.000 |
| IRIS_313-10092 | 0.000 | 0.000 | 0.000 | 0.000 | 1.000 | 0.000 | 0.000 |
| IRIS_313-10093 | 0.000 | 0.000 | 0.000 | 0.000 | 1.000 | 0.000 | 0.000 |
| IRIS_313-10094 | 0.000 | 0.710 | 0.000 | 0.000 | 0.215 | 0.075 | 0.000 |
| IRIS_313-10096 | 0.000 | 0.297 | 0.000 | 0.000 | 0.627 | 0.075 | 0.000 |
| IRIS_313-10097 | 0.000 | 0.000 | 0.000 | 0.000 | 1.000 | 0.000 | 0.000 |
| IRIS_313-10099 | 0.007 | 0.953 | 0.000 | 0.000 | 0.040 | 0.000 | 0.000 |
| IRIS_313-10102 | 0.266 | 0.006 | 0.000 | 0.694 | 0.000 | 0.034 | 0.000 |
| IRIS_313-10103 | 0.000 | 0.000 | 0.000 | 0.224 | 0.086 | 0.690 | 0.000 |
| IRIS_313-10109 | 0.000 | 0.000 | 0.000 | 0.448 | 0.007 | 0.000 | 0.546 |
| IRIS_313-10111 | 0.000 | 0.414 | 0.000 | 0.000 | 0.586 | 0.000 | 0.000 |
| IRIS_313-10113 | 0.000 | 0.000 | 0.000 | 0.000 | 0.000 | 0.000 | 1.000 |
| IRIS_313-10114 | 0.324 | 0.086 | 0.000 | 0.590 | 0.000 | 0.000 | 0.000 |
| IRIS_313-10119 | 0.000 | 0.320 | 0.000 | 0.000 | 0.680 | 0.000 | 0.000 |
| IRIS_313-10124 | 0.000 | 0.000 | 0.000 | 0.000 | 1.000 | 0.000 | 0.000 |
| IRIS_313-10129 | 0.000 | 0.000 | 0.000 | 0.009 | 0.000 | 0.991 | 0.000 |
| IRIS_313-10134 | 0.000 | 0.000 | 0.000 | 1.000 | 0.000 | 0.000 | 0.000 |
| IRIS_313-10147 | 0.000 | 0.000 | 0.000 | 1.000 | 0.000 | 0.000 | 0.000 |
| IRIS_313-10148 | 0.038 | 0.000 | 0.000 | 0.299 | 0.000 | 0.663 | 0.000 |
| IRIS_313-10150 | 1.000 | 0.000 | 0.000 | 0.000 | 0.000 | 0.000 | 0.000 |
| IRIS_313-10151 | 0.000 | 0.000 | 0.000 | 1.000 | 0.000 | 0.000 | 0.000 |
| IRIS_313-10152 | 0.000 | 1.000 | 0.000 | 0.000 | 0.000 | 0.000 | 0.000 |
| IRIS_313-10154 | 0.000 | 0.000 | 0.000 | 0.000 | 0.000 | 1.000 | 0.000 |
| IRIS_313-10158 | 0.016 | 0.156 | 0.000 | 0.446 | 0.000 | 0.000 | 0.382 |
| IRIS_313-10161 | 0.000 | 0.000 | 0.000 | 0.000 | 0.000 | 0.079 | 0.921 |
| IRIS_313-10164 | 0.000 | 0.000 | 0.000 | 0.000 | 1.000 | 0.000 | 0.000 |
| IRIS_313-10167 | 0.000 | 0.000 | 0.000 | 0.288 | 0.000 | 0.044 | 0.668 |
| IRIS_313-10168 | 0.000 | 0.000 | 0.000 | 0.393 | 0.000 | 0.607 | 0.000 |
| IRIS_313-10170 | 0.000 | 0.000 | 0.000 | 0.000 | 0.000 | 1.000 | 0.000 |
| IRIS_313-10171 | 0.000 | 0.000 | 0.018 | 0.081 | 0.000 | 0.901 | 0.000 |
| IRIS_313-10176 | 0.018 | 0.549 | 0.000 | 0.000 | 0.433 | 0.000 | 0.000 |
| IRIS_313-10177 | 0.012 | 0.014 | 0.000 | 0.621 | 0.000 | 0.352 | 0.000 |
| IRIS_313-10178 | 0.004 | 0.000 | 0.000 | 0.131 | 0.000 | 0.865 | 0.000 |

| | | | | | | | |
|---|---|---|---|---|---|---|---|
| IRIS_313-10179 | 0.000 | 0.000 | 0.000 | 0.560 | 0.000 | 0.435 | 0.004 |
| IRIS_313-10189 | 0.000 | 0.004 | 0.000 | 0.125 | 0.000 | 0.871 | 0.000 |
| IRIS_313-10190 | 0.020 | 0.000 | 0.000 | 0.102 | 0.000 | 0.878 | 0.000 |
| IRIS_313-10191 | 0.000 | 0.000 | 0.000 | 0.290 | 0.000 | 0.710 | 0.000 |
| IRIS_313-10196 | 0.025 | 0.845 | 0.000 | 0.000 | 0.105 | 0.026 | 0.000 |
| IRIS_313-10211 | 0.000 | 0.000 | 0.000 | 0.000 | 0.000 | 1.000 | 0.000 |
| IRIS_313-10220 | 0.000 | 0.000 | 0.000 | 0.012 | 0.000 | 0.240 | 0.748 |
| IRIS_313-10221 | 0.000 | 0.005 | 0.000 | 0.550 | 0.006 | 0.439 | 0.000 |
| IRIS_313-10224 | 0.000 | 0.000 | 0.000 | 0.131 | 0.000 | 0.869 | 0.000 |
| IRIS_313-10226 | 0.000 | 0.000 | 0.041 | 0.483 | 0.006 | 0.470 | 0.000 |
| IRIS_313-10228 | 0.000 | 0.000 | 0.072 | 0.000 | 0.909 | 0.013 | 0.007 |
| IRIS_313-10234 | 0.000 | 0.000 | 0.000 | 0.213 | 0.000 | 0.071 | 0.715 |
| IRIS_313-10235 | 0.000 | 0.000 | 0.000 | 0.471 | 0.000 | 0.050 | 0.479 |
| IRIS_313-10237 | 0.000 | 0.000 | 0.000 | 0.000 | 0.000 | 0.243 | 0.757 |
| IRIS_313-10238 | 0.000 | 0.000 | 0.000 | 0.000 | 0.000 | 0.241 | 0.759 |
| IRIS_313-10239 | 0.000 | 0.000 | 0.032 | 0.468 | 0.000 | 0.500 | 0.000 |
| IRIS_313-10242 | 0.000 | 0.000 | 0.000 | 0.000 | 1.000 | 0.000 | 0.000 |
| IRIS_313-10247 | 0.000 | 0.054 | 0.000 | 0.000 | 0.000 | 0.002 | 0.944 |
| IRIS_313-10257 | 0.000 | 0.026 | 0.000 | 0.273 | 0.037 | 0.000 | 0.664 |
| IRIS_313-10258 | 0.000 | 0.348 | 0.000 | 0.000 | 0.652 | 0.000 | 0.000 |
| IRIS_313-10260 | 0.000 | 0.000 | 0.000 | 0.010 | 0.000 | 0.244 | 0.746 |
| IRIS_313-10263 | 0.000 | 0.145 | 0.000 | 0.426 | 0.000 | 0.000 | 0.429 |
| IRIS_313-10268 | 0.000 | 0.151 | 0.000 | 0.000 | 0.000 | 0.081 | 0.768 |
| IRIS_313-10271 | 0.000 | 0.000 | 0.000 | 0.268 | 0.000 | 0.167 | 0.565 |
| IRIS_313-10272 | 0.000 | 0.607 | 0.000 | 0.255 | 0.138 | 0.000 | 0.000 |
| IRIS_313-10274 | 0.000 | 0.000 | 0.000 | 0.000 | 0.000 | 0.000 | 1.000 |
| IRIS_313-10275 | 0.000 | 0.007 | 0.000 | 0.309 | 0.031 | 0.180 | 0.473 |
| IRIS_313-10279 | 0.000 | 0.000 | 0.000 | 0.225 | 0.000 | 0.000 | 0.775 |
| IRIS_313-10285 | 0.000 | 0.069 | 0.000 | 0.249 | 0.000 | 0.413 | 0.269 |
| IRIS_313-10287 | 0.000 | 0.031 | 0.000 | 0.000 | 0.054 | 0.093 | 0.822 |
| IRIS_313-10289 | 0.000 | 0.000 | 0.007 | 0.000 | 0.005 | 0.155 | 0.833 |
| IRIS_313-10290 | 0.000 | 0.000 | 0.000 | 0.000 | 0.000 | 0.184 | 0.816 |
| IRIS_313-10293 | 0.000 | 0.050 | 0.000 | 0.000 | 0.000 | 0.000 | 0.950 |
| IRIS_313-10294 | 0.000 | 0.000 | 0.000 | 0.274 | 0.000 | 0.122 | 0.604 |
| IRIS_313-10295 | 0.063 | 0.279 | 0.231 | 0.000 | 0.427 | 0.000 | 0.000 |
| IRIS_313-10298 | 0.000 | 0.000 | 0.000 | 0.000 | 0.000 | 0.037 | 0.963 |
| IRIS_313-10300 | 0.000 | 0.068 | 0.000 | 0.000 | 0.066 | 0.045 | 0.822 |
| IRIS_313-10301 | 0.000 | 0.114 | 0.006 | 0.162 | 0.000 | 0.184 | 0.534 |
| IRIS_313-10307 | 0.000 | 0.000 | 0.000 | 0.000 | 0.000 | 0.270 | 0.730 |
| IRIS_313-10314 | 0.000 | 0.003 | 0.000 | 0.000 | 0.018 | 0.158 | 0.821 |
| IRIS_313-10318 | 0.000 | 0.128 | 0.000 | 0.430 | 0.023 | 0.000 | 0.419 |
| IRIS_313-10325 | 0.000 | 0.096 | 0.000 | 0.173 | 0.000 | 0.174 | 0.556 |
| IRIS_313-10327 | 0.000 | 1.000 | 0.000 | 0.000 | 0.000 | 0.000 | 0.000 |
| IRIS_313-10332 | 0.000 | 0.000 | 0.000 | 0.000 | 0.000 | 0.000 | 1.000 |
| IRIS_313-10333 | 0.023 | 0.000 | 0.029 | 0.113 | 0.000 | 0.000 | 0.835 |
| IRIS_313-10334 | 0.022 | 0.000 | 0.032 | 0.106 | 0.000 | 0.000 | 0.840 |
| IRIS_313-10336 | 0.000 | 0.971 | 0.000 | 0.000 | 0.000 | 0.029 | 0.000 |
| IRIS_313-10337 | 0.000 | 0.000 | 0.000 | 0.000 | 0.000 | 0.073 | 0.927 |
| IRIS_313-10340 | 0.000 | 0.000 | 0.000 | 0.663 | 0.000 | 0.020 | 0.317 |
| IRIS_313-10341 | 0.014 | 0.000 | 0.000 | 0.442 | 0.000 | 0.000 | 0.544 |
| IRIS_313-10348 | 0.095 | 0.000 | 0.002 | 0.510 | 0.000 | 0.070 | 0.323 |
| IRIS_313-10349 | 0.000 | 0.000 | 0.006 | 0.446 | 0.028 | 0.000 | 0.521 |
| IRIS_313-10352 | 0.049 | 0.070 | 0.000 | 0.105 | 0.000 | 0.000 | 0.776 |
| IRIS_313-10353 | 0.068 | 0.000 | 0.000 | 0.000 | 0.029 | 0.353 | 0.551 |
| IRIS_313-10355 | 0.063 | 0.000 | 0.000 | 0.192 | 0.000 | 0.080 | 0.665 |
| IRIS_313-10357 | 0.030 | 0.000 | 0.000 | 0.470 | 0.000 | 0.158 | 0.342 |
| IRIS_313-10359 | 0.306 | 0.000 | 0.000 | 0.063 | 0.000 | 0.185 | 0.446 |
| IRIS_313-10360 | 0.000 | 0.000 | 0.000 | 0.502 | 0.000 | 0.064 | 0.434 |
| IRIS_313-10361 | 0.068 | 0.000 | 0.000 | 0.285 | 0.000 | 0.344 | 0.303 |
| IRIS_313-10366 | 0.148 | 0.000 | 0.000 | 0.140 | 0.000 | 0.077 | 0.635 |
| IRIS_313-10371 | 0.235 | 0.000 | 0.000 | 0.060 | 0.000 | 0.160 | 0.545 |
| IRIS_313-10373 | 0.000 | 0.000 | 0.000 | 0.000 | 1.000 | 0.000 | 0.000 |
| IRIS_313-10374 | 0.000 | 0.000 | 0.000 | 0.398 | 0.000 | 0.000 | 0.602 |
| IRIS_313-10375 | 0.254 | 0.000 | 0.000 | 0.000 | 0.000 | 0.078 | 0.669 |
| IRIS_313-10379 | 0.000 | 0.000 | 0.000 | 0.000 | 1.000 | 0.000 | 0.000 |
| IRIS_313-10380 | 0.906 | 0.000 | 0.000 | 0.000 | 0.016 | 0.000 | 0.078 |
| IRIS_313-10385 | 0.015 | 0.000 | 0.000 | 0.000 | 0.000 | 0.107 | 0.878 |

| | | | | | | | |
|---|---|---|---|---|---|---|---|
| IRIS_313-10392 | 0.016 | 0.000 | 0.000 | 0.000 | 0.019 | 0.135 | 0.830 |
| IRIS_313-10394 | 0.026 | 0.000 | 0.000 | 0.000 | 0.015 | 0.000 | 0.958 |
| IRIS_313-10396 | 0.000 | 0.000 | 0.000 | 0.403 | 0.009 | 0.000 | 0.588 |
| IRIS_313-10397 | 0.000 | 0.000 | 0.000 | 0.000 | 0.000 | 0.235 | 0.765 |
| IRIS_313-10398 | 0.000 | 0.000 | 0.000 | 0.000 | 0.000 | 0.226 | 0.774 |
| IRIS_313-10399 | 0.000 | 0.000 | 0.000 | 0.152 | 0.000 | 0.000 | 0.848 |
| IRIS_313-10400 | 0.000 | 0.000 | 0.000 | 0.000 | 0.017 | 0.066 | 0.917 |
| IRIS_313-10401 | 0.000 | 0.000 | 0.000 | 0.000 | 0.000 | 0.223 | 0.777 |
| IRIS_313-10402 | 0.000 | 0.000 | 0.000 | 0.195 | 0.000 | 0.000 | 0.805 |
| IRIS_313-10403 | 0.000 | 0.000 | 0.000 | 0.171 | 0.000 | 0.000 | 0.829 |
| IRIS_313-10404 | 0.010 | 0.000 | 0.000 | 0.000 | 0.000 | 0.797 | 0.192 |
| IRIS_313-10412 | 0.000 | 0.000 | 0.000 | 0.266 | 0.000 | 0.000 | 0.734 |
| IRIS_313-10417 | 0.000 | 0.000 | 0.000 | 0.000 | 0.000 | 0.322 | 0.678 |
| IRIS_313-10421 | 0.000 | 0.000 | 0.000 | 0.394 | 0.000 | 0.028 | 0.578 |
| IRIS_313-10422 | 0.000 | 0.000 | 0.000 | 0.868 | 0.000 | 0.000 | 0.132 |
| IRIS_313-10423 | 0.000 | 0.000 | 0.000 | 0.000 | 0.000 | 0.000 | 1.000 |
| IRIS_313-10428 | 0.000 | 0.000 | 0.000 | 0.452 | 0.000 | 0.000 | 0.548 |
| IRIS_313-10429 | 0.000 | 0.000 | 0.000 | 0.000 | 1.000 | 0.000 | 0.000 |
| IRIS_313-10430 | 0.000 | 0.000 | 0.000 | 0.000 | 1.000 | 0.000 | 0.000 |
| IRIS_313-10433 | 0.000 | 0.000 | 0.000 | 0.455 | 0.000 | 0.000 | 0.545 |
| IRIS_313-10437 | 0.000 | 0.000 | 0.000 | 0.000 | 1.000 | 0.000 | 0.000 |
| IRIS_313-10440 | 0.000 | 0.000 | 0.064 | 0.000 | 0.936 | 0.000 | 0.000 |
| IRIS_313-10441 | 0.000 | 0.000 | 0.000 | 0.629 | 0.075 | 0.296 | 0.000 |
| IRIS_313-10444 | 0.000 | 0.703 | 0.000 | 0.061 | 0.000 | 0.236 | 0.000 |
| IRIS_313-10448 | 0.000 | 0.000 | 0.000 | 0.765 | 0.000 | 0.235 | 0.000 |
| IRIS_313-10449 | 0.000 | 0.000 | 0.000 | 0.042 | 0.176 | 0.783 | 0.000 |
| IRIS_313-10450 | 0.000 | 0.000 | 0.000 | 0.000 | 0.000 | 1.000 | 0.000 |
| IRIS_313-10452 | 0.000 | 0.000 | 0.000 | 0.000 | 0.000 | 1.000 | 0.000 |
| IRIS_313-10453 | 0.000 | 0.541 | 0.000 | 0.000 | 0.459 | 0.000 | 0.000 |
| IRIS_313-10458 | 0.000 | 0.000 | 0.000 | 0.063 | 0.000 | 0.937 | 0.000 |
| IRIS_313-10459 | 0.000 | 0.355 | 0.000 | 0.000 | 0.487 | 0.158 | 0.000 |
| IRIS_313-10469 | 0.000 | 0.000 | 0.000 | 0.000 | 1.000 | 0.000 | 0.000 |
| IRIS_313-10476 | 0.115 | 0.000 | 0.000 | 0.680 | 0.008 | 0.197 | 0.000 |
| IRIS_313-10477 | 0.000 | 0.000 | 0.030 | 0.463 | 0.000 | 0.507 | 0.000 |
| IRIS_313-10480 | 0.000 | 0.000 | 0.000 | 0.380 | 0.000 | 0.000 | 0.620 |
| IRIS_313-10484 | 0.000 | 0.406 | 0.000 | 0.000 | 0.000 | 0.594 | 0.000 |
| IRIS_313-10485 | 0.000 | 1.000 | 0.000 | 0.000 | 0.000 | 0.000 | 0.000 |
| IRIS_313-10489 | 0.000 | 1.000 | 0.000 | 0.000 | 0.000 | 0.000 | 0.000 |
| IRIS_313-10497 | 0.000 | 0.000 | 0.000 | 0.000 | 0.000 | 1.000 | 0.000 |
| IRIS_313-10502 | 0.000 | 0.000 | 0.000 | 0.021 | 0.000 | 0.979 | 0.000 |
| IRIS_313-10503 | 0.000 | 0.007 | 0.000 | 0.237 | 0.005 | 0.750 | 0.000 |
| IRIS_313-10504 | 0.000 | 0.000 | 0.000 | 0.000 | 0.000 | 1.000 | 0.000 |
| IRIS_313-10506 | 0.000 | 0.000 | 0.000 | 1.000 | 0.000 | 0.000 | 0.000 |
| IRIS_313-10507 | 0.000 | 0.704 | 0.000 | 0.000 | 0.226 | 0.071 | 0.000 |
| IRIS_313-10509 | 0.000 | 0.000 | 0.000 | 0.930 | 0.000 | 0.070 | 0.000 |
| IRIS_313-10510 | 0.000 | 0.114 | 0.000 | 0.637 | 0.000 | 0.249 | 0.000 |
| IRIS_313-10511 | 0.000 | 0.000 | 0.000 | 0.982 | 0.000 | 0.000 | 0.018 |
| IRIS_313-10513 | 0.000 | 0.000 | 0.000 | 0.748 | 0.000 | 0.252 | 0.000 |
| IRIS_313-10514 | 0.000 | 0.000 | 0.000 | 0.202 | 0.000 | 0.000 | 0.798 |
| IRIS_313-10515 | 0.000 | 0.000 | 0.000 | 0.000 | 0.000 | 1.000 | 0.000 |
| IRIS_313-10516 | 0.039 | 0.000 | 0.000 | 0.754 | 0.000 | 0.000 | 0.207 |
| IRIS_313-10517 | 0.000 | 0.014 | 0.000 | 0.835 | 0.000 | 0.151 | 0.000 |
| IRIS_313-10518 | 0.000 | 0.000 | 0.000 | 0.779 | 0.000 | 0.214 | 0.007 |
| IRIS_313-10519 | 0.142 | 0.000 | 0.000 | 0.858 | 0.000 | 0.000 | 0.000 |
| IRIS_313-10520 | 0.000 | 0.000 | 0.000 | 0.829 | 0.000 | 0.136 | 0.036 |
| IRIS_313-10521 | 0.000 | 0.000 | 1.000 | 0.000 | 0.000 | 0.000 | 0.000 |
| IRIS_313-10522 | 0.033 | 0.481 | 0.000 | 0.486 | 0.000 | 0.000 | 0.000 |
| IRIS_313-10523 | 0.145 | 0.000 | 0.022 | 0.823 | 0.010 | 0.000 | 0.000 |
| IRIS_313-10524 | 0.000 | 0.000 | 0.000 | 1.000 | 0.000 | 0.000 | 0.000 |
| IRIS_313-10525 | 0.000 | 0.000 | 0.000 | 0.434 | 0.000 | 0.000 | 0.567 |
| IRIS_313-10526 | 0.105 | 0.000 | 0.000 | 0.895 | 0.000 | 0.000 | 0.000 |
| IRIS_313-10527 | 0.154 | 0.002 | 0.002 | 0.804 | 0.000 | 0.000 | 0.037 |
| IRIS_313-10531 | 0.124 | 0.000 | 0.413 | 0.463 | 0.000 | 0.000 | 0.000 |
| IRIS_313-10534 | 1.000 | 0.000 | 0.000 | 0.000 | 0.000 | 0.000 | 0.000 |
| IRIS_313-10536 | 0.228 | 0.000 | 0.000 | 0.718 | 0.000 | 0.055 | 0.000 |
| IRIS_313-10537 | 0.000 | 0.143 | 0.547 | 0.000 | 0.157 | 0.040 | 0.113 |
| IRIS_313-10539 | 0.056 | 0.000 | 0.007 | 0.937 | 0.000 | 0.000 | 0.000 |

| | | | | | | | |
|---|---|---|---|---|---|---|---|
| IRIS_313-10541 | 0.000 | 1.000 | 0.000 | 0.000 | 0.000 | 0.000 | 0.000 |
| IRIS_313-10542 | 0.103 | 0.000 | 0.000 | 0.814 | 0.000 | 0.083 | 0.000 |
| IRIS_313-10543 | 0.983 | 0.000 | 0.000 | 0.000 | 0.017 | 0.000 | 0.000 |
| IRIS_313-10544 | 0.076 | 0.000 | 0.000 | 0.686 | 0.000 | 0.238 | 0.000 |
| IRIS_313-10545 | 0.921 | 0.000 | 0.000 | 0.000 | 0.000 | 0.000 | 0.079 |
| IRIS_313-10547 | 0.016 | 0.000 | 0.000 | 0.963 | 0.000 | 0.000 | 0.021 |
| IRIS_313-10549 | 1.000 | 0.000 | 0.000 | 0.000 | 0.000 | 0.000 | 0.000 |
| IRIS_313-10550 | 0.000 | 0.000 | 0.000 | 0.202 | 0.000 | 0.000 | 0.798 |
| IRIS_313-10552 | 0.000 | 1.000 | 0.000 | 0.000 | 0.000 | 0.000 | 0.000 |
| IRIS_313-10554 | 0.001 | 0.000 | 0.000 | 0.793 | 0.000 | 0.206 | 0.000 |
| IRIS_313-10555 | 0.000 | 0.000 | 0.000 | 0.994 | 0.006 | 0.000 | 0.000 |
| IRIS_313-10556 | 0.000 | 0.000 | 0.000 | 0.999 | 0.000 | 0.001 | 0.000 |
| IRIS_313-10557 | 0.000 | 0.000 | 0.000 | 0.972 | 0.000 | 0.000 | 0.028 |
| IRIS_313-10558 | 0.000 | 0.000 | 0.000 | 0.000 | 1.000 | 0.000 | 0.000 |
| IRIS_313-10559 | 0.000 | 0.000 | 0.000 | 0.000 | 1.000 | 0.000 | 0.000 |
| IRIS_313-10560 | 0.000 | 0.000 | 0.000 | 0.008 | 0.039 | 0.952 | 0.000 |
| IRIS_313-10561 | 0.000 | 0.000 | 0.000 | 0.000 | 0.000 | 1.000 | 0.000 |
| IRIS_313-10562 | 0.052 | 0.087 | 0.066 | 0.130 | 0.665 | 0.000 | 0.000 |
| IRIS_313-10563 | 0.000 | 0.000 | 0.000 | 0.000 | 1.000 | 0.000 | 0.000 |
| IRIS_313-10564 | 0.000 | 0.000 | 0.000 | 0.000 | 1.000 | 0.000 | 0.000 |
| IRIS_313-10565 | 0.000 | 0.000 | 0.000 | 0.000 | 0.000 | 0.000 | 1.000 |
| IRIS_313-10566 | 0.000 | 0.000 | 0.000 | 0.313 | 0.000 | 0.000 | 0.687 |
| IRIS_313-10567 | 0.000 | 0.000 | 0.000 | 0.000 | 1.000 | 0.000 | 0.000 |
| IRIS_313-10568 | 0.000 | 0.000 | 0.000 | 0.000 | 1.000 | 0.000 | 0.000 |
| IRIS_313-10569 | 0.000 | 0.000 | 0.000 | 0.000 | 1.000 | 0.000 | 0.000 |
| IRIS_313-10570 | 0.000 | 0.000 | 0.000 | 0.000 | 1.000 | 0.000 | 0.000 |
| IRIS_313-10572 | 0.074 | 0.000 | 0.000 | 0.602 | 0.013 | 0.078 | 0.233 |
| IRIS_313-10573 | 0.012 | 0.000 | 0.000 | 0.988 | 0.000 | 0.000 | 0.000 |
| IRIS_313-10574 | 0.018 | 0.000 | 0.000 | 0.982 | 0.000 | 0.000 | 0.000 |
| IRIS_313-10575 | 0.000 | 0.000 | 0.000 | 0.769 | 0.000 | 0.221 | 0.010 |
| IRIS_313-10576 | 0.033 | 0.000 | 0.000 | 0.966 | 0.000 | 0.000 | 0.000 |
| IRIS_313-10577 | 0.000 | 1.000 | 0.000 | 0.000 | 0.000 | 0.000 | 0.000 |
| IRIS_313-10578 | 0.000 | 1.000 | 0.000 | 0.000 | 0.000 | 0.000 | 0.000 |
| IRIS_313-10580 | 0.000 | 1.000 | 0.000 | 0.000 | 0.000 | 0.000 | 0.000 |
| IRIS_313-10581 | 0.000 | 1.000 | 0.000 | 0.000 | 0.000 | 0.000 | 0.000 |
| IRIS_313-10582 | 0.000 | 1.000 | 0.000 | 0.000 | 0.000 | 0.000 | 0.000 |
| IRIS_313-10583 | 0.000 | 0.000 | 0.000 | 0.000 | 1.000 | 0.000 | 0.000 |
| IRIS_313-10585 | 0.000 | 0.000 | 0.000 | 0.139 | 0.431 | 0.430 | 0.000 |
| IRIS_313-10587 | 1.000 | 0.000 | 0.000 | 0.000 | 0.000 | 0.000 | 0.000 |
| IRIS_313-10592 | 1.000 | 0.000 | 0.000 | 0.000 | 0.000 | 0.000 | 0.000 |
| IRIS_313-10593 | 1.000 | 0.000 | 0.000 | 0.000 | 0.000 | 0.000 | 0.000 |
| IRIS_313-10594 | 1.000 | 0.000 | 0.000 | 0.000 | 0.000 | 0.000 | 0.000 |
| IRIS_313-10595 | 1.000 | 0.000 | 0.000 | 0.000 | 0.000 | 0.000 | 0.000 |
| IRIS_313-10598 | 1.000 | 0.000 | 0.000 | 0.000 | 0.000 | 0.000 | 0.000 |
| IRIS_313-10600 | 1.000 | 0.000 | 0.000 | 0.000 | 0.000 | 0.000 | 0.000 |
| IRIS_313-10602 | 1.000 | 0.000 | 0.000 | 0.000 | 0.000 | 0.000 | 0.000 |
| IRIS_313-10603 | 1.000 | 0.000 | 0.000 | 0.000 | 0.000 | 0.000 | 0.000 |
| IRIS_313-10604 | 1.000 | 0.000 | 0.000 | 0.000 | 0.000 | 0.000 | 0.000 |
| IRIS_313-10605 | 1.000 | 0.000 | 0.000 | 0.000 | 0.000 | 0.000 | 0.000 |
| IRIS_313-10606 | 1.000 | 0.000 | 0.000 | 0.000 | 0.000 | 0.000 | 0.000 |
| IRIS_313-10608 | 1.000 | 0.000 | 0.000 | 0.000 | 0.000 | 0.000 | 0.000 |
| IRIS_313-10609 | 0.203 | 0.012 | 0.021 | 0.594 | 0.000 | 0.170 | 0.000 |
| IRIS_313-10610 | 0.123 | 0.000 | 0.000 | 0.659 | 0.000 | 0.217 | 0.000 |
| IRIS_313-10614 | 0.022 | 0.032 | 0.000 | 0.533 | 0.023 | 0.390 | 0.000 |
| IRIS_313-10617 | 0.000 | 0.055 | 0.000 | 0.000 | 0.945 | 0.000 | 0.000 |
| IRIS_313-10618 | 0.000 | 0.000 | 0.000 | 0.000 | 1.000 | 0.000 | 0.000 |
| IRIS_313-10619 | 0.000 | 1.000 | 0.000 | 0.000 | 0.000 | 0.000 | 0.000 |
| IRIS_313-10620 | 0.000 | 1.000 | 0.000 | 0.000 | 0.000 | 0.000 | 0.000 |
| IRIS_313-10623 | 0.983 | 0.000 | 0.000 | 0.017 | 0.000 | 0.000 | 0.000 |
| IRIS_313-10628 | 0.038 | 0.000 | 0.000 | 0.962 | 0.000 | 0.000 | 0.000 |
| IRIS_313-10631 | 0.000 | 0.000 | 0.000 | 0.000 | 1.000 | 0.000 | 0.000 |
| IRIS_313-10640 | 0.050 | 0.000 | 0.005 | 0.941 | 0.004 | 0.000 | 0.000 |
| IRIS_313-10642 | 0.000 | 0.000 | 0.000 | 0.000 | 1.000 | 0.000 | 0.000 |
| IRIS_313-10644 | 0.000 | 1.000 | 0.000 | 0.000 | 0.000 | 0.000 | 0.000 |
| IRIS_313-10645 | 0.000 | 1.000 | 0.000 | 0.000 | 0.000 | 0.000 | 0.000 |
| IRIS_313-10648 | 0.000 | 0.000 | 0.000 | 0.947 | 0.000 | 0.000 | 0.053 |
| IRIS_313-10649 | 0.000 | 0.984 | 0.000 | 0.006 | 0.000 | 0.007 | 0.003 |

| | | | | | | | |
|---|---|---|---|---|---|---|---|
| IRIS_313-10650 | 0.000 | 0.000 | 0.000 | 0.534 | 0.000 | 0.000 | 0.466 |
| IRIS_313-10651 | 0.000 | 0.000 | 0.000 | 0.848 | 0.000 | 0.152 | 0.000 |
| IRIS_313-10652 | 0.000 | 0.002 | 0.000 | 0.998 | 0.000 | 0.000 | 0.000 |
| IRIS_313-10653 | 0.000 | 0.697 | 0.000 | 0.000 | 0.303 | 0.000 | 0.000 |
| IRIS_313-10654 | 0.000 | 0.021 | 0.000 | 0.906 | 0.000 | 0.072 | 0.000 |
| IRIS_313-10655 | 0.000 | 0.010 | 0.000 | 0.819 | 0.000 | 0.172 | 0.000 |
| IRIS_313-10656 | 0.000 | 0.720 | 0.000 | 0.000 | 0.280 | 0.000 | 0.000 |
| IRIS_313-10657 | 0.000 | 0.654 | 0.000 | 0.000 | 0.346 | 0.000 | 0.000 |
| IRIS_313-10659 | 0.000 | 0.003 | 0.000 | 0.825 | 0.001 | 0.172 | 0.000 |
| IRIS_313-10660 | 0.000 | 0.670 | 0.000 | 0.000 | 0.330 | 0.000 | 0.000 |
| IRIS_313-10661 | 0.128 | 0.000 | 0.000 | 0.674 | 0.000 | 0.197 | 0.000 |
| IRIS_313-10662 | 0.155 | 0.000 | 0.000 | 0.697 | 0.000 | 0.147 | 0.000 |
| IRIS_313-10664 | 0.075 | 0.000 | 0.000 | 0.925 | 0.000 | 0.000 | 0.000 |
| IRIS_313-10666 | 0.000 | 0.000 | 0.000 | 1.000 | 0.000 | 0.000 | 0.000 |
| IRIS_313-10667 | 0.129 | 0.005 | 0.002 | 0.802 | 0.011 | 0.052 | 0.000 |
| IRIS_313-10668 | 0.080 | 0.000 | 0.000 | 0.920 | 0.000 | 0.000 | 0.000 |
| IRIS_313-10670 | 0.000 | 0.000 | 1.000 | 0.000 | 0.000 | 0.000 | 0.000 |
| IRIS_313-10671 | 0.228 | 0.000 | 0.000 | 0.412 | 0.000 | 0.360 | 0.000 |
| IRIS_313-10673 | 1.000 | 0.000 | 0.000 | 0.000 | 0.000 | 0.000 | 0.000 |
| IRIS_313-10674 | 0.000 | 0.000 | 0.000 | 0.873 | 0.000 | 0.127 | 0.000 |
| IRIS_313-10675 | 0.114 | 0.000 | 0.000 | 0.593 | 0.000 | 0.293 | 0.000 |
| IRIS_313-10676 | 0.391 | 0.000 | 0.019 | 0.241 | 0.000 | 0.349 | 0.000 |
| IRIS_313-10677 | 0.000 | 0.000 | 0.000 | 0.000 | 1.000 | 0.000 | 0.000 |
| IRIS_313-10678 | 0.000 | 0.000 | 0.000 | 1.000 | 0.000 | 0.000 | 0.000 |
| IRIS_313-10679 | 0.000 | 0.629 | 0.000 | 0.000 | 0.371 | 0.000 | 0.000 |
| IRIS_313-10680 | 0.000 | 0.000 | 0.000 | 0.904 | 0.000 | 0.096 | 0.000 |
| IRIS_313-10681 | 0.000 | 0.000 | 0.000 | 0.846 | 0.000 | 0.154 | 0.000 |
| IRIS_313-10682 | 0.000 | 0.000 | 0.000 | 0.939 | 0.000 | 0.061 | 0.000 |
| IRIS_313-10683 | 0.000 | 0.000 | 0.000 | 0.936 | 0.000 | 0.064 | 0.000 |
| IRIS_313-10684 | 0.000 | 0.000 | 0.000 | 0.435 | 0.000 | 0.000 | 0.565 |
| IRIS_313-10687 | 0.000 | 0.000 | 0.000 | 0.282 | 0.000 | 0.000 | 0.718 |
| IRIS_313-10688 | 0.000 | 0.000 | 0.000 | 0.540 | 0.000 | 0.000 | 0.460 |
| IRIS_313-10689 | 0.000 | 0.973 | 0.000 | 0.000 | 0.027 | 0.000 | 0.000 |
| IRIS_313-10690 | 0.000 | 0.154 | 0.000 | 0.300 | 0.000 | 0.000 | 0.546 |
| IRIS_313-10692 | 0.000 | 0.000 | 0.000 | 0.618 | 0.000 | 0.000 | 0.382 |
| IRIS_313-10693 | 0.000 | 1.000 | 0.000 | 0.000 | 0.000 | 0.000 | 0.000 |
| IRIS_313-10694 | 0.000 | 0.000 | 0.000 | 0.985 | 0.000 | 0.000 | 0.015 |
| IRIS_313-10695 | 0.000 | 0.005 | 0.000 | 0.972 | 0.000 | 0.000 | 0.023 |
| IRIS_313-10697 | 0.000 | 0.000 | 0.000 | 0.469 | 0.000 | 0.000 | 0.531 |
| IRIS_313-10698 | 0.000 | 0.000 | 0.000 | 0.600 | 0.000 | 0.000 | 0.400 |
| IRIS_313-10699 | 0.000 | 0.000 | 0.000 | 0.471 | 0.000 | 0.000 | 0.529 |
| IRIS_313-10700 | 0.000 | 0.000 | 0.000 | 0.447 | 0.001 | 0.000 | 0.551 |
| IRIS_313-10701 | 0.000 | 0.000 | 0.000 | 0.497 | 0.000 | 0.000 | 0.503 |
| IRIS_313-10702 | 0.000 | 0.006 | 0.000 | 0.528 | 0.000 | 0.000 | 0.466 |
| IRIS_313-10703 | 0.000 | 1.000 | 0.000 | 0.000 | 0.000 | 0.000 | 0.000 |
| IRIS_313-10704 | 0.000 | 1.000 | 0.000 | 0.000 | 0.000 | 0.000 | 0.000 |
| IRIS_313-10705 | 0.079 | 0.217 | 0.000 | 0.602 | 0.000 | 0.068 | 0.034 |
| IRIS_313-10706 | 0.000 | 0.000 | 0.000 | 0.596 | 0.000 | 0.000 | 0.404 |
| IRIS_313-10707 | 0.000 | 0.000 | 0.000 | 0.507 | 0.001 | 0.000 | 0.492 |
| IRIS_313-10708 | 0.000 | 1.000 | 0.000 | 0.000 | 0.000 | 0.000 | 0.000 |
| IRIS_313-10710 | 0.000 | 1.000 | 0.000 | 0.000 | 0.000 | 0.000 | 0.000 |
| IRIS_313-10711 | 0.000 | 1.000 | 0.000 | 0.000 | 0.000 | 0.000 | 0.000 |
| IRIS_313-10712 | 0.000 | 1.000 | 0.000 | 0.000 | 0.000 | 0.000 | 0.000 |
| IRIS_313-10715 | 0.134 | 0.013 | 0.017 | 0.594 | 0.000 | 0.243 | 0.000 |
| IRIS_313-10716 | 0.119 | 0.001 | 0.000 | 0.646 | 0.000 | 0.188 | 0.046 |
| IRIS_313-10717 | 0.186 | 0.000 | 0.000 | 0.812 | 0.000 | 0.000 | 0.002 |
| IRIS_313-10718 | 0.880 | 0.000 | 0.000 | 0.000 | 0.120 | 0.000 | 0.000 |
| IRIS_313-10719 | 0.131 | 0.013 | 0.000 | 0.694 | 0.000 | 0.163 | 0.000 |
| IRIS_313-10720 | 0.091 | 0.041 | 0.000 | 0.638 | 0.000 | 0.230 | 0.000 |
| IRIS_313-10721 | 0.128 | 0.005 | 0.000 | 0.653 | 0.000 | 0.214 | 0.000 |
| IRIS_313-10722 | 0.000 | 0.979 | 0.021 | 0.000 | 0.000 | 0.000 | 0.000 |
| IRIS_313-10723 | 0.030 | 0.000 | 0.017 | 0.953 | 0.000 | 0.000 | 0.000 |
| IRIS_313-10724 | 0.011 | 0.000 | 0.000 | 0.989 | 0.000 | 0.000 | 0.000 |
| IRIS_313-10725 | 0.044 | 0.000 | 0.000 | 0.956 | 0.000 | 0.000 | 0.000 |
| IRIS_313-10726 | 0.000 | 0.000 | 0.003 | 0.789 | 0.000 | 0.000 | 0.209 |
| IRIS_313-10727 | 0.000 | 0.000 | 0.000 | 0.996 | 0.000 | 0.000 | 0.004 |
| IRIS_313-10728 | 0.041 | 0.000 | 0.000 | 0.959 | 0.000 | 0.000 | 0.000 |

| | | | | | | | |
|---|---|---|---|---|---|---|---|
| IRIS_313-10729 | 0.298 | 0.000 | 0.120 | 0.241 | 0.340 | 0.000 | 0.000 |
| IRIS_313-10730 | 0.000 | 0.974 | 0.000 | 0.000 | 0.026 | 0.000 | 0.000 |
| IRIS_313-10731 | 0.000 | 0.000 | 0.000 | 0.007 | 0.000 | 0.993 | 0.000 |
| IRIS_313-10732 | 0.000 | 0.000 | 1.000 | 0.000 | 0.000 | 0.000 | 0.000 |
| IRIS_313-10733 | 0.009 | 0.000 | 0.000 | 0.776 | 0.000 | 0.215 | 0.000 |
| IRIS_313-10734 | 1.000 | 0.000 | 0.000 | 0.000 | 0.000 | 0.000 | 0.000 |
| IRIS_313-10735 | 1.000 | 0.000 | 0.000 | 0.000 | 0.000 | 0.000 | 0.000 |
| IRIS_313-10736 | 1.000 | 0.000 | 0.000 | 0.000 | 0.000 | 0.000 | 0.000 |
| IRIS_313-10737 | 0.946 | 0.020 | 0.000 | 0.000 | 0.000 | 0.000 | 0.035 |
| IRIS_313-10738 | 0.000 | 0.516 | 0.000 | 0.000 | 0.000 | 0.000 | 0.484 |
| IRIS_313-10739 | 0.000 | 0.359 | 0.000 | 0.000 | 0.000 | 0.415 | 0.226 |
| IRIS_313-10740 | 0.000 | 0.919 | 0.044 | 0.000 | 0.000 | 0.037 | 0.000 |
| IRIS_313-10741 | 0.000 | 0.919 | 0.046 | 0.000 | 0.000 | 0.036 | 0.000 |
| IRIS_313-10742 | 0.000 | 0.000 | 0.000 | 0.000 | 0.000 | 0.062 | 0.938 |
| IRIS_313-10743 | 0.000 | 0.972 | 0.000 | 0.000 | 0.028 | 0.000 | 0.000 |
| IRIS_313-10744 | 0.000 | 0.952 | 0.000 | 0.000 | 0.000 | 0.048 | 0.000 |
| IRIS_313-10745 | 0.000 | 0.928 | 0.072 | 0.000 | 0.000 | 0.000 | 0.000 |
| IRIS_313-10746 | 0.000 | 0.938 | 0.040 | 0.000 | 0.000 | 0.011 | 0.011 |
| IRIS_313-10747 | 0.000 | 1.000 | 0.000 | 0.000 | 0.000 | 0.000 | 0.000 |
| IRIS_313-10748 | 0.000 | 0.000 | 0.000 | 0.665 | 0.000 | 0.335 | 0.000 |
| IRIS_313-10749 | 0.000 | 0.000 | 0.000 | 0.989 | 0.000 | 0.000 | 0.011 |
| IRIS_313-10750 | 0.000 | 0.000 | 0.000 | 0.940 | 0.000 | 0.000 | 0.060 |
| IRIS_313-10751 | 0.029 | 0.000 | 0.017 | 0.388 | 0.000 | 0.345 | 0.220 |
| IRIS_313-10752 | 0.000 | 0.903 | 0.000 | 0.000 | 0.097 | 0.000 | 0.000 |
| IRIS_313-10753 | 0.000 | 0.000 | 0.000 | 0.000 | 0.045 | 0.164 | 0.791 |
| IRIS_313-10754 | 0.000 | 0.000 | 0.000 | 0.000 | 0.000 | 0.416 | 0.584 |
| IRIS_313-10755 | 0.000 | 0.000 | 0.000 | 0.901 | 0.000 | 0.000 | 0.099 |
| IRIS_313-10756 | 0.053 | 0.000 | 0.000 | 0.908 | 0.000 | 0.000 | 0.039 |
| IRIS_313-10757 | 0.000 | 0.000 | 0.000 | 0.584 | 0.000 | 0.000 | 0.416 |
| IRIS_313-10758 | 0.000 | 0.966 | 0.000 | 0.000 | 0.021 | 0.014 | 0.000 |
| IRIS_313-10759 | 0.000 | 0.916 | 0.039 | 0.000 | 0.000 | 0.045 | 0.000 |
| IRIS_313-10760 | 0.000 | 0.000 | 0.000 | 0.458 | 0.000 | 0.000 | 0.542 |
| IRIS_313-10761 | 0.000 | 1.000 | 0.000 | 0.000 | 0.000 | 0.000 | 0.000 |
| IRIS_313-10762 | 0.000 | 0.002 | 0.002 | 0.657 | 0.000 | 0.000 | 0.339 |
| IRIS_313-10765 | 0.000 | 0.846 | 0.030 | 0.002 | 0.076 | 0.045 | 0.000 |
| IRIS_313-10766 | 0.000 | 1.000 | 0.000 | 0.000 | 0.000 | 0.000 | 0.000 |
| IRIS_313-10767 | 0.000 | 1.000 | 0.000 | 0.000 | 0.000 | 0.000 | 0.000 |
| IRIS_313-10768 | 0.000 | 0.008 | 0.000 | 0.344 | 0.000 | 0.000 | 0.648 |
| IRIS_313-10769 | 0.000 | 0.000 | 0.000 | 0.282 | 0.000 | 0.000 | 0.718 |
| IRIS_313-10770 | 0.000 | 0.951 | 0.000 | 0.000 | 0.000 | 0.049 | 0.000 |
| IRIS_313-10771 | 0.000 | 0.698 | 0.000 | 0.293 | 0.009 | 0.000 | 0.000 |
| IRIS_313-10772 | 0.000 | 0.000 | 0.000 | 0.066 | 0.000 | 0.000 | 0.934 |
| IRIS_313-10773 | 0.000 | 0.977 | 0.023 | 0.000 | 0.000 | 0.000 | 0.000 |
| IRIS_313-10774 | 0.000 | 0.000 | 0.000 | 0.204 | 0.000 | 0.000 | 0.796 |
| IRIS_313-10775 | 0.000 | 0.000 | 0.000 | 0.351 | 0.000 | 0.000 | 0.649 |
| IRIS_313-10776 | 0.000 | 1.000 | 0.000 | 0.000 | 0.000 | 0.000 | 0.000 |
| IRIS_313-10777 | 0.000 | 0.023 | 0.000 | 0.250 | 0.000 | 0.000 | 0.727 |
| IRIS_313-10778 | 0.000 | 0.006 | 0.000 | 0.420 | 0.000 | 0.000 | 0.574 |
| IRIS_313-10779 | 0.000 | 0.000 | 0.000 | 0.263 | 0.000 | 0.000 | 0.737 |
| IRIS_313-10780 | 0.000 | 0.908 | 0.000 | 0.000 | 0.092 | 0.000 | 0.000 |
| IRIS_313-10781 | 0.000 | 0.955 | 0.045 | 0.000 | 0.000 | 0.000 | 0.000 |
| IRIS_313-10782 | 0.000 | 0.011 | 0.000 | 0.491 | 0.000 | 0.000 | 0.498 |
| IRIS_313-10783 | 0.000 | 0.880 | 0.000 | 0.000 | 0.057 | 0.063 | 0.000 |
| IRIS_313-10784 | 0.000 | 0.926 | 0.029 | 0.000 | 0.000 | 0.045 | 0.000 |
| IRIS_313-10785 | 0.000 | 0.897 | 0.056 | 0.000 | 0.000 | 0.047 | 0.000 |
| IRIS_313-10786 | 0.000 | 0.129 | 0.000 | 0.282 | 0.076 | 0.000 | 0.512 |
| IRIS_313-10787 | 0.000 | 0.421 | 0.000 | 0.314 | 0.000 | 0.000 | 0.265 |
| IRIS_313-10788 | 0.000 | 0.910 | 0.037 | 0.000 | 0.044 | 0.009 | 0.000 |
| IRIS_313-10789 | 0.000 | 0.961 | 0.039 | 0.000 | 0.000 | 0.000 | 0.000 |
| IRIS_313-10790 | 0.000 | 0.881 | 0.070 | 0.000 | 0.015 | 0.034 | 0.000 |
| IRIS_313-10791 | 0.000 | 0.016 | 0.000 | 0.372 | 0.000 | 0.000 | 0.613 |
| IRIS_313-10792 | 0.000 | 0.000 | 0.000 | 0.408 | 0.000 | 0.000 | 0.592 |
| IRIS_313-10793 | 0.000 | 0.978 | 0.022 | 0.000 | 0.000 | 0.000 | 0.000 |
| IRIS_313-10794 | 0.000 | 0.948 | 0.000 | 0.000 | 0.052 | 0.000 | 0.000 |
| IRIS_313-10795 | 0.000 | 1.000 | 0.000 | 0.000 | 0.000 | 0.000 | 0.000 |
| IRIS_313-10796 | 0.000 | 0.978 | 0.000 | 0.000 | 0.022 | 0.000 | 0.000 |
| IRIS_313-10797 | 0.000 | 0.006 | 0.000 | 0.269 | 0.000 | 0.000 | 0.725 |

| | | | | | | | |
|---|---|---|---|---|---|---|---|
| IRIS_313-10798 | 0.000 | 1.000 | 0.000 | 0.000 | 0.000 | 0.000 | 0.000 |
| IRIS_313-10799 | 0.000 | 1.000 | 0.000 | 0.000 | 0.000 | 0.000 | 0.000 |
| IRIS_313-10800 | 0.000 | 0.951 | 0.000 | 0.000 | 0.000 | 0.049 | 0.000 |
| IRIS_313-10801 | 0.000 | 0.897 | 0.000 | 0.000 | 0.103 | 0.000 | 0.000 |
| IRIS_313-10802 | 0.000 | 0.972 | 0.000 | 0.000 | 0.000 | 0.000 | 0.028 |
| IRIS_313-10803 | 0.000 | 0.893 | 0.000 | 0.000 | 0.059 | 0.048 | 0.000 |
| IRIS_313-10804 | 0.000 | 0.007 | 0.000 | 0.391 | 0.000 | 0.000 | 0.601 |
| IRIS_313-10805 | 0.000 | 0.961 | 0.000 | 0.004 | 0.000 | 0.035 | 0.000 |
| IRIS_313-10806 | 0.002 | 0.000 | 0.000 | 0.510 | 0.000 | 0.000 | 0.488 |
| IRIS_313-10807 | 0.000 | 0.021 | 0.000 | 0.463 | 0.000 | 0.000 | 0.516 |
| IRIS_313-10808 | 0.000 | 0.959 | 0.000 | 0.000 | 0.000 | 0.041 | 0.000 |
| IRIS_313-10809 | 0.000 | 0.805 | 0.000 | 0.000 | 0.058 | 0.089 | 0.048 |
| IRIS_313-10810 | 0.000 | 0.000 | 0.000 | 0.256 | 0.000 | 0.000 | 0.744 |
| IRIS_313-10811 | 0.000 | 0.000 | 0.000 | 0.386 | 0.000 | 0.000 | 0.614 |
| IRIS_313-10812 | 0.000 | 0.000 | 0.000 | 0.162 | 0.000 | 0.000 | 0.838 |
| IRIS_313-10813 | 0.000 | 0.015 | 0.000 | 0.499 | 0.000 | 0.000 | 0.486 |
| IRIS_313-10814 | 0.000 | 0.022 | 0.000 | 0.360 | 0.001 | 0.000 | 0.616 |
| IRIS_313-10815 | 0.000 | 0.930 | 0.000 | 0.000 | 0.036 | 0.034 | 0.000 |
| IRIS_313-10816 | 0.000 | 0.919 | 0.000 | 0.000 | 0.000 | 0.081 | 0.000 |
| IRIS_313-10817 | 0.000 | 0.872 | 0.076 | 0.004 | 0.000 | 0.049 | 0.000 |
| IRIS_313-10818 | 0.000 | 0.000 | 0.000 | 0.601 | 0.028 | 0.000 | 0.371 |
| IRIS_313-10819 | 0.000 | 0.008 | 0.008 | 0.467 | 0.011 | 0.000 | 0.505 |
| IRIS_313-10820 | 0.000 | 0.027 | 0.000 | 0.313 | 0.000 | 0.000 | 0.661 |
| IRIS_313-10821 | 0.000 | 0.000 | 0.000 | 0.545 | 0.000 | 0.000 | 0.455 |
| IRIS_313-10822 | 0.000 | 0.000 | 0.000 | 0.415 | 0.000 | 0.018 | 0.567 |
| IRIS_313-10823 | 0.000 | 0.010 | 0.000 | 0.454 | 0.004 | 0.000 | 0.531 |
| IRIS_313-10824 | 0.000 | 0.000 | 0.000 | 0.000 | 0.000 | 0.000 | 1.000 |
| IRIS_313-10825 | 0.038 | 0.000 | 0.006 | 0.955 | 0.000 | 0.000 | 0.000 |
| IRIS_313-10827 | 0.000 | 1.000 | 0.000 | 0.000 | 0.000 | 0.000 | 0.000 |
| IRIS_313-10828 | 0.000 | 1.000 | 0.000 | 0.000 | 0.000 | 0.000 | 0.000 |
| IRIS_313-10829 | 0.000 | 1.000 | 0.000 | 0.000 | 0.000 | 0.000 | 0.000 |
| IRIS_313-10830 | 0.000 | 1.000 | 0.000 | 0.000 | 0.000 | 0.000 | 0.000 |
| IRIS_313-10831 | 0.000 | 1.000 | 0.000 | 0.000 | 0.000 | 0.000 | 0.000 |
| IRIS_313-10832 | 0.000 | 1.000 | 0.000 | 0.000 | 0.000 | 0.000 | 0.000 |
| IRIS_313-10833 | 0.049 | 0.000 | 0.000 | 0.713 | 0.000 | 0.238 | 0.000 |
| IRIS_313-10834 | 0.000 | 0.495 | 0.039 | 0.000 | 0.466 | 0.000 | 0.000 |
| IRIS_313-10835 | 0.062 | 0.000 | 0.000 | 0.645 | 0.003 | 0.290 | 0.000 |
| IRIS_313-10836 | 0.039 | 0.000 | 0.000 | 0.694 | 0.000 | 0.267 | 0.000 |
| IRIS_313-10837 | 0.000 | 0.000 | 0.000 | 0.000 | 1.000 | 0.000 | 0.000 |
| IRIS_313-10838 | 0.000 | 0.000 | 0.000 | 0.000 | 1.000 | 0.000 | 0.000 |
| IRIS_313-10839 | 0.000 | 0.000 | 0.000 | 0.000 | 1.000 | 0.000 | 0.000 |
| IRIS_313-10840 | 0.000 | 0.000 | 0.000 | 0.000 | 1.000 | 0.000 | 0.000 |
| IRIS_313-10841 | 0.000 | 0.964 | 0.000 | 0.000 | 0.000 | 0.036 | 0.000 |
| IRIS_313-10842 | 0.000 | 0.000 | 0.000 | 0.302 | 0.000 | 0.000 | 0.698 |
| IRIS_313-10844 | 0.000 | 0.001 | 0.000 | 0.428 | 0.002 | 0.000 | 0.569 |
| IRIS_313-10845 | 0.905 | 0.000 | 0.000 | 0.094 | 0.000 | 0.001 | 0.000 |
| IRIS_313-10847 | 0.000 | 0.001 | 0.000 | 0.999 | 0.000 | 0.000 | 0.000 |
| IRIS_313-10848 | 0.032 | 0.590 | 0.022 | 0.002 | 0.350 | 0.000 | 0.005 |
| IRIS_313-10849 | 1.000 | 0.000 | 0.000 | 0.000 | 0.000 | 0.000 | 0.000 |
| IRIS_313-10850 | 0.166 | 0.479 | 0.000 | 0.327 | 0.028 | 0.000 | 0.000 |
| IRIS_313-10851 | 0.000 | 0.074 | 0.900 | 0.000 | 0.000 | 0.026 | 0.000 |
| IRIS_313-10852 | 1.000 | 0.000 | 0.000 | 0.000 | 0.000 | 0.000 | 0.000 |
| IRIS_313-10853 | 0.000 | 0.809 | 0.000 | 0.000 | 0.191 | 0.000 | 0.000 |
| IRIS_313-10854 | 0.884 | 0.011 | 0.000 | 0.002 | 0.000 | 0.066 | 0.039 |
| IRIS_313-10855 | 0.047 | 0.000 | 0.000 | 0.887 | 0.000 | 0.066 | 0.000 |
| IRIS_313-10856 | 0.343 | 0.010 | 0.000 | 0.476 | 0.000 | 0.172 | 0.000 |
| IRIS_313-10857 | 0.343 | 0.035 | 0.003 | 0.301 | 0.000 | 0.318 | 0.000 |
| IRIS_313-10858 | 0.109 | 0.004 | 0.005 | 0.680 | 0.000 | 0.202 | 0.000 |
| IRIS_313-10859 | 0.284 | 0.000 | 0.000 | 0.419 | 0.000 | 0.297 | 0.000 |
| IRIS_313-10861 | 0.911 | 0.014 | 0.000 | 0.002 | 0.000 | 0.037 | 0.037 |
| IRIS_313-10862 | 0.000 | 0.801 | 0.000 | 0.000 | 0.199 | 0.000 | 0.000 |
| IRIS_313-10863 | 0.047 | 0.000 | 0.000 | 0.953 | 0.000 | 0.000 | 0.000 |
| IRIS_313-10864 | 0.000 | 0.457 | 0.450 | 0.000 | 0.093 | 0.000 | 0.000 |
| IRIS_313-10865 | 0.001 | 0.635 | 0.010 | 0.000 | 0.348 | 0.006 | 0.000 |
| IRIS_313-10866 | 0.000 | 0.673 | 0.000 | 0.000 | 0.322 | 0.005 | 0.000 |
| IRIS_313-10867 | 0.000 | 0.724 | 0.000 | 0.000 | 0.276 | 0.000 | 0.000 |
| IRIS_313-10868 | 0.000 | 0.000 | 1.000 | 0.000 | 0.000 | 0.000 | 0.000 |

| | | | | | | | |
|---|---|---|---|---|---|---|---|
| IRIS_313-10869 | 1.000 | 0.000 | 0.000 | 0.000 | 0.000 | 0.000 | 0.000 |
| IRIS_313-10870 | 0.000 | 0.777 | 0.031 | 0.000 | 0.192 | 0.000 | 0.000 |
| IRIS_313-10871 | 0.957 | 0.043 | 0.000 | 0.000 | 0.000 | 0.000 | 0.000 |
| IRIS_313-10872 | 0.000 | 0.557 | 0.000 | 0.000 | 0.228 | 0.048 | 0.167 |
| IRIS_313-10873 | 0.748 | 0.166 | 0.000 | 0.000 | 0.038 | 0.000 | 0.048 |
| IRIS_313-10874 | 0.000 | 0.717 | 0.000 | 0.000 | 0.283 | 0.000 | 0.000 |
| IRIS_313-10875 | 1.000 | 0.000 | 0.000 | 0.000 | 0.000 | 0.000 | 0.000 |
| IRIS_313-10876 | 0.963 | 0.037 | 0.000 | 0.000 | 0.000 | 0.000 | 0.000 |
| IRIS_313-10877 | 0.969 | 0.031 | 0.000 | 0.000 | 0.000 | 0.000 | 0.000 |
| IRIS_313-10878 | 0.970 | 0.030 | 0.000 | 0.000 | 0.000 | 0.000 | 0.000 |
| IRIS_313-10879 | 0.605 | 0.000 | 0.000 | 0.269 | 0.000 | 0.126 | 0.000 |
| IRIS_313-10880 | 0.024 | 0.000 | 0.000 | 0.976 | 0.000 | 0.000 | 0.000 |
| IRIS_313-10881 | 0.373 | 0.400 | 0.000 | 0.000 | 0.080 | 0.147 | 0.000 |
| IRIS_313-10882 | 0.875 | 0.000 | 0.000 | 0.000 | 0.000 | 0.000 | 0.125 |
| IRIS_313-10883 | 0.333 | 0.503 | 0.000 | 0.000 | 0.058 | 0.106 | 0.000 |
| IRIS_313-10884 | 0.000 | 0.699 | 0.042 | 0.000 | 0.233 | 0.026 | 0.000 |
| IRIS_313-10885 | 0.000 | 0.000 | 0.000 | 0.955 | 0.000 | 0.045 | 0.000 |
| IRIS_313-10886 | 0.000 | 0.704 | 0.022 | 0.002 | 0.268 | 0.003 | 0.001 |
| IRIS_313-10887 | 0.000 | 0.000 | 0.000 | 0.828 | 0.000 | 0.172 | 0.000 |
| IRIS_313-10888 | 0.000 | 0.785 | 0.000 | 0.000 | 0.215 | 0.000 | 0.000 |
| IRIS_313-10889 | 0.004 | 0.696 | 0.000 | 0.000 | 0.297 | 0.002 | 0.001 |
| IRIS_313-10890 | 0.000 | 0.683 | 0.000 | 0.000 | 0.317 | 0.000 | 0.000 |
| IRIS_313-10891 | 0.795 | 0.000 | 0.001 | 0.000 | 0.000 | 0.204 | 0.000 |
| IRIS_313-10892 | 0.825 | 0.010 | 0.000 | 0.001 | 0.000 | 0.117 | 0.046 |
| IRIS_313-10893 | 0.000 | 0.716 | 0.000 | 0.000 | 0.284 | 0.000 | 0.000 |
| IRIS_313-10894 | 0.566 | 0.068 | 0.000 | 0.000 | 0.000 | 0.248 | 0.117 |
| IRIS_313-10895 | 0.000 | 0.781 | 0.000 | 0.000 | 0.219 | 0.000 | 0.000 |
| IRIS_313-10896 | 0.516 | 0.079 | 0.000 | 0.000 | 0.003 | 0.297 | 0.106 |
| IRIS_313-10897 | 0.000 | 0.000 | 0.000 | 0.997 | 0.003 | 0.000 | 0.000 |
| IRIS_313-10898 | 0.000 | 0.000 | 0.000 | 0.905 | 0.000 | 0.000 | 0.095 |
| IRIS_313-10899 | 0.000 | 0.000 | 0.000 | 1.000 | 0.000 | 0.000 | 0.000 |
| IRIS_313-10900 | 0.000 | 0.000 | 0.000 | 1.000 | 0.000 | 0.000 | 0.000 |
| IRIS_313-10901 | 0.000 | 0.000 | 0.000 | 1.000 | 0.000 | 0.000 | 0.000 |
| IRIS_313-10902 | 0.000 | 0.000 | 0.000 | 0.998 | 0.000 | 0.000 | 0.002 |
| IRIS_313-10903 | 0.000 | 0.000 | 0.000 | 0.915 | 0.000 | 0.000 | 0.085 |
| IRIS_313-10904 | 0.000 | 0.003 | 0.000 | 0.947 | 0.001 | 0.049 | 0.000 |
| IRIS_313-10905 | 0.000 | 0.000 | 0.000 | 0.996 | 0.004 | 0.000 | 0.000 |
| IRIS_313-10906 | 0.000 | 0.000 | 0.000 | 0.938 | 0.020 | 0.000 | 0.042 |
| IRIS_313-10907 | 0.000 | 0.000 | 0.000 | 0.921 | 0.000 | 0.079 | 0.000 |
| IRIS_313-10908 | 0.000 | 0.000 | 0.000 | 1.000 | 0.000 | 0.000 | 0.000 |
| IRIS_313-10909 | 0.000 | 0.000 | 0.000 | 1.000 | 0.000 | 0.000 | 0.000 |
| IRIS_313-10910 | 0.000 | 0.000 | 0.000 | 1.000 | 0.000 | 0.000 | 0.000 |
| IRIS_313-10911 | 0.000 | 0.000 | 0.000 | 0.964 | 0.000 | 0.036 | 0.000 |
| IRIS_313-10912 | 0.000 | 0.000 | 0.000 | 1.000 | 0.000 | 0.000 | 0.000 |
| IRIS_313-10913 | 0.000 | 0.000 | 0.000 | 0.938 | 0.000 | 0.000 | 0.062 |
| IRIS_313-10915 | 0.000 | 0.000 | 0.000 | 0.998 | 0.000 | 0.000 | 0.002 |
| IRIS_313-10916 | 0.000 | 0.230 | 0.000 | 0.000 | 0.770 | 0.000 | 0.000 |
| IRIS_313-10917 | 0.000 | 0.000 | 0.000 | 0.493 | 0.000 | 0.507 | 0.000 |
| IRIS_313-10918 | 0.000 | 0.853 | 0.000 | 0.000 | 0.000 | 0.000 | 0.147 |
| IRIS_313-10919 | 0.000 | 1.000 | 0.000 | 0.000 | 0.000 | 0.000 | 0.000 |
| IRIS_313-10920 | 0.000 | 0.986 | 0.000 | 0.006 | 0.000 | 0.008 | 0.000 |
| IRIS_313-10921 | 0.000 | 0.022 | 0.000 | 0.978 | 0.000 | 0.000 | 0.000 |
| IRIS_313-10922 | 0.000 | 0.710 | 0.000 | 0.000 | 0.290 | 0.000 | 0.000 |
| IRIS_313-10923 | 0.000 | 0.627 | 0.000 | 0.000 | 0.373 | 0.000 | 0.000 |
| IRIS_313-10924 | 0.000 | 0.000 | 0.000 | 0.745 | 0.000 | 0.255 | 0.000 |
| IRIS_313-10925 | 1.000 | 0.000 | 0.000 | 0.000 | 0.000 | 0.000 | 0.000 |
| IRIS_313-10926 | 0.000 | 0.000 | 1.000 | 0.000 | 0.000 | 0.000 | 0.000 |
| IRIS_313-10927 | 1.000 | 0.000 | 0.000 | 0.000 | 0.000 | 0.000 | 0.000 |
| IRIS_313-10928 | 0.000 | 0.000 | 0.000 | 0.883 | 0.001 | 0.041 | 0.075 |
| IRIS_313-10929 | 0.017 | 0.000 | 0.000 | 0.974 | 0.000 | 0.009 | 0.000 |
| IRIS_313-10930 | 1.000 | 0.000 | 0.000 | 0.000 | 0.000 | 0.000 | 0.000 |
| IRIS_313-10931 | 0.000 | 1.000 | 0.000 | 0.000 | 0.000 | 0.000 | 0.000 |
| IRIS_313-10932 | 0.000 | 0.000 | 0.000 | 0.000 | 0.000 | 1.000 | 0.000 |
| IRIS_313-10933 | 0.000 | 0.065 | 0.935 | 0.000 | 0.000 | 0.000 | 0.000 |
| IRIS_313-10934 | 0.000 | 0.000 | 0.000 | 1.000 | 0.000 | 0.000 | 0.000 |
| IRIS_313-10935 | 0.000 | 0.000 | 0.000 | 0.308 | 0.000 | 0.000 | 0.692 |
| IRIS_313-10936 | 0.000 | 0.918 | 0.082 | 0.000 | 0.000 | 0.000 | 0.000 |

| | | | | | | | |
|---|---|---|---|---|---|---|---|
| IRIS_313-10937 | 0.000 | 0.000 | 0.000 | 0.589 | 0.000 | 0.000 | 0.411 |
| IRIS_313-10938 | 0.000 | 0.000 | 0.000 | 0.336 | 0.000 | 0.000 | 0.664 |
| IRIS_313-10940 | 0.000 | 0.000 | 0.000 | 0.607 | 0.000 | 0.000 | 0.393 |
| IRIS_313-10941 | 0.000 | 0.000 | 0.000 | 0.124 | 0.000 | 0.000 | 0.876 |
| IRIS_313-10942 | 0.000 | 0.000 | 0.000 | 0.548 | 0.000 | 0.000 | 0.452 |
| IRIS_313-10943 | 0.000 | 0.000 | 0.000 | 0.222 | 0.000 | 0.000 | 0.778 |
| IRIS_313-10944 | 0.059 | 0.008 | 0.000 | 0.657 | 0.000 | 0.000 | 0.277 |
| IRIS_313-10945 | 0.000 | 1.000 | 0.000 | 0.000 | 0.000 | 0.000 | 0.000 |
| IRIS_313-10946 | 0.000 | 0.992 | 0.000 | 0.000 | 0.000 | 0.000 | 0.008 |
| IRIS_313-10947 | 0.000 | 0.043 | 0.000 | 0.660 | 0.000 | 0.059 | 0.239 |
| IRIS_313-10948 | 0.000 | 0.504 | 0.000 | 0.000 | 0.000 | 0.000 | 0.496 |
| IRIS_313-10949 | 0.000 | 1.000 | 0.000 | 0.000 | 0.000 | 0.000 | 0.000 |
| IRIS_313-10950 | 0.000 | 0.868 | 0.095 | 0.000 | 0.000 | 0.037 | 0.000 |
| IRIS_313-10951 | 0.000 | 0.000 | 0.000 | 0.439 | 0.000 | 0.000 | 0.561 |
| IRIS_313-10952 | 0.000 | 0.990 | 0.000 | 0.000 | 0.000 | 0.010 | 0.000 |
| IRIS_313-10953 | 0.000 | 0.993 | 0.007 | 0.000 | 0.000 | 0.000 | 0.000 |
| IRIS_313-10954 | 0.000 | 0.000 | 0.000 | 0.364 | 0.000 | 0.000 | 0.636 |
| IRIS_313-10955 | 0.000 | 0.010 | 0.000 | 0.626 | 0.000 | 0.000 | 0.364 |
| IRIS_313-10956 | 0.000 | 0.966 | 0.000 | 0.000 | 0.000 | 0.034 | 0.000 |
| IRIS_313-10957 | 0.000 | 0.991 | 0.000 | 0.000 | 0.000 | 0.004 | 0.004 |
| IRIS_313-10958 | 0.000 | 0.997 | 0.000 | 0.000 | 0.000 | 0.003 | 0.000 |
| IRIS_313-10959 | 0.000 | 1.000 | 0.000 | 0.000 | 0.000 | 0.000 | 0.000 |
| IRIS_313-10960 | 0.000 | 0.977 | 0.000 | 0.000 | 0.000 | 0.023 | 0.000 |
| IRIS_313-10961 | 0.000 | 0.000 | 0.000 | 0.168 | 0.000 | 0.000 | 0.832 |
| IRIS_313-10962 | 0.000 | 0.000 | 0.000 | 0.215 | 0.000 | 0.000 | 0.785 |
| IRIS_313-10963 | 1.000 | 0.000 | 0.000 | 0.000 | 0.000 | 0.000 | 0.000 |
| IRIS_313-10964 | 1.000 | 0.000 | 0.000 | 0.000 | 0.000 | 0.000 | 0.000 |
| IRIS_313-10965 | 1.000 | 0.000 | 0.000 | 0.000 | 0.000 | 0.000 | 0.000 |
| IRIS_313-10966 | 0.000 | 0.000 | 0.000 | 0.622 | 0.000 | 0.378 | 0.000 |
| IRIS_313-10967 | 0.000 | 0.208 | 0.000 | 0.000 | 0.792 | 0.000 | 0.000 |
| IRIS_313-10968 | 0.000 | 0.000 | 0.000 | 0.000 | 0.000 | 1.000 | 0.000 |
| IRIS_313-10969 | 1.000 | 0.000 | 0.000 | 0.000 | 0.000 | 0.000 | 0.000 |
| IRIS_313-10970 | 0.000 | 0.000 | 0.000 | 0.999 | 0.000 | 0.000 | 0.001 |
| IRIS_313-10971 | 0.035 | 0.000 | 0.002 | 0.857 | 0.000 | 0.107 | 0.000 |
| IRIS_313-10972 | 0.058 | 0.000 | 0.001 | 0.941 | 0.000 | 0.000 | 0.000 |
| IRIS_313-10973 | 0.070 | 0.000 | 0.019 | 0.911 | 0.000 | 0.000 | 0.000 |
| IRIS_313-10974 | 0.033 | 0.000 | 0.003 | 0.964 | 0.000 | 0.000 | 0.000 |
| IRIS_313-10975 | 0.000 | 0.000 | 0.000 | 1.000 | 0.000 | 0.000 | 0.000 |
| IRIS_313-10976 | 1.000 | 0.000 | 0.000 | 0.000 | 0.000 | 0.000 | 0.000 |
| IRIS_313-10977 | 0.078 | 0.000 | 0.011 | 0.753 | 0.000 | 0.158 | 0.000 |
| IRIS_313-10978 | 0.107 | 0.000 | 0.000 | 0.876 | 0.000 | 0.000 | 0.017 |
| IRIS_313-10979 | 0.845 | 0.000 | 0.019 | 0.136 | 0.000 | 0.000 | 0.000 |
| IRIS_313-10980 | 0.440 | 0.000 | 0.000 | 0.560 | 0.000 | 0.000 | 0.000 |
| IRIS_313-10981 | 0.894 | 0.000 | 0.020 | 0.086 | 0.000 | 0.000 | 0.000 |
| IRIS_313-10982 | 0.038 | 0.000 | 0.000 | 0.962 | 0.000 | 0.000 | 0.000 |
| IRIS_313-10983 | 0.000 | 0.000 | 0.000 | 0.931 | 0.000 | 0.069 | 0.000 |
| IRIS_313-10984 | 0.045 | 0.000 | 0.000 | 0.955 | 0.000 | 0.000 | 0.000 |
| IRIS_313-10985 | 0.057 | 0.000 | 0.000 | 0.943 | 0.000 | 0.000 | 0.000 |
| IRIS_313-10986 | 0.033 | 0.000 | 0.000 | 0.967 | 0.000 | 0.000 | 0.000 |
| IRIS_313-10987 | 0.887 | 0.000 | 0.000 | 0.109 | 0.000 | 0.003 | 0.001 |
| IRIS_313-10988 | 0.000 | 0.066 | 0.000 | 0.706 | 0.000 | 0.000 | 0.228 |
| IRIS_313-10989 | 0.012 | 0.000 | 0.000 | 0.988 | 0.000 | 0.000 | 0.000 |
| IRIS_313-10990 | 0.000 | 0.013 | 0.000 | 0.725 | 0.007 | 0.243 | 0.012 |
| IRIS_313-10991 | 0.000 | 1.000 | 0.000 | 0.000 | 0.000 | 0.000 | 0.000 |
| IRIS_313-10992 | 0.000 | 0.987 | 0.000 | 0.000 | 0.013 | 0.000 | 0.000 |
| IRIS_313-10993 | 0.000 | 1.000 | 0.000 | 0.000 | 0.000 | 0.000 | 0.000 |
| IRIS_313-10994 | 0.000 | 0.985 | 0.000 | 0.005 | 0.000 | 0.007 | 0.003 |
| IRIS_313-10995 | 0.000 | 0.000 | 0.000 | 0.752 | 0.000 | 0.000 | 0.248 |
| IRIS_313-10996 | 0.000 | 0.000 | 0.000 | 0.000 | 0.000 | 0.000 | 1.000 |
| IRIS_313-10997 | 0.000 | 0.000 | 0.000 | 0.104 | 0.000 | 0.000 | 0.896 |
| IRIS_313-10998 | 0.000 | 0.256 | 0.000 | 0.110 | 0.000 | 0.000 | 0.634 |
| IRIS_313-10999 | 0.000 | 1.000 | 0.000 | 0.000 | 0.000 | 0.000 | 0.000 |
| IRIS_313-11000 | 0.000 | 0.000 | 0.000 | 0.398 | 0.000 | 0.000 | 0.602 |
| IRIS_313-11001 | 0.000 | 1.000 | 0.000 | 0.000 | 0.000 | 0.000 | 0.000 |
| IRIS_313-11003 | 0.000 | 0.964 | 0.000 | 0.000 | 0.003 | 0.032 | 0.001 |
| IRIS_313-11004 | 0.000 | 1.000 | 0.000 | 0.000 | 0.000 | 0.000 | 0.000 |
| IRIS_313-11005 | 0.000 | 0.861 | 0.099 | 0.000 | 0.000 | 0.040 | 0.000 |

| | | | | | | | |
|---|---|---|---|---|---|---|---|
| IRIS_313-11006 | 0.000 | 0.005 | 0.000 | 0.642 | 0.000 | 0.000 | 0.353 |
| IRIS_313-11007 | 0.000 | 0.961 | 0.000 | 0.000 | 0.000 | 0.039 | 0.000 |
| IRIS_313-11008 | 0.000 | 0.965 | 0.013 | 0.010 | 0.000 | 0.011 | 0.001 |
| IRIS_313-11009 | 0.000 | 1.000 | 0.000 | 0.000 | 0.000 | 0.000 | 0.000 |
| IRIS_313-11010 | 0.000 | 0.000 | 0.000 | 0.510 | 0.000 | 0.000 | 0.490 |
| IRIS_313-11011 | 0.000 | 0.016 | 0.005 | 0.372 | 0.000 | 0.279 | 0.328 |
| IRIS_313-11012 | 0.000 | 0.529 | 0.000 | 0.000 | 0.000 | 0.000 | 0.471 |
| IRIS_313-11013 | 1.000 | 0.000 | 0.000 | 0.000 | 0.000 | 0.000 | 0.000 |
| IRIS_313-11014 | 1.000 | 0.000 | 0.000 | 0.000 | 0.000 | 0.000 | 0.000 |
| IRIS_313-11015 | 1.000 | 0.000 | 0.000 | 0.000 | 0.000 | 0.000 | 0.000 |
| IRIS_313-11016 | 1.000 | 0.000 | 0.000 | 0.000 | 0.000 | 0.000 | 0.000 |
| IRIS_313-11017 | 1.000 | 0.000 | 0.000 | 0.000 | 0.000 | 0.000 | 0.000 |
| IRIS_313-11018 | 1.000 | 0.000 | 0.000 | 0.000 | 0.000 | 0.000 | 0.000 |
| IRIS_313-11019 | 1.000 | 0.000 | 0.000 | 0.000 | 0.000 | 0.000 | 0.000 |
| IRIS_313-11020 | 1.000 | 0.000 | 0.000 | 0.000 | 0.000 | 0.000 | 0.000 |
| IRIS_313-11021 | 0.104 | 0.000 | 0.896 | 0.000 | 0.000 | 0.000 | 0.000 |
| IRIS_313-11022 | 0.126 | 0.000 | 0.874 | 0.000 | 0.000 | 0.000 | 0.000 |
| IRIS_313-11023 | 0.103 | 0.000 | 0.897 | 0.000 | 0.000 | 0.000 | 0.000 |
| IRIS_313-11024 | 1.000 | 0.000 | 0.000 | 0.000 | 0.000 | 0.000 | 0.000 |
| IRIS_313-11025 | 1.000 | 0.000 | 0.000 | 0.000 | 0.000 | 0.000 | 0.000 |
| IRIS_313-11026 | 0.000 | 0.000 | 1.000 | 0.000 | 0.000 | 0.000 | 0.000 |
| IRIS_313-11027 | 1.000 | 0.000 | 0.000 | 0.000 | 0.000 | 0.000 | 0.000 |
| IRIS_313-11028 | 1.000 | 0.000 | 0.000 | 0.000 | 0.000 | 0.000 | 0.000 |
| IRIS_313-11029 | 1.000 | 0.000 | 0.000 | 0.000 | 0.000 | 0.000 | 0.000 |
| IRIS_313-11030 | 0.278 | 0.000 | 0.722 | 0.000 | 0.000 | 0.000 | 0.000 |
| IRIS_313-11031 | 0.791 | 0.000 | 0.000 | 0.186 | 0.000 | 0.022 | 0.000 |
| IRIS_313-11032 | 0.274 | 0.000 | 0.646 | 0.000 | 0.000 | 0.052 | 0.028 |
| IRIS_313-11033 | 0.504 | 0.000 | 0.496 | 0.000 | 0.000 | 0.000 | 0.000 |
| IRIS_313-11034 | 1.000 | 0.000 | 0.000 | 0.000 | 0.000 | 0.000 | 0.000 |
| IRIS_313-11035 | 0.588 | 0.000 | 0.000 | 0.345 | 0.000 | 0.067 | 0.000 |
| IRIS_313-11036 | 0.715 | 0.000 | 0.285 | 0.000 | 0.000 | 0.000 | 0.000 |
| IRIS_313-11037 | 1.000 | 0.000 | 0.000 | 0.000 | 0.000 | 0.000 | 0.000 |
| IRIS_313-11038 | 0.000 | 0.000 | 0.000 | 0.000 | 0.000 | 1.000 | 0.000 |
| IRIS_313-11039 | 0.000 | 0.000 | 0.000 | 0.000 | 0.000 | 1.000 | 0.000 |
| IRIS_313-11040 | 0.133 | 0.000 | 0.000 | 0.867 | 0.000 | 0.000 | 0.000 |
| IRIS_313-11041 | 0.108 | 0.000 | 0.000 | 0.749 | 0.000 | 0.143 | 0.000 |
| IRIS_313-11042 | 0.171 | 0.000 | 0.000 | 0.773 | 0.000 | 0.056 | 0.000 |
| IRIS_313-11043 | 0.000 | 0.000 | 0.000 | 0.471 | 0.000 | 0.000 | 0.529 |
| IRIS_313-11044 | 0.010 | 0.849 | 0.049 | 0.046 | 0.045 | 0.000 | 0.000 |
| IRIS_313-11045 | 0.000 | 1.000 | 0.000 | 0.000 | 0.000 | 0.000 | 0.000 |
| IRIS_313-11046 | 0.014 | 0.919 | 0.000 | 0.067 | 0.000 | 0.000 | 0.000 |
| IRIS_313-11047 | 1.000 | 0.000 | 0.000 | 0.000 | 0.000 | 0.000 | 0.000 |
| IRIS_313-11048 | 1.000 | 0.000 | 0.000 | 0.000 | 0.000 | 0.000 | 0.000 |
| IRIS_313-11049 | 1.000 | 0.000 | 0.000 | 0.000 | 0.000 | 0.000 | 0.000 |
| IRIS_313-11050 | 1.000 | 0.000 | 0.000 | 0.000 | 0.000 | 0.000 | 0.000 |
| IRIS_313-11051 | 1.000 | 0.000 | 0.000 | 0.000 | 0.000 | 0.000 | 0.000 |
| IRIS_313-11052 | 0.939 | 0.025 | 0.002 | 0.002 | 0.000 | 0.029 | 0.004 |
| IRIS_313-11053 | 1.000 | 0.000 | 0.000 | 0.000 | 0.000 | 0.000 | 0.000 |
| IRIS_313-11054 | 1.000 | 0.000 | 0.000 | 0.000 | 0.000 | 0.000 | 0.000 |
| IRIS_313-11055 | 1.000 | 0.000 | 0.000 | 0.000 | 0.000 | 0.000 | 0.000 |
| IRIS_313-11056 | 1.000 | 0.000 | 0.000 | 0.000 | 0.000 | 0.000 | 0.000 |
| IRIS_313-11057 | 1.000 | 0.000 | 0.000 | 0.000 | 0.000 | 0.000 | 0.000 |
| IRIS_313-11058 | 1.000 | 0.000 | 0.000 | 0.000 | 0.000 | 0.000 | 0.000 |
| IRIS_313-11059 | 1.000 | 0.000 | 0.000 | 0.000 | 0.000 | 0.000 | 0.000 |
| IRIS_313-11060 | 1.000 | 0.000 | 0.000 | 0.000 | 0.000 | 0.000 | 0.000 |
| IRIS_313-11061 | 1.000 | 0.000 | 0.000 | 0.000 | 0.000 | 0.000 | 0.000 |
| IRIS_313-11062 | 0.137 | 0.224 | 0.640 | 0.000 | 0.000 | 0.000 | 0.000 |
| IRIS_313-11063 | 0.954 | 0.000 | 0.029 | 0.001 | 0.000 | 0.016 | 0.000 |
| IRIS_313-11064 | 0.977 | 0.000 | 0.023 | 0.000 | 0.000 | 0.000 | 0.000 |
| IRIS_313-11065 | 0.969 | 0.000 | 0.031 | 0.000 | 0.000 | 0.000 | 0.000 |
| IRIS_313-11066 | 0.201 | 0.255 | 0.543 | 0.000 | 0.000 | 0.000 | 0.000 |
| IRIS_313-11067 | 0.976 | 0.000 | 0.024 | 0.000 | 0.000 | 0.000 | 0.000 |
| IRIS_313-11068 | 0.066 | 0.207 | 0.727 | 0.000 | 0.000 | 0.000 | 0.000 |
| IRIS_313-11069 | 0.160 | 0.207 | 0.634 | 0.000 | 0.000 | 0.000 | 0.000 |
| IRIS_313-11070 | 0.176 | 0.270 | 0.553 | 0.000 | 0.001 | 0.000 | 0.000 |
| IRIS_313-11071 | 0.000 | 0.000 | 0.000 | 0.716 | 0.000 | 0.278 | 0.005 |
| IRIS_313-11072 | 0.000 | 0.000 | 0.000 | 1.000 | 0.000 | 0.000 | 0.000 |

| | | | | | | | |
|---|---|---|---|---|---|---|---|
| IRIS_313-11073 | 0.000 | 0.457 | 0.000 | 0.000 | 0.543 | 0.000 | 0.000 |
| IRIS_313-11074 | 0.000 | 0.000 | 0.000 | 0.949 | 0.000 | 0.000 | 0.051 |
| IRIS_313-11075 | 0.004 | 0.433 | 0.000 | 0.000 | 0.563 | 0.000 | 0.000 |
| IRIS_313-11076 | 0.000 | 0.000 | 0.000 | 0.926 | 0.000 | 0.074 | 0.000 |
| IRIS_313-11077 | 0.000 | 0.642 | 0.000 | 0.000 | 0.358 | 0.000 | 0.000 |
| IRIS_313-11078 | 0.000 | 0.000 | 0.000 | 0.136 | 0.000 | 0.112 | 0.752 |
| IRIS_313-11079 | 0.000 | 0.000 | 0.000 | 1.000 | 0.000 | 0.000 | 0.000 |
| IRIS_313-11080 | 0.000 | 0.023 | 0.000 | 0.977 | 0.000 | 0.000 | 0.000 |
| IRIS_313-11081 | 0.000 | 0.000 | 0.000 | 0.999 | 0.000 | 0.001 | 0.000 |
| IRIS_313-11082 | 0.000 | 0.000 | 0.000 | 1.000 | 0.000 | 0.000 | 0.000 |
| IRIS_313-11083 | 0.000 | 0.000 | 0.000 | 0.940 | 0.000 | 0.060 | 0.000 |
| IRIS_313-11084 | 0.000 | 0.000 | 0.000 | 0.999 | 0.000 | 0.000 | 0.001 |
| IRIS_313-11085 | 0.000 | 0.009 | 0.000 | 0.842 | 0.000 | 0.148 | 0.000 |
| IRIS_313-11086 | 0.000 | 0.000 | 0.000 | 0.928 | 0.000 | 0.072 | 0.000 |
| IRIS_313-11087 | 0.000 | 0.000 | 0.000 | 1.000 | 0.000 | 0.000 | 0.000 |
| IRIS_313-11088 | 0.000 | 0.000 | 0.000 | 1.000 | 0.000 | 0.000 | 0.000 |
| IRIS_313-11089 | 0.000 | 0.000 | 0.000 | 0.788 | 0.000 | 0.212 | 0.000 |
| IRIS_313-11090 | 0.000 | 0.000 | 0.000 | 0.973 | 0.000 | 0.027 | 0.000 |
| IRIS_313-11091 | 0.000 | 0.000 | 0.000 | 0.900 | 0.010 | 0.000 | 0.090 |
| IRIS_313-11092 | 0.000 | 0.351 | 0.000 | 0.422 | 0.162 | 0.010 | 0.055 |
| IRIS_313-11093 | 0.000 | 0.710 | 0.000 | 0.000 | 0.290 | 0.000 | 0.000 |
| IRIS_313-11094 | 0.015 | 0.693 | 0.000 | 0.000 | 0.260 | 0.032 | 0.000 |
| IRIS_313-11095 | 0.000 | 0.268 | 0.000 | 0.570 | 0.113 | 0.000 | 0.049 |
| IRIS_313-11096 | 0.000 | 0.000 | 0.000 | 0.998 | 0.000 | 0.002 | 0.000 |
| IRIS_313-11097 | 0.000 | 0.113 | 0.000 | 0.690 | 0.000 | 0.197 | 0.000 |
| IRIS_313-11098 | 0.000 | 0.034 | 0.000 | 0.418 | 0.000 | 0.000 | 0.548 |
| IRIS_313-11099 | 0.004 | 0.000 | 0.000 | 0.996 | 0.000 | 0.000 | 0.000 |
| IRIS_313-11100 | 0.000 | 1.000 | 0.000 | 0.000 | 0.000 | 0.000 | 0.000 |
| IRIS_313-11101 | 0.000 | 0.000 | 0.000 | 0.992 | 0.000 | 0.000 | 0.008 |
| IRIS_313-11102 | 0.000 | 1.000 | 0.000 | 0.000 | 0.000 | 0.000 | 0.000 |
| IRIS_313-11103 | 0.000 | 1.000 | 0.000 | 0.000 | 0.000 | 0.000 | 0.000 |
| IRIS_313-11104 | 0.000 | 1.000 | 0.000 | 0.000 | 0.000 | 0.000 | 0.000 |
| IRIS_313-11105 | 0.000 | 0.978 | 0.000 | 0.000 | 0.022 | 0.000 | 0.000 |
| IRIS_313-11106 | 0.000 | 1.000 | 0.000 | 0.000 | 0.000 | 0.000 | 0.000 |
| IRIS_313-11107 | 0.000 | 1.000 | 0.000 | 0.000 | 0.000 | 0.000 | 0.000 |
| IRIS_313-11108 | 0.000 | 0.013 | 0.000 | 0.407 | 0.000 | 0.000 | 0.580 |
| IRIS_313-11109 | 0.000 | 1.000 | 0.000 | 0.000 | 0.000 | 0.000 | 0.000 |
| IRIS_313-11110 | 0.189 | 0.019 | 0.016 | 0.536 | 0.000 | 0.231 | 0.009 |
| IRIS_313-11111 | 1.000 | 0.000 | 0.000 | 0.000 | 0.000 | 0.000 | 0.000 |
| IRIS_313-11112 | 0.854 | 0.000 | 0.009 | 0.128 | 0.000 | 0.000 | 0.009 |
| IRIS_313-11113 | 0.065 | 0.000 | 0.000 | 0.935 | 0.000 | 0.000 | 0.000 |
| IRIS_313-11114 | 0.009 | 0.000 | 0.000 | 0.991 | 0.000 | 0.000 | 0.000 |
| IRIS_313-11115 | 0.062 | 0.000 | 0.010 | 0.928 | 0.000 | 0.000 | 0.000 |
| IRIS_313-11116 | 0.864 | 0.000 | 0.000 | 0.136 | 0.000 | 0.000 | 0.000 |
| IRIS_313-11117 | 0.003 | 0.984 | 0.000 | 0.000 | 0.000 | 0.012 | 0.000 |
| IRIS_313-11118 | 0.000 | 0.000 | 0.000 | 1.000 | 0.000 | 0.000 | 0.000 |
| IRIS_313-11119 | 0.033 | 0.010 | 0.000 | 0.868 | 0.000 | 0.088 | 0.000 |
| IRIS_313-11120 | 0.039 | 0.000 | 0.000 | 0.961 | 0.000 | 0.000 | 0.000 |
| IRIS_313-11121 | 0.000 | 1.000 | 0.000 | 0.000 | 0.000 | 0.000 | 0.000 |
| IRIS_313-11122 | 0.027 | 0.000 | 0.000 | 0.030 | 0.000 | 0.186 | 0.757 |
| IRIS_313-11123 | 0.855 | 0.000 | 0.040 | 0.098 | 0.000 | 0.000 | 0.007 |
| IRIS_313-11124 | 0.654 | 0.000 | 0.088 | 0.227 | 0.032 | 0.000 | 0.000 |
| IRIS_313-11125 | 0.101 | 0.000 | 0.234 | 0.566 | 0.099 | 0.000 | 0.000 |
| IRIS_313-11126 | 0.000 | 0.000 | 0.000 | 1.000 | 0.000 | 0.000 | 0.000 |
| IRIS_313-11127 | 0.000 | 0.672 | 0.000 | 0.000 | 0.328 | 0.000 | 0.000 |
| IRIS_313-11128 | 0.000 | 0.003 | 0.000 | 0.997 | 0.001 | 0.000 | 0.000 |
| IRIS_313-11129 | 0.086 | 0.006 | 0.105 | 0.691 | 0.000 | 0.000 | 0.111 |
| IRIS_313-11130 | 0.000 | 0.006 | 0.016 | 0.903 | 0.000 | 0.000 | 0.074 |
| IRIS_313-11131 | 0.227 | 0.000 | 0.001 | 0.422 | 0.000 | 0.350 | 0.000 |
| IRIS_313-11132 | 0.000 | 0.000 | 0.000 | 0.972 | 0.000 | 0.028 | 0.000 |
| IRIS_313-11133 | 0.000 | 0.000 | 0.000 | 0.925 | 0.000 | 0.075 | 0.000 |
| IRIS_313-11134 | 0.033 | 0.000 | 0.023 | 0.606 | 0.000 | 0.339 | 0.000 |
| IRIS_313-11135 | 0.000 | 0.000 | 0.000 | 0.960 | 0.000 | 0.040 | 0.000 |
| IRIS_313-11136 | 0.000 | 0.150 | 0.392 | 0.458 | 0.000 | 0.000 | 0.000 |
| IRIS_313-11137 | 0.000 | 0.000 | 0.000 | 0.710 | 0.000 | 0.288 | 0.002 |
| IRIS_313-11138 | 0.006 | 0.000 | 0.000 | 0.928 | 0.000 | 0.000 | 0.067 |
| IRIS_313-11139 | 0.000 | 0.000 | 0.000 | 1.000 | 0.000 | 0.000 | 0.000 |

| | | | | | | | |
|---|---|---|---|---|---|---|---|
| IRIS_313-11140 | 0.028 | 0.000 | 0.000 | 0.972 | 0.000 | 0.000 | 0.000 |
| IRIS_313-11141 | 0.000 | 0.000 | 0.010 | 0.896 | 0.000 | 0.000 | 0.095 |
| IRIS_313-11142 | 0.000 | 0.001 | 0.001 | 0.997 | 0.001 | 0.000 | 0.000 |
| IRIS_313-11143 | 0.040 | 0.000 | 0.014 | 0.947 | 0.000 | 0.000 | 0.000 |
| IRIS_313-11144 | 0.027 | 0.000 | 0.000 | 0.959 | 0.000 | 0.000 | 0.013 |
| IRIS_313-11145 | 0.000 | 0.000 | 0.000 | 1.000 | 0.000 | 0.000 | 0.000 |
| IRIS_313-11146 | 0.000 | 0.000 | 0.000 | 0.959 | 0.000 | 0.000 | 0.041 |
| IRIS_313-11147 | 0.017 | 0.000 | 0.000 | 0.947 | 0.000 | 0.000 | 0.036 |
| IRIS_313-11148 | 0.017 | 0.000 | 0.000 | 0.967 | 0.000 | 0.000 | 0.017 |
| IRIS_313-11149 | 0.000 | 0.000 | 0.000 | 1.000 | 0.000 | 0.000 | 0.000 |
| IRIS_313-11150 | 0.000 | 0.000 | 0.000 | 1.000 | 0.000 | 0.000 | 0.000 |
| IRIS_313-11151 | 0.000 | 0.000 | 0.000 | 0.988 | 0.000 | 0.000 | 0.012 |
| IRIS_313-11152 | 0.000 | 0.000 | 0.000 | 0.000 | 0.000 | 1.000 | 0.000 |
| IRIS_313-11153 | 0.000 | 0.142 | 0.055 | 0.000 | 0.682 | 0.121 | 0.000 |
| IRIS_313-11154 | 1.000 | 0.000 | 0.000 | 0.000 | 0.000 | 0.000 | 0.000 |
| IRIS_313-11155 | 0.000 | 0.091 | 0.000 | 0.048 | 0.853 | 0.008 | 0.000 |
| IRIS_313-11156 | 0.000 | 0.522 | 0.000 | 0.000 | 0.478 | 0.000 | 0.000 |
| IRIS_313-11157 | 0.000 | 0.000 | 0.000 | 0.000 | 0.005 | 0.995 | 0.000 |
| IRIS_313-11158 | 0.000 | 0.000 | 0.000 | 0.000 | 0.000 | 1.000 | 0.000 |
| IRIS_313-11159 | 0.031 | 0.969 | 0.000 | 0.000 | 0.000 | 0.000 | 0.000 |
| IRIS_313-11160 | 0.002 | 0.000 | 0.000 | 0.503 | 0.016 | 0.006 | 0.473 |
| IRIS_313-11161 | 0.000 | 1.000 | 0.000 | 0.000 | 0.000 | 0.000 | 0.000 |
| IRIS_313-11162 | 0.000 | 0.000 | 0.000 | 0.000 | 0.000 | 0.278 | 0.722 |
| IRIS_313-11163 | 1.000 | 0.000 | 0.000 | 0.000 | 0.000 | 0.000 | 0.000 |
| IRIS_313-11164 | 1.000 | 0.000 | 0.000 | 0.000 | 0.000 | 0.000 | 0.000 |
| IRIS_313-11165 | 0.617 | 0.000 | 0.000 | 0.383 | 0.000 | 0.000 | 0.000 |
| IRIS_313-11166 | 0.964 | 0.000 | 0.000 | 0.028 | 0.000 | 0.000 | 0.008 |
| IRIS_313-11167 | 0.071 | 0.000 | 0.000 | 0.929 | 0.000 | 0.000 | 0.000 |
| IRIS_313-11168 | 1.000 | 0.000 | 0.000 | 0.000 | 0.000 | 0.000 | 0.000 |
| IRIS_313-11169 | 1.000 | 0.000 | 0.000 | 0.000 | 0.000 | 0.000 | 0.000 |
| IRIS_313-11170 | 1.000 | 0.000 | 0.000 | 0.000 | 0.000 | 0.000 | 0.000 |
| IRIS_313-11171 | 1.000 | 0.000 | 0.000 | 0.000 | 0.000 | 0.000 | 0.000 |
| IRIS_313-11172 | 1.000 | 0.000 | 0.000 | 0.000 | 0.000 | 0.000 | 0.000 |
| IRIS_313-11173 | 1.000 | 0.000 | 0.000 | 0.000 | 0.000 | 0.000 | 0.000 |
| IRIS_313-11174 | 1.000 | 0.000 | 0.000 | 0.000 | 0.000 | 0.000 | 0.000 |
| IRIS_313-11175 | 1.000 | 0.000 | 0.000 | 0.000 | 0.000 | 0.000 | 0.000 |
| IRIS_313-11176 | 0.357 | 0.000 | 0.000 | 0.254 | 0.000 | 0.000 | 0.389 |
| IRIS_313-11177 | 0.000 | 0.000 | 0.000 | 0.379 | 0.000 | 0.000 | 0.621 |
| IRIS_313-11178 | 0.000 | 0.038 | 0.000 | 0.474 | 0.000 | 0.000 | 0.489 |
| IRIS_313-11179 | 0.000 | 0.000 | 0.000 | 0.546 | 0.000 | 0.000 | 0.454 |
| IRIS_313-11180 | 0.000 | 0.000 | 0.000 | 0.513 | 0.065 | 0.000 | 0.421 |
| IRIS_313-11181 | 0.042 | 0.000 | 0.000 | 0.590 | 0.000 | 0.000 | 0.368 |
| IRIS_313-11182 | 0.000 | 0.000 | 0.000 | 0.384 | 0.000 | 0.000 | 0.616 |
| IRIS_313-11183 | 0.000 | 1.000 | 0.000 | 0.000 | 0.000 | 0.000 | 0.000 |
| IRIS_313-11189 | 0.415 | 0.000 | 0.585 | 0.000 | 0.000 | 0.000 | 0.000 |
| IRIS_313-11191 | 0.881 | 0.040 | 0.074 | 0.000 | 0.005 | 0.000 | 0.000 |
| IRIS_313-11192 | 0.000 | 0.000 | 0.000 | 0.575 | 0.000 | 0.000 | 0.425 |
| IRIS_313-11193 | 0.000 | 0.000 | 0.000 | 0.828 | 0.000 | 0.172 | 0.000 |
| IRIS_313-11194 | 0.000 | 0.000 | 0.000 | 0.998 | 0.000 | 0.000 | 0.002 |
| IRIS_313-11195 | 0.000 | 0.650 | 0.000 | 0.000 | 0.350 | 0.000 | 0.000 |
| IRIS_313-11196 | 0.000 | 0.018 | 0.000 | 0.914 | 0.000 | 0.000 | 0.068 |
| IRIS_313-11197 | 0.000 | 0.000 | 0.000 | 0.101 | 0.000 | 0.670 | 0.228 |
| IRIS_313-11198 | 0.000 | 0.000 | 0.000 | 0.000 | 1.000 | 0.000 | 0.000 |
| IRIS_313-11199 | 0.052 | 0.000 | 0.000 | 0.654 | 0.000 | 0.294 | 0.000 |
| IRIS_313-11200 | 0.000 | 0.000 | 0.000 | 0.172 | 0.000 | 0.000 | 0.828 |
| IRIS_313-11201 | 0.000 | 0.000 | 0.000 | 0.000 | 1.000 | 0.000 | 0.000 |
| IRIS_313-11202 | 0.000 | 0.000 | 0.000 | 0.000 | 1.000 | 0.000 | 0.000 |
| IRIS_313-11203 | 0.000 | 0.000 | 0.000 | 0.982 | 0.000 | 0.018 | 0.000 |
| IRIS_313-11204 | 0.268 | 0.000 | 0.000 | 0.728 | 0.000 | 0.004 | 0.000 |
| IRIS_313-11205 | 0.033 | 0.000 | 0.000 | 0.911 | 0.000 | 0.057 | 0.000 |
| IRIS_313-11206 | 0.019 | 0.000 | 0.015 | 0.639 | 0.000 | 0.000 | 0.327 |
| IRIS_313-11207 | 0.051 | 0.000 | 0.000 | 0.916 | 0.007 | 0.000 | 0.026 |
| IRIS_313-11208 | 0.056 | 0.000 | 0.010 | 0.933 | 0.001 | 0.000 | 0.000 |
| IRIS_313-11209 | 0.052 | 0.000 | 0.003 | 0.941 | 0.003 | 0.000 | 0.000 |
| IRIS_313-11210 | 1.000 | 0.000 | 0.000 | 0.000 | 0.000 | 0.000 | 0.000 |
| IRIS_313-11211 | 0.127 | 0.000 | 0.000 | 0.873 | 0.000 | 0.000 | 0.000 |
| IRIS_313-11212 | 0.096 | 0.000 | 0.000 | 0.904 | 0.000 | 0.000 | 0.000 |

| | | | | | | | |
|---|---|---|---|---|---|---|---|
| IRIS_313-11213 | 1.000 | 0.000 | 0.000 | 0.000 | 0.000 | 0.000 | 0.000 |
| IRIS_313-11214 | 0.891 | 0.000 | 0.004 | 0.074 | 0.000 | 0.028 | 0.004 |
| IRIS_313-11216 | 0.873 | 0.000 | 0.025 | 0.101 | 0.000 | 0.000 | 0.000 |
| IRIS_313-11217 | 0.000 | 0.165 | 0.588 | 0.000 | 0.120 | 0.127 | 0.000 |
| IRIS_313-11218 | 0.000 | 0.000 | 1.000 | 0.000 | 0.000 | 0.000 | 0.000 |
| IRIS_313-11219 | 0.118 | 0.001 | 0.013 | 0.808 | 0.001 | 0.058 | 0.000 |
| IRIS_313-11220 | 0.093 | 0.000 | 0.000 | 0.907 | 0.000 | 0.000 | 0.000 |
| IRIS_313-11221 | 0.000 | 0.000 | 0.000 | 1.000 | 0.000 | 0.000 | 0.000 |
| IRIS_313-11222 | 0.101 | 0.000 | 0.001 | 0.863 | 0.002 | 0.034 | 0.000 |
| IRIS_313-11223 | 0.000 | 0.000 | 0.648 | 0.352 | 0.000 | 0.000 | 0.000 |
| IRIS_313-11224 | 0.050 | 0.000 | 0.001 | 0.949 | 0.000 | 0.000 | 0.000 |
| IRIS_313-11225 | 0.052 | 0.000 | 0.000 | 0.947 | 0.000 | 0.000 | 0.001 |
| IRIS_313-11226 | 0.000 | 0.000 | 0.000 | 0.947 | 0.000 | 0.053 | 0.000 |
| IRIS_313-11227 | 0.095 | 0.000 | 0.000 | 0.904 | 0.002 | 0.000 | 0.000 |
| IRIS_313-11228 | 0.037 | 0.000 | 0.000 | 0.905 | 0.000 | 0.058 | 0.000 |
| IRIS_313-11229 | 0.040 | 0.000 | 0.000 | 0.960 | 0.000 | 0.000 | 0.000 |
| IRIS_313-11230 | 0.091 | 0.000 | 0.004 | 0.905 | 0.000 | 0.000 | 0.000 |
| IRIS_313-11231 | 0.020 | 0.000 | 0.015 | 0.964 | 0.002 | 0.000 | 0.000 |
| IRIS_313-11232 | 0.918 | 0.024 | 0.000 | 0.000 | 0.002 | 0.000 | 0.057 |
| IRIS_313-11233 | 0.000 | 0.000 | 0.000 | 0.026 | 0.000 | 0.153 | 0.821 |
| IRIS_313-11234 | 0.000 | 0.000 | 0.030 | 0.000 | 0.000 | 0.132 | 0.838 |
| IRIS_313-11235 | 0.024 | 0.000 | 0.000 | 0.971 | 0.006 | 0.000 | 0.000 |
| IRIS_313-11236 | 0.000 | 0.000 | 0.000 | 0.997 | 0.003 | 0.000 | 0.000 |
| IRIS_313-11237 | 0.000 | 0.000 | 0.000 | 0.000 | 0.046 | 0.225 | 0.729 |
| IRIS_313-11238 | 0.007 | 0.993 | 0.000 | 0.000 | 0.000 | 0.000 | 0.000 |
| IRIS_313-11239 | 0.000 | 0.000 | 0.000 | 0.227 | 0.000 | 0.000 | 0.773 |
| IRIS_313-11240 | 0.039 | 0.000 | 0.000 | 0.446 | 0.000 | 0.360 | 0.155 |
| IRIS_313-11241 | 0.082 | 0.000 | 0.006 | 0.790 | 0.000 | 0.122 | 0.000 |
| IRIS_313-11242 | 0.098 | 0.000 | 0.001 | 0.901 | 0.000 | 0.000 | 0.000 |
| IRIS_313-11243 | 0.548 | 0.000 | 0.000 | 0.000 | 0.000 | 0.452 | 0.000 |
| IRIS_313-11244 | 0.000 | 0.000 | 0.000 | 0.495 | 0.000 | 0.505 | 0.000 |
| IRIS_313-11245 | 0.000 | 0.000 | 0.000 | 0.041 | 0.000 | 0.220 | 0.740 |
| IRIS_313-11246 | 0.000 | 0.072 | 0.000 | 0.050 | 0.870 | 0.000 | 0.008 |
| IRIS_313-11247 | 0.021 | 0.000 | 0.000 | 0.267 | 0.000 | 0.712 | 0.000 |
| IRIS_313-11248 | 0.000 | 1.000 | 0.000 | 0.000 | 0.000 | 0.000 | 0.000 |
| IRIS_313-11249 | 0.007 | 0.000 | 0.000 | 0.505 | 0.000 | 0.083 | 0.405 |
| IRIS_313-11250 | 0.000 | 0.000 | 0.000 | 0.000 | 0.000 | 0.190 | 0.810 |
| IRIS_313-11251 | 0.000 | 0.000 | 0.000 | 0.000 | 0.000 | 0.169 | 0.831 |
| IRIS_313-11252 | 0.113 | 0.000 | 0.000 | 0.749 | 0.007 | 0.000 | 0.131 |
| IRIS_313-11253 | 0.021 | 0.135 | 0.000 | 0.000 | 0.000 | 0.221 | 0.623 |
| IRIS_313-11254 | 0.000 | 0.000 | 0.000 | 0.956 | 0.005 | 0.039 | 0.000 |
| IRIS_313-11255 | 0.061 | 0.000 | 0.000 | 0.480 | 0.000 | 0.459 | 0.000 |
| IRIS_313-11256 | 0.044 | 0.000 | 0.000 | 0.956 | 0.000 | 0.000 | 0.000 |
| IRIS_313-11257 | 0.019 | 0.217 | 0.000 | 0.758 | 0.006 | 0.000 | 0.000 |
| IRIS_313-11258 | 0.000 | 0.000 | 0.907 | 0.000 | 0.000 | 0.000 | 0.093 |
| IRIS_313-11259 | 0.000 | 0.088 | 0.912 | 0.000 | 0.000 | 0.000 | 0.000 |
| IRIS_313-11260 | 0.118 | 0.040 | 0.018 | 0.808 | 0.016 | 0.000 | 0.000 |
| IRIS_313-11261 | 0.012 | 0.000 | 0.000 | 0.987 | 0.001 | 0.000 | 0.000 |
| IRIS_313-11262 | 0.007 | 0.000 | 0.005 | 0.754 | 0.000 | 0.125 | 0.108 |
| IRIS_313-11263 | 0.000 | 0.000 | 0.095 | 0.378 | 0.087 | 0.000 | 0.440 |
| IRIS_313-11264 | 0.000 | 0.000 | 0.000 | 0.805 | 0.000 | 0.195 | 0.000 |
| IRIS_313-11265 | 1.000 | 0.000 | 0.000 | 0.000 | 0.000 | 0.000 | 0.000 |
| IRIS_313-11266 | 0.000 | 0.000 | 0.000 | 1.000 | 0.000 | 0.000 | 0.000 |
| IRIS_313-11267 | 0.018 | 0.000 | 0.000 | 0.982 | 0.000 | 0.000 | 0.000 |
| IRIS_313-11268 | 0.000 | 0.037 | 0.963 | 0.000 | 0.000 | 0.000 | 0.000 |
| IRIS_313-11269 | 0.000 | 0.000 | 0.458 | 0.526 | 0.016 | 0.000 | 0.000 |
| IRIS_313-11270 | 0.000 | 0.000 | 0.797 | 0.203 | 0.000 | 0.000 | 0.000 |
| IRIS_313-11271 | 0.032 | 0.000 | 0.018 | 0.949 | 0.000 | 0.000 | 0.000 |
| IRIS_313-11272 | 1.000 | 0.000 | 0.000 | 0.000 | 0.000 | 0.000 | 0.000 |
| IRIS_313-11273 | 0.041 | 0.000 | 0.000 | 0.959 | 0.000 | 0.000 | 0.000 |
| IRIS_313-11274 | 0.810 | 0.000 | 0.000 | 0.125 | 0.000 | 0.000 | 0.065 |
| IRIS_313-11275 | 0.107 | 0.000 | 0.000 | 0.591 | 0.000 | 0.302 | 0.000 |
| IRIS_313-11276 | 0.000 | 0.006 | 0.105 | 0.737 | 0.026 | 0.126 | 0.000 |
| IRIS_313-11277 | 0.858 | 0.000 | 0.015 | 0.128 | 0.000 | 0.000 | 0.000 |
| IRIS_313-11278 | 0.022 | 0.000 | 0.000 | 0.709 | 0.000 | 0.269 | 0.000 |
| IRIS_313-11279 | 0.000 | 0.040 | 0.043 | 0.725 | 0.042 | 0.150 | 0.000 |
| IRIS_313-11280 | 0.000 | 0.026 | 0.000 | 0.118 | 0.000 | 0.020 | 0.836 |

| | | | | | | | |
|---|---|---|---|---|---|---|---|
| IRIS_313-11281 | 0.000 | 0.000 | 0.000 | 0.000 | 0.000 | 1.000 | 0.000 |
| IRIS_313-11282 | 0.048 | 0.000 | 0.000 | 0.941 | 0.000 | 0.000 | 0.011 |
| IRIS_313-11283 | 0.022 | 0.000 | 0.000 | 0.520 | 0.019 | 0.440 | 0.000 |
| IRIS_313-11284 | 0.019 | 0.000 | 0.007 | 0.884 | 0.001 | 0.000 | 0.089 |
| IRIS_313-11285 | 0.097 | 0.000 | 0.082 | 0.821 | 0.000 | 0.000 | 0.000 |
| IRIS_313-11286 | 0.077 | 0.049 | 0.024 | 0.428 | 0.010 | 0.412 | 0.000 |
| IRIS_313-11287 | 0.000 | 0.000 | 0.000 | 1.000 | 0.000 | 0.000 | 0.000 |
| IRIS_313-11288 | 0.093 | 0.000 | 0.000 | 0.549 | 0.053 | 0.112 | 0.194 |
| IRIS_313-11289 | 0.000 | 0.057 | 0.943 | 0.000 | 0.000 | 0.000 | 0.000 |
| IRIS_313-11290 | 0.000 | 0.048 | 0.411 | 0.161 | 0.000 | 0.134 | 0.246 |
| IRIS_313-11291 | 1.000 | 0.000 | 0.000 | 0.000 | 0.000 | 0.000 | 0.000 |
| IRIS_313-11292 | 0.028 | 0.000 | 0.000 | 0.943 | 0.000 | 0.030 | 0.000 |
| IRIS_313-11293 | 0.000 | 0.037 | 0.963 | 0.000 | 0.000 | 0.000 | 0.000 |
| IRIS_313-11294 | 0.017 | 0.000 | 0.000 | 0.977 | 0.000 | 0.006 | 0.000 |
| IRIS_313-11295 | 0.175 | 0.148 | 0.039 | 0.278 | 0.060 | 0.300 | 0.000 |
| IRIS_313-11296 | 0.055 | 0.000 | 0.000 | 0.670 | 0.000 | 0.275 | 0.000 |
| IRIS_313-11297 | 0.000 | 0.328 | 0.456 | 0.000 | 0.216 | 0.000 | 0.000 |
| IRIS_313-11298 | 1.000 | 0.000 | 0.000 | 0.000 | 0.000 | 0.000 | 0.000 |
| IRIS_313-11299 | 0.034 | 0.000 | 0.000 | 0.734 | 0.026 | 0.206 | 0.000 |
| IRIS_313-11300 | 0.132 | 0.000 | 0.013 | 0.855 | 0.000 | 0.000 | 0.000 |
| IRIS_313-11301 | 0.045 | 0.000 | 0.000 | 0.955 | 0.000 | 0.000 | 0.000 |
| IRIS_313-11302 | 0.263 | 0.000 | 0.000 | 0.578 | 0.013 | 0.145 | 0.000 |
| IRIS_313-11303 | 0.073 | 0.000 | 0.000 | 0.687 | 0.013 | 0.228 | 0.000 |
| IRIS_313-11304 | 0.000 | 0.000 | 0.000 | 0.883 | 0.000 | 0.117 | 0.000 |
| IRIS_313-11305 | 0.014 | 0.110 | 0.015 | 0.790 | 0.000 | 0.070 | 0.000 |
| IRIS_313-11306 | 0.012 | 0.000 | 0.000 | 0.643 | 0.000 | 0.346 | 0.000 |
| IRIS_313-11307 | 0.000 | 0.000 | 0.000 | 1.000 | 0.000 | 0.000 | 0.000 |
| IRIS_313-11308 | 0.000 | 0.000 | 0.000 | 1.000 | 0.000 | 0.000 | 0.000 |
| IRIS_313-11309 | 0.000 | 0.000 | 0.000 | 0.705 | 0.000 | 0.295 | 0.000 |
| IRIS_313-11310 | 0.045 | 0.000 | 0.000 | 0.955 | 0.000 | 0.000 | 0.000 |
| IRIS_313-11311 | 0.000 | 0.889 | 0.000 | 0.000 | 0.111 | 0.000 | 0.000 |
| IRIS_313-11312 | 0.000 | 0.000 | 0.000 | 0.490 | 0.031 | 0.000 | 0.479 |
| IRIS_313-11313 | 0.000 | 0.947 | 0.000 | 0.000 | 0.000 | 0.053 | 0.000 |
| IRIS_313-11314 | 0.000 | 0.999 | 0.000 | 0.000 | 0.001 | 0.000 | 0.000 |
| IRIS_313-11315 | 0.000 | 0.000 | 0.000 | 0.550 | 0.000 | 0.000 | 0.450 |
| IRIS_313-11316 | 0.000 | 0.000 | 0.000 | 0.516 | 0.027 | 0.000 | 0.457 |
| IRIS_313-11317 | 0.000 | 0.000 | 0.000 | 0.291 | 0.000 | 0.000 | 0.709 |
| IRIS_313-11318 | 0.000 | 0.000 | 0.001 | 0.493 | 0.000 | 0.000 | 0.506 |
| IRIS_313-11319 | 0.000 | 0.000 | 0.000 | 0.257 | 0.000 | 0.000 | 0.743 |
| IRIS_313-11320 | 0.000 | 0.000 | 0.000 | 0.395 | 0.000 | 0.000 | 0.605 |
| IRIS_313-11321 | 0.053 | 0.000 | 0.000 | 0.917 | 0.000 | 0.030 | 0.000 |
| IRIS_313-11322 | 1.000 | 0.000 | 0.000 | 0.000 | 0.000 | 0.000 | 0.000 |
| IRIS_313-11323 | 0.993 | 0.000 | 0.000 | 0.000 | 0.007 | 0.000 | 0.000 |
| IRIS_313-11324 | 1.000 | 0.000 | 0.000 | 0.000 | 0.000 | 0.000 | 0.000 |
| IRIS_313-11325 | 0.056 | 0.000 | 0.000 | 0.469 | 0.000 | 0.474 | 0.000 |
| IRIS_313-11326 | 0.000 | 0.000 | 0.000 | 0.230 | 0.000 | 0.000 | 0.770 |
| IRIS_313-11327 | 0.000 | 0.000 | 0.000 | 1.000 | 0.000 | 0.000 | 0.000 |
| IRIS_313-11328 | 0.000 | 1.000 | 0.000 | 0.000 | 0.000 | 0.000 | 0.000 |
| IRIS_313-11329 | 0.000 | 1.000 | 0.000 | 0.000 | 0.000 | 0.000 | 0.000 |
| IRIS_313-11330 | 0.000 | 0.005 | 0.000 | 0.944 | 0.014 | 0.000 | 0.037 |
| IRIS_313-11331 | 0.000 | 0.000 | 0.000 | 0.838 | 0.000 | 0.134 | 0.028 |
| IRIS_313-11334 | 0.000 | 0.000 | 0.000 | 0.838 | 0.000 | 0.064 | 0.097 |
| IRIS_313-11335 | 0.099 | 0.009 | 0.000 | 0.687 | 0.000 | 0.179 | 0.026 |
| IRIS_313-11336 | 0.000 | 0.000 | 0.063 | 0.000 | 0.937 | 0.000 | 0.000 |
| IRIS_313-11337 | 0.000 | 1.000 | 0.000 | 0.000 | 0.000 | 0.000 | 0.000 |
| IRIS_313-11338 | 0.000 | 0.008 | 0.000 | 0.767 | 0.001 | 0.224 | 0.000 |
| IRIS_313-11339 | 0.069 | 0.000 | 0.000 | 0.931 | 0.000 | 0.000 | 0.000 |
| IRIS_313-11340 | 0.000 | 1.000 | 0.000 | 0.000 | 0.000 | 0.000 | 0.000 |
| IRIS_313-11341 | 0.000 | 0.000 | 0.000 | 0.997 | 0.000 | 0.000 | 0.003 |
| IRIS_313-11342 | 0.000 | 0.212 | 0.000 | 0.197 | 0.000 | 0.100 | 0.492 |
| IRIS_313-11343 | 0.000 | 0.016 | 0.000 | 0.978 | 0.000 | 0.000 | 0.006 |
| IRIS_313-11344 | 0.040 | 0.019 | 0.000 | 0.873 | 0.000 | 0.000 | 0.068 |
| IRIS_313-11345 | 0.000 | 0.017 | 0.000 | 0.704 | 0.000 | 0.280 | 0.000 |
| IRIS_313-11346 | 0.000 | 0.000 | 0.000 | 0.104 | 0.000 | 0.000 | 0.896 |
| IRIS_313-11347 | 0.000 | 0.000 | 0.000 | 0.700 | 0.010 | 0.291 | 0.000 |
| IRIS_313-11348 | 1.000 | 0.000 | 0.000 | 0.000 | 0.000 | 0.000 | 0.000 |
| IRIS_313-11349 | 0.023 | 0.000 | 0.000 | 0.977 | 0.000 | 0.000 | 0.000 |

| | | | | | | | |
|---|---|---|---|---|---|---|---|
| IRIS_313-11350 | 0.000 | 0.000 | 1.000 | 0.000 | 0.000 | 0.000 | 0.000 |
| IRIS_313-11351 | 0.023 | 0.000 | 0.000 | 0.891 | 0.000 | 0.086 | 0.000 |
| IRIS_313-11352 | 0.000 | 0.063 | 0.781 | 0.000 | 0.038 | 0.000 | 0.118 |
| IRIS_313-11353 | 0.894 | 0.000 | 0.000 | 0.106 | 0.000 | 0.000 | 0.000 |
| IRIS_313-11354 | 0.054 | 0.000 | 0.000 | 0.908 | 0.000 | 0.000 | 0.038 |
| IRIS_313-11355 | 0.358 | 0.000 | 0.000 | 0.000 | 0.000 | 0.642 | 0.000 |
| IRIS_313-11356 | 0.000 | 0.000 | 1.000 | 0.000 | 0.000 | 0.000 | 0.000 |
| IRIS_313-11357 | 0.013 | 0.000 | 0.000 | 0.000 | 0.000 | 0.352 | 0.634 |
| IRIS_313-11358 | 0.028 | 0.000 | 0.000 | 0.972 | 0.000 | 0.000 | 0.000 |
| IRIS_313-11359 | 0.028 | 0.000 | 0.000 | 0.972 | 0.000 | 0.000 | 0.000 |
| IRIS_313-11360 | 0.000 | 0.111 | 0.592 | 0.000 | 0.138 | 0.020 | 0.138 |
| IRIS_313-11361 | 0.056 | 0.000 | 0.000 | 0.944 | 0.000 | 0.000 | 0.000 |
| IRIS_313-11362 | 0.000 | 0.000 | 1.000 | 0.000 | 0.000 | 0.000 | 0.000 |
| IRIS_313-11363 | 0.032 | 0.000 | 0.000 | 0.966 | 0.000 | 0.002 | 0.000 |
| IRIS_313-11364 | 0.000 | 0.000 | 0.350 | 0.650 | 0.000 | 0.000 | 0.000 |
| IRIS_313-11365 | 0.040 | 0.000 | 0.000 | 0.960 | 0.000 | 0.000 | 0.000 |
| IRIS_313-11367 | 0.043 | 0.000 | 0.000 | 0.956 | 0.000 | 0.001 | 0.000 |
| IRIS_313-11368 | 0.017 | 0.000 | 0.002 | 0.981 | 0.001 | 0.000 | 0.000 |
| IRIS_313-11369 | 0.047 | 0.000 | 0.000 | 0.953 | 0.000 | 0.000 | 0.000 |
| IRIS_313-11370 | 0.037 | 0.000 | 0.000 | 0.963 | 0.000 | 0.000 | 0.000 |
| IRIS_313-11371 | 1.000 | 0.000 | 0.000 | 0.000 | 0.000 | 0.000 | 0.000 |
| IRIS_313-11372 | 0.060 | 0.000 | 0.000 | 0.940 | 0.000 | 0.000 | 0.000 |
| IRIS_313-11373 | 0.000 | 0.000 | 1.000 | 0.000 | 0.000 | 0.000 | 0.000 |
| IRIS_313-11374 | 1.000 | 0.000 | 0.000 | 0.000 | 0.000 | 0.000 | 0.000 |
| IRIS_313-11375 | 0.000 | 0.000 | 0.722 | 0.278 | 0.000 | 0.000 | 0.000 |
| IRIS_313-11376 | 0.000 | 1.000 | 0.000 | 0.000 | 0.000 | 0.000 | 0.000 |
| IRIS_313-11377 | 0.005 | 0.030 | 0.000 | 0.642 | 0.006 | 0.310 | 0.008 |
| IRIS_313-11379 | 0.017 | 0.983 | 0.000 | 0.000 | 0.000 | 0.000 | 0.000 |
| IRIS_313-11380 | 0.000 | 0.732 | 0.000 | 0.268 | 0.000 | 0.000 | 0.000 |
| IRIS_313-11381 | 0.000 | 0.000 | 0.000 | 0.748 | 0.000 | 0.251 | 0.001 |
| IRIS_313-11382 | 0.082 | 0.006 | 0.000 | 0.785 | 0.000 | 0.000 | 0.126 |
| IRIS_313-11383 | 0.026 | 0.023 | 0.004 | 0.531 | 0.019 | 0.398 | 0.000 |
| IRIS_313-11384 | 0.037 | 0.000 | 0.000 | 0.459 | 0.050 | 0.455 | 0.000 |
| IRIS_313-11385 | 0.019 | 0.000 | 0.000 | 0.000 | 0.000 | 0.363 | 0.618 |
| IRIS_313-11386 | 0.000 | 0.002 | 0.000 | 0.995 | 0.000 | 0.002 | 0.001 |
| IRIS_313-11387 | 0.064 | 0.000 | 0.000 | 0.820 | 0.012 | 0.061 | 0.044 |
| IRIS_313-11388 | 0.000 | 0.000 | 0.000 | 1.000 | 0.000 | 0.000 | 0.000 |
| IRIS_313-11389 | 0.000 | 0.000 | 0.000 | 1.000 | 0.000 | 0.000 | 0.000 |
| IRIS_313-11390 | 0.000 | 0.000 | 0.000 | 0.998 | 0.000 | 0.002 | 0.000 |
| IRIS_313-11391 | 0.000 | 0.000 | 0.000 | 0.975 | 0.000 | 0.025 | 0.000 |
| IRIS_313-11392 | 0.000 | 0.000 | 0.000 | 0.994 | 0.000 | 0.000 | 0.006 |
| IRIS_313-11393 | 0.000 | 0.000 | 0.000 | 1.000 | 0.000 | 0.000 | 0.000 |
| IRIS_313-11394 | 0.000 | 0.000 | 0.012 | 0.470 | 0.000 | 0.049 | 0.469 |
| IRIS_313-11395 | 0.000 | 0.025 | 0.000 | 0.502 | 0.000 | 0.000 | 0.473 |
| IRIS_313-11396 | 0.000 | 1.000 | 0.000 | 0.000 | 0.000 | 0.000 | 0.000 |
| IRIS_313-11397 | 0.000 | 1.000 | 0.000 | 0.000 | 0.000 | 0.000 | 0.000 |
| IRIS_313-11398 | 0.000 | 0.000 | 0.000 | 0.349 | 0.000 | 0.000 | 0.651 |
| IRIS_313-11399 | 0.035 | 0.000 | 0.000 | 0.854 | 0.000 | 0.057 | 0.055 |
| IRIS_313-11400 | 0.045 | 0.000 | 0.025 | 0.925 | 0.004 | 0.000 | 0.001 |
| IRIS_313-11401 | 0.000 | 0.085 | 0.915 | 0.000 | 0.000 | 0.000 | 0.000 |
| IRIS_313-11402 | 0.000 | 0.000 | 0.000 | 1.000 | 0.000 | 0.000 | 0.000 |
| IRIS_313-11403 | 0.054 | 0.000 | 0.000 | 0.723 | 0.030 | 0.193 | 0.000 |
| IRIS_313-11404 | 0.000 | 0.000 | 0.000 | 0.957 | 0.000 | 0.043 | 0.000 |
| IRIS_313-11405 | 0.038 | 0.000 | 0.000 | 0.662 | 0.000 | 0.000 | 0.301 |
| IRIS_313-11406 | 0.000 | 0.000 | 0.000 | 1.000 | 0.000 | 0.000 | 0.000 |
| IRIS_313-11407 | 0.010 | 0.000 | 0.069 | 0.921 | 0.000 | 0.000 | 0.000 |
| IRIS_313-11408 | 0.043 | 0.000 | 0.000 | 0.957 | 0.000 | 0.000 | 0.000 |
| IRIS_313-11409 | 0.161 | 0.000 | 0.000 | 0.771 | 0.000 | 0.068 | 0.000 |
| IRIS_313-11410 | 0.123 | 0.000 | 0.000 | 0.743 | 0.000 | 0.000 | 0.134 |
| IRIS_313-11411 | 0.104 | 0.000 | 0.005 | 0.708 | 0.002 | 0.181 | 0.000 |
| IRIS_313-11413 | 0.735 | 0.000 | 0.043 | 0.000 | 0.223 | 0.000 | 0.000 |
| IRIS_313-11414 | 0.110 | 0.000 | 0.000 | 0.627 | 0.000 | 0.179 | 0.084 |
| IRIS_313-11415 | 0.195 | 0.000 | 0.062 | 0.394 | 0.140 | 0.209 | 0.000 |
| IRIS_313-11416 | 0.029 | 0.000 | 0.007 | 0.965 | 0.000 | 0.000 | 0.000 |
| IRIS_313-11417 | 0.118 | 0.000 | 0.481 | 0.401 | 0.000 | 0.000 | 0.000 |
| IRIS_313-11418 | 0.000 | 0.000 | 0.000 | 1.000 | 0.000 | 0.000 | 0.000 |
| IRIS_313-11419 | 0.129 | 0.001 | 0.018 | 0.853 | 0.000 | 0.000 | 0.000 |

| | | | | | | | |
|---|---|---|---|---|---|---|---|
| IRIS_313-11420 | 0.172 | 0.000 | 0.000 | 0.825 | 0.003 | 0.000 | 0.000 |
| IRIS_313-11421 | 0.073 | 0.000 | 0.033 | 0.894 | 0.000 | 0.000 | 0.000 |
| IRIS_313-11422 | 0.000 | 0.000 | 0.000 | 0.018 | 0.007 | 0.975 | 0.000 |
| IRIS_313-11423 | 0.000 | 0.016 | 0.000 | 0.041 | 0.000 | 0.000 | 0.942 |
| IRIS_313-11424 | 0.000 | 1.000 | 0.000 | 0.000 | 0.000 | 0.000 | 0.000 |
| IRIS_313-11425 | 0.000 | 0.542 | 0.000 | 0.000 | 0.000 | 0.018 | 0.440 |
| IRIS_313-11426 | 0.000 | 1.000 | 0.000 | 0.000 | 0.000 | 0.000 | 0.000 |
| IRIS_313-11427 | 0.000 | 1.000 | 0.000 | 0.000 | 0.000 | 0.000 | 0.000 |
| IRIS_313-11428 | 0.000 | 1.000 | 0.000 | 0.000 | 0.000 | 0.000 | 0.000 |
| IRIS_313-11429 | 0.000 | 1.000 | 0.000 | 0.000 | 0.000 | 0.000 | 0.000 |
| IRIS_313-11431 | 0.000 | 0.028 | 0.000 | 0.000 | 0.000 | 0.000 | 0.972 |
| IRIS_313-11432 | 0.418 | 0.000 | 0.026 | 0.557 | 0.000 | 0.000 | 0.000 |
| IRIS_313-11433 | 0.000 | 0.000 | 0.000 | 0.003 | 0.000 | 0.754 | 0.243 |
| IRIS_313-11434 | 0.000 | 1.000 | 0.000 | 0.000 | 0.000 | 0.000 | 0.000 |
| IRIS_313-11435 | 0.000 | 1.000 | 0.000 | 0.000 | 0.000 | 0.000 | 0.000 |
| IRIS_313-11436 | 0.000 | 1.000 | 0.000 | 0.000 | 0.000 | 0.000 | 0.000 |
| IRIS_313-11437 | 0.000 | 0.000 | 0.000 | 0.000 | 0.000 | 0.062 | 0.938 |
| IRIS_313-11438 | 0.000 | 0.000 | 0.000 | 0.212 | 0.000 | 0.023 | 0.765 |
| IRIS_313-11439 | 0.000 | 1.000 | 0.000 | 0.000 | 0.000 | 0.000 | 0.000 |
| IRIS_313-11441 | 0.000 | 1.000 | 0.000 | 0.000 | 0.000 | 0.000 | 0.000 |
| IRIS_313-11442 | 0.019 | 0.005 | 0.000 | 0.680 | 0.000 | 0.296 | 0.000 |
| IRIS_313-11443 | 0.000 | 0.000 | 0.003 | 0.828 | 0.000 | 0.169 | 0.000 |
| IRIS_313-11445 | 0.079 | 0.000 | 0.000 | 0.921 | 0.000 | 0.000 | 0.000 |
| IRIS_313-11446 | 0.071 | 0.000 | 0.000 | 0.929 | 0.000 | 0.000 | 0.000 |
| IRIS_313-11447 | 0.000 | 0.000 | 0.000 | 0.782 | 0.000 | 0.218 | 0.000 |
| IRIS_313-11448 | 0.161 | 0.000 | 0.000 | 0.839 | 0.000 | 0.000 | 0.000 |
| IRIS_313-11449 | 0.344 | 0.000 | 0.000 | 0.655 | 0.000 | 0.001 | 0.000 |
| IRIS_313-11451 | 0.000 | 0.000 | 1.000 | 0.000 | 0.000 | 0.000 | 0.000 |
| IRIS_313-11452 | 0.755 | 0.063 | 0.000 | 0.181 | 0.001 | 0.000 | 0.000 |
| IRIS_313-11453 | 0.185 | 0.003 | 0.000 | 0.810 | 0.003 | 0.000 | 0.000 |
| IRIS_313-11454 | 1.000 | 0.000 | 0.000 | 0.000 | 0.000 | 0.000 | 0.000 |
| IRIS_313-11455 | 0.977 | 0.000 | 0.011 | 0.000 | 0.012 | 0.000 | 0.000 |
| IRIS_313-11456 | 1.000 | 0.000 | 0.000 | 0.000 | 0.000 | 0.000 | 0.000 |
| IRIS_313-11458 | 1.000 | 0.000 | 0.000 | 0.000 | 0.000 | 0.000 | 0.000 |
| IRIS_313-11460 | 0.180 | 0.000 | 0.000 | 0.820 | 0.000 | 0.000 | 0.000 |
| IRIS_313-11461 | 0.054 | 0.000 | 0.000 | 0.635 | 0.000 | 0.312 | 0.000 |
| IRIS_313-11462 | 1.000 | 0.000 | 0.000 | 0.000 | 0.000 | 0.000 | 0.000 |
| IRIS_313-11463 | 0.000 | 0.011 | 0.000 | 0.644 | 0.009 | 0.335 | 0.000 |
| IRIS_313-11464 | 0.000 | 0.998 | 0.000 | 0.000 | 0.000 | 0.002 | 0.000 |
| IRIS_313-11465 | 0.000 | 1.000 | 0.000 | 0.000 | 0.000 | 0.000 | 0.000 |
| IRIS_313-11467 | 0.000 | 0.023 | 0.000 | 0.362 | 0.000 | 0.000 | 0.615 |
| IRIS_313-11471 | 0.000 | 0.000 | 0.000 | 0.202 | 0.000 | 0.000 | 0.798 |
| IRIS_313-11472 | 0.000 | 0.003 | 0.000 | 0.775 | 0.000 | 0.172 | 0.050 |
| IRIS_313-11473 | 0.000 | 1.000 | 0.000 | 0.000 | 0.000 | 0.000 | 0.000 |
| IRIS_313-11476 | 0.766 | 0.070 | 0.000 | 0.163 | 0.000 | 0.000 | 0.000 |
| IRIS_313-11477 | 0.963 | 0.027 | 0.000 | 0.000 | 0.010 | 0.000 | 0.000 |
| IRIS_313-11478 | 0.318 | 0.000 | 0.000 | 0.682 | 0.000 | 0.000 | 0.000 |
| IRIS_313-11479 | 0.000 | 0.952 | 0.000 | 0.000 | 0.000 | 0.048 | 0.000 |
| IRIS_313-11480 | 0.000 | 0.000 | 0.000 | 0.000 | 0.000 | 0.000 | 1.000 |
| IRIS_313-11481 | 1.000 | 0.000 | 0.000 | 0.000 | 0.000 | 0.000 | 0.000 |
| IRIS_313-11482 | 0.855 | 0.000 | 0.000 | 0.143 | 0.000 | 0.001 | 0.001 |
| IRIS_313-11483 | 1.000 | 0.000 | 0.000 | 0.000 | 0.000 | 0.000 | 0.000 |
| IRIS_313-11484 | 0.753 | 0.000 | 0.014 | 0.233 | 0.000 | 0.000 | 0.000 |
| IRIS_313-11485 | 0.000 | 0.000 | 0.000 | 0.881 | 0.000 | 0.000 | 0.119 |
| IRIS_313-11486 | 0.407 | 0.000 | 0.001 | 0.592 | 0.000 | 0.000 | 0.000 |
| IRIS_313-11487 | 0.125 | 0.000 | 0.011 | 0.864 | 0.000 | 0.000 | 0.000 |
| IRIS_313-11489 | 0.751 | 0.000 | 0.029 | 0.000 | 0.000 | 0.000 | 0.220 |
| IRIS_313-11490 | 0.024 | 0.000 | 0.000 | 0.976 | 0.000 | 0.000 | 0.000 |
| IRIS_313-11491 | 0.063 | 0.000 | 0.000 | 0.937 | 0.000 | 0.000 | 0.000 |
| IRIS_313-11492 | 1.000 | 0.000 | 0.000 | 0.000 | 0.000 | 0.000 | 0.000 |
| IRIS_313-11493 | 0.000 | 0.000 | 0.000 | 0.534 | 0.000 | 0.129 | 0.336 |
| IRIS_313-11494 | 0.000 | 0.807 | 0.000 | 0.000 | 0.193 | 0.000 | 0.000 |
| IRIS_313-11495 | 0.000 | 1.000 | 0.000 | 0.000 | 0.000 | 0.000 | 0.000 |
| IRIS_313-11496 | 0.000 | 0.901 | 0.000 | 0.000 | 0.076 | 0.023 | 0.000 |
| IRIS_313-11497 | 0.187 | 0.000 | 0.000 | 0.800 | 0.000 | 0.013 | 0.000 |
| IRIS_313-11498 | 0.000 | 0.920 | 0.000 | 0.000 | 0.000 | 0.080 | 0.000 |
| IRIS_313-11499 | 0.000 | 1.000 | 0.000 | 0.000 | 0.000 | 0.000 | 0.000 |

| | | | | | | | |
|---|---|---|---|---|---|---|---|
| IRIS_313-11500 | 0.000 | 0.000 | 0.000 | 0.989 | 0.011 | 0.000 | 0.000 |
| IRIS_313-11505 | 0.048 | 0.000 | 0.000 | 0.633 | 0.000 | 0.320 | 0.000 |
| IRIS_313-11506 | 0.022 | 0.000 | 0.000 | 0.800 | 0.000 | 0.000 | 0.178 |
| IRIS_313-11507 | 0.000 | 0.000 | 0.000 | 0.000 | 0.000 | 0.045 | 0.955 |
| IRIS_313-11508 | 0.000 | 0.000 | 0.000 | 1.000 | 0.000 | 0.000 | 0.000 |
| IRIS_313-11509 | 0.000 | 0.000 | 0.000 | 1.000 | 0.000 | 0.000 | 0.000 |
| IRIS_313-11510 | 0.040 | 0.000 | 0.000 | 0.390 | 0.000 | 0.113 | 0.457 |
| IRIS_313-11511 | 0.000 | 1.000 | 0.000 | 0.000 | 0.000 | 0.000 | 0.000 |
| IRIS_313-11512 | 0.000 | 0.005 | 0.000 | 0.676 | 0.000 | 0.319 | 0.000 |
| IRIS_313-11513 | 0.000 | 0.000 | 0.000 | 0.000 | 0.000 | 0.348 | 0.652 |
| IRIS_313-11515 | 0.041 | 0.000 | 0.000 | 0.453 | 0.000 | 0.025 | 0.481 |
| IRIS_313-11516 | 0.000 | 0.000 | 0.000 | 0.225 | 0.000 | 0.000 | 0.775 |
| IRIS_313-11517 | 0.000 | 0.000 | 0.000 | 0.113 | 0.000 | 0.143 | 0.744 |
| IRIS_313-11521 | 0.000 | 0.000 | 0.004 | 0.689 | 0.000 | 0.307 | 0.000 |
| IRIS_313-11522 | 0.000 | 0.127 | 0.000 | 0.000 | 0.873 | 0.000 | 0.000 |
| IRIS_313-11523 | 0.000 | 0.000 | 0.000 | 1.000 | 0.000 | 0.000 | 0.000 |
| IRIS_313-11524 | 0.000 | 1.000 | 0.000 | 0.000 | 0.000 | 0.000 | 0.000 |
| IRIS_313-11525 | 0.000 | 0.000 | 0.000 | 0.902 | 0.000 | 0.098 | 0.000 |
| IRIS_313-11526 | 0.000 | 1.000 | 0.000 | 0.000 | 0.000 | 0.000 | 0.000 |
| IRIS_313-11527 | 0.000 | 1.000 | 0.000 | 0.000 | 0.000 | 0.000 | 0.000 |
| IRIS_313-11528 | 0.000 | 0.000 | 0.000 | 1.000 | 0.000 | 0.000 | 0.000 |
| IRIS_313-11530 | 0.000 | 0.000 | 0.000 | 1.000 | 0.000 | 0.000 | 0.000 |
| IRIS_313-11532 | 0.000 | 1.000 | 0.000 | 0.000 | 0.000 | 0.000 | 0.000 |
| IRIS_313-11534 | 0.000 | 0.003 | 0.000 | 0.567 | 0.002 | 0.000 | 0.428 |
| IRIS_313-11535 | 0.000 | 0.003 | 0.000 | 0.284 | 0.007 | 0.000 | 0.706 |
| IRIS_313-11536 | 0.000 | 0.000 | 0.000 | 0.000 | 1.000 | 0.000 | 0.000 |
| IRIS_313-11537 | 0.000 | 0.702 | 0.000 | 0.000 | 0.298 | 0.000 | 0.000 |
| IRIS_313-11538 | 0.000 | 0.000 | 0.000 | 0.000 | 0.081 | 0.209 | 0.710 |
| IRIS_313-11539 | 0.000 | 0.818 | 0.000 | 0.000 | 0.182 | 0.000 | 0.000 |
| IRIS_313-11540 | 0.000 | 1.000 | 0.000 | 0.000 | 0.000 | 0.000 | 0.000 |
| IRIS_313-11541 | 0.000 | 0.000 | 0.000 | 0.800 | 0.000 | 0.195 | 0.005 |
| IRIS_313-11542 | 0.000 | 0.128 | 0.580 | 0.000 | 0.131 | 0.013 | 0.148 |
| IRIS_313-11543 | 0.000 | 0.000 | 0.000 | 0.958 | 0.000 | 0.042 | 0.000 |
| IRIS_313-11544 | 0.000 | 0.003 | 0.000 | 0.899 | 0.000 | 0.098 | 0.000 |
| IRIS_313-11545 | 0.000 | 0.004 | 0.000 | 0.898 | 0.000 | 0.098 | 0.000 |
| IRIS_313-11546 | 0.000 | 0.000 | 0.000 | 0.701 | 0.000 | 0.299 | 0.000 |
| IRIS_313-11547 | 0.000 | 0.042 | 0.000 | 0.729 | 0.000 | 0.090 | 0.139 |
| IRIS_313-11548 | 0.000 | 0.000 | 0.004 | 0.723 | 0.000 | 0.272 | 0.000 |
| IRIS_313-11549 | 0.000 | 0.000 | 0.000 | 0.795 | 0.000 | 0.205 | 0.000 |
| IRIS_313-11551 | 0.076 | 0.000 | 0.000 | 0.721 | 0.015 | 0.000 | 0.189 |
| IRIS_313-11554 | 0.000 | 0.000 | 0.000 | 0.553 | 0.000 | 0.000 | 0.447 |
| IRIS_313-11555 | 0.028 | 0.000 | 0.000 | 0.966 | 0.000 | 0.006 | 0.000 |
| IRIS_313-11556 | 0.000 | 0.000 | 0.000 | 0.330 | 0.000 | 0.000 | 0.670 |
| IRIS_313-11557 | 1.000 | 0.000 | 0.000 | 0.000 | 0.000 | 0.000 | 0.000 |
| IRIS_313-11558 | 0.049 | 0.000 | 0.000 | 0.858 | 0.000 | 0.092 | 0.000 |
| IRIS_313-11561 | 0.133 | 0.000 | 0.002 | 0.865 | 0.000 | 0.000 | 0.000 |
| IRIS_313-11563 | 0.143 | 0.000 | 0.000 | 0.857 | 0.000 | 0.000 | 0.000 |
| IRIS_313-11564 | 0.000 | 0.109 | 0.891 | 0.000 | 0.000 | 0.000 | 0.000 |
| IRIS_313-11565 | 0.139 | 0.000 | 0.000 | 0.861 | 0.000 | 0.000 | 0.000 |
| IRIS_313-11566 | 0.082 | 0.000 | 0.000 | 0.852 | 0.000 | 0.066 | 0.000 |
| IRIS_313-11567 | 0.000 | 0.000 | 1.000 | 0.000 | 0.000 | 0.000 | 0.000 |
| IRIS_313-11568 | 0.000 | 0.000 | 0.000 | 0.010 | 0.000 | 0.990 | 0.000 |
| IRIS_313-11570 | 0.000 | 0.000 | 0.000 | 0.000 | 0.067 | 0.643 | 0.290 |
| IRIS_313-11571 | 0.000 | 0.000 | 0.000 | 0.000 | 1.000 | 0.000 | 0.000 |
| IRIS_313-11572 | 0.000 | 0.000 | 0.000 | 0.285 | 0.000 | 0.715 | 0.000 |
| IRIS_313-11573 | 0.000 | 0.000 | 0.000 | 0.300 | 0.000 | 0.658 | 0.042 |
| IRIS_313-11574 | 0.000 | 0.000 | 0.000 | 0.000 | 1.000 | 0.000 | 0.000 |
| IRIS_313-11575 | 0.000 | 0.000 | 0.000 | 0.000 | 1.000 | 0.000 | 0.000 |
| IRIS_313-11576 | 0.000 | 0.000 | 0.033 | 0.380 | 0.000 | 0.587 | 0.000 |
| IRIS_313-11577 | 0.000 | 0.001 | 0.011 | 0.248 | 0.000 | 0.693 | 0.047 |
| IRIS_313-11578 | 0.000 | 0.000 | 0.000 | 0.000 | 1.000 | 0.000 | 0.000 |
| IRIS_313-11579 | 0.000 | 0.000 | 0.000 | 0.000 | 0.000 | 1.000 | 0.000 |
| IRIS_313-11580 | 0.000 | 0.000 | 0.012 | 0.488 | 0.026 | 0.474 | 0.000 |
| IRIS_313-11582 | 0.000 | 0.000 | 0.000 | 0.002 | 0.979 | 0.020 | 0.000 |
| IRIS_313-11584 | 0.000 | 0.000 | 0.000 | 0.000 | 1.000 | 0.000 | 0.000 |
| IRIS_313-11585 | 0.000 | 0.000 | 0.000 | 0.000 | 0.000 | 0.782 | 0.218 |
| IRIS_313-11586 | 0.000 | 0.000 | 0.000 | 0.000 | 1.000 | 0.000 | 0.000 |

| | | | | | | | |
|---|---|---|---|---|---|---|---|
| IRIS_313-11588 | 0.000 | 0.000 | 0.000 | 0.474 | 0.010 | 0.000 | 0.516 |
| IRIS_313-11591 | 0.000 | 0.391 | 0.000 | 0.609 | 0.000 | 0.000 | 0.000 |
| IRIS_313-11594 | 0.000 | 0.000 | 0.000 | 0.667 | 0.000 | 0.333 | 0.000 |
| IRIS_313-11595 | 0.987 | 0.000 | 0.000 | 0.000 | 0.013 | 0.000 | 0.000 |
| IRIS_313-11596 | 0.441 | 0.000 | 0.000 | 0.000 | 0.000 | 0.559 | 0.000 |
| IRIS_313-11597 | 0.080 | 0.000 | 0.000 | 0.831 | 0.000 | 0.089 | 0.000 |
| IRIS_313-11598 | 0.572 | 0.000 | 0.000 | 0.289 | 0.000 | 0.139 | 0.000 |
| IRIS_313-11599 | 0.006 | 0.000 | 0.000 | 0.994 | 0.000 | 0.000 | 0.000 |
| IRIS_313-11600 | 0.056 | 0.000 | 0.000 | 0.911 | 0.000 | 0.033 | 0.000 |
| IRIS_313-11602 | 1.000 | 0.000 | 0.000 | 0.000 | 0.000 | 0.000 | 0.000 |
| IRIS_313-11603 | 0.992 | 0.000 | 0.000 | 0.000 | 0.008 | 0.000 | 0.000 |
| IRIS_313-11604 | 1.000 | 0.000 | 0.000 | 0.000 | 0.000 | 0.000 | 0.000 |
| IRIS_313-11606 | 0.052 | 0.000 | 0.000 | 0.948 | 0.000 | 0.000 | 0.000 |
| IRIS_313-11607 | 0.035 | 0.000 | 0.000 | 0.965 | 0.000 | 0.000 | 0.000 |
| IRIS_313-11608 | 0.000 | 0.000 | 0.000 | 0.444 | 0.000 | 0.000 | 0.556 |
| IRIS_313-11609 | 0.000 | 0.000 | 0.000 | 0.652 | 0.000 | 0.000 | 0.348 |
| IRIS_313-11610 | 0.000 | 0.000 | 0.000 | 0.478 | 0.000 | 0.000 | 0.522 |
| IRIS_313-11611 | 0.022 | 0.927 | 0.000 | 0.000 | 0.000 | 0.049 | 0.002 |
| IRIS_313-11615 | 0.000 | 0.000 | 0.000 | 0.950 | 0.000 | 0.050 | 0.000 |
| IRIS_313-11616 | 0.491 | 0.000 | 0.000 | 0.509 | 0.000 | 0.000 | 0.000 |
| IRIS_313-11617 | 0.955 | 0.000 | 0.001 | 0.000 | 0.000 | 0.000 | 0.045 |
| IRIS_313-11618 | 0.987 | 0.000 | 0.000 | 0.013 | 0.000 | 0.000 | 0.000 |
| IRIS_313-11619 | 1.000 | 0.000 | 0.000 | 0.000 | 0.000 | 0.000 | 0.000 |
| IRIS_313-11620 | 0.978 | 0.000 | 0.000 | 0.000 | 0.000 | 0.022 | 0.000 |
| IRIS_313-11621 | 0.000 | 0.000 | 0.000 | 0.249 | 0.000 | 0.000 | 0.751 |
| IRIS_313-11622 | 0.000 | 0.000 | 0.000 | 0.000 | 0.000 | 1.000 | 0.000 |
| IRIS_313-11623 | 0.000 | 0.739 | 0.000 | 0.000 | 0.261 | 0.000 | 0.000 |
| IRIS_313-11624 | 0.000 | 0.000 | 0.000 | 0.000 | 0.000 | 1.000 | 0.000 |
| IRIS_313-11625 | 0.000 | 0.000 | 0.864 | 0.000 | 0.000 | 0.054 | 0.081 |
| IRIS_313-11626 | 0.000 | 0.116 | 0.884 | 0.000 | 0.000 | 0.000 | 0.000 |
| IRIS_313-11627 | 0.000 | 0.000 | 1.000 | 0.000 | 0.000 | 0.000 | 0.000 |
| IRIS_313-11628 | 0.298 | 0.000 | 0.377 | 0.325 | 0.000 | 0.000 | 0.000 |
| IRIS_313-11629 | 1.000 | 0.000 | 0.000 | 0.000 | 0.000 | 0.000 | 0.000 |
| IRIS_313-11630 | 0.000 | 0.000 | 1.000 | 0.000 | 0.000 | 0.000 | 0.000 |
| IRIS_313-11632 | 0.000 | 0.000 | 1.000 | 0.000 | 0.000 | 0.000 | 0.000 |
| IRIS_313-11635 | 0.000 | 0.000 | 0.000 | 0.956 | 0.000 | 0.044 | 0.000 |
| IRIS_313-11636 | 1.000 | 0.000 | 0.000 | 0.000 | 0.000 | 0.000 | 0.000 |
| IRIS_313-11638 | 0.058 | 0.000 | 0.000 | 0.942 | 0.000 | 0.000 | 0.000 |
| IRIS_313-11639 | 0.054 | 0.000 | 0.000 | 0.329 | 0.000 | 0.617 | 0.000 |
| IRIS_313-11640 | 0.061 | 0.000 | 0.000 | 0.939 | 0.000 | 0.000 | 0.000 |
| IRIS_313-11641 | 0.064 | 0.000 | 0.000 | 0.936 | 0.000 | 0.000 | 0.000 |
| IRIS_313-11642 | 0.046 | 0.000 | 0.000 | 0.954 | 0.000 | 0.000 | 0.000 |
| IRIS_313-11643 | 0.213 | 0.000 | 0.000 | 0.000 | 0.000 | 0.000 | 0.787 |
| IRIS_313-11644 | 0.163 | 0.000 | 0.007 | 0.577 | 0.000 | 0.253 | 0.000 |
| IRIS_313-11645 | 0.116 | 0.000 | 0.000 | 0.739 | 0.000 | 0.145 | 0.000 |
| IRIS_313-11646 | 0.021 | 0.000 | 0.000 | 0.856 | 0.000 | 0.000 | 0.123 |
| IRIS_313-11647 | 0.155 | 0.000 | 0.000 | 0.845 | 0.000 | 0.000 | 0.000 |
| IRIS_313-11648 | 0.033 | 0.000 | 0.000 | 0.967 | 0.000 | 0.000 | 0.000 |
| IRIS_313-11650 | 0.065 | 0.000 | 0.180 | 0.755 | 0.000 | 0.000 | 0.000 |
| IRIS_313-11651 | 0.000 | 0.055 | 0.000 | 0.000 | 0.945 | 0.000 | 0.000 |
| IRIS_313-11652 | 0.000 | 0.000 | 0.000 | 0.000 | 1.000 | 0.000 | 0.000 |
| IRIS_313-11653 | 0.000 | 0.000 | 0.000 | 0.000 | 1.000 | 0.000 | 0.000 |
| IRIS_313-11654 | 0.000 | 0.082 | 0.000 | 0.000 | 0.918 | 0.000 | 0.000 |
| IRIS_313-11655 | 0.000 | 0.197 | 0.000 | 0.000 | 0.803 | 0.000 | 0.000 |
| IRIS_313-11656 | 0.000 | 0.011 | 0.005 | 0.000 | 0.020 | 0.035 | 0.929 |
| IRIS_313-11657 | 0.058 | 0.000 | 0.004 | 0.938 | 0.000 | 0.000 | 0.000 |
| IRIS_313-11658 | 0.000 | 1.000 | 0.000 | 0.000 | 0.000 | 0.000 | 0.000 |
| IRIS_313-11659 | 0.000 | 1.000 | 0.000 | 0.000 | 0.000 | 0.000 | 0.000 |
| IRIS_313-11660 | 0.000 | 1.000 | 0.000 | 0.000 | 0.000 | 0.000 | 0.000 |
| IRIS_313-11661 | 0.000 | 0.005 | 0.000 | 0.000 | 0.995 | 0.000 | 0.000 |
| IRIS_313-11663 | 0.411 | 0.000 | 0.000 | 0.589 | 0.000 | 0.000 | 0.000 |
| IRIS_313-11664 | 0.000 | 0.000 | 0.000 | 0.533 | 0.000 | 0.467 | 0.000 |
| IRIS_313-11665 | 0.000 | 0.000 | 0.000 | 0.052 | 0.000 | 0.948 | 0.000 |
| IRIS_313-11666 | 0.000 | 0.000 | 0.000 | 0.257 | 0.014 | 0.729 | 0.000 |
| IRIS_313-11667 | 0.000 | 0.000 | 0.000 | 0.194 | 0.000 | 0.806 | 0.000 |
| IRIS_313-11668 | 0.000 | 0.000 | 0.019 | 0.000 | 0.000 | 0.981 | 0.000 |
| IRIS_313-11669 | 0.000 | 0.000 | 0.000 | 0.000 | 0.061 | 0.679 | 0.260 |

| | | | | | | | |
|---|---|---|---|---|---|---|---|
| IRIS_313-11671 | 0.120 | 0.000 | 0.000 | 0.880 | 0.000 | 0.000 | 0.000 |
| IRIS_313-11672 | 0.000 | 0.000 | 0.000 | 0.000 | 1.000 | 0.000 | 0.000 |
| IRIS_313-11673 | 0.000 | 1.000 | 0.000 | 0.000 | 0.000 | 0.000 | 0.000 |
| IRIS_313-11674 | 0.000 | 0.000 | 0.000 | 1.000 | 0.000 | 0.000 | 0.000 |
| IRIS_313-11676 | 0.000 | 0.000 | 0.000 | 0.998 | 0.002 | 0.000 | 0.000 |
| IRIS_313-11677 | 0.000 | 0.000 | 0.000 | 0.838 | 0.000 | 0.104 | 0.059 |
| IRIS_313-11678 | 0.000 | 0.000 | 0.000 | 1.000 | 0.000 | 0.000 | 0.000 |
| IRIS_313-11679 | 0.000 | 0.000 | 0.000 | 0.996 | 0.000 | 0.000 | 0.004 |
| IRIS_313-11680 | 0.000 | 0.000 | 0.000 | 1.000 | 0.000 | 0.000 | 0.000 |
| IRIS_313-11681 | 0.061 | 0.015 | 0.000 | 0.821 | 0.000 | 0.000 | 0.103 |
| IRIS_313-11682 | 0.000 | 0.000 | 0.000 | 0.959 | 0.004 | 0.037 | 0.000 |
| IRIS_313-11683 | 0.000 | 0.000 | 0.000 | 1.000 | 0.000 | 0.000 | 0.000 |
| IRIS_313-11684 | 0.000 | 0.000 | 0.000 | 1.000 | 0.000 | 0.000 | 0.000 |
| IRIS_313-11685 | 0.000 | 0.000 | 0.000 | 1.000 | 0.000 | 0.000 | 0.000 |
| IRIS_313-11686 | 0.000 | 0.000 | 0.000 | 1.000 | 0.000 | 0.000 | 0.000 |
| IRIS_313-11687 | 0.000 | 0.000 | 0.000 | 1.000 | 0.000 | 0.000 | 0.000 |
| IRIS_313-11688 | 0.000 | 0.000 | 0.000 | 0.975 | 0.000 | 0.025 | 0.000 |
| IRIS_313-11689 | 0.000 | 0.000 | 0.000 | 0.000 | 1.000 | 0.000 | 0.000 |
| IRIS_313-11690 | 0.000 | 0.282 | 0.070 | 0.016 | 0.188 | 0.444 | 0.000 |
| IRIS_313-11691 | 0.000 | 0.539 | 0.064 | 0.000 | 0.397 | 0.000 | 0.000 |
| IRIS_313-11692 | 0.000 | 0.000 | 0.000 | 0.002 | 0.000 | 0.998 | 0.000 |
| IRIS_313-11693 | 0.001 | 0.026 | 0.000 | 0.559 | 0.008 | 0.399 | 0.008 |
| IRIS_313-11694 | 0.000 | 0.038 | 0.000 | 0.438 | 0.000 | 0.492 | 0.032 |
| IRIS_313-11695 | 0.000 | 0.000 | 0.000 | 0.277 | 0.013 | 0.711 | 0.000 |
| IRIS_313-11698 | 0.000 | 0.000 | 1.000 | 0.000 | 0.000 | 0.000 | 0.000 |
| IRIS_313-11700 | 0.000 | 0.000 | 0.000 | 0.956 | 0.000 | 0.042 | 0.002 |
| IRIS_313-11702 | 0.000 | 0.105 | 0.000 | 0.000 | 0.895 | 0.000 | 0.000 |
| IRIS_313-11704 | 0.043 | 0.015 | 0.001 | 0.815 | 0.000 | 0.047 | 0.080 |
| IRIS_313-11705 | 0.000 | 0.001 | 0.002 | 0.997 | 0.000 | 0.000 | 0.000 |
| IRIS_313-11706 | 0.000 | 0.000 | 0.036 | 0.507 | 0.002 | 0.000 | 0.455 |
| IRIS_313-11707 | 0.000 | 0.000 | 0.000 | 0.959 | 0.000 | 0.000 | 0.041 |
| IRIS_313-11708 | 0.000 | 0.000 | 0.000 | 1.000 | 0.000 | 0.000 | 0.000 |
| IRIS_313-11709 | 0.000 | 0.000 | 0.000 | 1.000 | 0.000 | 0.000 | 0.000 |
| IRIS_313-11710 | 0.000 | 0.000 | 0.000 | 1.000 | 0.000 | 0.000 | 0.000 |
| IRIS_313-11711 | 0.000 | 0.000 | 0.000 | 0.964 | 0.000 | 0.000 | 0.036 |
| IRIS_313-11712 | 1.000 | 0.000 | 0.000 | 0.000 | 0.000 | 0.000 | 0.000 |
| IRIS_313-11713 | 0.000 | 0.000 | 0.000 | 0.445 | 0.008 | 0.000 | 0.547 |
| IRIS_313-11714 | 0.000 | 0.152 | 0.000 | 0.000 | 0.108 | 0.229 | 0.511 |
| IRIS_313-11716 | 0.000 | 0.025 | 0.000 | 0.829 | 0.000 | 0.000 | 0.146 |
| IRIS_313-11717 | 0.000 | 0.000 | 0.000 | 0.223 | 0.000 | 0.000 | 0.777 |
| IRIS_313-11719 | 0.000 | 0.000 | 0.000 | 0.999 | 0.000 | 0.001 | 0.000 |
| IRIS_313-11720 | 0.000 | 0.000 | 0.000 | 1.000 | 0.000 | 0.000 | 0.000 |
| IRIS_313-11721 | 0.000 | 0.000 | 0.000 | 1.000 | 0.000 | 0.000 | 0.000 |
| IRIS_313-11722 | 0.000 | 0.000 | 0.000 | 0.501 | 0.000 | 0.499 | 0.000 |
| IRIS_313-11723 | 0.049 | 0.000 | 0.000 | 0.951 | 0.000 | 0.000 | 0.000 |
| IRIS_313-11724 | 0.000 | 0.000 | 0.000 | 0.996 | 0.000 | 0.004 | 0.000 |
| IRIS_313-11725 | 0.000 | 0.000 | 0.000 | 0.000 | 1.000 | 0.000 | 0.000 |
| IRIS_313-11726 | 0.000 | 0.000 | 0.000 | 0.293 | 0.012 | 0.695 | 0.000 |
| IRIS_313-11727 | 0.000 | 0.000 | 0.000 | 0.000 | 0.000 | 1.000 | 0.000 |
| IRIS_313-11728 | 0.000 | 0.000 | 0.000 | 0.904 | 0.000 | 0.096 | 0.000 |
| IRIS_313-11729 | 0.000 | 0.000 | 0.000 | 0.000 | 0.000 | 0.713 | 0.287 |
| IRIS_313-11730 | 0.000 | 0.000 | 0.000 | 0.029 | 0.000 | 0.389 | 0.582 |
| IRIS_313-11731 | 0.000 | 0.000 | 0.000 | 0.146 | 0.000 | 0.622 | 0.232 |
| IRIS_313-11732 | 0.000 | 0.000 | 0.000 | 0.000 | 0.000 | 0.932 | 0.068 |
| IRIS_313-11733 | 0.000 | 0.000 | 0.000 | 0.000 | 0.000 | 0.810 | 0.190 |
| IRIS_313-11734 | 0.000 | 0.000 | 0.000 | 0.003 | 0.000 | 0.822 | 0.175 |
| IRIS_313-11735 | 0.043 | 0.000 | 0.002 | 0.476 | 0.000 | 0.447 | 0.032 |
| IRIS_313-11736 | 0.000 | 1.000 | 0.000 | 0.000 | 0.000 | 0.000 | 0.000 |
| IRIS_313-11737 | 0.927 | 0.000 | 0.000 | 0.000 | 0.000 | 0.000 | 0.073 |
| IRIS_313-11738 | 0.088 | 0.000 | 0.000 | 0.912 | 0.000 | 0.000 | 0.000 |
| IRIS_313-11739 | 0.000 | 1.000 | 0.000 | 0.000 | 0.000 | 0.000 | 0.000 |
| IRIS_313-11740 | 0.000 | 0.000 | 0.000 | 0.780 | 0.000 | 0.220 | 0.000 |
| IRIS_313-11741 | 0.139 | 0.000 | 0.000 | 0.651 | 0.000 | 0.210 | 0.000 |
| IRIS_313-11742 | 1.000 | 0.000 | 0.000 | 0.000 | 0.000 | 0.000 | 0.000 |
| IRIS_313-11743 | 0.000 | 0.045 | 0.955 | 0.000 | 0.000 | 0.000 | 0.000 |
| IRIS_313-11744 | 0.000 | 0.000 | 0.020 | 0.414 | 0.000 | 0.566 | 0.000 |
| IRIS_313-11745 | 0.000 | 0.000 | 0.000 | 0.075 | 0.000 | 0.925 | 0.000 |

| | | | | | | | |
|---|---|---|---|---|---|---|---|
| IRIS_313-11746 | 0.000 | 0.000 | 0.000 | 0.198 | 0.000 | 0.802 | 0.000 |
| IRIS_313-11747 | 0.000 | 0.000 | 0.000 | 0.272 | 0.000 | 0.728 | 0.000 |
| IRIS_313-11748 | 0.000 | 0.000 | 0.027 | 0.381 | 0.000 | 0.591 | 0.001 |
| IRIS_313-11750 | 0.000 | 0.000 | 0.000 | 0.073 | 0.000 | 0.455 | 0.472 |
| IRIS_313-11751 | 0.000 | 0.000 | 0.000 | 0.040 | 0.000 | 0.396 | 0.564 |
| IRIS_313-11752 | 0.000 | 0.000 | 0.000 | 0.000 | 0.000 | 1.000 | 0.000 |
| IRIS_313-11753 | 0.000 | 0.000 | 0.000 | 0.000 | 0.000 | 1.000 | 0.000 |
| IRIS_313-11754 | 0.154 | 0.512 | 0.000 | 0.262 | 0.000 | 0.073 | 0.000 |
| IRIS_313-11755 | 0.000 | 1.000 | 0.000 | 0.000 | 0.000 | 0.000 | 0.000 |
| IRIS_313-11756 | 0.000 | 1.000 | 0.000 | 0.000 | 0.000 | 0.000 | 0.000 |
| IRIS_313-11757 | 0.000 | 0.000 | 0.000 | 0.955 | 0.006 | 0.000 | 0.039 |
| IRIS_313-11758 | 0.004 | 0.000 | 0.000 | 0.996 | 0.000 | 0.000 | 0.000 |
| IRIS_313-11759 | 0.000 | 1.000 | 0.000 | 0.000 | 0.000 | 0.000 | 0.000 |
| IRIS_313-11760 | 0.061 | 0.000 | 0.000 | 0.939 | 0.000 | 0.000 | 0.000 |
| IRIS_313-11761 | 0.114 | 0.000 | 0.000 | 0.886 | 0.000 | 0.000 | 0.000 |
| IRIS_313-11762 | 0.073 | 0.000 | 0.000 | 0.925 | 0.000 | 0.002 | 0.000 |
| IRIS_313-11763 | 0.000 | 0.000 | 0.000 | 0.000 | 0.000 | 0.415 | 0.585 |
| IRIS_313-11764 | 0.000 | 0.000 | 0.000 | 0.943 | 0.000 | 0.000 | 0.057 |
| IRIS_313-11765 | 0.000 | 0.000 | 1.000 | 0.000 | 0.000 | 0.000 | 0.000 |
| IRIS_313-11766 | 0.581 | 0.049 | 0.000 | 0.370 | 0.000 | 0.000 | 0.000 |
| IRIS_313-11767 | 0.159 | 0.170 | 0.022 | 0.628 | 0.021 | 0.001 | 0.000 |
| IRIS_313-11772 | 0.317 | 0.068 | 0.000 | 0.616 | 0.000 | 0.000 | 0.000 |
| IRIS_313-11773 | 0.016 | 0.000 | 0.000 | 0.984 | 0.000 | 0.000 | 0.000 |
| IRIS_313-11779 | 0.000 | 0.003 | 0.000 | 0.936 | 0.002 | 0.059 | 0.000 |
| IRIS_313-11780 | 0.000 | 0.000 | 0.000 | 1.000 | 0.000 | 0.000 | 0.000 |
| IRIS_313-11782 | 0.184 | 0.000 | 0.022 | 0.794 | 0.000 | 0.000 | 0.000 |
| IRIS_313-11783 | 0.000 | 0.000 | 0.000 | 1.000 | 0.000 | 0.000 | 0.000 |
| IRIS_313-11784 | 0.000 | 0.056 | 0.000 | 0.174 | 0.000 | 0.000 | 0.771 |
| IRIS_313-11785 | 0.000 | 0.000 | 0.000 | 1.000 | 0.000 | 0.000 | 0.000 |
| IRIS_313-11786 | 0.000 | 0.000 | 0.000 | 0.917 | 0.000 | 0.083 | 0.000 |
| IRIS_313-11787 | 0.055 | 0.000 | 0.000 | 0.945 | 0.000 | 0.000 | 0.000 |
| IRIS_313-11788 | 0.000 | 0.000 | 0.065 | 0.000 | 0.935 | 0.000 | 0.000 |
| IRIS_313-11789 | 0.113 | 0.000 | 0.000 | 0.887 | 0.000 | 0.000 | 0.000 |
| IRIS_313-11790 | 0.023 | 0.890 | 0.087 | 0.000 | 0.000 | 0.000 | 0.000 |
| IRIS_313-11791 | 0.276 | 0.098 | 0.000 | 0.626 | 0.000 | 0.000 | 0.000 |
| IRIS_313-11792 | 0.000 | 1.000 | 0.000 | 0.000 | 0.000 | 0.000 | 0.000 |
| IRIS_313-11793 | 0.200 | 0.000 | 0.000 | 0.798 | 0.003 | 0.000 | 0.000 |
| IRIS_313-11794 | 0.129 | 0.000 | 0.000 | 0.871 | 0.000 | 0.000 | 0.000 |
| IRIS_313-11795 | 0.000 | 0.000 | 0.000 | 0.000 | 0.000 | 1.000 | 0.000 |
| IRIS_313-11796 | 0.000 | 0.000 | 0.000 | 0.067 | 0.000 | 0.933 | 0.000 |
| IRIS_313-11797 | 0.000 | 0.000 | 0.000 | 0.207 | 0.029 | 0.763 | 0.000 |
| IRIS_313-11798 | 0.000 | 0.000 | 0.000 | 0.181 | 0.000 | 0.052 | 0.767 |
| IRIS_313-11799 | 0.000 | 0.000 | 0.000 | 0.000 | 0.000 | 1.000 | 0.000 |
| IRIS_313-11800 | 0.000 | 0.000 | 0.000 | 0.000 | 1.000 | 0.000 | 0.000 |
| IRIS_313-11801 | 0.000 | 0.000 | 0.000 | 0.000 | 0.000 | 0.500 | 0.500 |
| IRIS_313-11802 | 0.000 | 0.000 | 0.000 | 0.006 | 0.000 | 0.983 | 0.010 |
| IRIS_313-11803 | 0.000 | 0.000 | 0.000 | 0.000 | 1.000 | 0.000 | 0.000 |
| IRIS_313-11804 | 0.000 | 0.000 | 0.000 | 0.000 | 0.000 | 1.000 | 0.000 |
| IRIS_313-11805 | 0.000 | 0.000 | 0.000 | 0.011 | 0.000 | 0.989 | 0.000 |
| IRIS_313-11806 | 0.000 | 0.000 | 0.000 | 0.000 | 0.000 | 1.000 | 0.000 |
| IRIS_313-11807 | 0.006 | 0.053 | 0.000 | 0.000 | 0.027 | 0.247 | 0.667 |
| IRIS_313-11808 | 0.000 | 0.097 | 0.000 | 0.248 | 0.028 | 0.000 | 0.627 |
| IRIS_313-11809 | 1.000 | 0.000 | 0.000 | 0.000 | 0.000 | 0.000 | 0.000 |
| IRIS_313-11810 | 0.000 | 0.000 | 0.000 | 1.000 | 0.000 | 0.000 | 0.000 |
| IRIS_313-11811 | 0.074 | 0.000 | 0.000 | 0.926 | 0.000 | 0.000 | 0.000 |
| IRIS_313-11812 | 0.000 | 0.000 | 0.000 | 1.000 | 0.000 | 0.000 | 0.000 |
| IRIS_313-11813 | 0.040 | 0.000 | 0.000 | 0.960 | 0.000 | 0.000 | 0.000 |
| IRIS_313-11814 | 0.000 | 0.000 | 0.000 | 1.000 | 0.000 | 0.000 | 0.000 |
| IRIS_313-11815 | 0.000 | 0.000 | 0.000 | 1.000 | 0.000 | 0.000 | 0.000 |
| IRIS_313-11816 | 0.046 | 0.000 | 0.002 | 0.604 | 0.000 | 0.349 | 0.000 |
| IRIS_313-11817 | 0.000 | 0.000 | 0.000 | 0.609 | 0.000 | 0.391 | 0.000 |
| IRIS_313-11818 | 0.061 | 0.000 | 0.006 | 0.650 | 0.000 | 0.283 | 0.000 |
| IRIS_313-11819 | 0.000 | 0.000 | 0.000 | 0.907 | 0.000 | 0.070 | 0.023 |
| IRIS_313-11820 | 0.000 | 0.000 | 0.000 | 1.000 | 0.000 | 0.000 | 0.000 |
| IRIS_313-11821 | 0.073 | 0.000 | 0.009 | 0.918 | 0.000 | 0.000 | 0.000 |
| IRIS_313-11822 | 0.039 | 0.000 | 0.000 | 0.961 | 0.000 | 0.000 | 0.000 |
| IRIS_313-11823 | 0.071 | 0.000 | 0.000 | 0.929 | 0.000 | 0.000 | 0.000 |

| | | | | | | | |
|---|---|---|---|---|---|---|---|
| IRIS_313-11824 | 0.036 | 0.000 | 0.000 | 0.964 | 0.000 | 0.000 | 0.000 |
| IRIS_313-11825 | 0.000 | 0.000 | 1.000 | 0.000 | 0.000 | 0.000 | 0.000 |
| IRIS_313-11826 | 0.000 | 0.000 | 1.000 | 0.000 | 0.000 | 0.000 | 0.000 |
| IRIS_313-11827 | 0.086 | 0.000 | 0.000 | 0.914 | 0.000 | 0.000 | 0.000 |
| IRIS_313-11828 | 0.095 | 0.000 | 0.000 | 0.905 | 0.000 | 0.000 | 0.000 |
| IRIS_313-11829 | 0.000 | 0.000 | 0.000 | 0.000 | 1.000 | 0.000 | 0.000 |
| IRIS_313-11830 | 0.000 | 0.000 | 0.000 | 0.378 | 0.000 | 0.000 | 0.622 |
| IRIS_313-11831 | 0.000 | 0.621 | 0.000 | 0.000 | 0.379 | 0.000 | 0.000 |
| IRIS_313-11832 | 0.000 | 0.810 | 0.000 | 0.000 | 0.190 | 0.000 | 0.000 |
| IRIS_313-11833 | 0.000 | 0.000 | 0.000 | 0.542 | 0.000 | 0.000 | 0.458 |
| IRIS_313-11834 | 0.003 | 0.742 | 0.022 | 0.000 | 0.232 | 0.000 | 0.000 |
| IRIS_313-11835 | 0.000 | 0.000 | 0.000 | 0.592 | 0.000 | 0.408 | 0.000 |
| IRIS_313-11836 | 0.000 | 0.000 | 0.000 | 1.000 | 0.000 | 0.000 | 0.000 |
| IRIS_313-11837 | 0.000 | 0.000 | 0.000 | 1.000 | 0.000 | 0.000 | 0.000 |
| IRIS_313-11838 | 0.000 | 0.001 | 0.000 | 0.975 | 0.005 | 0.000 | 0.020 |
| IRIS_313-11839 | 0.000 | 0.000 | 0.000 | 0.998 | 0.002 | 0.000 | 0.000 |
| IRIS_313-11840 | 0.000 | 0.000 | 0.000 | 0.481 | 0.000 | 0.000 | 0.519 |
| IRIS_313-11841 | 0.000 | 0.000 | 0.000 | 0.965 | 0.000 | 0.000 | 0.035 |
| IRIS_313-11842 | 0.000 | 0.000 | 0.000 | 0.515 | 0.000 | 0.000 | 0.485 |
| IRIS_313-11843 | 0.000 | 0.000 | 0.000 | 0.945 | 0.000 | 0.000 | 0.055 |
| IRIS_313-11844 | 0.000 | 0.000 | 0.000 | 0.774 | 0.000 | 0.133 | 0.093 |
| IRIS_313-11845 | 0.000 | 0.682 | 0.000 | 0.000 | 0.318 | 0.000 | 0.000 |
| IRIS_313-11846 | 0.000 | 0.769 | 0.000 | 0.000 | 0.222 | 0.009 | 0.000 |
| IRIS_313-11848 | 0.000 | 0.000 | 0.000 | 0.772 | 0.000 | 0.000 | 0.228 |
| IRIS_313-11849 | 0.000 | 0.018 | 0.001 | 0.707 | 0.000 | 0.000 | 0.274 |
| IRIS_313-11850 | 0.000 | 0.000 | 0.000 | 0.559 | 0.000 | 0.000 | 0.441 |
| IRIS_313-11851 | 0.000 | 0.961 | 0.000 | 0.000 | 0.039 | 0.000 | 0.000 |
| IRIS_313-11852 | 0.000 | 0.000 | 0.000 | 0.170 | 0.010 | 0.820 | 0.000 |
| IRIS_313-11853 | 0.000 | 0.000 | 0.000 | 0.206 | 0.000 | 0.794 | 0.000 |
| IRIS_313-11854 | 0.000 | 0.000 | 0.000 | 0.139 | 0.000 | 0.861 | 0.000 |
| IRIS_313-11855 | 0.000 | 0.000 | 0.000 | 0.042 | 0.000 | 0.837 | 0.121 |
| IRIS_313-11856 | 0.000 | 0.000 | 0.000 | 0.153 | 0.000 | 0.847 | 0.000 |
| IRIS_313-11857 | 0.000 | 0.000 | 0.000 | 0.162 | 0.000 | 0.838 | 0.000 |
| IRIS_313-11858 | 0.000 | 0.000 | 0.000 | 0.027 | 0.000 | 0.973 | 0.000 |
| IRIS_313-11859 | 0.000 | 0.000 | 0.000 | 0.000 | 0.000 | 1.000 | 0.000 |
| IRIS_313-11860 | 0.000 | 0.000 | 0.000 | 0.000 | 0.000 | 1.000 | 0.000 |
| IRIS_313-11861 | 0.000 | 0.000 | 0.000 | 0.083 | 0.000 | 0.917 | 0.000 |
| IRIS_313-11862 | 0.000 | 0.000 | 0.000 | 0.007 | 0.000 | 0.993 | 0.000 |
| IRIS_313-11863 | 0.000 | 0.000 | 0.000 | 0.157 | 0.000 | 0.843 | 0.000 |
| IRIS_313-11865 | 0.000 | 0.000 | 0.000 | 0.000 | 0.015 | 0.985 | 0.000 |
| IRIS_313-11866 | 0.000 | 0.000 | 0.000 | 0.000 | 0.000 | 1.000 | 0.000 |
| IRIS_313-11867 | 0.000 | 0.000 | 0.000 | 0.138 | 0.000 | 0.862 | 0.000 |
| IRIS_313-11868 | 0.000 | 0.000 | 0.000 | 0.165 | 0.000 | 0.835 | 0.000 |
| IRIS_313-11869 | 0.000 | 0.000 | 0.000 | 0.000 | 0.000 | 1.000 | 0.000 |
| IRIS_313-11870 | 0.000 | 0.000 | 0.000 | 0.000 | 0.000 | 1.000 | 0.000 |
| IRIS_313-11871 | 0.000 | 0.000 | 0.000 | 0.000 | 0.000 | 0.877 | 0.123 |
| IRIS_313-11872 | 0.000 | 0.000 | 0.023 | 0.072 | 0.000 | 0.851 | 0.055 |
| IRIS_313-11875 | 0.000 | 0.081 | 0.000 | 0.000 | 0.919 | 0.000 | 0.000 |
| IRIS_313-11876 | 0.000 | 0.061 | 0.000 | 0.143 | 0.084 | 0.703 | 0.010 |
| IRIS_313-11877 | 0.000 | 0.000 | 0.000 | 0.000 | 0.000 | 1.000 | 0.000 |
| IRIS_313-11878 | 0.000 | 0.000 | 0.000 | 0.102 | 0.000 | 0.898 | 0.000 |
| IRIS_313-11881 | 0.000 | 0.000 | 0.000 | 0.013 | 0.005 | 0.982 | 0.000 |
| IRIS_313-11882 | 0.000 | 0.000 | 0.000 | 0.000 | 0.000 | 1.000 | 0.000 |
| IRIS_313-11884 | 0.000 | 0.000 | 0.000 | 0.000 | 0.000 | 1.000 | 0.000 |
| IRIS_313-11885 | 0.000 | 0.000 | 0.000 | 0.000 | 0.000 | 1.000 | 0.000 |
| IRIS_313-11886 | 0.000 | 0.000 | 0.000 | 0.000 | 0.000 | 0.134 | 0.866 |
| IRIS_313-11887 | 0.000 | 0.000 | 0.000 | 0.199 | 0.000 | 0.083 | 0.718 |
| IRIS_313-11888 | 1.000 | 0.000 | 0.000 | 0.000 | 0.000 | 0.000 | 0.000 |
| IRIS_313-11889 | 0.122 | 0.003 | 0.000 | 0.000 | 0.000 | 0.238 | 0.637 |
| IRIS_313-11890 | 0.009 | 0.000 | 0.000 | 0.003 | 0.972 | 0.016 | 0.000 |
| IRIS_313-11892 | 0.000 | 0.735 | 0.000 | 0.000 | 0.260 | 0.005 | 0.000 |
| IRIS_313-11893 | 0.000 | 0.019 | 0.000 | 0.458 | 0.000 | 0.358 | 0.165 |
| IRIS_313-11894 | 0.000 | 0.002 | 0.000 | 0.569 | 0.008 | 0.367 | 0.054 |
| IRIS_313-11895 | 0.000 | 0.000 | 0.000 | 0.989 | 0.000 | 0.011 | 0.000 |
| IRIS_313-11896 | 0.000 | 0.000 | 0.000 | 0.966 | 0.000 | 0.000 | 0.034 |
| IRIS_313-11897 | 0.000 | 0.654 | 0.000 | 0.000 | 0.346 | 0.000 | 0.000 |
| IRIS_313-11898 | 0.000 | 0.000 | 0.000 | 0.890 | 0.000 | 0.110 | 0.000 |

| | | | | | | | |
|---|---|---|---|---|---|---|---|
| IRIS_313-11899 | 0.000 | 0.009 | 0.000 | 0.787 | 0.000 | 0.204 | 0.000 |
| IRIS_313-11900 | 0.000 | 0.580 | 0.000 | 0.000 | 0.420 | 0.000 | 0.000 |
| IRIS_313-11901 | 0.000 | 0.000 | 0.000 | 0.467 | 0.000 | 0.000 | 0.533 |
| IRIS_313-11902 | 0.000 | 0.000 | 0.000 | 0.508 | 0.000 | 0.000 | 0.492 |
| IRIS_313-11903 | 0.000 | 1.000 | 0.000 | 0.000 | 0.000 | 0.000 | 0.000 |
| IRIS_313-11904 | 0.000 | 0.000 | 0.000 | 0.347 | 0.000 | 0.000 | 0.653 |
| IRIS_313-11905 | 0.000 | 0.000 | 0.000 | 0.533 | 0.000 | 0.000 | 0.467 |
| IRIS_313-11906 | 0.000 | 0.986 | 0.000 | 0.000 | 0.000 | 0.014 | 0.000 |
| IRIS_313-11907 | 0.000 | 0.952 | 0.000 | 0.000 | 0.048 | 0.000 | 0.000 |
| IRIS_313-11908 | 0.000 | 0.000 | 0.022 | 0.000 | 0.978 | 0.000 | 0.000 |
| IRIS_313-11909 | 0.000 | 0.000 | 0.000 | 0.000 | 0.000 | 1.000 | 0.000 |
| IRIS_313-11910 | 0.017 | 0.000 | 0.000 | 0.457 | 0.000 | 0.503 | 0.023 |
| IRIS_313-11911 | 0.000 | 0.000 | 0.000 | 0.076 | 0.000 | 0.924 | 0.000 |
| IRIS_313-11912 | 0.000 | 0.000 | 0.000 | 0.521 | 0.000 | 0.000 | 0.479 |
| IRIS_313-11913 | 0.000 | 1.000 | 0.000 | 0.000 | 0.000 | 0.000 | 0.000 |
| IRIS_313-11914 | 0.288 | 0.000 | 0.014 | 0.698 | 0.000 | 0.000 | 0.000 |
| IRIS_313-11915 | 0.000 | 0.954 | 0.000 | 0.000 | 0.046 | 0.000 | 0.000 |
| IRIS_313-11916 | 0.152 | 0.018 | 0.014 | 0.718 | 0.000 | 0.097 | 0.000 |
| IRIS_313-11917 | 1.000 | 0.000 | 0.000 | 0.000 | 0.000 | 0.000 | 0.000 |
| IRIS_313-11918 | 0.061 | 0.000 | 0.000 | 0.939 | 0.000 | 0.000 | 0.000 |
| IRIS_313-11919 | 0.062 | 0.000 | 0.000 | 0.938 | 0.000 | 0.000 | 0.000 |
| IRIS_313-11920 | 0.000 | 0.000 | 0.000 | 0.975 | 0.000 | 0.025 | 0.000 |
| IRIS_313-11921 | 0.000 | 0.000 | 0.000 | 0.996 | 0.004 | 0.000 | 0.000 |
| IRIS_313-11922 | 0.000 | 0.472 | 0.000 | 0.000 | 0.528 | 0.000 | 0.000 |
| IRIS_313-11923 | 0.000 | 0.662 | 0.000 | 0.000 | 0.338 | 0.000 | 0.000 |
| IRIS_313-11924 | 0.000 | 0.533 | 0.000 | 0.000 | 0.467 | 0.000 | 0.000 |
| IRIS_313-11925 | 0.000 | 0.000 | 0.000 | 0.943 | 0.000 | 0.000 | 0.057 |
| IRIS_313-11926 | 0.000 | 0.666 | 0.000 | 0.000 | 0.334 | 0.000 | 0.000 |
| IRIS_313-11927 | 0.000 | 0.000 | 0.000 | 1.000 | 0.000 | 0.000 | 0.000 |
| IRIS_313-11928 | 0.000 | 1.000 | 0.000 | 0.000 | 0.000 | 0.000 | 0.000 |
| IRIS_313-11929 | 0.000 | 1.000 | 0.000 | 0.000 | 0.000 | 0.000 | 0.000 |
| IRIS_313-11930 | 0.081 | 0.224 | 0.000 | 0.000 | 0.000 | 0.000 | 0.695 |
| IRIS_313-11931 | 0.000 | 0.029 | 0.000 | 0.000 | 0.971 | 0.000 | 0.000 |
| IRIS_313-11932 | 0.186 | 0.000 | 0.020 | 0.794 | 0.000 | 0.000 | 0.000 |
| IRIS_313-11934 | 0.000 | 0.000 | 0.063 | 0.177 | 0.000 | 0.158 | 0.602 |
| IRIS_313-11935 | 0.000 | 0.000 | 0.000 | 1.000 | 0.000 | 0.000 | 0.000 |
| IRIS_313-11936 | 0.000 | 0.000 | 0.000 | 1.000 | 0.000 | 0.000 | 0.000 |
| IRIS_313-11937 | 0.009 | 0.000 | 0.000 | 0.054 | 0.000 | 0.511 | 0.427 |
| IRIS_313-11938 | 0.030 | 0.000 | 0.000 | 0.970 | 0.000 | 0.000 | 0.000 |
| IRIS_313-11939 | 0.000 | 0.000 | 0.000 | 1.000 | 0.000 | 0.000 | 0.000 |
| IRIS_313-11940 | 0.001 | 0.000 | 0.000 | 0.992 | 0.000 | 0.000 | 0.007 |
| IRIS_313-11941 | 0.000 | 0.000 | 0.000 | 0.000 | 0.000 | 0.000 | 1.000 |
| IRIS_313-11943 | 0.000 | 0.000 | 0.000 | 0.000 | 0.000 | 1.000 | 0.000 |
| IRIS_313-11944 | 0.000 | 0.000 | 0.000 | 0.000 | 0.000 | 1.000 | 0.000 |
| IRIS_313-11945 | 0.069 | 0.000 | 0.007 | 0.874 | 0.000 | 0.050 | 0.000 |
| IRIS_313-11946 | 0.000 | 1.000 | 0.000 | 0.000 | 0.000 | 0.000 | 0.000 |
| IRIS_313-11947 | 0.000 | 0.000 | 0.011 | 0.608 | 0.013 | 0.368 | 0.000 |
| IRIS_313-11948 | 0.000 | 0.000 | 0.000 | 0.052 | 0.000 | 0.948 | 0.000 |
| IRIS_313-11949 | 0.000 | 0.000 | 0.000 | 0.000 | 0.000 | 1.000 | 0.000 |
| IRIS_313-11950 | 0.000 | 0.000 | 0.000 | 0.110 | 0.000 | 0.890 | 0.000 |
| IRIS_313-11951 | 0.000 | 0.000 | 0.000 | 0.000 | 0.000 | 0.966 | 0.034 |
| IRIS_313-11952 | 0.000 | 0.000 | 0.000 | 0.003 | 0.004 | 0.993 | 0.000 |
| IRIS_313-11953 | 0.000 | 0.000 | 0.000 | 0.175 | 0.039 | 0.785 | 0.000 |
| IRIS_313-11954 | 0.000 | 0.000 | 0.000 | 0.253 | 0.000 | 0.747 | 0.000 |
| IRIS_313-11955 | 0.000 | 0.000 | 0.000 | 0.172 | 0.000 | 0.828 | 0.000 |
| IRIS_313-11956 | 0.084 | 0.277 | 0.138 | 0.000 | 0.176 | 0.325 | 0.000 |
| IRIS_313-11957 | 0.021 | 0.000 | 0.000 | 0.603 | 0.000 | 0.376 | 0.000 |
| IRIS_313-11958 | 0.043 | 0.000 | 0.000 | 0.874 | 0.000 | 0.000 | 0.083 |
| IRIS_313-11959 | 0.005 | 0.016 | 0.000 | 0.877 | 0.004 | 0.000 | 0.097 |
| IRIS_313-11960 | 0.000 | 0.000 | 0.000 | 0.040 | 0.000 | 0.086 | 0.875 |
| IRIS_313-11961 | 0.000 | 0.000 | 0.000 | 1.000 | 0.000 | 0.000 | 0.000 |
| IRIS_313-11962 | 0.000 | 0.000 | 0.000 | 1.000 | 0.000 | 0.000 | 0.000 |
| IRIS_313-11963 | 1.000 | 0.000 | 0.000 | 0.000 | 0.000 | 0.000 | 0.000 |
| IRIS_313-11964 | 0.031 | 0.000 | 0.000 | 0.918 | 0.000 | 0.052 | 0.000 |
| IRIS_313-11965 | 0.000 | 0.000 | 0.000 | 0.000 | 0.000 | 1.000 | 0.000 |
| IRIS_313-11966 | 0.000 | 0.000 | 0.000 | 0.056 | 0.000 | 0.944 | 0.000 |
| IRIS_313-11967 | 0.000 | 0.000 | 0.000 | 0.596 | 0.033 | 0.372 | 0.000 |

| | | | | | | | |
|---|---|---|---|---|---|---|---|
| IRIS_313-11968 | 0.000 | 0.000 | 0.000 | 0.000 | 0.000 | 1.000 | 0.000 |
| IRIS_313-11969 | 0.082 | 0.000 | 0.000 | 0.918 | 0.000 | 0.000 | 0.000 |
| IRIS_313-11970 | 0.000 | 1.000 | 0.000 | 0.000 | 0.000 | 0.000 | 0.000 |
| IRIS_313-11971 | 0.000 | 0.984 | 0.000 | 0.000 | 0.000 | 0.000 | 0.016 |
| IRIS_313-11973 | 0.000 | 0.000 | 0.000 | 0.000 | 1.000 | 0.000 | 0.000 |
| IRIS_313-11974 | 0.000 | 0.000 | 0.000 | 0.227 | 0.000 | 0.000 | 0.773 |
| IRIS_313-11975 | 0.034 | 0.000 | 0.000 | 0.966 | 0.000 | 0.000 | 0.000 |
| IRIS_313-11976 | 0.000 | 0.000 | 0.000 | 0.370 | 0.000 | 0.000 | 0.630 |
| IRIS_313-11977 | 0.026 | 0.000 | 0.000 | 0.972 | 0.000 | 0.000 | 0.002 |
| IRIS_313-11978 | 0.000 | 0.004 | 0.000 | 0.000 | 0.000 | 0.127 | 0.870 |
| IRIS_313-11979 | 0.029 | 0.000 | 0.000 | 0.000 | 0.000 | 0.001 | 0.969 |
| IRIS_313-11980 | 0.000 | 0.000 | 0.000 | 0.000 | 0.000 | 1.000 | 0.000 |
| IRIS_313-11981 | 0.000 | 0.000 | 0.000 | 0.000 | 1.000 | 0.000 | 0.000 |
| IRIS_313-11982 | 1.000 | 0.000 | 0.000 | 0.000 | 0.000 | 0.000 | 0.000 |
| IRIS_313-11983 | 0.000 | 0.000 | 0.000 | 1.000 | 0.000 | 0.000 | 0.000 |
| IRIS_313-11984 | 0.000 | 0.000 | 0.032 | 0.000 | 0.968 | 0.000 | 0.000 |
| IRIS_313-11985 | 0.011 | 0.011 | 0.000 | 0.625 | 0.005 | 0.349 | 0.000 |
| IRIS_313-11986 | 0.000 | 0.000 | 0.000 | 0.000 | 0.000 | 1.000 | 0.000 |
| IRIS_313-11987 | 0.000 | 0.022 | 0.000 | 0.000 | 0.000 | 0.066 | 0.911 |
| IRIS_313-11988 | 0.016 | 0.000 | 0.000 | 0.983 | 0.000 | 0.000 | 0.000 |
| IRIS_313-11989 | 0.000 | 0.000 | 0.000 | 0.583 | 0.000 | 0.000 | 0.417 |
| IRIS_313-11990 | 0.000 | 0.000 | 0.000 | 0.000 | 0.014 | 0.109 | 0.878 |
| IRIS_313-11991 | 0.000 | 0.000 | 0.000 | 1.000 | 0.000 | 0.000 | 0.000 |
| IRIS_313-11992 | 0.000 | 0.000 | 0.000 | 1.000 | 0.000 | 0.000 | 0.000 |
| IRIS_313-11993 | 0.049 | 0.000 | 0.000 | 0.882 | 0.000 | 0.069 | 0.000 |
| IRIS_313-11994 | 0.000 | 1.000 | 0.000 | 0.000 | 0.000 | 0.000 | 0.000 |
| IRIS_313-11995 | 0.000 | 1.000 | 0.000 | 0.000 | 0.000 | 0.000 | 0.000 |
| IRIS_313-11996 | 0.000 | 0.009 | 0.000 | 0.745 | 0.012 | 0.185 | 0.048 |
| IRIS_313-11997 | 0.000 | 0.000 | 0.000 | 0.997 | 0.003 | 0.000 | 0.000 |
| IRIS_313-11998 | 0.000 | 0.000 | 0.000 | 1.000 | 0.000 | 0.000 | 0.000 |
| IRIS_313-11999 | 0.000 | 0.000 | 0.000 | 1.000 | 0.000 | 0.000 | 0.000 |
| IRIS_313-12000 | 0.000 | 0.000 | 0.000 | 1.000 | 0.000 | 0.000 | 0.000 |
| IRIS_313-12002 | 0.950 | 0.004 | 0.045 | 0.000 | 0.000 | 0.000 | 0.000 |
| IRIS_313-12003 | 0.000 | 0.000 | 0.000 | 0.000 | 1.000 | 0.000 | 0.000 |
| IRIS_313-12004 | 0.150 | 0.641 | 0.000 | 0.209 | 0.000 | 0.000 | 0.000 |
| IRIS_313-12005 | 0.000 | 0.000 | 0.000 | 0.462 | 0.000 | 0.000 | 0.538 |
| IRIS_313-12006 | 0.006 | 0.980 | 0.000 | 0.001 | 0.000 | 0.001 | 0.012 |
| IRIS_313-12007 | 0.000 | 0.000 | 0.030 | 0.572 | 0.000 | 0.394 | 0.004 |
| IRIS_313-12009 | 0.000 | 0.000 | 0.000 | 0.100 | 0.000 | 0.900 | 0.000 |
| IRIS_313-12010 | 0.000 | 0.000 | 0.000 | 0.000 | 0.000 | 1.000 | 0.000 |
| IRIS_313-12011 | 0.000 | 0.000 | 0.019 | 0.000 | 0.027 | 0.663 | 0.291 |
| IRIS_313-12012 | 0.000 | 0.000 | 0.000 | 0.070 | 0.000 | 0.930 | 0.000 |
| IRIS_313-12013 | 0.128 | 0.000 | 0.000 | 0.872 | 0.000 | 0.000 | 0.000 |
| IRIS_313-12014 | 0.000 | 0.001 | 0.000 | 0.997 | 0.002 | 0.000 | 0.000 |
| IRIS_313-12015 | 0.112 | 0.000 | 0.000 | 0.888 | 0.000 | 0.000 | 0.000 |
| IRIS_313-12016 | 0.015 | 0.055 | 0.000 | 0.211 | 0.036 | 0.683 | 0.000 |
| IRIS_313-12017 | 0.000 | 0.000 | 0.000 | 0.772 | 0.000 | 0.228 | 0.000 |
| IRIS_313-12018 | 0.000 | 1.000 | 0.000 | 0.000 | 0.000 | 0.000 | 0.000 |
| IRIS_313-12021 | 0.000 | 1.000 | 0.000 | 0.000 | 0.000 | 0.000 | 0.000 |
| IRIS_313-12024 | 0.000 | 0.000 | 0.000 | 0.000 | 0.000 | 0.000 | 1.000 |
| IRIS_313-12027 | 0.000 | 0.074 | 0.000 | 0.622 | 0.001 | 0.000 | 0.302 |
| IRIS_313-12028 | 0.000 | 0.000 | 0.000 | 0.571 | 0.000 | 0.429 | 0.000 |
| IRIS_313-12029 | 0.000 | 0.652 | 0.000 | 0.000 | 0.348 | 0.000 | 0.000 |
| IRIS_313-12030 | 0.000 | 0.668 | 0.000 | 0.129 | 0.000 | 0.000 | 0.203 |
| IRIS_313-12031 | 0.000 | 0.957 | 0.000 | 0.000 | 0.043 | 0.000 | 0.000 |
| IRIS_313-12033 | 0.000 | 0.000 | 0.000 | 0.000 | 0.000 | 1.000 | 0.000 |
| IRIS_313-12036 | 0.000 | 0.000 | 0.000 | 1.000 | 0.000 | 0.000 | 0.000 |
| IRIS_313-12037 | 0.000 | 0.000 | 0.000 | 1.000 | 0.000 | 0.000 | 0.000 |
| IRIS_313-12038 | 0.000 | 0.000 | 0.000 | 0.969 | 0.000 | 0.031 | 0.000 |
| IRIS_313-12039 | 0.000 | 0.000 | 0.000 | 0.998 | 0.000 | 0.000 | 0.002 |
| IRIS_313-12040 | 0.000 | 0.000 | 0.000 | 0.995 | 0.000 | 0.000 | 0.005 |
| IRIS_313-12041 | 0.000 | 0.000 | 0.000 | 1.000 | 0.000 | 0.000 | 0.000 |
| IRIS_313-12042 | 0.000 | 0.000 | 0.000 | 0.968 | 0.000 | 0.032 | 0.000 |
| IRIS_313-12043 | 0.000 | 0.000 | 0.000 | 0.923 | 0.000 | 0.000 | 0.077 |
| IRIS_313-12044 | 0.000 | 0.000 | 0.000 | 1.000 | 0.000 | 0.000 | 0.000 |
| IRIS_313-12045 | 0.000 | 1.000 | 0.000 | 0.000 | 0.000 | 0.000 | 0.000 |
| IRIS_313-12046 | 0.000 | 0.000 | 0.000 | 0.546 | 0.000 | 0.000 | 0.454 |

| | | | | | | | |
|---|---|---|---|---|---|---|---|
| IRIS_313-12047 | 0.000 | 0.008 | 0.000 | 0.641 | 0.000 | 0.000 | 0.350 |
| IRIS_313-12048 | 0.000 | 0.000 | 0.000 | 0.654 | 0.000 | 0.000 | 0.346 |
| IRIS_313-12049 | 0.000 | 0.000 | 0.000 | 0.935 | 0.000 | 0.065 | 0.000 |
| IRIS_313-12050 | 0.000 | 0.000 | 0.002 | 0.557 | 0.000 | 0.441 | 0.000 |
| IRIS_313-12051 | 0.000 | 1.000 | 0.000 | 0.000 | 0.000 | 0.000 | 0.000 |
| IRIS_313-12052 | 0.237 | 0.000 | 0.021 | 0.594 | 0.004 | 0.144 | 0.000 |
| IRIS_313-12053 | 0.000 | 0.000 | 0.000 | 0.345 | 0.000 | 0.000 | 0.655 |
| IRIS_313-12054 | 0.000 | 0.000 | 0.000 | 0.000 | 0.970 | 0.030 | 0.000 |
| IRIS_313-12055 | 0.923 | 0.000 | 0.000 | 0.000 | 0.000 | 0.000 | 0.077 |
| IRIS_313-12057 | 0.000 | 0.000 | 0.000 | 0.282 | 0.068 | 0.650 | 0.000 |
| IRIS_313-12058 | 0.000 | 0.000 | 0.000 | 1.000 | 0.000 | 0.000 | 0.000 |
| IRIS_313-12059 | 0.000 | 0.000 | 0.000 | 0.000 | 1.000 | 0.000 | 0.000 |
| IRIS_313-12060 | 0.000 | 0.130 | 0.000 | 0.000 | 0.870 | 0.000 | 0.000 |
| IRIS_313-12061 | 0.000 | 0.000 | 0.000 | 0.000 | 1.000 | 0.000 | 0.000 |
| IRIS_313-12063 | 0.000 | 0.608 | 0.000 | 0.000 | 0.392 | 0.000 | 0.000 |
| IRIS_313-12065 | 0.538 | 0.000 | 0.000 | 0.447 | 0.000 | 0.015 | 0.000 |
| IRIS_313-12066 | 0.000 | 0.000 | 0.000 | 1.000 | 0.000 | 0.000 | 0.000 |
| IRIS_313-12067 | 0.034 | 0.008 | 0.000 | 0.314 | 0.000 | 0.000 | 0.644 |
| IRIS_313-12068 | 0.000 | 0.975 | 0.000 | 0.000 | 0.007 | 0.018 | 0.000 |
| IRIS_313-12069 | 0.000 | 1.000 | 0.000 | 0.000 | 0.000 | 0.000 | 0.000 |
| IRIS_313-12070 | 0.000 | 1.000 | 0.000 | 0.000 | 0.000 | 0.000 | 0.000 |
| IRIS_313-12071 | 0.000 | 0.569 | 0.000 | 0.000 | 0.431 | 0.000 | 0.000 |
| IRIS_313-12072 | 0.000 | 0.000 | 0.000 | 0.566 | 0.000 | 0.434 | 0.000 |
| IRIS_313-12073 | 0.000 | 1.000 | 0.000 | 0.000 | 0.000 | 0.000 | 0.000 |
| IRIS_313-12074 | 0.000 | 0.135 | 0.801 | 0.000 | 0.064 | 0.000 | 0.000 |
| IRIS_313-12076 | 0.000 | 0.583 | 0.000 | 0.000 | 0.417 | 0.000 | 0.000 |
| IRIS_313-12077 | 0.000 | 0.599 | 0.000 | 0.000 | 0.401 | 0.000 | 0.000 |
| IRIS_313-12078 | 0.003 | 0.000 | 0.000 | 0.606 | 0.000 | 0.390 | 0.000 |
| IRIS_313-12079 | 0.000 | 0.000 | 0.000 | 0.000 | 0.000 | 0.000 | 1.000 |
| IRIS_313-12080 | 0.000 | 0.000 | 0.000 | 0.000 | 0.000 | 0.000 | 1.000 |
| IRIS_313-12081 | 0.000 | 0.000 | 0.000 | 0.332 | 0.000 | 0.000 | 0.668 |
| IRIS_313-12082 | 0.174 | 0.000 | 0.045 | 0.781 | 0.000 | 0.000 | 0.000 |
| IRIS_313-12083 | 0.207 | 0.000 | 0.000 | 0.789 | 0.004 | 0.000 | 0.000 |
| IRIS_313-12093 | 0.000 | 0.000 | 0.000 | 0.796 | 0.000 | 0.204 | 0.000 |
| IRIS_313-12094 | 0.000 | 0.000 | 1.000 | 0.000 | 0.000 | 0.000 | 0.000 |
| IRIS_313-12096 | 0.055 | 0.000 | 0.000 | 0.829 | 0.000 | 0.027 | 0.089 |
| IRIS_313-12097 | 0.000 | 0.000 | 0.000 | 1.000 | 0.000 | 0.000 | 0.000 |
| IRIS_313-12101 | 0.000 | 0.000 | 0.000 | 0.997 | 0.000 | 0.000 | 0.003 |
| IRIS_313-12102 | 0.000 | 0.000 | 0.000 | 0.971 | 0.000 | 0.029 | 0.000 |
| IRIS_313-12108 | 0.000 | 1.000 | 0.000 | 0.000 | 0.000 | 0.000 | 0.000 |
| IRIS_313-12109 | 0.000 | 0.000 | 0.000 | 0.977 | 0.000 | 0.023 | 0.000 |
| IRIS_313-12118 | 0.199 | 0.000 | 0.000 | 0.801 | 0.000 | 0.000 | 0.000 |
| IRIS_313-12121 | 0.000 | 0.000 | 0.000 | 1.000 | 0.000 | 0.000 | 0.000 |
| IRIS_313-12127 | 0.000 | 0.000 | 0.000 | 0.984 | 0.000 | 0.016 | 0.000 |
| IRIS_313-12128 | 0.000 | 0.000 | 0.000 | 1.000 | 0.000 | 0.000 | 0.000 |
| IRIS_313-12129 | 0.000 | 0.720 | 0.000 | 0.000 | 0.280 | 0.000 | 0.000 |
| IRIS_313-12130 | 0.000 | 0.000 | 0.000 | 0.908 | 0.000 | 0.088 | 0.004 |
| IRIS_313-12131 | 0.000 | 0.000 | 0.000 | 0.899 | 0.000 | 0.000 | 0.101 |
| IRIS_313-12133 | 0.000 | 0.003 | 0.000 | 0.997 | 0.000 | 0.000 | 0.000 |
| IRIS_313-12134 | 0.000 | 0.748 | 0.009 | 0.000 | 0.243 | 0.000 | 0.000 |
| IRIS_313-12135 | 0.000 | 0.000 | 0.000 | 0.156 | 0.000 | 0.000 | 0.844 |
| IRIS_313-12138 | 0.000 | 0.000 | 0.000 | 0.749 | 0.000 | 0.251 | 0.000 |
| IRIS_313-12139 | 1.000 | 0.000 | 0.000 | 0.000 | 0.000 | 0.000 | 0.000 |
| IRIS_313-12141 | 1.000 | 0.000 | 0.000 | 0.000 | 0.000 | 0.000 | 0.000 |
| IRIS_313-12142 | 0.000 | 0.000 | 0.000 | 1.000 | 0.000 | 0.000 | 0.000 |
| IRIS_313-12143 | 0.000 | 0.000 | 0.000 | 1.000 | 0.000 | 0.000 | 0.000 |
| IRIS_313-12144 | 0.000 | 1.000 | 0.000 | 0.000 | 0.000 | 0.000 | 0.000 |
| IRIS_313-12146 | 0.000 | 0.010 | 0.000 | 0.798 | 0.003 | 0.124 | 0.065 |
| IRIS_313-12147 | 0.000 | 0.000 | 0.000 | 0.956 | 0.000 | 0.000 | 0.044 |
| IRIS_313-12148 | 0.000 | 0.000 | 0.000 | 0.988 | 0.000 | 0.000 | 0.012 |
| IRIS_313-12151 | 0.000 | 0.000 | 0.000 | 1.000 | 0.000 | 0.000 | 0.000 |
| IRIS_313-12152 | 0.000 | 0.000 | 0.000 | 1.000 | 0.000 | 0.000 | 0.000 |
| IRIS_313-12161 | 0.000 | 0.000 | 0.000 | 1.000 | 0.000 | 0.000 | 0.000 |
| IRIS_313-12164 | 0.000 | 0.946 | 0.024 | 0.000 | 0.031 | 0.000 | 0.000 |
| IRIS_313-12180 | 0.000 | 0.000 | 0.000 | 0.744 | 0.000 | 0.256 | 0.000 |
| IRIS_313-12182 | 0.000 | 0.000 | 0.000 | 0.772 | 0.000 | 0.228 | 0.000 |
| IRIS_313-12183 | 0.962 | 0.000 | 0.000 | 0.000 | 0.000 | 0.038 | 0.000 |

| | | | | | | | |
|---|---|---|---|---|---|---|---|
| IRIS_313-12185 | 0.000 | 0.000 | 0.000 | 0.779 | 0.000 | 0.221 | 0.000 |
| IRIS_313-12186 | 0.013 | 0.062 | 0.000 | 0.200 | 0.044 | 0.681 | 0.000 |
| IRIS_313-12187 | 0.000 | 0.000 | 0.000 | 0.777 | 0.000 | 0.223 | 0.000 |
| IRIS_313-12188 | 0.000 | 0.000 | 0.000 | 1.000 | 0.000 | 0.000 | 0.000 |
| IRIS_313-12190 | 0.000 | 0.000 | 0.000 | 0.576 | 0.000 | 0.424 | 0.000 |
| IRIS_313-12193 | 0.000 | 0.000 | 0.000 | 0.997 | 0.000 | 0.003 | 0.000 |
| IRIS_313-12194 | 0.000 | 0.000 | 0.000 | 0.972 | 0.000 | 0.028 | 0.000 |
| IRIS_313-12200 | 0.000 | 0.560 | 0.000 | 0.000 | 0.440 | 0.000 | 0.000 |
| IRIS_313-12207 | 0.014 | 0.000 | 0.000 | 0.533 | 0.000 | 0.452 | 0.000 |
| IRIS_313-12210 | 0.000 | 0.003 | 0.000 | 0.639 | 0.001 | 0.338 | 0.019 |
| IRIS_313-12217 | 0.000 | 0.000 | 0.000 | 0.000 | 1.000 | 0.000 | 0.000 |
| IRIS_313-12220 | 0.000 | 0.000 | 0.000 | 0.869 | 0.000 | 0.131 | 0.000 |
| IRIS_313-12221 | 0.000 | 0.000 | 0.000 | 0.937 | 0.000 | 0.055 | 0.008 |
| IRIS_313-12222 | 0.000 | 0.000 | 0.000 | 0.876 | 0.000 | 0.124 | 0.000 |
| IRIS_313-12224 | 0.000 | 0.000 | 0.000 | 0.994 | 0.000 | 0.006 | 0.000 |
| IRIS_313-12225 | 0.000 | 0.000 | 0.000 | 1.000 | 0.000 | 0.000 | 0.000 |
| IRIS_313-12226 | 0.000 | 0.686 | 0.000 | 0.000 | 0.314 | 0.000 | 0.000 |
| IRIS_313-12227 | 0.000 | 0.691 | 0.000 | 0.000 | 0.309 | 0.000 | 0.000 |
| IRIS_313-12228 | 0.000 | 0.665 | 0.000 | 0.000 | 0.335 | 0.000 | 0.000 |
| IRIS_313-12229 | 0.000 | 0.000 | 0.000 | 0.819 | 0.000 | 0.181 | 0.000 |
| IRIS_313-12231 | 0.000 | 0.000 | 0.000 | 1.000 | 0.000 | 0.000 | 0.000 |
| IRIS_313-12232 | 0.001 | 0.000 | 0.000 | 0.000 | 0.000 | 0.545 | 0.455 |
| IRIS_313-12234 | 0.000 | 0.000 | 0.000 | 0.000 | 0.000 | 0.315 | 0.685 |
| IRIS_313-12236 | 0.000 | 0.000 | 0.000 | 0.071 | 0.000 | 0.929 | 0.000 |
| IRIS_313-12237 | 0.000 | 0.000 | 0.000 | 0.007 | 0.000 | 0.993 | 0.000 |
| IRIS_313-12240 | 0.000 | 0.000 | 0.000 | 0.422 | 0.000 | 0.000 | 0.578 |
| IRIS_313-12241 | 0.000 | 1.000 | 0.000 | 0.000 | 0.000 | 0.000 | 0.000 |
| IRIS_313-12242 | 0.000 | 0.597 | 0.000 | 0.000 | 0.403 | 0.000 | 0.000 |
| IRIS_313-12244 | 0.000 | 1.000 | 0.000 | 0.000 | 0.000 | 0.000 | 0.000 |
| IRIS_313-12246 | 0.000 | 0.000 | 0.000 | 0.990 | 0.000 | 0.010 | 0.000 |
| IRIS_313-12247 | 0.000 | 0.000 | 0.000 | 0.999 | 0.000 | 0.001 | 0.000 |
| IRIS_313-12249 | 0.000 | 0.000 | 0.000 | 0.881 | 0.000 | 0.042 | 0.076 |
| IRIS_313-12251 | 0.000 | 0.000 | 0.000 | 1.000 | 0.000 | 0.000 | 0.000 |
| IRIS_313-12252 | 0.000 | 0.743 | 0.008 | 0.000 | 0.249 | 0.000 | 0.000 |
| IRIS_313-12254 | 0.000 | 0.639 | 0.000 | 0.000 | 0.361 | 0.000 | 0.000 |
| IRIS_313-12257 | 0.000 | 0.612 | 0.000 | 0.000 | 0.388 | 0.000 | 0.000 |
| IRIS_313-12258 | 0.000 | 0.590 | 0.000 | 0.000 | 0.410 | 0.000 | 0.000 |
| IRIS_313-12259 | 0.000 | 0.000 | 0.000 | 0.988 | 0.000 | 0.012 | 0.000 |
| IRIS_313-12260 | 0.000 | 0.000 | 0.000 | 1.000 | 0.000 | 0.000 | 0.000 |
| IRIS_313-12261 | 0.000 | 0.000 | 0.000 | 1.000 | 0.000 | 0.000 | 0.000 |
| IRIS_313-12262 | 0.000 | 0.591 | 0.000 | 0.000 | 0.409 | 0.000 | 0.000 |
| IRIS_313-12263 | 0.000 | 0.000 | 0.000 | 1.000 | 0.000 | 0.000 | 0.000 |
| IRIS_313-12265 | 0.000 | 0.678 | 0.000 | 0.000 | 0.322 | 0.000 | 0.000 |
| IRIS_313-12266 | 0.007 | 0.651 | 0.000 | 0.000 | 0.341 | 0.000 | 0.000 |
| IRIS_313-12268 | 0.000 | 0.000 | 0.000 | 0.774 | 0.000 | 0.226 | 0.000 |
| IRIS_313-12269 | 0.000 | 0.000 | 0.000 | 0.765 | 0.000 | 0.092 | 0.144 |
| IRIS_313-12270 | 0.000 | 0.000 | 0.000 | 0.816 | 0.000 | 0.184 | 0.000 |
| IRIS_313-12271 | 0.000 | 1.000 | 0.000 | 0.000 | 0.000 | 0.000 | 0.000 |
| IRIS_313-12272 | 0.000 | 1.000 | 0.000 | 0.000 | 0.000 | 0.000 | 0.000 |
| IRIS_313-12273 | 0.000 | 0.000 | 0.000 | 0.010 | 0.000 | 0.904 | 0.086 |
| IRIS_313-12275 | 0.010 | 0.000 | 0.000 | 0.508 | 0.010 | 0.459 | 0.012 |
| IRIS_313-12278 | 0.175 | 0.000 | 0.006 | 0.819 | 0.000 | 0.000 | 0.000 |
| IRIS_313-12279 | 0.000 | 1.000 | 0.000 | 0.000 | 0.000 | 0.000 | 0.000 |
| IRIS_313-12280 | 0.000 | 0.000 | 0.000 | 0.998 | 0.002 | 0.000 | 0.000 |
| IRIS_313-12281 | 0.000 | 1.000 | 0.000 | 0.000 | 0.000 | 0.000 | 0.000 |
| IRIS_313-12282 | 0.079 | 0.000 | 0.000 | 0.742 | 0.000 | 0.001 | 0.178 |
| IRIS_313-12283 | 0.076 | 0.000 | 0.000 | 0.756 | 0.000 | 0.001 | 0.166 |
| IRIS_313-12284 | 0.311 | 0.000 | 0.000 | 0.689 | 0.000 | 0.000 | 0.000 |
| IRIS_313-12285 | 0.000 | 1.000 | 0.000 | 0.000 | 0.000 | 0.000 | 0.000 |
| IRIS_313-12286 | 0.000 | 0.000 | 0.000 | 0.959 | 0.000 | 0.000 | 0.041 |
| IRIS_313-12287 | 0.000 | 0.000 | 0.000 | 1.000 | 0.000 | 0.000 | 0.000 |
| IRIS_313-12288 | 0.000 | 0.000 | 0.000 | 0.964 | 0.000 | 0.036 | 0.000 |
| IRIS_313-12289 | 0.000 | 0.635 | 0.016 | 0.000 | 0.349 | 0.000 | 0.000 |
| IRIS_313-12290 | 0.000 | 0.014 | 0.000 | 0.815 | 0.000 | 0.161 | 0.010 |
| IRIS_313-12291 | 0.000 | 0.000 | 0.001 | 0.801 | 0.000 | 0.198 | 0.000 |
| IRIS_313-12292 | 0.010 | 0.002 | 0.000 | 0.813 | 0.002 | 0.022 | 0.150 |
| IRIS_313-12296 | 0.000 | 0.000 | 0.000 | 0.806 | 0.013 | 0.000 | 0.180 |

| | | | | | | | |
|---|---|---|---|---|---|---|---|
| IRIS_313-12297 | 0.000 | 0.010 | 0.014 | 0.605 | 0.000 | 0.371 | 0.000 |
| IRIS_313-12299 | 0.015 | 0.690 | 0.000 | 0.000 | 0.264 | 0.032 | 0.000 |
| IRIS_313-12300 | 0.000 | 0.000 | 0.000 | 0.568 | 0.000 | 0.432 | 0.000 |
| IRIS_313-12301 | 0.000 | 0.000 | 0.000 | 0.766 | 0.004 | 0.229 | 0.000 |
| IRIS_313-12302 | 0.000 | 0.000 | 0.000 | 0.893 | 0.000 | 0.107 | 0.000 |
| IRIS_313-12303 | 0.000 | 0.000 | 0.000 | 0.813 | 0.000 | 0.187 | 0.000 |
| IRIS_313-12305 | 0.000 | 0.002 | 0.000 | 0.929 | 0.001 | 0.068 | 0.000 |
| IRIS_313-12307 | 0.000 | 0.554 | 0.000 | 0.000 | 0.446 | 0.000 | 0.000 |
| IRIS_313-12308 | 0.000 | 0.001 | 0.000 | 0.997 | 0.001 | 0.000 | 0.000 |
| IRIS_313-12309 | 0.000 | 0.001 | 0.000 | 0.966 | 0.002 | 0.032 | 0.000 |
| IRIS_313-12311 | 0.000 | 0.541 | 0.000 | 0.000 | 0.459 | 0.000 | 0.000 |
| IRIS_313-12312 | 0.000 | 0.594 | 0.000 | 0.000 | 0.406 | 0.000 | 0.000 |
| IRIS_313-12313 | 0.000 | 0.668 | 0.000 | 0.000 | 0.332 | 0.000 | 0.000 |
| IRIS_313-12319 | 0.000 | 0.000 | 0.000 | 0.131 | 0.000 | 0.125 | 0.744 |
| IRIS_313-12321 | 0.000 | 0.751 | 0.000 | 0.000 | 0.249 | 0.000 | 0.000 |
| IRIS_313-12322 | 0.000 | 0.667 | 0.000 | 0.000 | 0.333 | 0.000 | 0.000 |
| IRIS_313-12323 | 0.000 | 0.617 | 0.000 | 0.000 | 0.383 | 0.000 | 0.000 |
| IRIS_313-12324 | 0.000 | 0.602 | 0.000 | 0.000 | 0.398 | 0.000 | 0.000 |
| IRIS_313-12325 | 0.021 | 0.000 | 0.000 | 0.978 | 0.001 | 0.000 | 0.000 |
| IRIS_313-12329 | 0.018 | 0.008 | 0.000 | 0.583 | 0.002 | 0.389 | 0.000 |
| IRIS_313-12330 | 0.000 | 0.635 | 0.000 | 0.000 | 0.365 | 0.000 | 0.000 |
| IRIS_313-12332 | 0.000 | 0.623 | 0.000 | 0.000 | 0.377 | 0.000 | 0.000 |
| IRIS_313-12334 | 0.000 | 0.000 | 0.000 | 0.964 | 0.000 | 0.002 | 0.035 |
| IRIS_313-12336 | 0.000 | 0.626 | 0.000 | 0.000 | 0.374 | 0.000 | 0.000 |
| IRIS_313-12337 | 0.000 | 0.590 | 0.000 | 0.000 | 0.410 | 0.000 | 0.000 |
| IRIS_313-12340 | 0.000 | 0.560 | 0.000 | 0.000 | 0.440 | 0.000 | 0.000 |
| IRIS_313-12341 | 0.000 | 0.598 | 0.000 | 0.000 | 0.402 | 0.000 | 0.000 |
| IRIS_313-12342 | 0.000 | 0.628 | 0.000 | 0.000 | 0.372 | 0.000 | 0.000 |
| IRIS_313-12344 | 0.000 | 0.616 | 0.000 | 0.000 | 0.384 | 0.000 | 0.000 |
| IRIS_313-12345 | 0.000 | 0.666 | 0.000 | 0.000 | 0.295 | 0.039 | 0.000 |
| IRIS_313-12346 | 0.000 | 0.651 | 0.000 | 0.000 | 0.349 | 0.000 | 0.000 |
| IRIS_313-12347 | 0.000 | 0.593 | 0.000 | 0.000 | 0.407 | 0.000 | 0.000 |
| IRIS_313-12348 | 0.000 | 0.621 | 0.000 | 0.000 | 0.379 | 0.000 | 0.000 |
| IRIS_313-12349 | 0.000 | 0.643 | 0.000 | 0.000 | 0.357 | 0.000 | 0.000 |
| IRIS_313-12350 | 0.000 | 0.644 | 0.000 | 0.000 | 0.356 | 0.000 | 0.000 |
| IRIS_313-12351 | 0.000 | 0.606 | 0.000 | 0.000 | 0.394 | 0.000 | 0.000 |
| IRIS_313-12352 | 0.000 | 0.603 | 0.000 | 0.000 | 0.397 | 0.000 | 0.000 |
| IRIS_313-12353 | 0.000 | 0.523 | 0.000 | 0.000 | 0.477 | 0.000 | 0.000 |
| IRIS_313-12354 | 0.000 | 0.000 | 0.000 | 0.000 | 0.000 | 0.216 | 0.784 |
| IRIS_313-12355 | 0.000 | 0.000 | 0.000 | 0.362 | 0.000 | 0.280 | 0.358 |
| IRIS_313-15896 | 0.067 | 0.000 | 0.000 | 0.000 | 0.000 | 0.125 | 0.808 |
| IRIS_313-15897 | 0.000 | 0.000 | 0.000 | 0.128 | 0.000 | 0.444 | 0.428 |
| IRIS_313-15898 | 0.000 | 0.000 | 0.000 | 0.465 | 0.000 | 0.000 | 0.535 |
| IRIS_313-15899 | 0.120 | 0.000 | 0.000 | 0.303 | 0.008 | 0.175 | 0.394 |
| IRIS_313-15900 | 0.000 | 0.000 | 0.000 | 0.399 | 0.000 | 0.039 | 0.562 |
| IRIS_313-15901 | 0.008 | 0.000 | 0.060 | 0.000 | 0.000 | 0.136 | 0.796 |
| IRIS_313-15902 | 0.000 | 0.000 | 0.000 | 0.000 | 0.023 | 0.236 | 0.741 |
| IRIS_313-15903 | 0.303 | 0.031 | 0.000 | 0.120 | 0.000 | 0.000 | 0.546 |
| IRIS_313-15904 | 0.000 | 0.000 | 0.000 | 0.000 | 1.000 | 0.000 | 0.000 |
| IRIS_313-15905 | 0.000 | 0.941 | 0.000 | 0.000 | 0.000 | 0.000 | 0.059 |
| IRIS_313-15906 | 0.000 | 0.000 | 0.000 | 0.000 | 0.123 | 0.175 | 0.702 |
| IRIS_313-15907 | 0.000 | 0.974 | 0.000 | 0.000 | 0.000 | 0.026 | 0.000 |
| IRIS_313-15908 | 1.000 | 0.000 | 0.000 | 0.000 | 0.000 | 0.000 | 0.000 |
| IRIS_313-15909 | 0.119 | 0.000 | 0.881 | 0.000 | 0.000 | 0.000 | 0.000 |
| IRIS_313-15910 | 0.000 | 0.973 | 0.000 | 0.000 | 0.000 | 0.027 | 0.000 |
| LGR | 0.044 | 0.000 | 0.000 | 0.257 | 0.698 | 0.000 | 0.000 |
| PB 1121 | 0.178 | 0.000 | 0.179 | 0.001 | 0.402 | 0.043 | 0.198 |
| SONASAL | 0.123 | 0.010 | 0.217 | 0.000 | 0.295 | 0.116 | 0.239 |
| BINDLI | 0.116 | 0.000 | 0.214 | 0.148 | 0.312 | 0.000 | 0.209 |
