## Supplemental Table 11 for "Genome-wide analysis of polymorphisms identified domestication-associated polymorphism desert carrying important rice grain size/weight QTL"

**Supplemental Table 11: Genome fraction estimation of aromatic group genotypes through fastStructure (K=3) meanQ analysis**

| | Gene pool | | |
|---|---|---|---|
| Accession number | *indica* | *japonica* | *aus* |
| CX104 | 0.000 | 0.582 | 0.418 |
| CX110 | 0.000 | 0.581 | 0.419 |
| CX112 | 0.000 | 0.549 | 0.451 |
| CX143 | 0.013 | 0.546 | 0.441 |
| CX149 | 0.000 | 0.435 | 0.565 |
| IRIS_313-8215 | 0.958 | 0.038 | 0.004 |
| IRIS_313-8326 | 0.000 | 0.620 | 0.380 |
| IRIS_313-8656 | 0.000 | 0.454 | 0.546 |
| IRIS_313-8664 | 0.024 | 0.936 | 0.039 |
| IRIS_313-8712 | 0.000 | 0.639 | 0.361 |
| IRIS_313-8747 | 0.000 | 0.561 | 0.439 |
| IRIS_313-8765 | 0.005 | 0.555 | 0.440 |
| IRIS_313-8813 | 0.000 | 0.560 | 0.440 |
| IRIS_313-8814 | 0.000 | 0.523 | 0.477 |
| IRIS_313-8923 | 0.000 | 1.000 | 0.000 |
| IRIS_313-9083 | 0.000 | 0.509 | 0.491 |
| IRIS_313-9170 | 0.017 | 0.624 | 0.359 |
| IRIS_313-9558 | 0.005 | 0.995 | 0.000 |
| IRIS_313-9682 | 0.020 | 0.615 | 0.365 |
| IRIS_313-9939 | 0.574 | 0.414 | 0.013 |
| IRIS_313-10016 | 1.000 | 0.000 | 0.000 |
| IRIS_313-10051 | 0.006 | 0.994 | 0.000 |
| IRIS_313-10258 | 0.000 | 1.000 | 0.000 |
| IRIS_313-10521 | 0.006 | 0.523 | 0.470 |
| IRIS_313-10670 | 0.000 | 0.610 | 0.390 |
| IRIS_313-10732 | 0.000 | 0.537 | 0.463 |
| IRIS_313-10851 | 0.022 | 0.558 | 0.420 |
| IRIS_313-10864 | 0.000 | 0.803 | 0.197 |
| IRIS_313-10868 | 0.000 | 0.562 | 0.438 |
| IRIS_313-10926 | 0.011 | 0.535 | 0.453 |
| IRIS_313-10933 | 0.010 | 0.610 | 0.381 |
| IRIS_313-11021 | 0.000 | 0.469 | 0.531 |
| IRIS_313-11022 | 0.000 | 0.439 | 0.561 |
| IRIS_313-11023 | 0.000 | 0.474 | 0.526 |
| IRIS_313-11026 | 0.000 | 0.511 | 0.489 |
| IRIS_313-11062 | 0.000 | 0.571 | 0.429 |
| IRIS_313-11066 | 0.000 | 0.552 | 0.448 |
| IRIS_313-11068 | 0.000 | 0.594 | 0.406 |
| IRIS_313-11069 | 0.000 | 0.549 | 0.451 |
| IRIS_313-11070 | 0.000 | 0.574 | 0.426 |
| IRIS_313-11218 | 0.000 | 0.565 | 0.435 |
| IRIS_313-11258 | 0.067 | 0.506 | 0.427 |
| IRIS_313-11259 | 0.000 | 0.673 | 0.327 |
| IRIS_313-11268 | 0.000 | 0.622 | 0.378 |
| IRIS_313-11289 | 0.000 | 0.616 | 0.384 |

| | | | |
|---|---|---|---|
| IRIS_313-11293 | 0.000 | 0.616 | 0.384 |
| IRIS_313-11297 | 0.000 | 0.799 | 0.201 |
| IRIS_313-11350 | 0.019 | 0.525 | 0.456 |
| IRIS_313-11352 | 0.102 | 0.511 | 0.387 |
| IRIS_313-11356 | 0.000 | 0.551 | 0.449 |
| IRIS_313-11362 | 0.000 | 0.532 | 0.468 |
| IRIS_313-11373 | 0.000 | 0.570 | 0.430 |
| IRIS_313-11401 | 0.000 | 0.620 | 0.380 |
| IRIS_313-11563 | 0.859 | 0.000 | 0.141 |
| IRIS_313-11566 | 0.912 | 0.000 | 0.088 |
| IRIS_313-11606 | 0.940 | 0.000 | 0.060 |
| IRIS_313-11625 | 0.060 | 0.436 | 0.504 |
| IRIS_313-11626 | 0.000 | 0.582 | 0.418 |
| IRIS_313-11629 | 0.000 | 0.000 | 1.000 |
| IRIS_313-11630 | 0.000 | 0.507 | 0.493 |
| IRIS_313-11695 | 0.978 | 0.022 | 0.000 |
| IRIS_313-11742 | 0.000 | 0.000 | 1.000 |
| IRIS_313-11764 | 1.000 | 0.000 | 0.000 |
| IRIS_313-11824 | 0.948 | 0.000 | 0.052 |
| IRIS_313-11825 | 0.000 | 0.536 | 0.464 |
| IRIS_313-12074 | 0.000 | 0.672 | 0.328 |
| IRIS_313-12094 | 0.000 | 0.534 | 0.466 |
| IRIS_313-15908 | 0.001 | 0.002 | 0.997 |
| PB 1121 | 0.194 | 0.501 | 0.305 |
| SONASAL | 0.256 | 0.404 | 0.340 |
| BINDLI | 0.257 | 0.409 | 0.334 |
