## Supplemental Table 24 for "Genome-wide analysis of polymorphisms identified domestication-associated polymorphism desert carrying important rice grain size/weight QTL"

**Supplemental Table 24: Distribution of Desert and Non-Desert genotypes for their origin country among 3004 rice accessions**

| Origin_country | Total_genotypes | Desert_genotypes | Non-Desert_genotypes |
|---|---|---|---|
| China | 481 | 452 | 29 |
| India | 439 | 333 | 106 |
| Indonesia | 248 | 205 | 43 |
| Philippines | 229 | 214 | 15 |
| Bangladesh | 186 | 73 | 113 |
| Thailand | 147 | 133 | 14 |
| Laos | 126 | 115 | 11 |
| Malaysia | 75 | 70 | 5 |
| Myanmar | 75 | 64 | 11 |
| Madagascar | 66 | 28 | 38 |
| Cambodia | 59 | 53 | 6 |
| Japan | 55 | 55 | 0 |
| Vietnam | 55 | 48 | 7 |
| Sri_Lanka | 54 | 36 | 18 |
| United_States | 51 | 49 | 2 |
| Nepal | 44 | 38 | 6 |
| Italy | 38 | 38 | 0 |
| Taiwan | 35 | 31 | 4 |
| No_Data | 34 | 31 | 3 |
| Pakistan | 34 | 20 | 14 |
| South_Korea | 34 | 33 | 1 |
| Brazil | 28 | 24 | 4 |
| Ivory_Coast | 26 | 25 | 1 |
| Colombia | 24 | 21 | 3 |
| Portugal | 24 | 24 | 0 |
| Senegal | 22 | 21 | 1 |
| Liberia | 20 | 18 | 2 |
| Bhutan | 19 | 18 | 1 |
| Sierra_Leone | 19 | 17 | 2 |
| Australia | 14 | 14 | 0 |
| Nigeria | 13 | 10 | 3 |
| Africa | 11 | 10 | 1 |
| Argentina | 11 | 11 | 0 |
| Bulgaria | 10 | 10 | 0 |
| France | 10 | 10 | 0 |
| Iran | 10 | 9 | 1 |
| Spain | 10 | 10 | 0 |
| Suriname | 9 | 9 | 0 |
| Burkina_Fasso | 8 | 8 | 0 |
| Egypt | 8 | 7 | 1 |
| Guinea | 8 | 7 | 1 |
| Kenya | 8 | 4 | 4 |
| Gambia | 7 | 6 | 1 |
| Hungary | 7 | 7 | 0 |
| Mali | 7 | 7 | 0 |
| North_Korea | 7 | 7 | 0 |

| | | | |
|---|---|---|---|
| Cuba | 6 | 6 | 0 |
| Mexico | 6 | 6 | 0 |
| Peru | 6 | 5 | 1 |
| Tanzania | 6 | 6 | 0 |
| Ghana | 5 | 5 | 0 |
| Ecuador | 4 | 3 | 1 |
| Greece | 4 | 4 | 0 |
| Guinea-Bissau | 4 | 4 | 0 |
| Venezuela | 4 | 3 | 1 |
| Burundi | 3 | 1 | 2 |
| Guatemala | 3 | 3 | 0 |
| Guyana | 3 | 2 | 1 |
| Romania | 3 | 3 | 0 |
| Brunei_Darussalam | 2 | 1 | 1 |
| Cameroon | 2 | 1 | 1 |
| Georgia | 2 | 2 | 0 |
| Russia | 2 | 2 | 0 |
| Uruguay | 2 | 2 | 0 |
| USSR | 2 | 2 | 0 |
| Zambia | 2 | 1 | 1 |
| Afghanistan | 1 | 0 | 1 |
| Albania | 1 | 1 | 0 |
| Belgium | 1 | 1 | 0 |
| Benin | 1 | 1 | 0 |
| Bolivia | 1 | 1 | 0 |
| Central_America | 1 | 1 | 0 |
| Chad | 1 | 1 | 0 |
| Chile | 1 | 1 | 0 |
| Dominican_Republic | 1 | 1 | 0 |
| Fiji | 1 | 1 | 0 |
| French_Guiana | 1 | 1 | 0 |
| Haiti | 1 | 1 | 0 |
| Hong_Kong | 1 | 0 | 1 |
| Korea | 1 | 1 | 0 |
| Malawi | 1 | 1 | 0 |
| Mozambique | 1 | 1 | 0 |
| Netherlands | 1 | 0 | 1 |
| New_Zealand | 1 | 1 | 0 |
| Nicaragua | 1 | 1 | 0 |
| Niger | 1 | 1 | 0 |
| Norway | 1 | 1 | 0 |
| Panama | 1 | 1 | 0 |
| Paraguay | 1 | 1 | 0 |
| Solomon_Islands | 1 | 1 | 0 |
| Soviet_Union | 1 | 1 | 0 |
| Turkey | 1 | 1 | 0 |
| Uganda | 1 | 1 | 0 |
| Zimbabwe | 1 | 0 | 1 |
