## Supplemental Table 25 for "Genome-wide analysis of polymorphisms identified domestication-associated polymorphism desert carrying important rice grain size/weight QTL"

**Supplemental Table 25: Details of Chr 5 desert region SNPs found to be significantly associated (FDR adjusted p-value < 0.05) with grain size/weight and other domesticated related traits across the 2270 rice accessions.**

Grain_length_mm

| SNP | Chr | Position | maf | nobs | $R^2$ with SNP | FDR_Adjusted_P-values | allele |
|---|---|---|---|---|---|---|---|
| SNP_A.K_5_8172473 | 5 | 8172473 | 0.015 | 2266 | 0.117722466 | 0.006859582 | G/A |
| SNP_A.K_5_8541630 | 5 | 8541630 | 0.026 | 2266 | 0.117536146 | 0.006859582 | G/A |
| SNP_A.K_5_10795164 | 5 | 10795164 | 0.024 | 2266 | 0.117600003 | 0.006859582 | T/C |
| SNP_A.K_5_12779064 | 5 | 12779064 | 0.018 | 2266 | 0.117820167 | 0.006859582 | C/T |
| SNP_A.K_5_13003372 | 5 | 13003372 | 0.022 | 2266 | 0.117462282 | 0.006859582 | T/C |
| SNP_A.K_5_8027721 | 5 | 8027721 | 0.022 | 2266 | 0.116998549 | 0.010455834 | G/A |
| SNP_A.K_5_8449552 | 5 | 8449552 | 0.017 | 2266 | 0.116701162 | 0.013209492 | C/T |
| SNP_A.K_5_10341545 | 5 | 10341545 | 0.023 | 2266 | 0.116568351 | 0.013747145 | C/T |
| SNP_A.K_5_12911427 | 5 | 12911427 | 0.242 | 2266 | 0.11636128 | 0.016017461 | G/A |
| SNP_A.K_5_12099957 | 5 | 12099957 | 0.021 | 2266 | 0.116131741 | 0.017895912 | A/T |
| SNP_A.K_5_12827141 | 5 | 12827141 | 0.024 | 2266 | 0.116123168 | 0.017895912 | G/C |
| SNP_A.K_5_13254319 | 5 | 13254319 | 0.036 | 2266 | 0.115751574 | 0.026698136 | G/A |
| SNP_A.K_5_11666786 | 5 | 11666786 | 0.023 | 2266 | 0.115624128 | 0.029131692 | G/A |
| SNP_A.K_5_10906284 | 5 | 10906284 | 0.024 | 2266 | 0.115450645 | 0.029728112 | T/C |
| SNP_A.K_5_13231852 | 5 | 13231852 | 0.031 | 2266 | 0.115484892 | 0.029728112 | A/T |
| SNP_A.K_5_13316066 | 5 | 13316066 | 0.023 | 2266 | 0.115515065 | 0.029728112 | C/T |
| SNP_A.K_5_12711437 | 5 | 12711437 | 0.024 | 2266 | 0.115387033 | 0.030419876 | T/A |
| SNP_A.K_5_8933914 | 5 | 8933914 | 0.019 | 2266 | 0.115283327 | 0.032928717 | G/A |
| SNP_A.K_5_9056325 | 5 | 9056325 | 0.023 | 2266 | 0.115120384 | 0.037858172 | T/C |
| SNP_A.K_5_11765957 | 5 | 11765957 | 0.015 | 2266 | 0.115097338 | 0.037858172 | G/C |
| SNP_A.K_5_11763085 | 5 | 11763085 | 0.017 | 2266 | 0.114956561 | 0.043405843 | A/T |
| SNP_A.K_5_10807497 | 5 | 10807497 | 0.482 | 2266 | 0.114861601 | 0.046961379 | G/A |
| SNP_A.K_5_8236608 | 5 | 8236608 | 0.141 | 2266 | 0.114760143 | 0.047333154 | G/A |
| SNP_A.K_5_11765307 | 5 | 11765307 | 0.015 | 2266 | 0.114710678 | 0.047333154 | C/T |
| SNP_A.K_5_12908239 | 5 | 12908239 | 0.117 | 2266 | 0.1147005 | 0.047333154 | C/T |
| SNP_A.K_5_13767065 | 5 | 13767065 | 0.029 | 2266 | 0.114741194 | 0.047333154 | G/A |
| SNP_A.K_5_13771364 | 5 | 13771364 | 0.031 | 2266 | 0.114708032 | 0.047333154 | C/T |
| SNP_A.K_5_13589891 | 5 | 13589891 | 0.026 | 2266 | 0.114654559 | 0.048498675 | G/T |
| SNP_A.K_5_12873567 | 5 | 12873567 | 0.196 | 2266 | 0.114457672 | 0.058726175 | C/T |
| SNP_A.K_5_12914772 | 5 | 12914772 | 0.211 | 2266 | 0.114468012 | 0.058726175 | A/G |

Grain_width_mm

| SNP | Chr | Position | maf | nobs | $R^2$ with SNP | FDR_Adjusted_P-values | allele |
|---|---|---|---|---|---|---|---|
| SNP_A.K_5_8024799 | 5 | 8024799 | 0.017 | 2266 | 0.279721988 | 7.87148E-05 | C/T |
| SNP_A.K_5_8382703 | 5 | 8382703 | 0.015 | 2266 | 0.279992873 | 7.87148E-05 | C/T |
| SNP_A.K_5_11691286 | 5 | 11691286 | 0.295 | 2266 | 0.279402773 | 7.87148E-05 | T/C |
| SNP_A.K_5_12349684 | 5 | 12349684 | 0.246 | 2266 | 0.279337917 | 7.87148E-05 | A/G |
| SNP_A.K_5_8150444 | 5 | 8150444 | 0.095 | 2266 | 0.278560822 | 0.000212991 | G/A |
| SNP_A.K_5_8097699 | 5 | 8097699 | 0.017 | 2266 | 0.278301485 | 0.000266776 | G/A |
| SNP_A.K_5_8287113 | 5 | 8287113 | 0.018 | 2266 | 0.276556869 | 0.003588464 | A/T |
| SNP_A.K_5_12783182 | 5 | 12783182 | 0.014 | 2266 | 0.27612532 | 0.006226838 | A/T |
| SNP_A.K_5_12350390 | 5 | 12350390 | 0.023 | 2266 | 0.275811467 | 0.009116328 | G/A |
| SNP_A.K_5_13980526 | 5 | 13980526 | 0.022 | 2266 | 0.275541016 | 0.012621875 | G/C |
| SNP_A.K_5_8196979 | 5 | 8196979 | 0.016 | 2266 | 0.275157521 | 0.016640011 | G/A |
| SNP_A.K_5_8360226 | 5 | 8360226 | 0.018 | 2266 | 0.275302059 | 0.016640011 | C/T |
| SNP_A.K_5_8903375 | 5 | 8903375 | 0.021 | 2266 | 0.275157004 | 0.016640011 | C/T |
| SNP_A.K_5_12010348 | 5 | 12010348 | 0.035 | 2266 | 0.275196866 | 0.016640011 | G/A |
| SNP_A.K_5_13691228 | 5 | 13691228 | 0.016 | 2266 | 0.275065061 | 0.017989385 | C/T |
| SNP_A.K_5_12089202 | 5 | 12089202 | 0.016 | 2266 | 0.27500901 | 0.018446661 | C/T |
| SNP_A.K_5_8400748 | 5 | 8400748 | 0.034 | 2266 | 0.274873635 | 0.019767291 | C/T |
| SNP_A.K_5_9052758 | 5 | 9052758 | 0.013 | 2266 | 0.274624845 | 0.019767291 | C/T |
| SNP_A.K_5_9099952 | 5 | 9099952 | 0.015 | 2266 | 0.274732862 | 0.019767291 | C/T |
| SNP_A.K_5_9393809 | 5 | 9393809 | 0.120 | 2266 | 0.274616482 | 0.019767291 | G/A |
| SNP_A.K_5_10122094 | 5 | 10122094 | 0.014 | 2266 | 0.27467834 | 0.019767291 | T/C |
| SNP_A.K_5_10276942 | 5 | 10276942 | 0.178 | 2266 | 0.274630522 | 0.019767291 | G/A |
| SNP_A.K_5_12628190 | 5 | 12628190 | 0.015 | 2266 | 0.274818379 | 0.019767291 | G/A |
| SNP_A.K_5_12705831 | 5 | 12705831 | 0.150 | 2266 | 0.274870969 | 0.019767291 | G/T |
| SNP_A.K_5_12706192 | 5 | 12706192 | 0.143 | 2266 | 0.274761446 | 0.019767291 | T/A |
| SNP_A.K_5_12931498 | 5 | 12931498 | 0.015 | 2266 | 0.274648966 | 0.019767291 | T/A |
| SNP_A.K_5_13509846 | 5 | 13509846 | 0.015 | 2266 | 0.274637039 | 0.019767291 | T/A |

| | | | | | | | |
|---|---|---|---|---|---|---|---|
| SNP_A.K_5_13553961 | 5 | 13553961 | 0.023 | 2266 | 0.274716821 | 0.019767291 | G/T |
| SNP_A.K_5_12706139 | 5 | 12706139 | 0.137 | 2266 | 0.274560541 | 0.02087796 | T/C |
| SNP_A.K_5_9360148 | 5 | 9360148 | 0.017 | 2266 | 0.274400329 | 0.026103492 | T/A |
| SNP_A.K_5_12355842 | 5 | 12355842 | 0.028 | 2266 | 0.274338924 | 0.027881748 | C/T |
| SNP_A.K_5_9723039 | 5 | 9723039 | 0.015 | 2266 | 0.274296596 | 0.028036674 | T/C |
| SNP_A.K_5_11691721 | 5 | 11691721 | 0.288 | 2266 | 0.274297577 | 0.028036674 | A/G |
| SNP_A.K_5_10343410 | 5 | 10343410 | 0.017 | 2266 | 0.274178219 | 0.02915295 | G/T |
| SNP_A.K_5_10358047 | 5 | 10358047 | 0.019 | 2266 | 0.274212756 | 0.02915295 | A/G |
| SNP_A.K_5_11428687 | 5 | 11428687 | 0.023 | 2266 | 0.274186067 | 0.02915295 | C/T |
| SNP_A.K_5_11430717 | 5 | 11430717 | 0.014 | 2266 | 0.274181946 | 0.02915295 | G/A |
| SNP_A.K_5_11740554 | 5 | 11740554 | 0.014 | 2266 | 0.274190674 | 0.02915295 | C/T |
| SNP_A.K_5_12876839 | 5 | 12876839 | 0.014 | 2266 | 0.274168513 | 0.02915295 | G/A |
| SNP_A.K_5_8848238 | 5 | 8848238 | 0.015 | 2266 | 0.274060656 | 0.032302745 | G/A |
| SNP_A.K_5_12333962 | 5 | 12333962 | 0.014 | 2266 | 0.274050166 | 0.032302745 | C/T |
| SNP_A.K_5_12706191 | 5 | 12706191 | 0.142 | 2266 | 0.274044206 | 0.032302745 | G/A |
| SNP_A.K_5_12873886 | 5 | 12873886 | 0.016 | 2266 | 0.274062167 | 0.032302745 | C/T |
| SNP_A.K_5_9492708 | 5 | 9492708 | 0.017 | 2266 | 0.273981968 | 0.034900392 | A/G |
| SNP_A.K_5_13656006 | 5 | 13656006 | 0.014 | 2266 | 0.273932749 | 0.03694434 | T/C |
| SNP_A.K_5_12705372 | 5 | 12705372 | 0.149 | 2266 | 0.273917263 | 0.037055544 | T/C |
| SNP_A.K_5_9054962 | 5 | 9054962 | 0.021 | 2266 | 0.273779037 | 0.045335205 | C/T |
| SNP_A.K_5_8027721 | 5 | 8027721 | 0.022 | 2266 | 0.273734762 | 0.047682997 | G/A |
| SNP_A.K_5_11435845 | 5 | 11435845 | 0.014 | 2266 | 0.273673256 | 0.04985135 | C/T |
| SNP_A.K_5_11737955 | 5 | 11737955 | 0.029 | 2266 | 0.273668988 | 0.04985135 | G/A |
| SNP_A.K_5_11739970 | 5 | 11739970 | 0.024 | 2266 | 0.273647952 | 0.04985135 | G/A |
| SNP_A.K_5_12653598 | 5 | 12653598 | 0.015 | 2266 | 0.273645976 | 0.04985135 | G/A |
| SNP_A.K_5_13684306 | 5 | 13684306 | 0.015 | 2266 | 0.273646559 | 0.04985135 | C/A |
| SNP_A.K_5_13595120 | 5 | 13595120 | 0.017 | 2266 | 0.273633701 | 0.049909494 | G/A |

HGW_gm

| SNP | Chr | Position | maf | nobs | $R^2$ with SNP | FDR_Adjusted_P-values | allele |
|---|---|---|---|---|---|---|---|
| SNP_A.K_5_8449552 | 5 | 8449552 | 0.017 | 2264 | 0.156400425 | 1.42347E-11 | C/T |
| SNP_A.K_5_8027721 | 5 | 8027721 | 0.022 | 2264 | 0.155013016 | 4.21312E-11 | G/A |
| SNP_A.K_5_8172473 | 5 | 8172473 | 0.015 | 2264 | 0.153236193 | 2.75564E-10 | G/A |
| SNP_A.K_5_8933914 | 5 | 8933914 | 0.019 | 2264 | 0.151603923 | 1.69466E-09 | G/A |
| SNP_A.K_5_8541630 | 5 | 8541630 | 0.027 | 2264 | 0.14760889 | 2.40446E-07 | G/A |
| SNP_A.K_5_9084127 | 5 | 9084127 | 0.019 | 2264 | 0.146619079 | 6.23812E-07 | G/A |
| SNP_A.K_5_9218977 | 5 | 9218977 | 0.018 | 2264 | 0.146724952 | 6.23812E-07 | T/A |
| SNP_A.K_5_9212444 | 5 | 9212444 | 0.019 | 2264 | 0.143297425 | 4.23776E-05 | C/T |
| SNP_A.K_5_9045676 | 5 | 9045676 | 0.020 | 2264 | 0.143106129 | 4.84588E-05 | G/A |
| SNP_A.K_5_11666786 | 5 | 11666786 | 0.023 | 2264 | 0.14282942 | 6.28002E-05 | G/A |
| SNP_A.K_5_12779064 | 5 | 12779064 | 0.018 | 2264 | 0.141951278 | 0.000181982 | C/T |
| SNP_A.K_5_9212909 | 5 | 9212909 | 0.017 | 2264 | 0.14134149 | 0.00037389 | G/A |
| SNP_A.K_5_9212097 | 5 | 9212097 | 0.052 | 2264 | 0.141088906 | 0.000482389 | G/T |
| SNP_A.K_5_9173352 | 5 | 9173352 | 0.021 | 2264 | 0.140880714 | 0.000516637 | T/A |
| SNP_A.K_5_9389374 | 5 | 9389374 | 0.017 | 2264 | 0.140901461 | 0.000516637 | G/C |
| SNP_A.K_5_13093127 | 5 | 13093127 | 0.175 | 2264 | 0.140886613 | 0.000516637 | C/T |
| SNP_A.K_5_12789820 | 5 | 12789820 | 0.014 | 2264 | 0.140690898 | 0.000625631 | A/T |
| SNP_A.K_5_10807497 | 5 | 10807497 | 0.482 | 2264 | 0.140102685 | 0.001288875 | G/A |
| SNP_A.K_5_13565167 | 5 | 13565167 | 0.015 | 2264 | 0.140063868 | 0.001288875 | G/A |
| SNP_A.K_5_12509109 | 5 | 12509109 | 0.120 | 2264 | 0.139937732 | 0.001418825 | T/A |
| SNP_A.K_5_12661806 | 5 | 12661806 | 0.015 | 2264 | 0.139916644 | 0.001418825 | C/T |
| SNP_A.K_5_9356298 | 5 | 9356298 | 0.013 | 2264 | 0.139767443 | 0.001580539 | A/T |
| SNP_A.K_5_12415856 | 5 | 12415856 | 0.156 | 2264 | 0.13978015 | 0.001580539 | C/T |
| SNP_A.K_5_8915413 | 5 | 8915413 | 0.025 | 2264 | 0.139624061 | 0.001833972 | C/A |
| SNP_A.K_5_9289192 | 5 | 9289192 | 0.014 | 2264 | 0.139534146 | 0.001838073 | C/T |
| SNP_A.K_5_9356299 | 5 | 9356299 | 0.013 | 2264 | 0.13953569 | 0.001838073 | A/T |
| SNP_A.K_5_10410809 | 5 | 10410809 | 0.013 | 2264 | 0.139550798 | 0.001838073 | C/A |
| SNP_A.K_5_9139104 | 5 | 9139104 | 0.046 | 2264 | 0.139455888 | 0.001899816 | G/A |
| SNP_A.K_5_11870967 | 5 | 11870967 | 0.023 | 2264 | 0.139466811 | 0.001899816 | C/T |
| SNP_A.K_5_10443983 | 5 | 10443983 | 0.029 | 2264 | 0.139386681 | 0.002014357 | G/A |

Days_to_first_flower

| SNP | Chr | Position | maf | nobs | $R^2$ with SNP | FDR_Adjusted_P-values | allele |
|---|---|---|---|---|---|---|---|
| SNP_A.K_5_11378339 | 5 | 11378339 | 0.024 | 743 | 0.441061305 | 0.036700749 | C/T |
| SNP_A.K_5_13002048 | 5 | 13002048 | 0.266 | 743 | 0.441055468 | 0.036700749 | G/A |
| SNP_A.K_5_12092645 | 5 | 12092645 | 0.023 | 743 | 0.440807702 | 0.036700749 | C/G |

Days_to_80_percent_Heading

| SNP | Chr | Position | maf | nobs | $R^2$ with SNP | FDR_Adjusted_P-values | allele |
|---|---|---|---|---|---|---|---|
| SNP_A.K_5_11063392 | 5 | 11063392 | 0.292 | 2270 | 0.451428834 | 0.001989403 | A/G |
| SNP_A.K_5_8894243 | 5 | 8894243 | 0.054 | 2270 | 0.451308017 | 0.001989403 | A/G |
| SNP_A.K_5_11690459 | 5 | 11690459 | 0.239 | 2270 | 0.451116514 | 0.001989403 | A/G |
| SNP_A.K_5_13002048 | 5 | 13002048 | 0.289 | 2270 | 0.450140574 | 0.011439548 | G/A |

Panicle_axis

| SNP | Chr | Position | maf | nobs | $R^2$ with SNP | FDR_Adjusted_P-values | allele |
|---|---|---|---|---|---|---|---|
| SNP_A.K_5_10728963 | 5 | 10728963 | 0.029 | 1527 | 0.024798739 | 0.000115725 | C/T |
| SNP_A.K_5_10867318 | 5 | 10867318 | 0.029 | 1527 | 0.024839405 | 0.000115725 | C/T |
| SNP_A.K_5_13332632 | 5 | 13332632 | 0.021 | 1527 | 0.02619532 | 0.000115725 | T/A |
| SNP_A.K_5_9659462 | 5 | 9659462 | 0.037 | 1527 | 0.023263545 | 0.000188552 | G/A |
| SNP_A.K_5_10841323 | 5 | 10841323 | 0.030 | 1527 | 0.023501733 | 0.000188552 | G/A |
| SNP_A.K_5_11661531 | 5 | 11661531 | 0.028 | 1527 | 0.023298007 | 0.000188552 | T/C |
| SNP_A.K_5_8449398 | 5 | 8449398 | 0.029 | 1527 | 0.021968467 | 0.000439201 | C/T |
| SNP_A.K_5_11360824 | 5 | 11360824 | 0.043 | 1527 | 0.021657992 | 0.000488606 | T/A |
| SNP_A.K_5_13466183 | 5 | 13466183 | 0.024 | 1527 | 0.019958276 | 0.001622538 | C/T |
| SNP_A.K_5_8359650 | 5 | 8359650 | 0.025 | 1527 | 0.01862565 | 0.003974221 | T/G |
| SNP_A.K_5_10620861 | 5 | 10620861 | 0.036 | 1527 | 0.018550176 | 0.003974221 | T/A |
| SNP_A.K_5_12241979 | 5 | 12241979 | 0.050 | 1527 | 0.018271645 | 0.004527889 | A/G |
| SNP_A.K_5_10284936 | 5 | 10284936 | 0.051 | 1527 | 0.01690316 | 0.010877265 | G/A |
| SNP_A.K_5_10574641 | 5 | 10574641 | 0.024 | 1527 | 0.016979088 | 0.010877265 | A/G |
| SNP_A.K_5_12705812 | 5 | 12705812 | 0.056 | 1527 | 0.016866945 | 0.010877265 | C/T |
| SNP_A.K_5_10396542 | 5 | 10396542 | 0.046 | 1527 | 0.016691407 | 0.011015307 | G/A |
| SNP_A.K_5_10812190 | 5 | 10812190 | 0.014 | 1527 | 0.016696201 | 0.011015307 | T/A |
| SNP_A.K_5_11353753 | 5 | 11353753 | 0.039 | 1527 | 0.016486612 | 0.01221877 | C/T |
| SNP_A.K_5_9303187 | 5 | 9303187 | 0.038 | 1527 | 0.016366901 | 0.012717542 | C/T |
| SNP_A.K_5_12318571 | 5 | 12318571 | 0.060 | 1527 | 0.016166896 | 0.014139301 | C/T |
| SNP_A.K_5_8114117 | 5 | 8114117 | 0.038 | 1527 | 0.015915059 | 0.01641772 | G/A |
| SNP_A.K_5_9969602 | 5 | 9969602 | 0.046 | 1527 | 0.015626658 | 0.016697959 | C/T |
| SNP_A.K_5_12241994 | 5 | 12241994 | 0.047 | 1527 | 0.015624478 | 0.016697959 | C/T |
| SNP_A.K_5_12242140 | 5 | 12242140 | 0.070 | 1527 | 0.015800181 | 0.016697959 | C/T |
| SNP_A.K_5_13123673 | 5 | 13123673 | 0.098 | 1527 | 0.015664017 | 0.016697959 | C/T |
| SNP_A.K_5_13901045 | 5 | 13901045 | 0.029 | 1527 | 0.015622477 | 0.016697959 | G/T |
| SNP_A.K_5_13906932 | 5 | 13906932 | 0.040 | 1527 | 0.015526002 | 0.017350276 | G/T |
| SNP_A.K_5_8158846 | 5 | 8158846 | 0.023 | 1527 | 0.015362832 | 0.019028784 | G/A |
| SNP_A.K_5_13617349 | 5 | 13617349 | 0.047 | 1527 | 0.015182081 | 0.021190268 | T/C |
| SNP_A.K_5_12248762 | 5 | 12248762 | 0.055 | 1527 | 0.014954593 | 0.024516954 | C/T |
| SNP_A.K_5_9582256 | 5 | 9582256 | 0.042 | 1527 | 0.0148652 | 0.025463508 | G/A |
| SNP_A.K_5_10398368 | 5 | 10398368 | 0.059 | 1527 | 0.014761481 | 0.025965202 | A/G |
| SNP_A.K_5_10488519 | 5 | 10488519 | 0.096 | 1527 | 0.01478473 | 0.025965202 | A/G |
| SNP_A.K_5_11013083 | 5 | 11013083 | 0.053 | 1527 | 0.014482573 | 0.030180196 | T/C |
| SNP_A.K_5_12067074 | 5 | 12067074 | 0.012 | 1527 | 0.014484273 | 0.030180196 | C/T |
| SNP_A.K_5_12921253 | 5 | 12921253 | 0.012 | 1527 | 0.014461529 | 0.030180196 | G/A |
| SNP_A.K_5_8138291 | 5 | 8138291 | 0.018 | 1527 | 0.014386627 | 0.030608037 | G/A |
| SNP_A.K_5_8380161 | 5 | 8380161 | 0.015 | 1527 | 0.014342741 | 0.030608037 | C/T |
| SNP_A.K_5_13117915 | 5 | 13117915 | 0.007 | 1527 | 0.014352335 | 0.030608037 | C/T |
| SNP_A.K_5_8848629 | 5 | 8848629 | 0.048 | 1527 | 0.014268792 | 0.030872611 | T/A |
| SNP_A.K_5_12301094 | 5 | 12301094 | 0.019 | 1527 | 0.014284218 | 0.030872611 | C/T |
| SNP_A.K_5_12279367 | 5 | 12279367 | 0.055 | 1527 | 0.014036455 | 0.036236632 | G/A |
| SNP_A.K_5_10894849 | 5 | 10894849 | 0.102 | 1527 | 0.014003507 | 0.036331745 | A/G |
| SNP_A.K_5_8549310 | 5 | 8549310 | 0.024 | 1527 | 0.013960996 | 0.036724719 | C/T |
| SNP_A.K_5_11214681 | 5 | 11214681 | 0.051 | 1527 | 0.013676315 | 0.045022183 | C/T |
| SNP_A.K_5_10722486 | 5 | 10722486 | 0.012 | 1527 | 0.013612409 | 0.046339384 | T/A |
| SNP_A.K_5_10330298 | 5 | 10330298 | 0.048 | 1527 | 0.013575224 | 0.046714855 | C/T |
| SNP_A.K_5_13426145 | 5 | 13426145 | 0.048 | 1527 | 0.013489193 | 0.048982154 | C/A |
| SNP_A.K_5_8180679 | 5 | 8180679 | 0.047 | 1527 | 0.013447224 | 0.049612306 | G/C |

Panicle_length_cm

| SNP | Chr | Position | maf | nobs | $R^2$ with SNP | FDR_Adjusted_P-values | allele |
|---|---|---|---|---|---|---|---|
| SNP_A.K_5_13702562 | 5 | 13702562 | 0.025 | 1897 | 0.241251019 | 4.29532E-06 | A/G |
| SNP_A.K_5_13886106 | 5 | 13886106 | 0.085 | 1897 | 0.236967455 | 0.00041182 | G/A |

| | | | | | | | |
|---|---|---|---|---|---|---|---|
| SNP_A.K_5_13810557 | 5 | 13810557 | 0.028 | 1897 | 0.23557961 | 0.001270231 | T/A |
| SNP_A.K_5_13893047 | 5 | 13893047 | 0.087 | 1897 | 0.23549756 | 0.001270231 | C/T |
| SNP_A.K_5_8005349 | 5 | 8005349 | 0.017 | 1897 | 0.234850356 | 0.001622258 | G/A |
| SNP_A.K_5_8730426 | 5 | 8730426 | 0.017 | 1897 | 0.235084287 | 0.001622258 | C/T |
| SNP_A.K_5_13697935 | 5 | 13697935 | 0.017 | 1897 | 0.234867941 | 0.001622258 | C/A |
| SNP_A.K_5_8558841 | 5 | 8558841 | 0.019 | 1897 | 0.234695749 | 0.001720665 | T/A |
| SNP_A.K_5_13972981 | 5 | 13972981 | 0.344 | 1897 | 0.234390288 | 0.002237726 | G/A |
| SNP_A.K_5_9385636 | 5 | 9385636 | 0.016 | 1897 | 0.234200114 | 0.002320887 | G/A |
| SNP_A.K_5_13778114 | 5 | 13778114 | 0.021 | 1897 | 0.234201117 | 0.002320887 | C/T |
| SNP_A.K_5_9475849 | 5 | 9475849 | 0.017 | 1897 | 0.234014109 | 0.00268341 | A/G |
| SNP_A.K_5_8236422 | 5 | 8236422 | 0.016 | 1897 | 0.233874847 | 0.002947567 | G/A |
| SNP_A.K_5_8063107 | 5 | 8063107 | 0.019 | 1897 | 0.233734194 | 0.00312677 | G/A |
| SNP_A.K_5_8158363 | 5 | 8158363 | 0.020 | 1897 | 0.233567354 | 0.00312677 | T/G |
| SNP_A.K_5_9015056 | 5 | 9015056 | 0.021 | 1897 | 0.233578862 | 0.00312677 | C/T |
| SNP_A.K_5_9521534 | 5 | 9521534 | 0.018 | 1897 | 0.23365442 | 0.00312677 | C/T |
| SNP_A.K_5_13699096 | 5 | 13699096 | 0.016 | 1897 | 0.233581758 | 0.00312677 | A/T |
| SNP_A.K_5_9933669 | 5 | 9933669 | 0.132 | 1897 | 0.233491262 | 0.003258124 | T/G |
| SNP_A.K_5_9933649 | 5 | 9933649 | 0.131 | 1897 | 0.233281673 | 0.00402413 | T/C |
| SNP_A.K_5_8532924 | 5 | 8532924 | 0.016 | 1897 | 0.23319849 | 0.004253554 | G/A |
| SNP_A.K_5_9052838 | 5 | 9052838 | 0.016 | 1897 | 0.23315828 | 0.004270105 | G/A |
| SNP_A.K_5_8539086 | 5 | 8539086 | 0.018 | 1897 | 0.233045567 | 0.004704521 | T/G |
| SNP_A.K_5_8576755 | 5 | 8576755 | 0.020 | 1897 | 0.232943171 | 0.00512656 | G/A |
| SNP_A.K_5_8064846 | 5 | 8064846 | 0.018 | 1897 | 0.232826767 | 0.005695863 | C/G |
| SNP_A.K_5_9314112 | 5 | 9314112 | 0.020 | 1897 | 0.232555405 | 0.00770231 | C/T |
| SNP_A.K_5_8108060 | 5 | 8108060 | 0.015 | 1897 | 0.232473811 | 0.008218697 | A/T |
| SNP_A.K_5_8464406 | 5 | 8464406 | 0.021 | 1897 | 0.232331144 | 0.009386949 | G/A |
| SNP_A.K_5_13689625 | 5 | 13689625 | 0.016 | 1897 | 0.232311443 | 0.009386949 | T/A |
| SNP_A.K_5_13770442 | 5 | 13770442 | 0.019 | 1897 | 0.231988108 | 0.013639297 | G/A |
| SNP_A.K_5_8463563 | 5 | 8463563 | 0.022 | 1897 | 0.231945877 | 0.01392173 | C/T |
| SNP_A.K_5_9014486 | 5 | 9014486 | 0.022 | 1897 | 0.231907514 | 0.014155725 | G/T |
| SNP_A.K_5_12435636 | 5 | 12435636 | 0.020 | 1897 | 0.231776492 | 0.016196441 | G/A |
| SNP_A.K_5_8345394 | 5 | 8345394 | 0.078 | 1897 | 0.231566464 | 0.019914341 | G/A |
| SNP_A.K_5_8461806 | 5 | 8461806 | 0.017 | 1897 | 0.231575777 | 0.019914341 | A/T |
| SNP_A.K_5_11353548 | 5 | 11353548 | 0.013 | 1897 | 0.231221674 | 0.029960809 | G/A |
| SNP_A.K_5_8564207 | 5 | 8564207 | 0.060 | 1897 | 0.231171959 | 0.030391184 | C/T |
| SNP_A.K_5_10490501 | 5 | 10490501 | 0.018 | 1897 | 0.231167788 | 0.030391184 | C/T |
| SNP_A.K_5_8393214 | 5 | 8393214 | 0.016 | 1897 | 0.231000085 | 0.036634644 | C/G |
| SNP_A.K_5_8391829 | 5 | 8391829 | 0.022 | 1897 | 0.230950233 | 0.038053345 | A/T |
| SNP_A.K_5_9350565 | 5 | 9350565 | 0.298 | 1897 | 0.230902428 | 0.039449845 | G/T |
| SNP_A.K_5_12099863 | 5 | 12099863 | 0.018 | 1897 | 0.230874133 | 0.039920495 | T/A |

Panicle_shattering

| | | | | | | | |
|---|---|---|---|---|---|---|---|
| SNP_A.K_5_8507521 | 5 | 8507521 | 0.023 | 1526 | 0.023998006 | 0.012629134 | G/T |

Seed_coat_color

| SNP | Chr | Position | maf | nobs | $R^2$ with SNP | FDR_Adjusted_P-values | allele |
|---|---|---|---|---|---|---|---|
| SNP_A.K_5_8322836 | 5 | 8322836 | 0.015 | 2110 | 0.042214027 | 0.000929448 | A/C |
| SNP_A.K_5_12334319 | 5 | 12334319 | 0.016 | 2110 | 0.04203224 | 0.000929448 | C/T |
| SNP_A.K_5_8352246 | 5 | 8352246 | 0.016 | 2110 | 0.040839078 | 0.002310631 | G/A |
| SNP_A.K_5_8362611 | 5 | 8362611 | 0.017 | 2110 | 0.039905924 | 0.004868431 | G/T |
| SNP_A.K_5_8224456 | 5 | 8224456 | 0.015 | 2110 | 0.038820635 | 0.013005993 | G/A |
| SNP_A.K_5_8051440 | 5 | 8051440 | 0.018 | 2110 | 0.038017496 | 0.014377455 | G/A |
| SNP_A.K_5_8097341 | 5 | 8097341 | 0.018 | 2110 | 0.038264013 | 0.014377455 | C/A |
| SNP_A.K_5_8132974 | 5 | 8132974 | 0.040 | 2110 | 0.038478615 | 0.014377455 | A/T |
| SNP_A.K_5_8300172 | 5 | 8300172 | 0.025 | 2110 | 0.03786913 | 0.014377455 | C/G |
| SNP_A.K_5_8327062 | 5 | 8327062 | 0.025 | 2110 | 0.038182543 | 0.014377455 | T/A |
| SNP_A.K_5_13317000 | 5 | 13317000 | 0.018 | 2110 | 0.038395933 | 0.014377455 | G/A |
| SNP_A.K_5_13992770 | 5 | 13992770 | 0.098 | 2110 | 0.037808062 | 0.014377455 | A/T |
| SNP_A.K_5_13993609 | 5 | 13993609 | 0.097 | 2110 | 0.038054663 | 0.014377455 | C/G |
| SNP_A.K_5_13994902 | 5 | 13994902 | 0.100 | 2110 | 0.037921771 | 0.014377455 | G/A |
| SNP_A.K_5_8328585 | 5 | 8328585 | 0.030 | 2110 | 0.03768306 | 0.014468497 | C/T |
| SNP_A.K_5_13994594 | 5 | 13994594 | 0.101 | 2110 | 0.037707077 | 0.014468497 | A/T |
| SNP_A.K_5_8320771 | 5 | 8320771 | 0.012 | 2110 | 0.037265349 | 0.016070571 | C/T |
| SNP_A.K_5_8320900 | 5 | 8320900 | 0.012 | 2110 | 0.037339026 | 0.016070571 | C/T |

| | | | | | | | |
|---|---|---|---|---|---|---|---|
| SNP_A.K_5_12356652 | 5 | 12356652 | 0.022 | 2110 | 0.037405238 | 0.016070571 | C/T |
| SNP_A.K_5_12509109 | 5 | 12509109 | 0.120 | 2110 | 0.03740774 | 0.016070571 | T/A |
| SNP_A.K_5_13989752 | 5 | 13989752 | 0.098 | 2110 | 0.037308924 | 0.016070571 | C/A |
| SNP_A.K_5_13994635 | 5 | 13994635 | 0.102 | 2110 | 0.037381771 | 0.016070571 | A/C |
| SNP_A.K_5_13998802 | 5 | 13998802 | 0.098 | 2110 | 0.037293751 | 0.016070571 | T/C |
| SNP_A.K_5_8212050 | 5 | 8212050 | 0.043 | 2110 | 0.037017744 | 0.016345663 | G/A |
| SNP_A.K_5_9541458 | 5 | 9541458 | 0.091 | 2110 | 0.037036434 | 0.016345663 | C/T |
| SNP_A.K_5_12175257 | 5 | 12175257 | 0.105 | 2110 | 0.037013435 | 0.016345663 | T/C |
| SNP_A.K_5_12175974 | 5 | 12175974 | 0.109 | 2110 | 0.037111972 | 0.016345663 | T/C |
| SNP_A.K_5_13858426 | 5 | 13858426 | 0.083 | 2110 | 0.037031148 | 0.016345663 | G/T |
| SNP_A.K_5_13994230 | 5 | 13994230 | 0.099 | 2110 | 0.037204667 | 0.016345663 | G/T |
| SNP_A.K_5_13998522 | 5 | 13998522 | 0.097 | 2110 | 0.037068406 | 0.016345663 | A/G |
| SNP_A.K_5_8080053 | 5 | 8080053 | 0.034 | 2110 | 0.036956971 | 0.016515048 | G/A |
| SNP_A.K_5_13995154 | 5 | 13995154 | 0.101 | 2110 | 0.036946788 | 0.016515048 | C/T |
| SNP_A.K_5_12075722 | 5 | 12075722 | 0.109 | 2110 | 0.036807993 | 0.017946127 | C/T |
| SNP_A.K_5_13988408 | 5 | 13988408 | 0.096 | 2110 | 0.036808612 | 0.017946127 | G/A |
| SNP_A.K_5_13989319 | 5 | 13989319 | 0.099 | 2110 | 0.03679309 | 0.017946127 | G/A |
| SNP_A.K_5_12176149 | 5 | 12176149 | 0.111 | 2110 | 0.036767354 | 0.017959829 | A/G |
| SNP_A.K_5_13988556 | 5 | 13988556 | 0.089 | 2110 | 0.036731303 | 0.018197364 | C/T |
| SNP_A.K_5_8107030 | 5 | 8107030 | 0.023 | 2110 | 0.036529128 | 0.022244616 | T/G |
| SNP_A.K_5_9819377 | 5 | 9819377 | 0.120 | 2110 | 0.036439719 | 0.023970256 | T/A |
| SNP_A.K_5_8104424 | 5 | 8104424 | 0.029 | 2110 | 0.036365748 | 0.025226426 | G/T |
| SNP_A.K_5_8894823 | 5 | 8894823 | 0.112 | 2110 | 0.036343845 | 0.025226426 | C/T |
| SNP_A.K_5_13995589 | 5 | 13995589 | 0.103 | 2110 | 0.036328622 | 0.025226426 | A/T |
| SNP_A.K_5_12175994 | 5 | 12175994 | 0.114 | 2110 | 0.036267843 | 0.026387315 | A/G |
| SNP_A.K_5_13996896 | 5 | 13996896 | 0.097 | 2110 | 0.036172895 | 0.028227221 | C/T |
| SNP_A.K_5_13998803 | 5 | 13998803 | 0.116 | 2110 | 0.03616778 | 0.028227221 | G/T |
| SNP_A.K_5_13974845 | 5 | 13974845 | 0.120 | 2110 | 0.036147289 | 0.028259474 | G/A |
| SNP_A.K_5_9585535 | 5 | 9585535 | 0.088 | 2110 | 0.036059684 | 0.030532925 | C/T |
| SNP_A.K_5_10741429 | 5 | 10741429 | 0.059 | 2110 | 0.036023896 | 0.031129719 | G/A |
| SNP_A.K_5_13166749 | 5 | 13166749 | 0.089 | 2110 | 0.035921191 | 0.034245871 | C/A |
| SNP_A.K_5_8323766 | 5 | 8323766 | 0.013 | 2110 | 0.035854256 | 0.035531452 | G/A |
| SNP_A.K_5_8512705 | 5 | 8512705 | 0.148 | 2110 | 0.035849893 | 0.035531452 | G/A |
| SNP_A.K_5_12262414 | 5 | 12262414 | 0.181 | 2110 | 0.035836018 | 0.035531452 | A/G |
| SNP_A.K_5_13993814 | 5 | 13993814 | 0.091 | 2110 | 0.035724726 | 0.039537474 | C/A |
| SNP_A.K_5_8396638 | 5 | 8396638 | 0.095 | 2110 | 0.035673047 | 0.040714651 | C/T |
| SNP_A.K_5_13588026 | 5 | 13588026 | 0.024 | 2110 | 0.035666071 | 0.040714651 | C/T |
| SNP_A.K_5_8149346 | 5 | 8149346 | 0.021 | 2110 | 0.035642682 | 0.041060486 | G/A |
| SNP_A.K_5_11773732 | 5 | 11773732 | 0.088 | 2110 | 0.035599064 | 0.04238246 | C/T |
| SNP_A.K_5_9637309 | 5 | 9637309 | 0.154 | 2110 | 0.035527381 | 0.045174838 | T/A |
| SNP_A.K_5_8358988 | 5 | 8358988 | 0.037 | 2110 | 0.03550429 | 0.045586412 | C/T |
| SNP_A.K_5_8296512 | 5 | 8296512 | 0.023 | 2110 | 0.035461278 | 0.047065997 | C/T |

Culm_length_cm

| SNP | Chr | Position | maf | nobs | $R^2$ with SNP | FDR_Adjusted_P-values | allele |
|---|---|---|---|---|---|---|---|
| SNP_A.K_5_8114117 | 5 | 8114117 | 0.032 | 1901 | 0.305396463 | 2.06592E-06 | G/A |
| SNP_A.K_5_9303187 | 5 | 9303187 | 0.031 | 1901 | 0.305357006 | 2.06592E-06 | C/T |
| SNP_A.K_5_11764205 | 5 | 11764205 | 0.044 | 1901 | 0.305607069 | 2.06592E-06 | C/T |
| SNP_A.K_5_10398368 | 5 | 10398368 | 0.051 | 1901 | 0.303989555 | 9.67813E-06 | A/G |
| SNP_A.K_5_10620861 | 5 | 10620861 | 0.030 | 1901 | 0.303606217 | 1.29563E-05 | T/A |
| SNP_A.K_5_8950730 | 5 | 8950730 | 0.036 | 1901 | 0.302941615 | 2.27015E-05 | G/A |
| SNP_A.K_5_10174842 | 5 | 10174842 | 0.138 | 1901 | 0.302939192 | 2.27015E-05 | C/T |
| SNP_A.K_5_10330298 | 5 | 10330298 | 0.040 | 1901 | 0.302708821 | 2.70923E-05 | C/T |
| SNP_A.K_5_9809816 | 5 | 9809816 | 0.046 | 1901 | 0.302361092 | 3.84884E-05 | G/C |
| SNP_A.K_5_13886974 | 5 | 13886974 | 0.013 | 1901 | 0.302096744 | 4.94925E-05 | A/G |
| SNP_A.K_5_10284936 | 5 | 10284936 | 0.046 | 1901 | 0.301671935 | 7.98868E-05 | G/A |
| SNP_A.K_5_9969602 | 5 | 9969602 | 0.040 | 1901 | 0.300640615 | 0.000296208 | C/T |
| SNP_A.K_5_12248762 | 5 | 12248762 | 0.052 | 1901 | 0.300510764 | 0.000304547 | C/T |
| SNP_A.K_5_12334304 | 5 | 12334304 | 0.014 | 1901 | 0.300506664 | 0.000304547 | G/A |
| SNP_A.K_5_10396542 | 5 | 10396542 | 0.039 | 1901 | 0.300313365 | 0.000369636 | G/A |
| SNP_A.K_5_9213498 | 5 | 9213498 | 0.038 | 1901 | 0.300228847 | 0.000388737 | C/T |
| SNP_A.K_5_9887574 | 5 | 9887574 | 0.124 | 1901 | 0.300071962 | 0.000452917 | C/T |
| SNP_A.K_5_9165947 | 5 | 9165947 | 0.013 | 1901 | 0.299730144 | 0.000681339 | A/C |
| SNP_A.K_5_9870500 | 5 | 9870500 | 0.023 | 1901 | 0.299348689 | 0.001086065 | A/T |
| SNP_A.K_5_11353753 | 5 | 11353753 | 0.033 | 1901 | 0.299252915 | 0.001175921 | C/T |

| | | | | | | | |
|---|---|---|---|---|---|---|---|
| SNP_A.K_5_13002048 | 5 | 13002048 | 0.301 | 1901 | 0.299011283 | 0.001558124 | G/A |
| SNP_A.K_5_12631510 | 5 | 12631510 | 0.011 | 1901 | 0.298594029 | 0.002632975 | G/A |
| SNP_A.K_5_8892384 | 5 | 8892384 | 0.052 | 1901 | 0.298509264 | 0.002701267 | A/G |
| SNP_A.K_5_10205527 | 5 | 10205527 | 0.092 | 1901 | 0.298517509 | 0.002701267 | A/C |
| SNP_A.K_5_13123673 | 5 | 13123673 | 0.110 | 1901 | 0.298482085 | 0.002701267 | C/T |
| SNP_A.K_5_11267126 | 5 | 11267126 | 0.013 | 1901 | 0.298178229 | 0.003941046 | C/T |
| SNP_A.K_5_8892372 | 5 | 8892372 | 0.052 | 1901 | 0.297967226 | 0.004890428 | C/T |
| SNP_A.K_5_13549188 | 5 | 13549188 | 0.009 | 1901 | 0.297971502 | 0.004890428 | C/T |
| SNP_A.K_5_11655029 | 5 | 11655029 | 0.017 | 1901 | 0.297918255 | 0.00505072 | C/A |
| SNP_A.K_5_8350184 | 5 | 8350184 | 0.086 | 1901 | 0.29778369 | 0.005481002 | G/A |
| SNP_A.K_5_9920736 | 5 | 9920736 | 0.125 | 1901 | 0.29774324 | 0.005481002 | G/A |
| SNP_A.K_5_11160794 | 5 | 11160794 | 0.076 | 1901 | 0.297783672 | 0.005481002 | C/T |
| SNP_A.K_5_11353548 | 5 | 11353548 | 0.013 | 1901 | 0.297744377 | 0.005481002 | G/A |
| SNP_A.K_5_12875278 | 5 | 12875278 | 0.011 | 1901 | 0.297760182 | 0.005481002 | G/A |
| SNP_A.K_5_9707642 | 5 | 9707642 | 0.010 | 1901 | 0.297699397 | 0.00565572 | G/C |
| SNP_A.K_5_9817650 | 5 | 9817650 | 0.022 | 1901 | 0.29763988 | 0.00596835 | C/T |
| SNP_A.K_5_8547189 | 5 | 8547189 | 0.093 | 1901 | 0.297393712 | 0.00815341 | C/A |
| SNP_A.K_5_8894243 | 5 | 8894243 | 0.054 | 1901 | 0.297270991 | 0.009404092 | A/G |
| SNP_A.K_5_8126750 | 5 | 8126750 | 0.229 | 1901 | 0.297211132 | 0.009952606 | C/T |
| SNP_A.K_5_12292313 | 5 | 12292313 | 0.016 | 1901 | 0.297170705 | 0.010261136 | C/T |
| SNP_A.K_5_8180679 | 5 | 8180679 | 0.044 | 1901 | 0.297083158 | 0.010532103 | G/C |
| SNP_A.K_5_8386731 | 5 | 8386731 | 0.010 | 1901 | 0.297074043 | 0.010532103 | C/A |
| SNP_A.K_5_10261680 | 5 | 10261680 | 0.017 | 1901 | 0.297076991 | 0.010532103 | C/T |
| SNP_A.K_5_12681700 | 5 | 12681700 | 0.009 | 1901 | 0.297120332 | 0.010532103 | T/C |
| SNP_A.K_5_13505084 | 5 | 13505084 | 0.011 | 1901 | 0.297066625 | 0.010532103 | T/A |
| SNP_A.K_5_11064424 | 5 | 11064424 | 0.009 | 1901 | 0.296994874 | 0.011377901 | G/A |
| SNP_A.K_5_9810456 | 5 | 9810456 | 0.229 | 1901 | 0.296902763 | 0.012385959 | T/C |
| SNP_A.K_5_13835724 | 5 | 13835724 | 0.010 | 1901 | 0.296907394 | 0.012385959 | G/C |
| SNP_A.K_5_13886106 | 5 | 13886106 | 0.086 | 1901 | 0.296798738 | 0.014012883 | G/A |
| SNP_A.K_5_11743015 | 5 | 11743015 | 0.018 | 1901 | 0.296781484 | 0.014064769 | G/A |
| SNP_A.K_5_10365722 | 5 | 10365722 | 0.019 | 1901 | 0.296747754 | 0.014448667 | C/T |
| SNP_A.K_5_13421498 | 5 | 13421498 | 0.017 | 1901 | 0.296715566 | 0.014817187 | C/T |
| SNP_A.K_5_12391883 | 5 | 12391883 | 0.018 | 1901 | 0.296643423 | 0.01606671 | C/T |
| SNP_A.K_5_10333786 | 5 | 10333786 | 0.020 | 1901 | 0.296596969 | 0.016818552 | C/T |
| SNP_A.K_5_11662961 | 5 | 11662961 | 0.010 | 1901 | 0.296580354 | 0.016897768 | T/C |
| SNP_A.K_5_9185677 | 5 | 9185677 | 0.018 | 1901 | 0.296508592 | 0.018036706 | C/T |
| SNP_A.K_5_11431442 | 5 | 11431442 | 0.012 | 1901 | 0.296507608 | 0.018036706 | C/T |
| SNP_A.K_5_11184322 | 5 | 11184322 | 0.010 | 1901 | 0.296470078 | 0.0186737 | G/A |
| SNP_A.K_5_13802799 | 5 | 13802799 | 0.017 | 1901 | 0.296416422 | 0.019777262 | C/T |
| SNP_A.K_5_12316757 | 5 | 12316757 | 0.017 | 1901 | 0.296362967 | 0.020946854 | G/A |
| SNP_A.K_5_11063392 | 5 | 11063392 | 0.297 | 1901 | 0.296330289 | 0.021212947 | A/G |
| SNP_A.K_5_11073795 | 5 | 11073795 | 0.012 | 1901 | 0.296341958 | 0.021212947 | C/T |
| SNP_A.K_5_9658912 | 5 | 9658912 | 0.018 | 1901 | 0.296284897 | 0.022235984 | G/A |
| SNP_A.K_5_8875108 | 5 | 8875108 | 0.017 | 1901 | 0.296228161 | 0.023321146 | C/T |
| SNP_A.K_5_10120085 | 5 | 10120085 | 0.226 | 1901 | 0.296236038 | 0.023321146 | C/T |
| SNP_A.K_5_12557039 | 5 | 12557039 | 0.013 | 1901 | 0.296210062 | 0.023553427 | C/A |
| SNP_A.K_5_13466183 | 5 | 13466183 | 0.021 | 1901 | 0.296179024 | 0.024225789 | C/T |
| SNP_A.K_5_8279845 | 5 | 8279845 | 0.011 | 1901 | 0.296107697 | 0.026361009 | G/C |
| SNP_A.K_5_9447947 | 5 | 9447947 | 0.011 | 1901 | 0.296038956 | 0.02787359 | A/G |
| SNP_A.K_5_9634028 | 5 | 9634028 | 0.122 | 1901 | 0.296043746 | 0.02787359 | C/T |
| SNP_A.K_5_10432968 | 5 | 10432968 | 0.058 | 1901 | 0.296026599 | 0.02787359 | C/T |
| SNP_A.K_5_10644859 | 5 | 10644859 | 0.008 | 1901 | 0.296034904 | 0.02787359 | G/T |
| SNP_A.K_5_8357605 | 5 | 8357605 | 0.011 | 1901 | 0.296003641 | 0.028385457 | G/C |
| SNP_A.K_5_9891368 | 5 | 9891368 | 0.084 | 1901 | 0.295950426 | 0.030157817 | C/T |
| SNP_A.K_5_10193142 | 5 | 10193142 | 0.102 | 1901 | 0.29588063 | 0.032797616 | C/A |
| SNP_A.K_5_11351209 | 5 | 11351209 | 0.015 | 1901 | 0.295841774 | 0.034169135 | C/T |
| SNP_A.K_5_8430912 | 5 | 8430912 | 0.017 | 1901 | 0.295812744 | 0.035119823 | C/T |
| SNP_A.K_5_11024872 | 5 | 11024872 | 0.022 | 1901 | 0.295786756 | 0.035950383 | A/T |
| SNP_A.K_5_12659358 | 5 | 12659358 | 0.247 | 1901 | 0.29577534 | 0.036065634 | T/C |
| SNP_A.K_5_8401168 | 5 | 8401168 | 0.014 | 1901 | 0.295711585 | 0.038931519 | C/T |
| SNP_A.K_5_8891367 | 5 | 8891367 | 0.055 | 1901 | 0.295661209 | 0.040610173 | C/T |
| SNP_A.K_5_11423163 | 5 | 11423163 | 0.012 | 1901 | 0.295655031 | 0.040610173 | G/C |
| SNP_A.K_5_12818958 | 5 | 12818958 | 0.212 | 1901 | 0.295663728 | 0.040610173 | T/C |
| SNP_A.K_5_9268631 | 5 | 9268631 | 0.014 | 1901 | 0.295613206 | 0.042542959 | G/A |
| SNP_A.K_5_12170302 | 5 | 12170302 | 0.020 | 1901 | 0.29556615 | 0.044902386 | G/A |
| SNP_A.K_5_13700894 | 5 | 13700894 | 0.011 | 1901 | 0.295554828 | 0.045088813 | G/T |
| SNP_A.K_5_11948092 | 5 | 11948092 | 0.018 | 1901 | 0.295529606 | 0.045781974 | G/A |

| | | | | | | | |
|---|---|---|---|---|---|---|---|
| SNP_A.K_5_12778324 | 5 | 12778324 | 0.009 | 1901 | 0.295527494 | 0.045781974 | G/A |
| SNP_A.K_5_8057492 | 5 | 8057492 | 0.450 | 1901 | 0.295513654 | 0.046152921 | T/C |
| SNP_A.K_5_11004608 | 5 | 11004608 | 0.219 | 1901 | 0.29549819 | 0.046638703 | A/G |
| SNP_A.K_5_10971057 | 5 | 10971057 | 0.015 | 1901 | 0.295455625 | 0.048958543 | G/A |
| SNP_A.K_5_11180536 | 5 | 11180536 | 0.018 | 1901 | 0.295439369 | 0.04954149 | T/C |
| SNP_A.K_5_11756975 | 5 | 11756975 | 0.016 | 1901 | 0.295425995 | 0.049935474 | G/A |
